# Panvascular-JEPA: Unseen-Structure Evaluation of Factorized Predictive States for Hemodynamic Networks

**DOI:** 10.64898/2026.10.12.758273

**Authors:** Lingsen You, Zijie Feng, Chang Gao, Yingcheng Wu, Zhenfei Yin, Ling Yang, Li Shen, Junbo Ge

## Abstract

Predictive representations for vascular simulation should be tested on unseen network structures and compared with the governing equations before being interpreted as general-purpose world models. We constructed a new, dimensioned but uncalibrated nodal resistance–inertance–compliance benchmark containing all 38 connected, non-isomorphic six-node graphs with five to seven edges. Twenty-four structures supplied training, validation, and new-background testing; 14 structures were wholly withheld. Across 648 parameter backgrounds and two prescribed action programs, 1,296 trajectories contained 96 forecast steps over three cycles. Four graph predictors received equal six-candidate optimization searches, followed by five fixed-seed fits. Selected configurations and analysis code were locked before test trajectories were generated. Comparators included known-equation numerical integrators, an inertia-free reduction, and a five-coefficient model fitted to training trajectories.

Unseen-structure mean standardized RMSE was 8.46 × 10^9^ for four-factor OPF, 329 for single-factor OPF, and 1.83 × 10^31^ for supervised graph prediction, compared with 1.09 × 10^−4^ for four-substep known-equation integration and 6.77 × 10^−4^ for the fitted mechanism. The primary OPF-4 minus OPF-1 difference was 8.46 × 10^9^, with a paired stratified-graph 95% percentile interval of [2.23 × 10^5^, 2.54 × 10^10^]. Neural errors were strongly dependent on training seed and network structure, with extreme forecasts dominating the means. All four neural means exceeded persistence; relative performance within this comparison did not establish a usable forecasting model.

The study supplies a distinct numerical test of structure transfer, factor-count control, and recursive stability. Its conclusions concern the implemented models, optimization budget, and finite graph catalogue. It establishes neither patient-specific performance nor anatomical transfer, biological remodeling, device efficacy, or clinical readiness.

## 1 Introduction

Reduced-order cardiovascular simulation makes pressure and flow predictions tractable while exposing assumptions about topology, boundary conditions, and constitutive behavior [1, 2]. Learned graph surrogates already predict cardiovascular states recursively across different anatomies and boundary conditions [3]; neural, Gaussian-process, and polynomial metamodels have also been compared for zero-dimensional cardiovascular models [4]. Thus, learning hemodynamics on a graph is an established direction. A useful new experiment must identify what its representation adds beyond supervised prediction and available physical structure.

Joint-Embedding Predictive Architectures learn to predict encoded targets rather than reconstruct all observations during representation training [5]. JEPA-Anything introduces orthogonal predictive factorization (OPF), which assigns predictors to complementary learned target coordinates and synthesizes their outputs [6]. Variance regularization can discourage degenerate embeddings [7], but algebraic rank, active coordinates, and small one-step losses do not establish stable autonomous forecasts. Repeated prediction supplies the model with its own outputs, which can differ from encoded observations encountered during training.

Our earlier exploratory Panvascular-JEPA pilot used one resistor bifurcation and two RCR outlets. Its four-factor implementation did not outperform a regularized single predictor. That experiment could not assess transfer to a new graph and did not include a sufficiently explicit known-equation comparison. Here we ask a narrower, independently testable question: after equal-budget tuning, does four-factor prediction improve 96-step forecasts over a single-factor model on completely withheld network structures? We separate this primary contrast from comparisons with an identity-basis monolithic JEPA, a directly supervised graph predictor, and numerical mechanisms. All planned runs, including very large or nonfinite forecast errors, are retained.

The new study introduces a fresh graph catalogue, parameter draws, repeated controls, simulation outputs, and protected structure-level tests. It does not reuse the GEO expression data or predictive results of the authors’ related transcriptomic manuscripts. The word “panvascular” denotes the surrounding research programme; the present networks are abstract circuits rather than named arterial beds. The authors’ *Biomarker Research* IABP study [8] provides clinical context for distinguishing intervention exposures and endpoints. It supplies neither data nor validation for this numerical experiment.

## 2 Methods

### 2.1 Locally locked design and independent units

The design, simulation code, neural implementations, and development dataset were hashed before the first optimization search. That 1,000-update search showed large recursive validation errors across model classes and did not demonstrate adequate convergence. Before generating any test trajectories, we recorded a development-driven version-2 amendment: every arm’s full six-candidate grid was rerun from initialization for 6,000 updates, with five final seeds trained for 12,000 updates. All original outcomes were retained as budget-development evidence. Architecture, objective, data, hyperparameter grid, seeds, primary contrast, and test definitions were unchanged. The stopping rule prohibited repeatedly extending training until a preferred outcome appeared. A separate final selection lock recorded the selected hyperparameters, evaluation code, and analysis script before test generation. This locally recorded sequence is not external preregistration, and the revision was informed by development failures. Candidate selection uses validation data only, addressing a recognized source of performance-selection bias [9].

NetworkX 3.6.1 [10] supplied all connected simple undirected six-node graphs with five, six, or seven edges: 6, 13, and 19 structures, respectively. Pairwise graph-isomorphism checks verified that all 38 were distinct. A fixed random ordering within edge-count strata allocated 4/8/12 structures to the seen group and 2/5/7 to the withheld group. The split unit is the *unrooted isomorphism class*; changes in parameter values, vertex labels, edge order, or action program cannot move an instance of the same structure into the withheld group. All graphs have the same number of nodes. This tests unseen structures within a finite catalogue, without establishing size extrapolation or a physiological distribution shift.

Each of the 24 seen structures contributed 12 training, four validation, and four test backgrounds. Each of the 14 withheld structures contributed 12 test backgrounds. Each background generated two action programs with identical initial states and parameters. There were therefore 288/96/96/168 backgrounds and 576/192/192/336 episodes in training, validation, seen-structure testing, and unseen-structure testing, respectively. Node labels other than the inlet and edge order were randomly permuted for each background, consistently across all associated arrays. Graph identity was never a model input. Figure 1 summarizes the design and Figure 2 shows the complete structure split.

**Figure 1:**
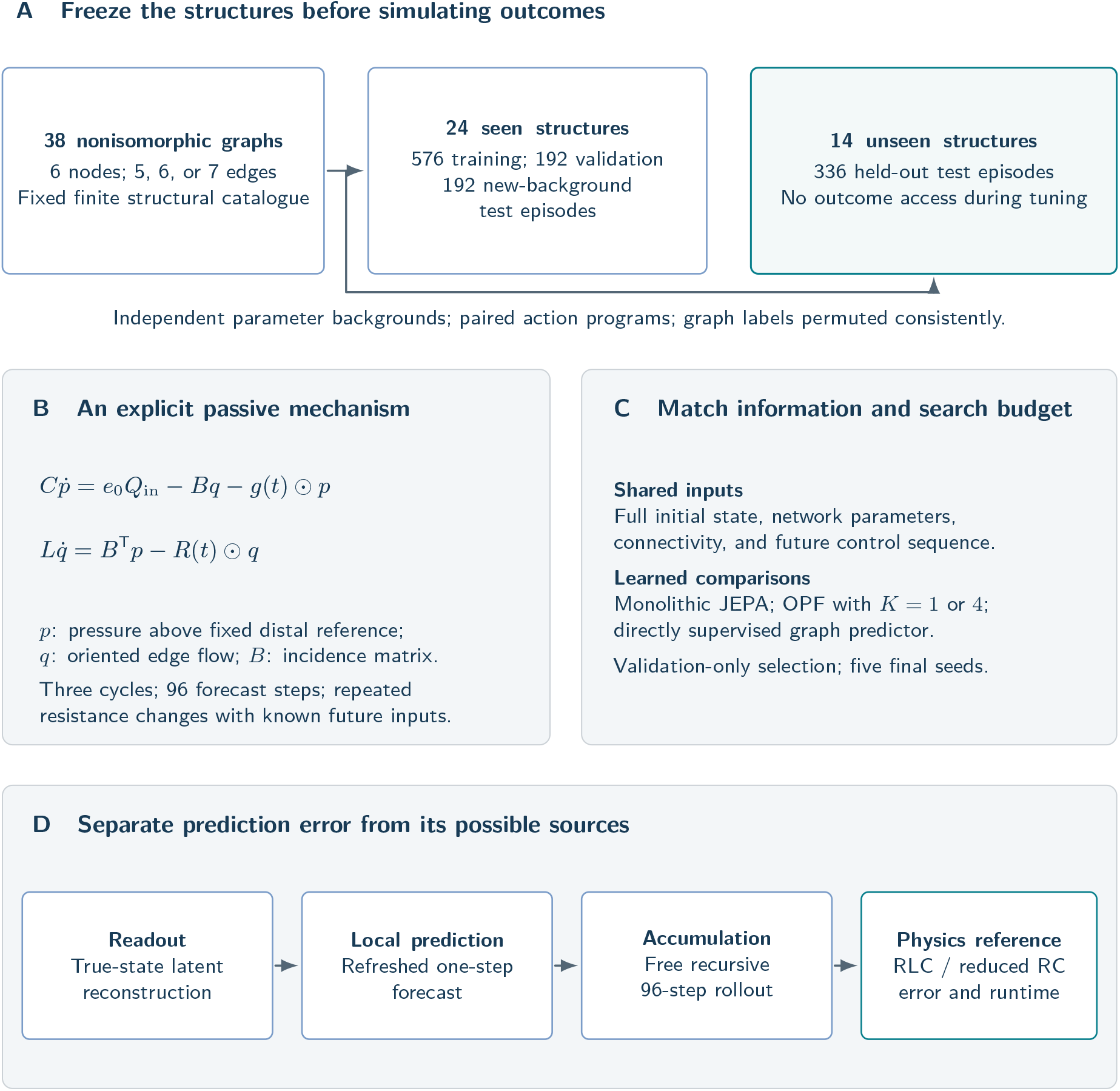
Design of the independent graph-structure experiment. Every episode is newly simulated; parameter backgrounds and action programs are separated from structural split units. Nodes carry compliance and edges carry resistance and inertance in a custom nodal RLC network. The pressure reference is 5 mmHg, and stored pressures are relative to that reference. Known-parameter mechanistic predictors receive the same information supplied to the learned models. The diagram describes the fixed experimental design, not an observed performance ranking or a patient-specific anatomy.

**Figure 2:**
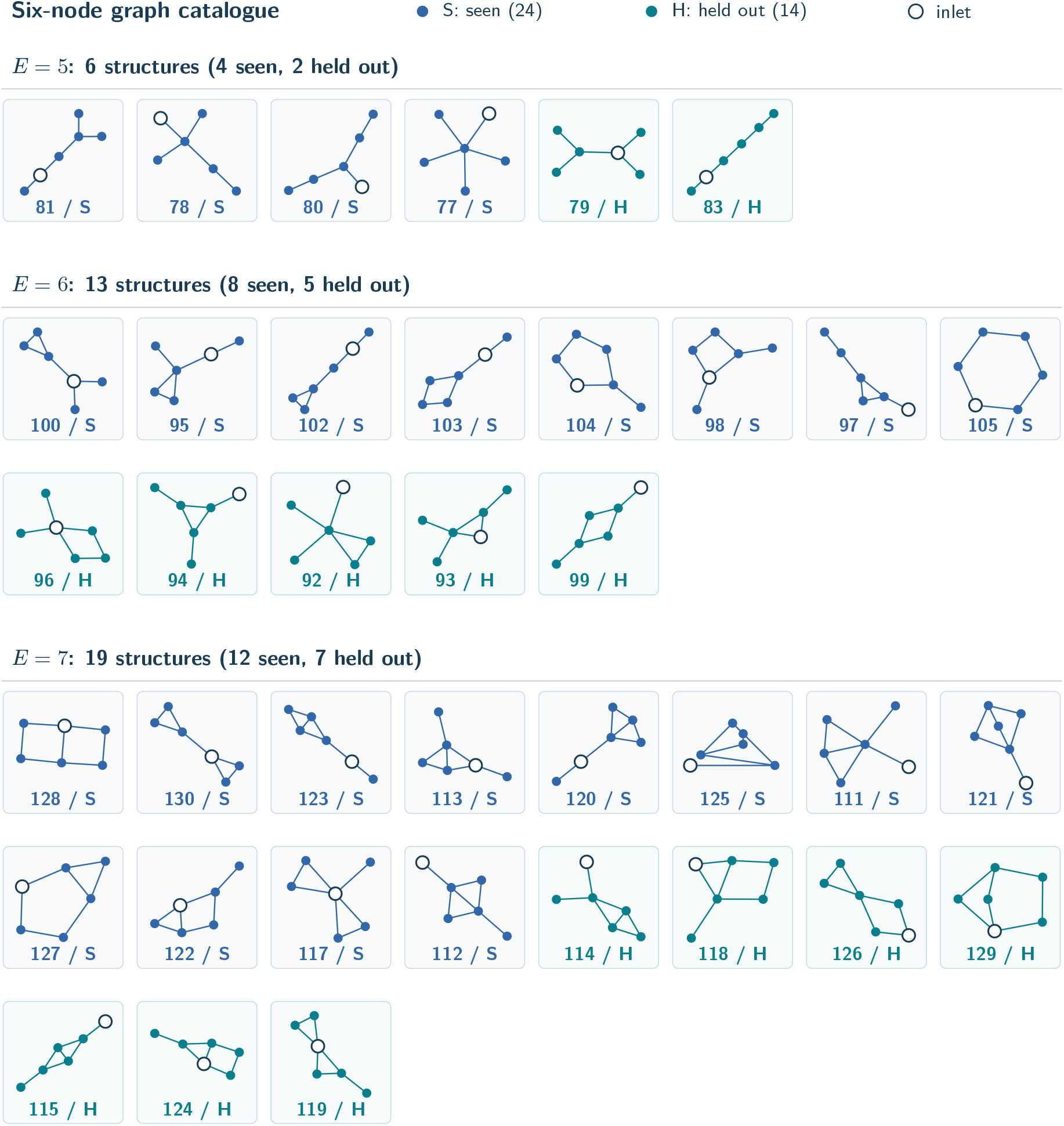
The complete fixed catalogue of 38 connected, pairwise nonisomorphic, six-node graphs with five, six, or seven edges. Colours and letter tags identify the 24 seen structures (S) and 14 held-out structures (H); numbers are NetworkX graph-atlas indices. An outlined node marks the inlet. Edge orientation is only a sign convention and is omitted here. The held-out structures are not used in model fitting or hyperparameter selection. These are abstract circuit topologies, not reconstructed vascular anatomies.

### 2.2 Numerical model and prescribed perturbations

This study implements a custom passive nodal RLC network, not a direct reproduction of a svZeroDSolver blood-vessel block. The software family and its reduced-order formulations provide methodological context [1, 2]; the executable custom equations are given here. Each node has positive compliance *C*_*i*_, and each edge has positive resistance *R*_*e*_ and inertance *L*_*e*_. With incidence matrix *B* positive at an edge’s source and negative at its destination,

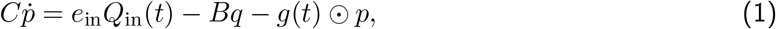

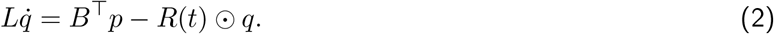

Here *C* and *L* are diagonal matrices, *e*_in_ marks the inlet, and *g* is nonzero at two distal runoff nodes. Pressures are stored as gauge pressures relative to a fixed distal pressure of 5 mmHg; adding 5 restores pressure on the specified reference scale, rather than thermodynamic absolute pressure. Edge direction is a bookkeeping convention, so negative edge flow does not itself indicate failed simulation. Two non-inlet nodes with the largest shortest-path distance from the inlet are selected as runoff sites, with deterministic tie-breaking. This rule is structural, not an anatomical assertion.

Mean inflow is sampled uniformly over 60–100 mL/s, period over 0.75–1.05 s, total compliance over 0.8–1.5 mL/mmHg, edge resistance over 0.03–0.10 mmHg s/mL, and edge inertance over 0.002– 0.010 mmHg s^2^/mL. Nodal compliance weights are sampled over 0.75–1.25 and rescaled to the sampled total. Each terminal resistance is twice a shared runoff scale of 0.7–1.2 mmHg s/mL, multiplied by independent 0.85–1.15 jitter. Inflow includes two sinusoidal harmonics with fractions sampled over 0.25–0.55 and 0.05–0.15. These ranges describe idealized, dimensioned engineering circuits; they are not calibrated to patients or vascular beds.

The common initial state is the exact harmonic periodic response of the baseline linear system to continuous sinusoidal inflow, calculated through complex linear solves. The forecast then uses the piecewise-linear interpolation of supplied control samples; the initializer is not claimed to be its exact periodic response. Each trajectory lasts three cycles with 32 intervals per cycle. Four perturbations begin at 0.25, 1.00, 1.75, and 2.50 cycles, with cubic-smoothstep control values sampled over 0.25 cycle. Program A uses edge-resistance targets 0.65/1.30/0.85/1.15 and runoff-resistance targets 0.80/1.20/0.90/1.10 times baseline. Program B uses 1.40/0.80/1.25/0.90 and 1.25/0.75/1.15/0.85. Two distinct target edges are sampled without replacement using the background seed, one per program. The two structurally selected runoff sites are assigned to programs A and B in their fixed distance-based order. Every comparator receives the same sampled future controls, including their interval endpoints; all numerical methods use the same linear interpolation between samples. These actions are resistance perturbations, not simulated device deployment or clinical treatment effects.

The state comprises six pressures and five to seven edge flows, with unused coordinates padded to 13 and masked. Reference trajectories use DOP853 with relative tolerance 10^*−*10^ and absolute tolerance 10^*−*11^, restarting at every observed interval. Five fixed development backgrounds were checked using Radau, analytic stationary and harmonic solutions, and the passive energy identity

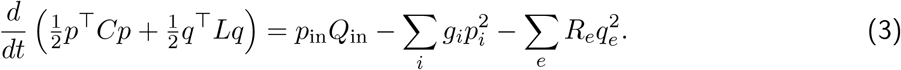

Midpoint integration with one, two, four, and eight substeps checked numerical convergence. Generation failures were to be recorded, investigated, and disclosed, without outcome-based exclusion or resimulation until a preferred model ranking appeared.

### 2.3 Representation, factor count, and supervised control

All neural models use the same graph interface. An entity is either a nodal pressure or an oriented edge flow. The encoder takes its standardized scalar state, two oriented sums of neighbouring entity states, and two entity-type flags. Shared 5–32–16 SiLU layers produce a 16-dimensional state per entity. Predictors receive the entity’s full current latent vector, two oriented sums of neighbouring latent vectors, six static features, and seven interval-control features. Static features encode the applicable log-compliance, log-runoff conductance, log-resistance, log-inertance, runoff flag, and inlet flag; controls encode inlet flow and resistance multipliers at both interval endpoints plus interval duration. Controls enter locally; there is no global control broadcast or graph pooling. The shared local message radius can therefore limit the immediate response to distant inputs. Padded states and inapplicable log-parameter, inlet-flow, and edge-control channels are masked. Runoff multipliers are supplied at every node, including their baseline value of one where runoff conductance is zero. State, log-parameter, and control normalizers use training episodes only, with pressure and flow scaled separately; binary type, inlet, and runoff flags are retained.

For online embedding *z*_*t*_ = *f*_*θ*_(*x*_*t*_) and an exponential-moving-average target encoder 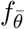, the factorized transition is

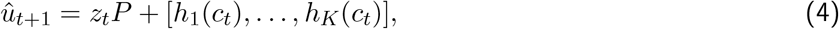

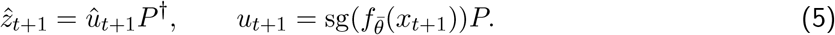

The context *c*_*t*_ contains the full available local context described above. Every head sees that same context: the factorization partitions the predicted coordinates, not independent causal mechanisms. The target encoder is updated with momentum 0.99. The loss combines equal-type latent mean squared error, 0.1 mean[(*P* ^*⊤*^*P −I*)^2^], a target-factor activity penalty of weight 0.05, and an online-embedding activity penalty of weight 0.02. For an embedding array *v*, activity is the equal-type average of 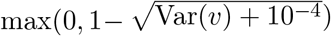 across coordinates, with variance computed over active entities of that type in the minibatch. This penalty is applied to *u*_*t*+1_ and *z*_*t*_, respectively; latent vectors are not normalized to unit length. The full-Gram orthogonality normalization is identical for *K* = 1 and *K* = 4, and orthogonality is a soft penalty rather than an exact constraint. This is a disclosed adaptation inspired by JEPA-Anything, not a reproduction of its complete training system or a new OPF algorithm.

The primary comparison is OPF-4 versus OPF-1. Both learn a 16 × 16 basis *P* and use the same objective. OPF-4 has four four-dimensional outputs, each with hidden width 32; OPF-1 uses one 16-dimensional head of hidden width 108 to match capacity. The secondary monolithic JEPA fixes *P* = *I* and uses one predictor of hidden width 112. Its target activity term is constant with respect to trainable weights because the target embedding is stop-gradient and its basis is fixed; therefore this secondary comparison changes more than factor count. The supervised graph model uses the same encoder/context interface with a hidden-width-138 head and directly predicts a scalar physical-state residual. Neural trainable counts are 9,472/9,416/9,440/9,415 for monolithic JEPA/OPF-1/OPF-4/supervised prediction, a maximum difference below 0.7%. JEPA models additionally have 720 non-optimized EMA parameters and 34 fitted readout coefficients.

Two frozen affine ridge readouts, one for pressure and one for flow, map training target-encoder latents to standardized physical states, with an unpenalized intercept. Ridge strengths 10^*−*6^, 10^*−*4^, 10^*−*2^, 1, 100 are chosen by reconstruction error pooled over active validation coordinates separately for each type. This additional readout selection is disclosed and is not performed for the supervised model. Pure latent rollout uses the online encoder on the observed initial state once, then recursively predicts latents and decodes them without future observations. The supervised model instead feeds back its scalar physical-state predictions and recomputes its encoder at each step. Pseudoinversion uses relative tolerance 10^*−*5^. Prespecified secondary diagnostics comprise target-encoding reconstruction, observation-refreshed one-step prediction, and decoded-state re-encoding between forecast steps. Re-encoding still receives no future truth, but changes the recursive map and is not the primary result.

### 2.4 Equal-budget selection and mechanistic baselines

All four neural arms receive the same six candidates: learning rates 3 × 10^*−*4^, 10^*−*3^, 3 × 10^*−*3^ crossed with weight decays 0 and 10^*−*4^. Every version-2 candidate uses 6,000 updates, seed 101, batch size 128, and gradient-norm clipping at 5. Selection minimizes final validation 96-step structure-macro RMSE, not training or one-step loss. Validation uses new backgrounds on *seen* structures, so it is not a structural transfer validation set. The selected configuration is refitted on training data only with seeds 211, 223, 227, 229, and 233 for exactly 12,000 updates; only the final checkpoint supplies test predictions. Fixed validation checks at 2,000/4,000/6,000 search updates and 4,000/8,000/12,000 final-fit updates diagnose budget sensitivity without best-checkpoint selection. The two budgets correspond to approximately 13.9 and 27.8 training-set-equivalent draws, sampled with replacement rather than complete epochs. No candidate, seed, or topology is removed for poor performance. The matching concerns candidate counts, updates, batch sizes, data, and approximate trainable capacity, not equal floating-point operations or wall-clock cost. Equal search counts and parameter budgets reduce specific comparison imbalances, but six candidates, one selection seed, and a fixed budget do not establish convergence or globally optimal tuning for any model.

The known-equation RLC baselines integrate Equations 1–2 by implicit midpoint with one or four substeps per observation interval. These are different numerical solvers from the DOP853 reference. The quasi-steady RC reduction sets edge inertia to zero and advances the resulting pressure system with four substeps. It starts from the supplied pressures and subsequently determines flows algebraically, so the independently observed inertial flow state is not preserved as a dynamic state in this reduction. Persistence repeats the initial observations. A further structured baseline fits five global coefficients multiplying inlet flow, incidence flow, runoff, pressure gradient, and resistive flow terms. Training-only regression uses midpoint states/controls and interval differences, with equal pressure/flow weighting after scaling. It has no intercept and no hard coefficient constraints. Five ridge strengths match the readout grid; selection uses validation 96-step rollout under four-substep midpoint integration. Unit coefficients recover the known equations. Fitted deviations may absorb coarse-time differencing bias and do not constitute discovery of new physiology.

Every method receives the full initial state, true topology and parameters, and prescribed future control schedule. Observations and supplied parameters are noiseless. The mechanistic methods additionally encode the correct equation structure. They are an explicitly favorable known-physics comparator, useful for evaluating whether learning is necessary in this information setting. No claim of matched algorithmic knowledge or unknown-parameter clinical inference is made.

### 2.5 Endpoints, uncertainty, and failure analysis

For background *b*, let *e*^*p*^ and *e*^*q*^ be pressure and active-edge flow errors divided by the corresponding training standard deviations. The primary error is

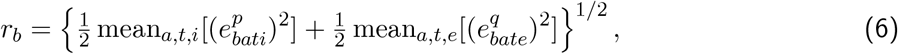

where the means include both action programs and all 96 future steps. Errors are averaged equally across backgrounds within each graph, then equally across graphs. Training-seed means and standard deviations are separate quantities. The primary contrast is OPF-4 minus OPF-1 on the 14 withheld graphs after averaging the five seed results for each graph. Positive values favor OPF-1. A paired graph bootstrap draws within the 5/6/7-edge strata, retaining 2/5/7 graph draws per replicate, with 10,000 replicates and fixed seed 2026101131. Reported 95% percentile intervals are conditional on this catalogue, sampled backgrounds, split, and fixed training seeds. They describe sensitivity to resampling these structures, not confidence about an external anatomical population. Fourteen graphs, not 336 episodes or five training seeds, are the resampling units. Other contrasts are exploratory.

Secondary measures include physical-unit MAE, horizon-specific and per-structure error, nonfinite forecasts, projection singular values, coordinate activity, and permutation-equivariance checks. The latter test simultaneous node/edge relabeling; they do not establish equivariance to reversing edge orientation and flow signs, nor do they prove physical conservation. Conservation diagnostics evaluate trapezoidal finite-difference residuals of Equations 1–2; even the reference has a discretization residual, which is reported alongside model residuals. They are not exact continuous conservation errors. A displayed example was fixed before testing: the lowest atlas-index withheld graph, its lowest-index background, program A, and seed 211. No favorable trajectory is selected after inspecting outcomes.

Neural runs use PyTorch 2.11.0+cu128 on an NVIDIA RTX 5070 Ti, with two PyTorch CPU threads and a process memory cap. Search-stage NumPy used its default 24-thread OpenBLAS configuration; final fits explicitly limited BLAS/OMP/MKL to two threads. This resource-only change, including its possible effect on timing and low-order floating-point reductions, is recorded separately and did not trigger reselection or exclusion. Other user computation shared the GPU. Numerical baselines use sequential CPU integration with one-thread BLAS. Raw timings, batch sizes, and host-transfer scope are archived; these measurements do not support a matched-hardware speedup claim.

## 3 Results

### 3.1 Numerical verification and complete selection

The reference-generation checks supported numerical consistency of the specified circuit. Across five fixed development fixtures, the maximum native-coordinate difference between DOP853 and Radau was 2.97 × 10^*−*8^; stationary residuals were below 7.3 × 10^*−*12^ and the continuous-harmonic check differed by less than 6.3 × 10^*−*9^. The passive-energy identity held to relative residual below 2.2 × 10^*−*15^. These are numerical checks of the implemented equations, not physiological validation. All 1,296 planned development and test episodes were generated, with no generation failures.

All 24 amended search runs completed 6,000 updates, with no training failures or nonfinite final selection scores. Their first 1,000-update model tensors matched the archived lower-budget runs exactly. All 72 intermediate evaluation checks preserved the model, optimizer, and random-number state. The selected learning-rate/weight-decay pairs were 3 × 10^*−*4^*/*10^*−*4^ for monolithic JEPA, 3 × 10^*−*3^*/*10^*−*4^ for OPF-1, 10^*−*3^*/*10^*−*4^ for OPF-4, and 3 × 10^*−*4^*/*10^*−*4^ for supervised prediction. Their validation RMSEs were, respectively, 3.05 × 10^17^, 1.38 × 10^8^, 3.83 × 10^10^, and 7.30 × 10^5^. Finite scores of this size are severe prediction errors, not evidence of stable or usable dynamics. Figure 3 and Table 3 retain the complete search, and Figure 7 reports the fixed-checkpoint validation trajectories.

**Figure 3:**
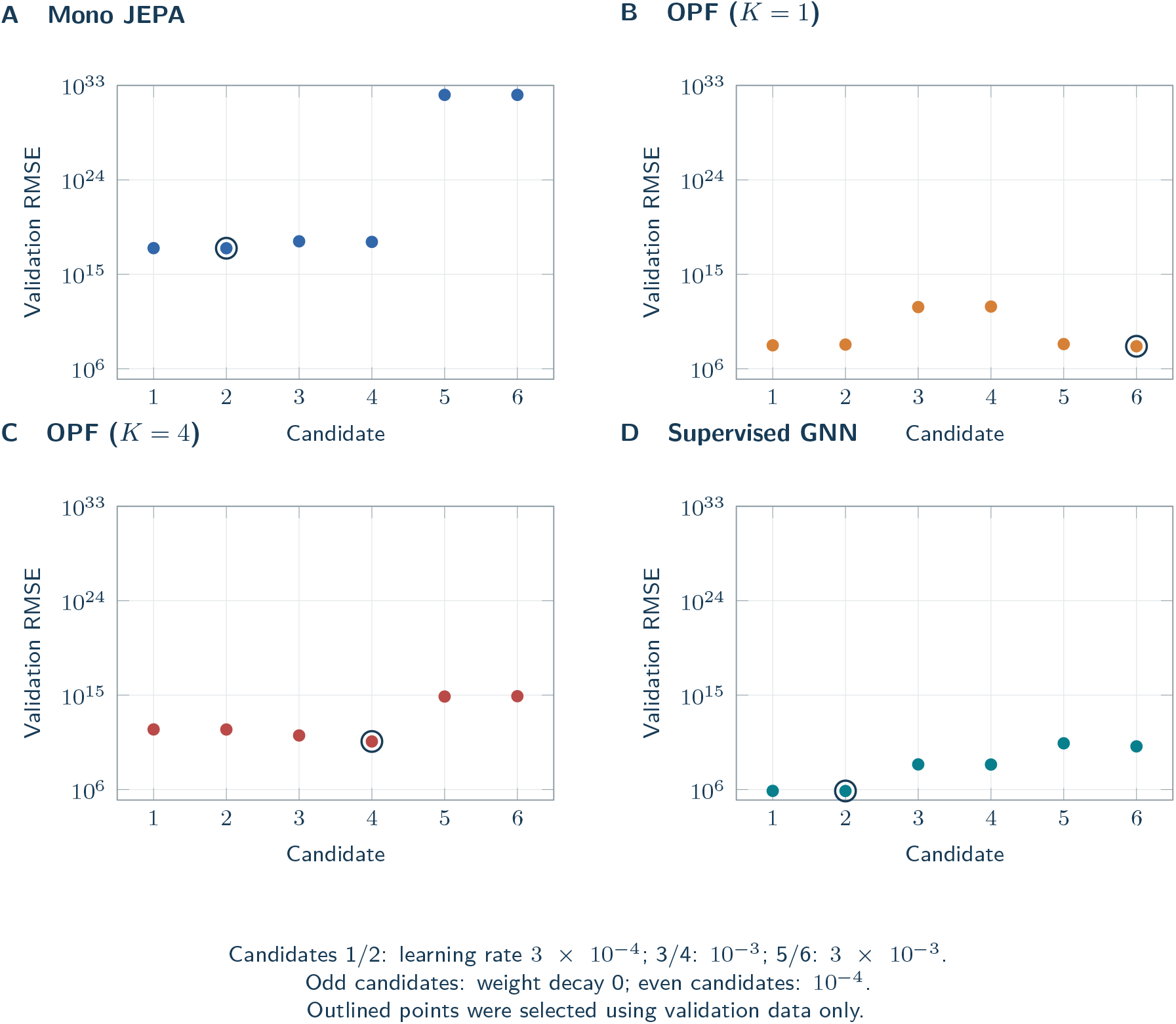
Equal-budget development search. Each architecture used the same six learning-rate/weight-decay combinations, 6,000 updates per candidate and development seed 101. The vertical log scale is identical across panels and displays the complete finite range. Crosses, if present, denote nonfinite or failed candidates rather than omitted trials. Selection uses the 96-step equal-structure macro-RMSE on new parameter backgrounds from seen structures; these are not held-out-structure test results.

All 20 final fits completed the prescribed 12,000 updates and produced both test splits. There were 0 nonfinite active scalar predictions across the primary neural rollouts. No seed was excluded or retrained after testing. Numerical finiteness is reported separately from the magnitude of prediction error.

### 3.2 Autonomous prediction on withheld structures

Table 1 reports the full comparison. On unseen structures, the five-seed means were Monolithic JEPA 1.49 × 10^12^, OPF-1 329, OPF-4 8.46 × 10^9^, Supervised graph 1.83 × 10^31^. The primary OPF-4 minus OPF-1 difference was 8.46 × 10^9^, with a paired stratified-graph 95% percentile interval of [2.23 × 10^5^, 2.54 × 10^10^]. All four neural means exceeded persistence. Thus, a relative advantage within this comparison would not establish a usable forecasting model. The resampling interval is conditional on 14 graph structures and the fixed set of training seeds, rather than an uncertainty estimate for a clinical population.

**Table 1:**
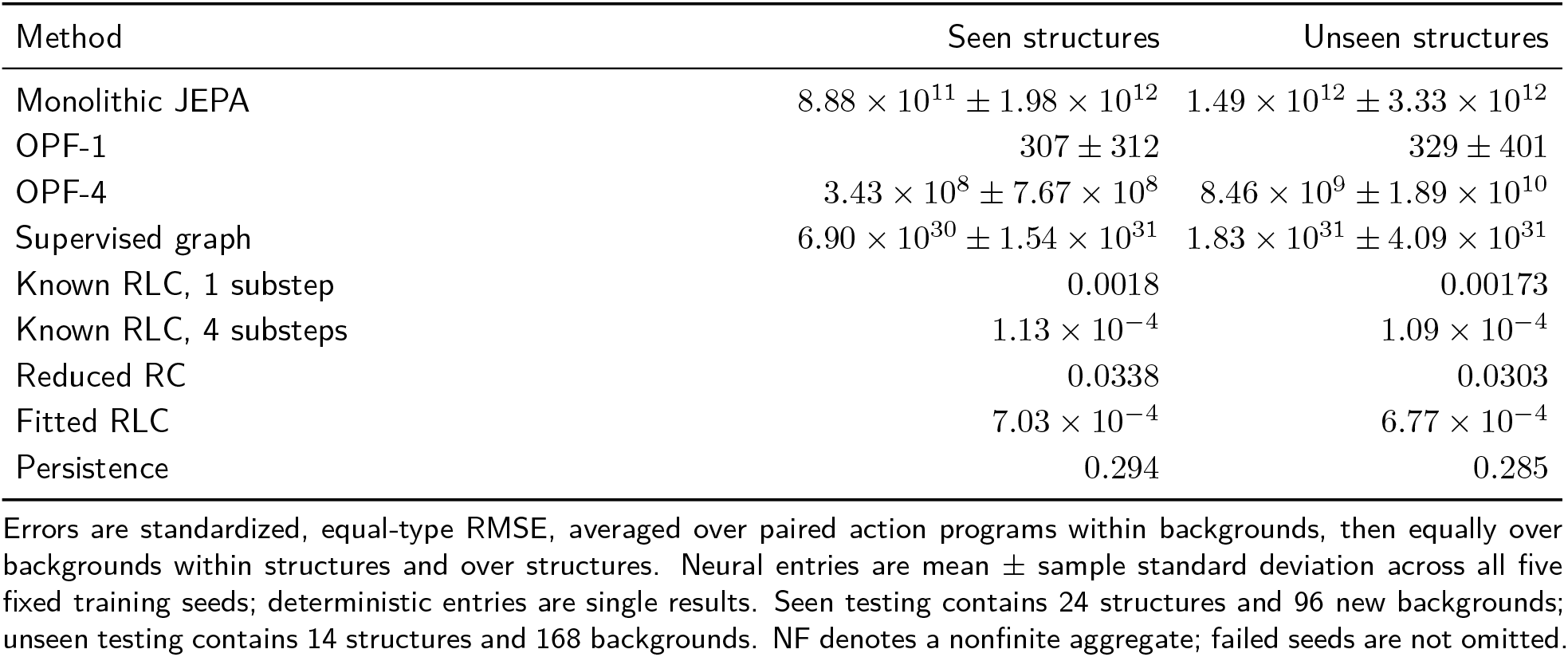
Autonomous 96-step prediction after frozen model selection.

| Method | Seen structures | Unseen structures |
| --- | --- | --- |
| Monolithic JEPA | $8.88 \times 10^{11} \pm 1.98 \times 10^{12}$ | $1.49 \times 10^{12} \pm 3.33 \times 10^{12}$ |
| OPF-1 | $307 \pm 312$ | $329 \pm 401$ |
| OPF-4 | $3.43 \times 10^8 \pm 7.67 \times 10^8$ | $8.46 \times 10^9 \pm 1.89 \times 10^{10}$ |
| Supervised graph | $6.90 \times 10^{30} \pm 1.54 \times 10^{31}$ | $1.83 \times 10^{31} \pm 4.09 \times 10^{31}$ |
| Known RLC, 1 substep | 0.0018 | 0.00173 |
| Known RLC, 4 substeps | $1.13 \times 10^{-4}$ | $1.09 \times 10^{-4}$ |
| Reduced RC | 0.0338 | 0.0303 |
| Fitted RLC | $7.03 \times 10^{-4}$ | $6.77 \times 10^{-4}$ |
| Persistence | 0.294 | 0.285 |

Errors are standardized, equal-type RMSE, averaged over paired action programs within backgrounds, then equally over backgrounds within structures and over structures. Neural entries are mean ± sample standard deviation across all five fixed training seeds; deterministic entries are single results. Seen testing contains 24 structures and 96 new backgrounds; unseen testing contains 14 structures and 168 backgrounds. NF denotes a nonfinite aggregate; failed seeds are not omitted.

**Table 2:** Physical-coordinate errors on the 14 unseen structures.

| Method | Pressure MAE | Flow MAE | Nonfinite |
| --- | --- | --- | --- |
| Monolithic JEPa | $1.89 \times 10^{12}$ | $1.52 \times 10^{13}$ | 0 |
| OPF-1 | $1.69 \times 10^3$ | $4.52 \times 10^3$ | 0 |
| OPF-4 | $1.65 \times 10^{10}$ | $8.34 \times 10^{10}$ | 0 |
| Supervised graph | $3.28 \times 10^{31}$ | $1.03 \times 10^{32}$ | 0 |
| Known RLC, 1 substep | 0.0112 | 0.0655 | 0 |
| Known RLC, 4 substeps | $6.93 \times 10^{-4}$ | 0.00414 | 0 |
| Reduced RC | 0.287 | 1.03 | 0 |
| Fitted RLC | 0.0129 | 0.0106 | 0 |
| Persistence | 4.75 | 6.66 | 0 |

Pressure is in mmHg and flow in mL/s. MAEs use background-then-structure averaging and, for neural methods, the mean of all five seeds. The last column counts nonfinite active scalar predictions across those seeds; a zero count does not imply that a very large finite forecast is stable or useful.

Descriptive inspection of the retained graph and seed results showed a positive OPF-4 minus OPF-1 difference for each of the 14 graph means and each of the five paired seed means. Its magnitude, however, was dominated by graph atlas_92 and OPF-4 seed 229. The very large aggregate difference therefore should not be interpreted as a typical per-structure effect. This observation does not change the prespecified estimator or interval; every graph and seed remains in the analysis.

### 3.3 Known and fitted mechanisms

On the 14 unseen structures, known-parameter midpoint RLC integration gave standardized RMSE 0.001730 with one substep and 0.0001089 with four substeps, a 15.9-fold reduction consistent with the independent numerical-convergence check. The four-substep model’s pressure and flow MAEs were 0.000693 mmHg and 0.00414 mL/s. The zero-inertance RC reduction had RMSE 0.03032, pressure MAE 0.287 mmHg, and flow MAE 1.031 mL/s. This measures the cost of that specified simplification for these trajectories; it does not establish the importance of inertia in an unmeasured clinical setting.

The training-fitted RLC model selected ridge strength 10^*−*4^ and coefficients 1.00104, 1.00133, 1.00132, 1.00599, and 1.00597 for inlet, incidence, runoff, pressure-gradient, and resistance terms. Its unseen-structure RMSE was 0.0006773, with pressure MAE 0.01290 mmHg and flow MAE 0.01063 mL/s. Persistence had RMSE 0.2845. The fitted mechanism transferred across this graph catalogue but did not improve on the matched four-substep integration with the supplied exact coefficients. These comparisons have the disclosed advantage of correct governing structure.

### 3.4 Recursive-error and representation diagnostics

For monolithic JEPA, OPF-1, and OPF-4, respectively, mean unseen-structure reconstruction RMSEs were 0.0112, 0.0267, 0.012; observation-refreshed one-step RMSEs were 0.043, 0.0491, 0.0395. The corresponding decoded-state re-encoding RMSEs were 9.14 × 10^29^, NF, NF. These diagnostics do not replace the primary free latent rollout. The supervised model had one-step RMSE 0.0145; its physical-state rollout is also its re-encoding diagnostic by construction. Training-only normalization standard deviations were 16.9566 mmHg for pressure and 38.8960 mL/s for flow. Nonfinite re-encoding forecasts occurred for OPF-1 (2/5 fits), OPF-4 (2/5 fits); their all-seed aggregates are reported as NF, without averaging only the remaining finite runs. All ten learned projection matrices retained all 16 singular directions at the declared pseudoinverse tolerance; their condition numbers ranged from 1.16–1.43. Algebraic completeness alone therefore did not certify accurate recursive prediction. Coordinate activity and effective ranks are reported in the implementation diagnostics appendix.

Figure 4 displays all methods, seeds, and paired per-structure contrasts; Figure 5 separates forecast horizons and diagnostic tasks. Figure 6 uses the case fixed before test generation, retaining full-range errors rather than selecting a favorable trajectory. Table 4 reports every seed and Table 5 includes the reference’s discretization residual alongside model residuals.

**Figure 4:**
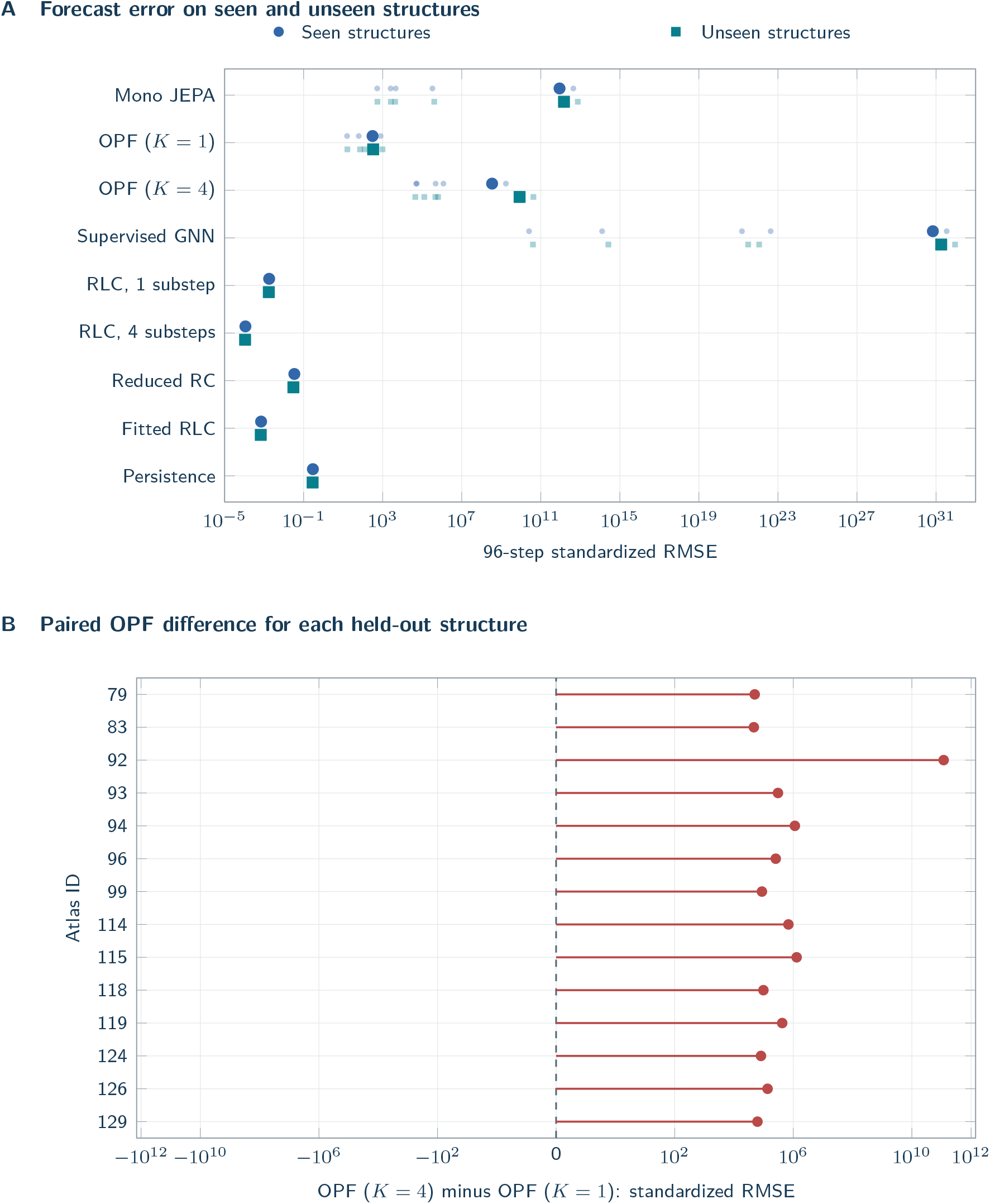
Primary and structure-specific comparisons. (A) Large points are means across the five fixed neural training seeds; small points show the individual seeds. Deterministic comparators have one result. Both axes include the complete finite range without clipping divergent runs. (B) Differences are computed after averaging seeds within each held-out structure; positive values favour *K* = 1. To retain both sign and large magnitudes, horizontal coordinates use sign(*d*) log_10_(1 + |*d*|*/*0.01), with ticks labelled in original RMSE units. These are paired differences for the fixed catalogue, not fourteen independent patient cohorts.

**Figure 5:**
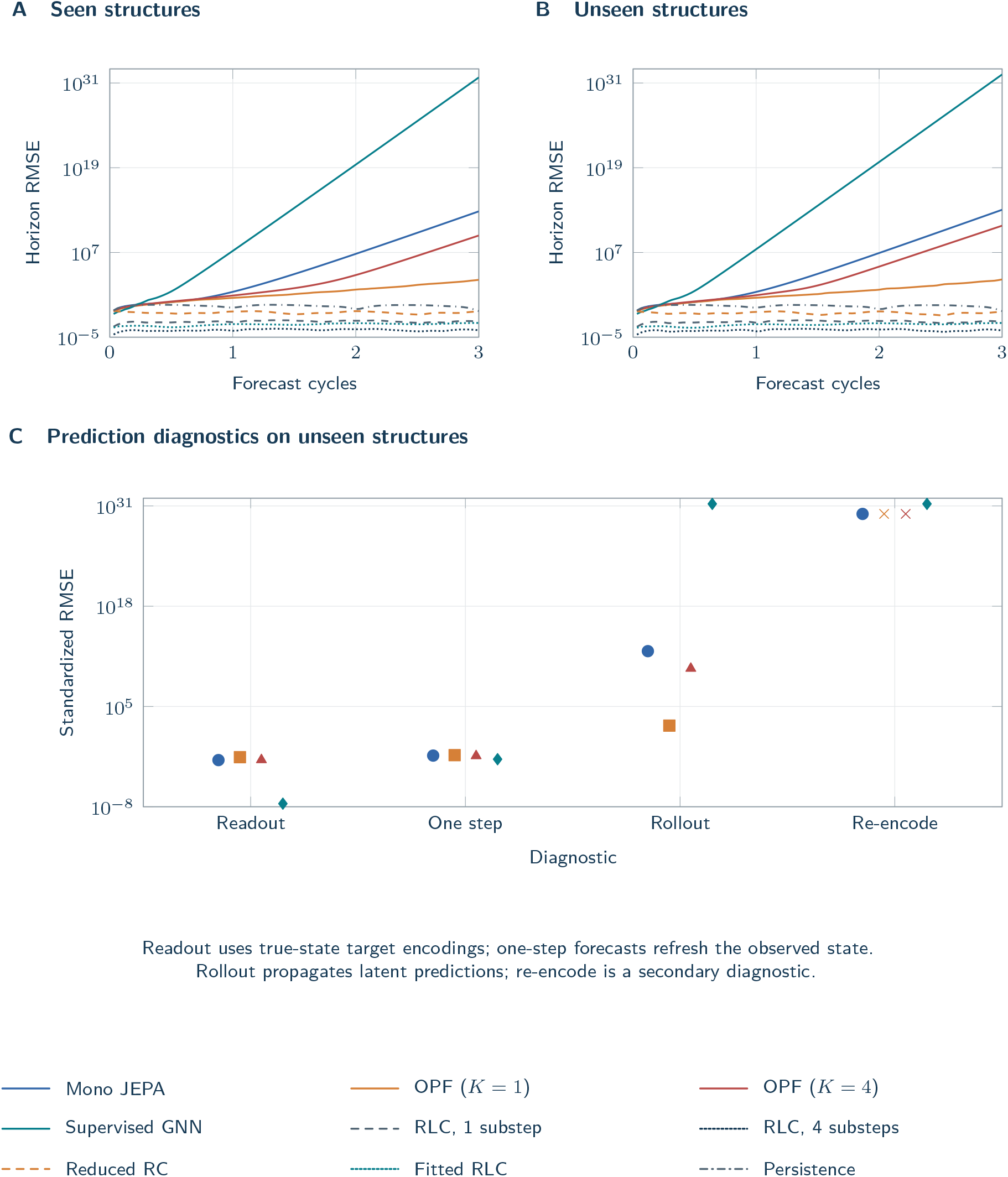
Forecast horizon and representation/rollout diagnostics. (A,B) Equal-structure mean errors at each of 96 forecast steps use identical logarithmic axes; every planned model is included. (C) Means across the five neural seeds separate readout reconstruction, observation-refreshed one-step prediction, primary free rollout, and secondary decoded-state re-encoding. A label of zero denotes an exact identity/zero error displayed at the log-axis floor; crosses denote nonfinite or unavailable diagnostics. Large rollout errors are retained, not replaced by a more favourable diagnostic.

**Figure 6:**
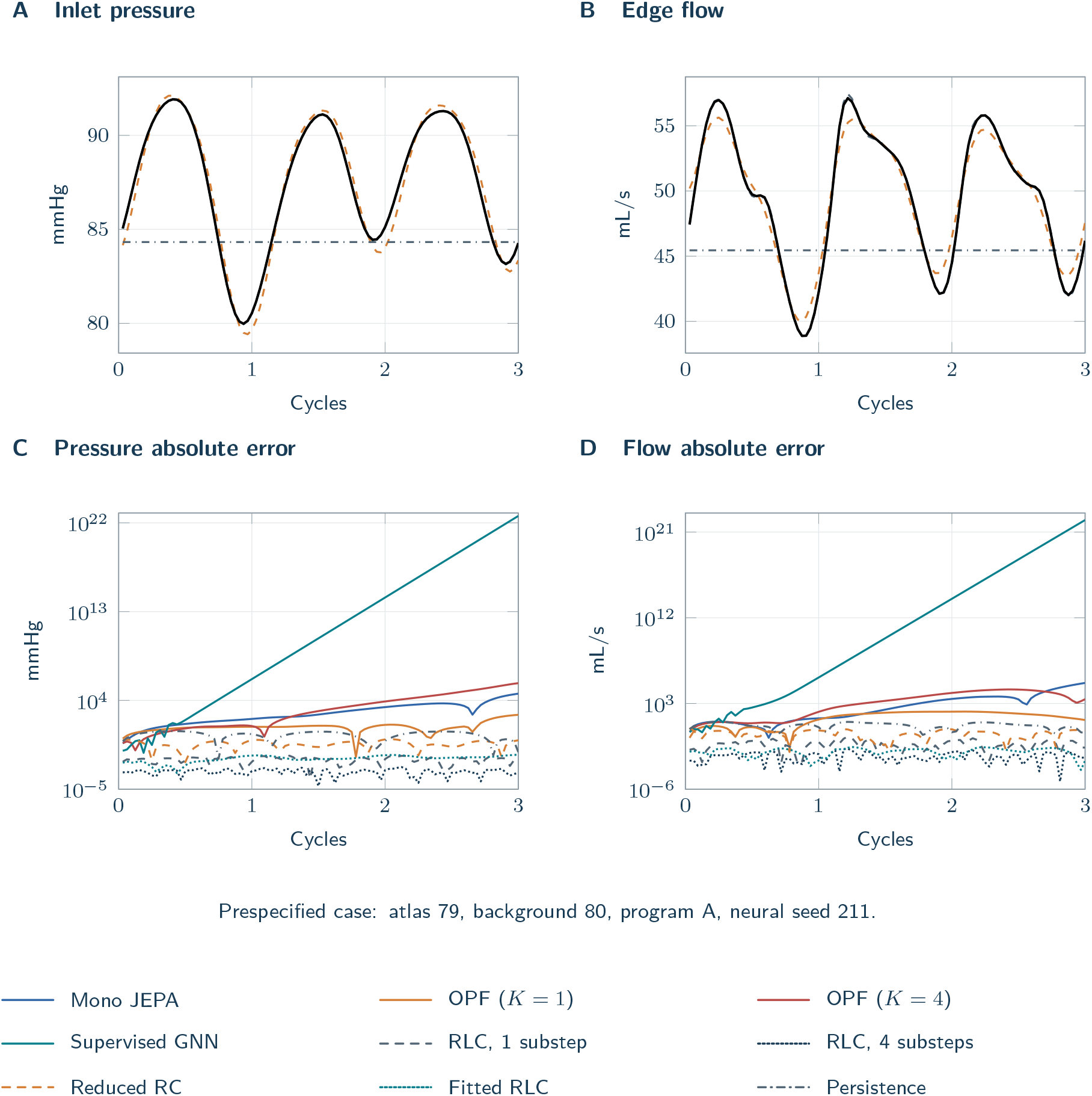
A case selected before viewing test outputs. (A,B) The black line is the reference; known-parameter RLC, reduced RC, fitted RLC and persistence predictions are shown on physical axes. Pressure is shown on the specified reference scale, including the 5 mmHg distal datum. (C,D) Absolute errors include every method, including all neural predictions, over the complete finite logarithmic range. This separates readable reference dynamics from potentially very large neural errors without truncating unstable predictions. No representative or favourable example was selected retrospectively. Zero errors, if any, are displayed at the recorded positive log floor; nonfinite values are flagged explicitly.

## 4 Discussion

The fixed five-seed mean error of OPF-4 was higher than that of OPF-1 on the withheld structures. All four neural means exceeded persistence. Thus, a relative advantage within this comparison would not establish a usable forecasting model. The defensible conclusion is specific to these local graph architectures, data, one-step objectives, and fixed optimization/rollout protocol. It does not establish a universal ranking of factorized representations, a failure of JEPA-Anything in its published settings, or superiority of numerical integration when the true equations and parameters are unavailable.

The distinction between coordinate geometry and dynamical stability is important. For fixed parameters and interval controls, when the pseudoinverse retains all 16 singular directions, the implemented transition on active entities has the residual form *F*_*t*_(*z*) = *z* + *H*_*t*_(*z*), up to floating-point arithmetic, where *H*_*t*_ includes the learned residual and basis synthesis. Consequently, its state Jacobian is *D*_*z*_*F*_*t*_ = *I* + *D*_*z*_*H*_*t*_. Making the basis well conditioned does not constrain the spectrum of this recurrent map or enforce the circuit’s mass balance and passive energy relation. This algebraic observation explains why projection diagnostics alone cannot certify a usable world model; we did not estimate Jacobian spectra or prove that a particular instability mechanism caused every observed error.

Reconstruction and observation-refreshed one-step scores answer different questions from autonomous forecasting. The latter repeatedly evaluates a learned map on its own outputs, which can leave the set of encoded observations encountered during training. During training, each JEPA predictor consumes online-encoder states and predicts the next EMA-encoder state; pure latent rollout starts with the online encoder but, from its second step, consumes its own predictions of target-space states. The EMA update does not enforce exact coordinate equality between the two encoders, and the readout is fitted to EMA states. This interface is shared by all three JEPA arms. Encoder lag, prediction error, departure from the training representation distribution, and the absence of a multistep objective may coexist; without targeted interventions, the diagnostics do not identify a unique cause or form an additive error decomposition. Decoding and re-encoding changes the recurrent map, but does not enforce the equations and is not a general stability guarantee. Conversely, a model that performs poorly on both seen and unseen structures has not isolated structural novelty as the source of its error. The protected structure split prevents an easy form of leakage; it does not make an inaccurate model useful.

These internal comparisons are not tests of the best available neural simulator. The cardiovascular graph surrogate of Pegolotti et al. uses five-step training strides, input-noise augmentation, and additional boundary-to-interior connections [3]; the compact one-step models here do not reproduce that design. Our OPF adaptation also does not reproduce the domain-specific backbones and base-plus-OPF objectives of JEPA-Anything [6]. Results on this circuit benchmark therefore do not overturn performance reported for those distinct implementations and datasets.

The known-equation comparison also sets the appropriate interpretation of the fitted five-coefficient model. Its equation library is correct by construction and its coefficients are shared across every network. The observed near-unit estimates and small prediction errors support accurate numerical fitting within that supplied equation family; they do not establish discovery of unknown vascular biology or preservation of the original energy identity by unconstrained coefficients. The selected positive coefficients do admit a modified continuous-time storage inequality, derived as a post hoc interpretation in Appendix B.4. This illustrates the structural advantage supplied to the mechanistic comparator without asserting stepwise energy decrease in the driven numerical trajectories. When equations, parameters, initial states, and future controls are already supplied, numerical integration is a relevant alternative to a neural surrogate. Learning must justify its role in a different information or computational setting; unmatched CPU/GPU timings cannot provide that justification here.

### 4.1 Independence from related work

The related author manuscript *PV-JEPA: A Capacity-Controlled Predictive-State Benchmark for Cross-Cohort Vascular Transcriptomics* was submitted to *European Heart Journal – Digital Health* before this revision. The related VascFlux-GEO manuscript analyzes overlapping public vascular transcriptomic cohorts. Those manuscripts require their own overlap disclosure and are not independent replications of each other. No GEO participants, gene modules, expression matrices, out-of-fold predictions, clinical effect estimates, or result tables from either manuscript enter the new graph experiment. Shared authorship, predictive-state motivation, and parts of the general machine-learning methodology are acknowledged. The current work’s distinct evidence is the newly generated structure-held-out RLC experiment, its locked model search, and its known-equation comparisons. Reusing the JEPA name or obtaining another negative comparison would not alone create a separate scientific contribution.

The older single-bifurcation pilot is part of this study’s development history, not a second independent confirmation. It is summarized in Appendix A; its source, data, and analyses are preserved. The former broad prospective clinical schematics have been removed from the empirical argument. These disclosures should accompany any later editorial assessment of the revised manuscript.

### 4.2 Limits and next experiments

All 38 structures are small idealized circuits, including cycles that need not map to a particular vascular bed. The 14 withheld classes are sampled from one finite six-node catalogue, and rooted boundary placements are determined by a simple rule. The same two relative action programs occur in development and testing; transfer to an unseen action schedule was not evaluated. Network size, patient anatomy, nonlinear stenosis, pressure-dependent material behavior, incomplete sensing, uncertain parameters, changing topology, and unobserved future actions remain untested. Repeated prescribed resistance changes are not counterfactual patient treatment effects. Accurate acute pressure/flow prediction would not validate endothelial repair, thrombosis, or clinical benefit; the IABP observational study [8] cannot supply that missing evidence.

The neural objectives optimize one-step transitions, whereas the principal endpoint is a 96-step autonomous forecast. Learning-rate and weight-decay selection at 6,000 updates need not identify the best configuration at the 12,000-update final-fit horizon. No validation-based early stopping was used. Matching parameter counts also required different head widths, so the primary contrast concerns the specified matched-capacity implementations, not factor count isolated from every architectural difference. Multistep training, invariant-preserving maps, adaptive temporal resolution, richer message passing, and larger searches may change the outcome; none was added after seeing test performance. The structured baseline has the correct equation library and the known-parameter solver has the correct coefficients, advantages unavailable when real mechanisms or parameters are unknown. A future clinical or anatomy-derived study must define a new dataset and locked evaluation, rather than reinterpreting these synthetic results as patient evidence.

## 5 Conclusion

This newly generated, structure-held-out circuit benchmark separates predictive-coordinate design from recurrent forecasting performance. Four-factor OPF did not improve the prespecified comparison with single-factor OPF, and all four neural means exceeded persistence. These negative results support a bounded assessment of the implemented models, factor count, fixed-budget tuning, and supplied mechanisms; they do not establish a clinically validated vascular world model. Future claims of vascular applicability require stable prediction under an appropriate training design and validation on independently acquired anatomical or biological data.

## Supporting information

Supplementary Data - Compact audit, summary tables, code, and figure sources

## Ethics and data availability

This work analyzes newly generated numerical trajectories. It involves no recruitment, identifiable patient records, tissue experiment, or animal experiment, and claims no ethics approval or exemption. Ethics and governance identifiers from cited publications are not approvals for this study.

The author-review reproducibility package includes the locked protocol and manifests, generator and model code, numerical checks, all search outcomes, selected configurations, test predictions, fitted checkpoints, independent recalculations, figure-generation code, and manuscript source. No separate public repository or public deposition is claimed at this review stage. The previous pilot remains archived with its original svZeroDSolver provenance and license. No restricted patient data are included. The current revision has not been uploaded or submitted.

## Author contributions

Lingsen You, Zijie Feng, Chang Gao, Yingcheng Wu, Zhenfei Yin, and Ling Yang share first authorship and contributed equally to this work.

## DECLARATION OF GENERATIVE AI AND AI-ASSISTED TECHNOLOGIES

Codex assisted with LaTeX assembly, prose editing, and consistency checks. The authors reviewed and revised all AI-assisted text and take responsibility for the submitted manuscript.

## A Exploratory pilot retained for provenance

The earlier single-topology experiment comprised 320 parameter backgrounds and 3,040 trajectories using archived svZeroDSolver components, release 2021.09.30, commit 53eefab47e36ddc433abbac3 6250eb7239929591 [11]. It used one resistor bifurcation, two RCR outlets, two independent dynamic states, and 20 forecast steps. In-distribution RMSE was 0.0716*±*0.0079 for the full OPF implementation, 0.0478*±*0.0074 for the regularized monolithic JEPA, and 0.0357*±*0.0037 for a supervised MLP. OPF minus monolithic JEPA was 0.0239 with a background-bootstrap 95% interval of 0.0218–0.0261. These are historical pilot results from the archived analysis, not recomputed or pooled with the new graph experiment. The old objective, topology, capacity control, and horizon differ, so numerical differences between the studies do not estimate improvement caused by the revision.

## B Complete selection and implementation diagnostics

### B.1 Complete hyperparameter search and budget sensitivity

The initial 1,000-update search is retained in Figure 8. It was development evidence that motivated the documented budget amendment, not an independent confirmation. The complete amended search appears in Table 3; Figure 7 shows all six search candidates and all five final-fit seeds at every fixed validation checkpoint. No checkpoint was selected retrospectively by early stopping, and no budget was extended after test inspection.

**Figure 7:**
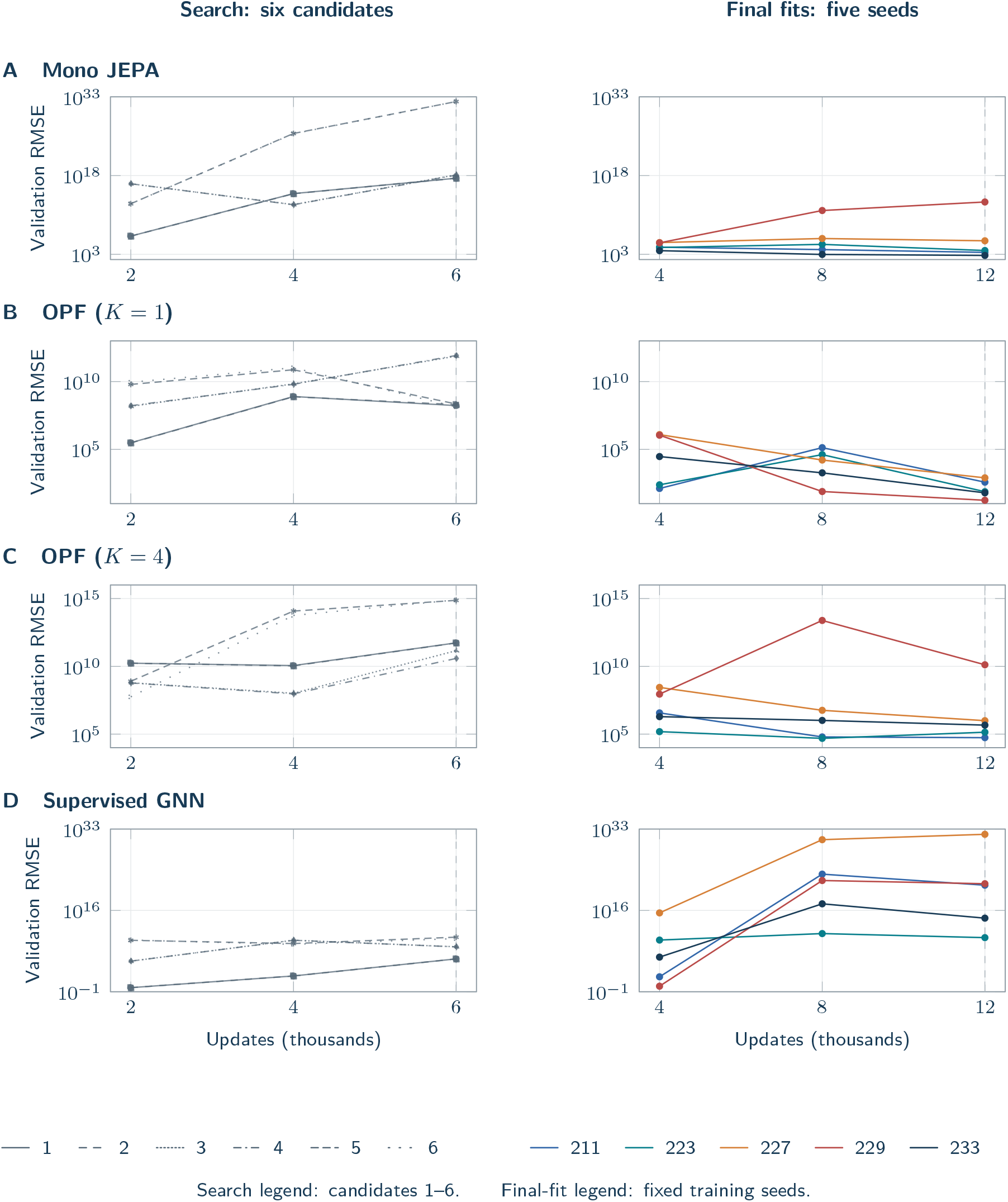
Development error as training budget increases. Each row is one architecture, with all six search candidates (left: 2,000, 4,000 and 6,000 updates) and all five final-fit training seeds (right: 4,000, 8,000 and 12,000 updates). Both columns share the same complete finite logarithmic range within each row; ranges may differ between rows. The displayed values are 96-step validation rollout errors on seen structures. The fixed 6,000-update candidate scores determined hyperparameter selection; the fixed 12,000-update models generated test forecasts. Intermediate checkpoints are descriptive and were not selected by early stopping. Crosses indicate observed checkpoints with nonfinite rollout error; F marks a checkpoint unavailable after a recorded training failure. Adjacent line segments are omitted rather than interpolated through failures. Candidate numbers match the search-comparison figure; no run is removed from this display.

**Figure 8:**
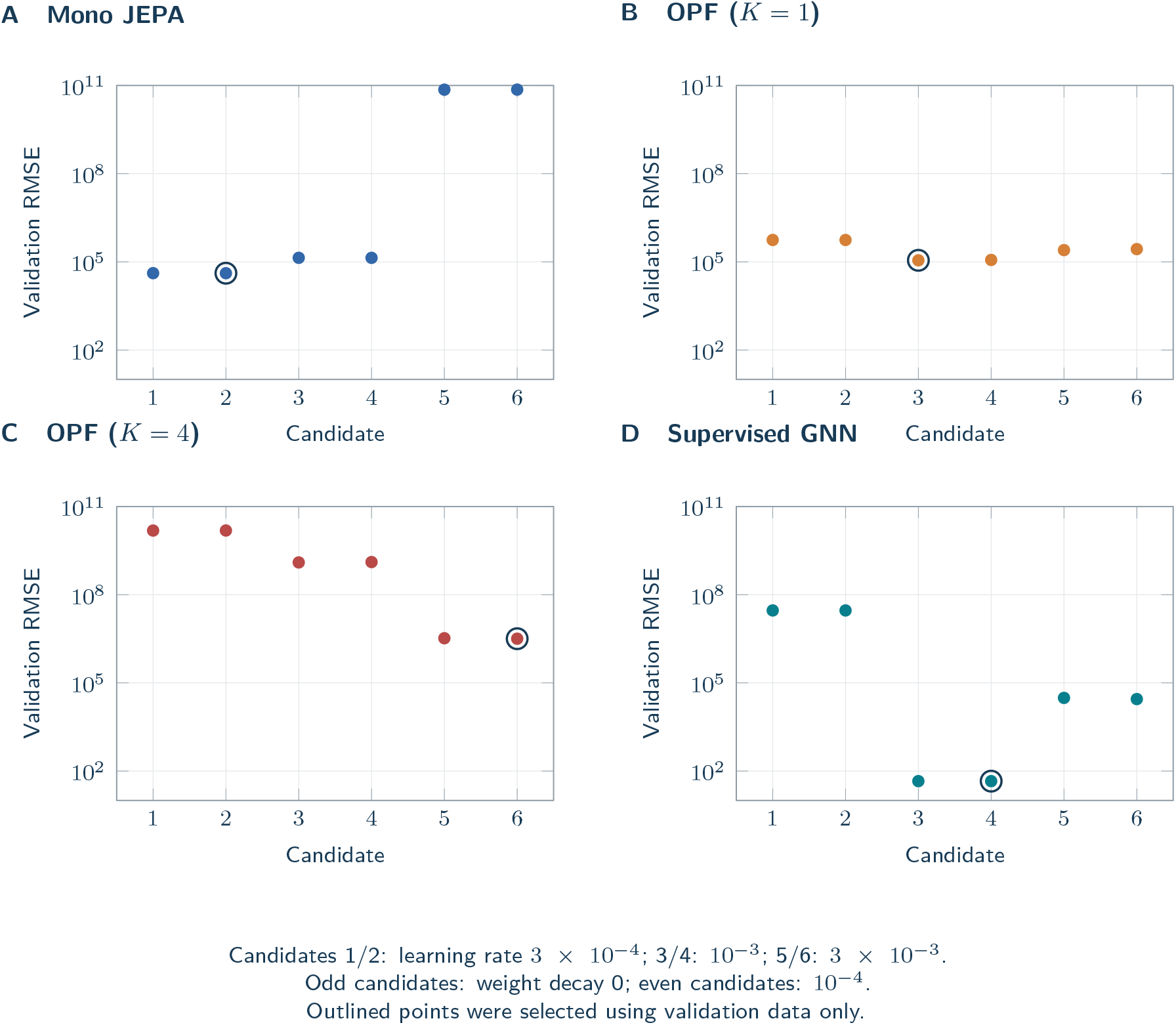
Initial lower-budget development search, retained as an explicit budget comparison. Each architecture used the same six learning-rate/weight-decay combinations, 1,000 updates per candidate and development seed 101. The vertical log scale is identical across panels and displays the complete finite range. Crosses, if present, denote nonfinite or failed candidates rather than omitted trials. Selection uses the 96-step equal-structure macro-RMSE on new parameter backgrounds from seen structures; these are not held-out-structure test results.

**Table 3:** Complete six-candidate search at the fixed 6,000-update checkpoint.

| ID | LR | WD | Mono | OPF-1 | OPF-4 | Supervised |
| --- | --- | --- | --- | --- | --- | --- |
| 1 | $3.00 \times 10^{-4}$ | 0 | $3.12 \times 10^{17}$ | $1.70 \times 10^8$ | $5.37 \times 10^{11}$ | $7.46 \times 10^5$ |
| 2 | $3.00 \times 10^{-4}$ | $1.00 \times 10^{-4}$ | $3.05 \times 10^{17*}$ | $2.00 \times 10^8$ | $5.28 \times 10^{11}$ | $7.30 \times 10^{5*}$ |
| 3 | 0.001 | 0 | $1.39 \times 10^{18}$ | $7.53 \times 10^{11}$ | $1.43 \times 10^{11}$ | $2.49 \times 10^8$ |
| 4 | 0.001 | $1.00 \times 10^{-4}$ | $1.23 \times 10^{18}$ | $8.42 \times 10^{11}$ | $3.83 \times 10^{10*}$ | $2.42 \times 10^8$ |
| 5 | 0.003 | 0 | $1.26 \times 10^{32}$ | $2.23 \times 10^8$ | $7.34 \times 10^{14}$ | $2.55 \times 10^{10}$ |
| 6 | 0.003 | $1.00 \times 10^{-4}$ | $1.26 \times 10^{32}$ | $1.38 \times 10^{8*}$ | $7.92 \times 10^{14}$ | $1.30 \times 10^{10}$ |

LR: learning rate; WD: weight decay. Entries are validation 96-step structure-macro RMSE at seed 101; asterisks mark the prespecified minimum-score selection. No earlier checkpoint was substituted. Values are rounded for display; full precision and every checkpoint are in the archived selection files.

**Table 4:**
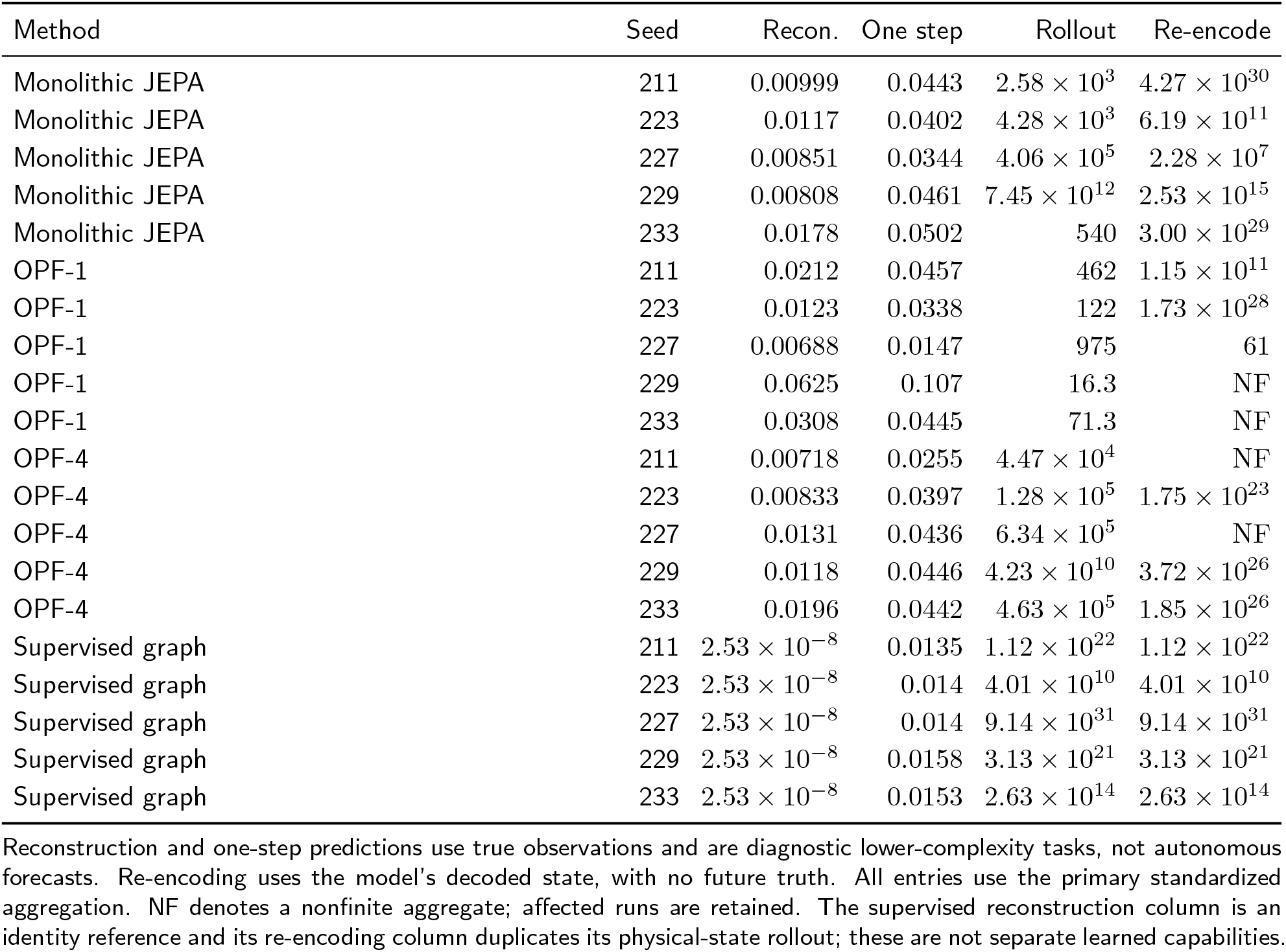
Every fixed-seed diagnostic on unseen structures.

**Table 5:** Discrete balance residuals on unseen structures.

| Method | Mass balance | Momentum balance |
| --- | --- | --- |
| Monolithic JEPa | $1.23 \times 10^{14}$ | $3.27 \times 10^{13}$ |
| OPF-1 | $2.29 \times 10^4$ | $6.11 \times 10^3$ |
| OPF-4 | $7.35 \times 10^{11}$ | $1.65 \times 10^{11}$ |
| Supervised graph | $2.51 \times 10^{33}$ | $4.16 \times 10^{32}$ |
| Known RLC, 1 substep | 0.00717 | $4.02 \times 10^{-4}$ |
| Known RLC, 4 substeps | 0.0471 | 0.00371 |
| Reduced RC | 0.351 | 0.348 |
| Fitted RLC | 0.051 | 0.00545 |
| Persistence | 10.3 | 0.706 |
| DOP853 reference | 0.0506 | 0.00393 |

Mass-balance residual is in mL/s and momentum-balance residual in mmHg. Trapezoidal differences at the observation interval have a discretization floor, including for the DOP853 reference. Entries average trajectory RMS residuals within each graph and then equally across graphs and fixed seeds. They do not certify continuous-time conservation or clinical plausibility.

### B.2 Every fixed-seed diagnostic

Development-set representation checks covered all 15 JEPA fits. The numerically retained projection ranks were 16 at relative pseudoinverse tolerance 10^*−*5^. Learned-basis condition numbers ranged from 1.16–1.43. Pressure-coordinate standard deviations spanned 1.05–2.12, and flow-coordinate standard deviations spanned 1.06–1.98. The entropy-based effective ranks of centered EMA embeddings were 2.88–4.82 for pressure and 2.08–3.08 for flow. Effective rank is exp[–∑_*j*_ *w*_*j*_ log *w*_*j*_], with *w*_*j*_ the normalized singular values, not a count of causal factors. Nonzero coordinate activity and an algebraically complete projection do not establish 16 independently informative physical directions. The maximum absolute change under the tested relabeling was 0, within the specified 10^*−*4^ tolerance.

### B.3 Physical balance diagnostics

The residual calculations use the observed time grid and a trapezoidal approximation. They must be compared with the same calculation on the reference trajectory; neither a nonzero residual nor a small residual by itself establishes a continuous-time physical defect or guarantee. Prediction accuracy, numerical finiteness, and physical balance are reported as separate properties.

### B.4 Post hoc storage identity for the fitted mechanism

This algebraic interpretation was added after coefficient selection; it is not a prespecified evaluation endpoint or a new simulation. Write the five fitted coefficients, in the order given in Methods, as (*a, b, c, d, e*). The implemented equations are

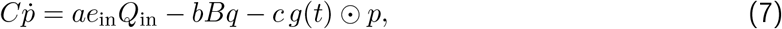

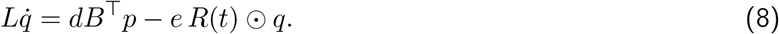

For the selected model all five coefficients are positive, although positivity was not imposed by regression. With fixed positive *C, L*, define *H* = (*dp*^*⊤*^*Cp* + *bq*^*⊤*^*Lq*)*/*2 on active physical coordinates. The coupling terms cancel, giving

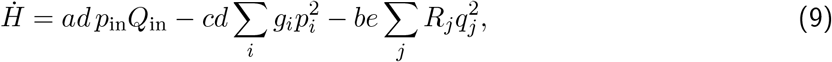

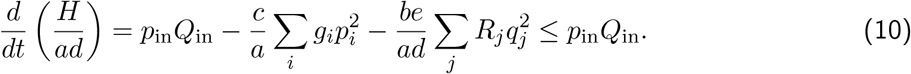

The time-varying positive resistances enter dissipation, not the storage function. This continuous-time inequality follows from the supplied equation family and this particular positive coefficient vector. It does not retain the original unweighted storage, guarantee positivity for future unconstrained fits, bound numerical forecast error, or imply that driven discrete trajectories decrease in energy at every step. No additional model selection or test endpoint used this derivation.

### B.5 Reproducibility and provenance

The local author-review package supplies all 50 final prediction files, comprising two test splits for four neural arms with five seeds each and five deterministic comparators. It also retains the original and amended search outcomes, checkpoint files, generator, analysis scripts, and plotting sources. Identity checks align each prediction with its exact graph, background, action program, mask, and reference trajectory. Canonical floating-point reference states, rather than rounded printed table entries, supply the reported physical-coordinate errors.

Independent analysis recomputed all 50 prediction files from saved physical-coordinate arrays and the original reference states, using separate metric code. It checked graph/background/action identities, all primary and diagnostic errors, physical MAEs, seed summaries, and all six paired stratified-graph contrasts. Agreement was required with absolute tolerance 10^*−*10^ and relative tolerance 5 × 10^*−*12^; nonfinite values had to agree in position rather than be dropped. The reports and exact file hashes are retained in the audit directory.

Complete commands, software versions, resource configurations, units, seeds, and regeneration safeguards appear in the packaged `reproducibility_README.txt`. The full-author reference audit identifies two team-overlapping works among 11 references: JEPA-Anything supplies necessary method attribution, and the IABP study supplies clinical context only. No transcriptomic team paper is added merely to increase citations. This package remains local pending author review; it is not described as an already deposited public dataset.

