## Supplementary Data - Compact audit, summary tables, code, and figure sources for "Panvascular-JEPA: Unseen-Structure Evaluation of Factorized Predictive States for Hemodynamic Networks": final_scientific_review.txt

Independent final scientific review -- 2026-10-11

Disposition
The implemented study supports a distinct, narrowly framed computational benchmark and negative-methodological report for author review. It does not support a newly successful panvascular world model, a new OPF algorithm, a clinically validated predictor, or a claim to have tested the best achievable JEPA or graph-neural simulator. Independence of data and analysis is necessary but not sufficient for publication worthiness; journal suitability and scientific impact remain editorial judgments. Submission remains on hold under the user's explicit instruction.

Independence from related work
The new experiment creates 38 non-isomorphic six-node circuit structures, newly sampled parameter backgrounds, actions and reference trajectories. Fourteen entire unrooted isomorphism classes are withheld. These trajectories and this protected structural split differ from the earlier one-bifurcation pilot and from the team's GEO expression studies. The manuscript discloses those relationships. It does not reuse those papers' predictive outcomes or claim that their clinical/transcriptomic findings validate the present circuit. OPF comes from cited prior work; the new contribution is the controlled experimental comparison, protected graph evaluation, mechanistic controls and complete negative-results reporting.

Reproducibility and arithmetic
audit/confirmation_independent_recomputation.json reports PASS for all 50 saved prediction files and 3,849 comparisons. Separate metric code checked physical-coordinate predictions against canonical reference truth, exact sample identities, active-coordinate masks, training-only scales, pressure/flow-balanced background RMSE, equal graph weighting, five-seed summaries, physical MAE, diagnostic metrics and six stratified graph-bootstrap contrasts. Maximum scaled numerical difference was 5.81992805563247e-16. Truth export discrepancy was zero. This confirms arithmetic and archived input consistency, not biological validity or universal correctness of every modeling assumption.

Nonfinite predictions are retained. The comparison adapter requires identical finite/nonfinite positions, accommodating frozen JSON null serialization of either NaN or infinity; it never filters failed runs from any aggregate. Diagnostic forecasts have genuine nonfinite outcomes even though every primary neural rollout is finite.

Protocol chronology
Separate local lock and dataset-segregation reports verify development, version-2 amendment and final configuration/analysis locks, plus disjoint background identities and non-isomorphic withheld structures. All six hyperparameter candidates per neural arm were rerun after the pre-test amendment, rather than extending only favorable candidates. The 24 original 1,000-update outcomes remain archived. The first 1,000 tensors of every amended run reproduce their original counterpart exactly. All 20 final fits use fixed seeds and 12,000 updates. No test-informed tuning or exclusion was authorized or performed. These are locally recorded hashes and timestamps, not external preregistration.

Primary result and its interpretation
The independently recomputed unseen-structure OPF-4 minus OPF-1 difference is 8,455,843,558.773528 standardized RMSE units; its prespecified E-stratified paired graph percentile interval is [223,137.70418444107, 25,366,824,665.73234]. The positive direction is present in all 14 graph means and all five paired seed means. However, atlas_92 contributes approximately 99.9961% of the summed per-graph difference, and OPF-4 seed 229 contributes approximately 99.9970% of the sum of its five seed RMSEs. These are explicitly post hoc descriptive observations about the already reported results, not replacement endpoints or additional hypothesis tests. The prespecified mean and interval are retained without exclusions. Their huge magnitude is not a typical graph effect. The bootstrap is conditional on the 14 graphs, sampled backgrounds and fixed training seeds; it is not a clinical-population or training-seed population interval.

Every individual primary neural seed on withheld structures is worse than persistence (persistence RMSE 0.28451366091784375; best neural seed, OPF-1 seed 229, 16.3166011614). Neural models also fail badly on seen structures. Consequently, these results do not isolate structural novelty as the source of poor forecasting. Small one-step and reconstruction errors, active latent coordinates, full retained projection rank, and finite numbers do not establish stable or useful autonomous prediction.

Known-equation midpoint integration is accurate, as expected given correct equations, all parameters, initial states and future controls. The fitted model also has the correct equation library by construction. Neither comparator demonstrates discovery of unknown vascular mechanisms. The post hoc modified-storage derivation for the fitted positive coefficients is algebraically valid in continuous time; positivity was not constrained during fitting, the storage is not the original unweighted circuit energy, and the derivation does not establish per-step decrease under forcing or accuracy of every discrete integrator.

Architecture and comparison limitations retained in the manuscript
1. Controls and message context are local; the immediate response to distant inputs is restricted.
2. Predictors use one-step losses and are recursively iterated for 96 steps. The study does not reproduce published graph-surrogate multistep/noise/boundary-connection designs or JEPA-Anything's domain-specific backbones and full objectives.
3. Online-encoder states are used as predictor inputs during training, EMA states as targets, and predicted target-space states as recursive inputs after the first step. EMA does not force coordinate identity. Encoder lag, off-distribution recursive inputs and lack of multistep training are possible concurrent contributors, not proven isolated causes.
4. Equal candidate counts, update budgets and approximately matched parameter counts are documented. They do not imply converged models, optimal hyperparameters, equal FLOPs or an exhaustive best-achievable comparison. The development curves document severe instability rather than convergence.
5. Four-factor versus single-factor OPF is the primary contrast; the identity-basis monolithic objective has a constant target-activity term and is secondary. This difference is disclosed.
6. The catalogue consists of small dimensioned but uncalibrated engineering circuits, with two fixed perturbation programs. It does not demonstrate arterial anatomy transfer, size extrapolation, unseen-action generalization, cell biology, healing, device efficacy or clinical benefit.

Manuscript review and changes
Actual numeric Results, Discussion, Supplement, Abstract result and Conclusion fragments were reviewed against frozen outputs. No result table, graph coordinates, frozen source or primary manuscript file was edited by this audit. The following prose-only corrections were saved:
- Abstract now states that neural errors depend strongly on seed and structure, with extreme forecasts dominating the means.
- Results distinguishes a fixed set of training seeds from a prediction ensemble and explicitly describes atlas_92/seed-229 magnitude dominance without changing the estimator.
- A figure caption now says representation/rollout diagnostics, avoiding causal error-source wording.
- Conclusion directly reports failure to improve the prespecified comparison and failure to outperform persistence.
These fragments require parent assembly and visual/LaTeX QA; this report does not certify final PDF layout.

Authorship, declarations and references
The assembled bibliography contains exactly the 11 planned entries checked against independently obtained full author bylines. Team overlap occurs in two necessary references: JEPA-Anything (method source) and the Biomarker Research IABP study (user-required clinical context). The latter is not used as data or validation. Full-byline evidence and rationales are saved in audit/final_full_author_self_citation_audit.json. The exact user-required AI declaration is present in the assembled source. This check does not independently certify author approval, funding applicability or institutional affiliation.

Research-worthiness boundary
This revision is substantively independent enough to be considered as a reproducible computational stress test with negative outcomes, provided the title, abstract and discussion keep the above limits. It should not be promoted as an improved clinical AI system. A stronger model-development paper would need a separately specified, defensible training/stability intervention and new protected confirmation data after the present test has been inspected. Retuning on the revealed catalogue and relabeling the same test as untouched would invalidate that distinction. Such future work is outside the current locked experiment and must not silently replace the present results.

Final state: experiments complete, independent recomputation passed, manuscript prose reviewed, pending parent assembly/layout and author review. No submission authorized by this audit.
