## Supplementary Data - Compact audit, summary tables, code, and figure sources for "Panvascular-JEPA: Unseen-Structure Evaluation of Factorized Predictive States for Hemodynamic Networks": budget_v2.py

```
"""Audited development-budget amendment; original v1 implementation unchanged.

Every arm receives all six original LR/weight-decay combinations for 6,000
updates. Only the last checkpoint selects hyperparameters. Final fitting uses
12,000 fixed updates. Intermediate validation diagnostics are descriptive only.
"""
from __future__ import annotations
import argparse
import hashlib
import json
import pickle
import random
import time
from pathlib import Path
import numpy as np
import torch
from graph_models import GraphModel
from train_graph_models import (ROOT, Data, CANDIDATES, FINAL_SEEDS, seed_all, save, sha,
                                fit_readout, diagnostics, evaluate, clean)


def source_hashes():
    return {n: sha(ROOT / n) for n in ["graph_models.py", "train_graph_models.py", "budget_v2.py"]}


def rng_hash():
    parts = [torch.get_rng_state().numpy().tobytes(), pickle.dumps(np.random.get_state()),
             pickle.dumps(random.getstate())]
    if torch.cuda.is_available():
        parts.extend(state.cpu().numpy().tobytes() for state in torch.cuda.get_rng_state_all())
    return hashlib.sha256(b"".join(parts)).hexdigest()


def state_hash(value):
    digest = hashlib.sha256()
    def visit(x):
        if isinstance(x, torch.Tensor):
            y = x.detach().cpu().numpy()
            digest.update(str(y.dtype).encode()); digest.update(str(y.shape).encode()); digest.update(y.tobytes())
        elif isinstance(x, dict):
            for k, v in x.items(): digest.update(repr(k).encode()); visit(v)
        elif isinstance(x, (tuple, list)):
            for v in x: visit(v)
        else:
            digest.update(repr(x).encode())
    visit(value)
    return digest.hexdigest()


def fit(name, candidate, seed, steps, milestones, data, output, device,
        old_checkpoint=None, batch=128):
    output = Path(output); output.mkdir(parents=True, exist_ok=True)
    seed_all(seed)
    model = GraphModel(name).to(device)
    opt = torch.optim.AdamW(model.parameters(), lr=candidate["learning_rate"], weight_decay=candidate["weight_decay"])
    t = data.tensors(device)
    train_rows = torch.as_tensor(np.flatnonzero(data.split == "train"), device=device)
    curve = []; fail = None; completed = 0; checkpoints = []; training_time = 0.
    if device == "cuda": torch.cuda.synchronize(); torch.cuda.reset_peak_memory_stats()
    segment_start = time.perf_counter()
    weights = None
    equivalence = None
    for step in range(steps):
        rr = train_rows[torch.randint(len(train_rows), (batch,), device=device)]
        tt = torch.randint(data.nt, (batch,), device=device)
        opt.zero_grad(set_to_none=True)
        loss, parts = model.objective(t["s"][rr, tt], t["s"][rr, tt + 1], t["adj"][rr],
                                      t["etype"][rr], t["mask"][rr], t["static"][rr], t["ctrl"][rr, tt])
        if not torch.isfinite(loss):
            fail = f"Nonfinite training loss at update {step + 1}"; break
        loss.backward()
        norm = torch.nn.utils.clip_grad_norm_(model.parameters(), 5.0)
        if not torch.isfinite(norm):
            fail = f"Nonfinite gradient norm at update {step + 1}"; break
        opt.step(); model.update_target(); completed += 1
        if (step + 1) % 500 == 0 or step == 0 or step + 1 == steps:
            curve.append({"update": step + 1, "loss": float(loss.detach().cpu()),
                          "gradient_norm_pre_clip": float(norm.detach().cpu()),
                          **{k: float(v.cpu()) for k, v in parts.items()}})
        if step + 1 == 1000 and old_checkpoint is not None:
            old = torch.load(old_checkpoint, map_location="cpu", weights_only=False)["state_dict"]
            differences = {k: float((v.detach().cpu() - old[k]).abs().max()) for k, v in model.state_dict().items()}
            equivalence = {"old_checkpoint": str(old_checkpoint), "old_checkpoint_sha256": sha(old_checkpoint),
                           "max_abs_difference": max(differences.values()), "per_tensor_difference": differences}
            save(output / "v1_update1000_equivalence.json", equivalence)
            if equivalence["max_abs_difference"] != 0:
                raise AssertionError("v2 training semantics differ from original first 1,000 updates")
        if step + 1 in milestones:
            if device == "cuda": torch.cuda.synchronize()
            training_time += time.perf_counter() - segment_start
            rng_before = rng_hash()
            model_before = state_hash(model.state_dict())
            optimizer_before = state_hash(opt.state_dict())
            checkpoint = {"update": step + 1, "readout": None}
            if name != "supervised_graph":
                try:
                    weights, checkpoint["readout"] = fit_readout(model, data, device)
                except np.linalg.LinAlgError as exc:
                    fail = "Readout linear algebra failed: " + str(exc)
                    checkpoint["failure"] = fail
            if fail is None:
                checkpoint["diagnostics"] = diagnostics(model, data, device)
                pred_path = output / "validation_predictions.npz" if step + 1 == steps else None
                checkpoint["validation"] = evaluate(model, weights, data, "val", device, pred_path)
            checkpoint["evaluation_state_audit"] = {
                "rng_unchanged": rng_before == rng_hash(),
                "model_unchanged": model_before == state_hash(model.state_dict()),
                "optimizer_unchanged": optimizer_before == state_hash(opt.state_dict())}
            if not all(checkpoint["evaluation_state_audit"].values()):
                raise AssertionError("Intermediate validation mutated model, optimizer, or RNG")
            save(output / f"validation_at_{step + 1}.json", checkpoint)
            torch.save({"name": name, "state_dict": model.state_dict(), "readout": weights,
                        "optimizer_state": opt.state_dict(), "update": step + 1}, output / f"checkpoint_{step + 1}.pt")
            checkpoints.append(checkpoint)
            if device == "cuda": torch.cuda.synchronize()
            segment_start = time.perf_counter()
            if fail: break
    if device == "cuda": torch.cuda.synchronize()
    training_time += time.perf_counter() - segment_start
    report = {"name": name, "candidate": candidate, "seed": seed, "planned_updates": steps,
              "completed_updates": completed, "capacity": model.capacity(), "training_seconds": training_time,
              "training_curve": curve, "validation_milestones": list(milestones), "failure": fail,
              "data_sha256": data.hash, "source_hashes": source_hashes(),
              "peak_cuda_memory_bytes": torch.cuda.max_memory_allocated() if device == "cuda" else 0,
              "batch_size": batch, "gradient_clip_norm": 5., "ema_momentum": .99,
              "orthogonality_weight": .1, "target_activity_weight": .05, "online_activity_weight": .02,
              "v1_first1000_equivalence": equivalence,
              "checkpoint_selection": "Final fixed update only; intermediate validation never selects checkpoint"}
    if checkpoints:
        report["readout"] = checkpoints[-1].get("readout")
        report["diagnostics"] = checkpoints[-1].get("diagnostics")
    torch.save({"state_dict": model.state_dict(), "name": name, "readout": weights}, output / "model.pt")
    save(output / "training.json", report)
    save(output / "scalers.json", data.scaler)
    return model, weights, report, checkpoints


def merge(out):
    out = Path(out)
    records = [json.loads((out / name / "arm_selection.json").read_text(encoding="utf-8")) for name in GraphModel.ARMS]
    selection = {"version": "budget-v2", "development_sha256": records[0]["development_sha256"],
                 "source_hashes": records[0]["source_hashes"], "selection_seed": 101,
                 "selection_metric": "96-step pure-latent rollout equal-structure macro-RMSE at fixed final update6000",
                 "updates_per_candidate": 6000, "candidates_per_arm": 6,
                 "final_updates": 12000, "final_seeds": FINAL_SEEDS, "arms": {}}
    for expected_name, record in zip(GraphModel.ARMS, records):
        if record["name"] != expected_name:
            raise ValueError("Arm selection record is stored under the wrong arm")
        if record["source_hashes"] != selection["source_hashes"] or record["development_sha256"] != selection["development_sha256"]:
            raise ValueError("Search arms used different locked code or development data")
        if record["source_hashes"] != source_hashes():
            raise ValueError("Current code differs from the code used by a search arm")
        if record["updates_per_candidate"] != 6000 or record["milestones"] != [2000, 4000, 6000] or record["seed"] != 101:
            raise ValueError("Search budget/diagnostic schedule/seed differs between arms")
        if len(record["all_candidates"]) != 6 or sorted(c["candidate_index"] for c in record["all_candidates"]) != list(range(6)):
            raise ValueError("Search record lacks the complete six-candidate grid")
        for candidate in record["all_candidates"]:
            expected = CANDIDATES[candidate["candidate_index"]]
            if any(candidate[k] != expected[k] for k in expected):
                raise ValueError("Candidate grid differs from the protocol")
            score = candidate["validation_macro_rmse"]
            if candidate["finite"] and (score is None or not np.isfinite(score)):
                raise ValueError("Finite candidate contains a missing/nonfinite score")
            training = json.loads((out / expected_name / f"candidate_{candidate['candidate_index']}" / "training.json").read_text(encoding="utf-8"))
            if training["planned_updates"] != 6000 or (training["failure"] is None and training["completed_updates"] != 6000):
                raise ValueError("Candidate did not receive its prescribed training budget")
            if training["source_hashes"] != source_hashes() or training["data_sha256"] != selection["development_sha256"]:
                raise ValueError("Candidate training provenance differs from selection record")
        finite = [c for c in record["all_candidates"] if c["finite"]]
        if not finite or record["winner"] != min(finite, key=lambda c: c["validation_macro_rmse"]):
            raise ValueError("Stored winner is not the final-checkpoint validation minimum")
        selection["arms"][record["name"]] = {"all_candidates": record["all_candidates"], "winner": record["winner"]}
    if set(selection["arms"]) != set(GraphModel.ARMS):
        raise ValueError("Merged search does not contain all four arms")
    save(out / "selected_hyperparameters.json", selection)
    (out / "selected_hyperparameters.sha256").write_text(sha(out / "selected_hyperparameters.json"), encoding="utf-8")


def main():
    parser = argparse.ArgumentParser()
    parser.add_argument("mode", choices=["search", "merge", "confirm"])
    parser.add_argument("--development")
    parser.add_argument("--confirmation")
    parser.add_argument("--output", required=True)
    parser.add_argument("--selection")
    parser.add_argument("--arm", choices=GraphModel.ARMS)
    parser.add_argument("--seed", type=int)
    parser.add_argument("--old-search", default=str(ROOT / "search"))
    parser.add_argument("--device", choices=["cuda", "cpu"], default="cuda")
    args = parser.parse_args()
    if args.mode == "merge": merge(args.output); return
    if not args.arm: raise ValueError("Run one arm per process for bounded concurrency")
    torch.set_num_threads(2); torch.set_num_interop_threads(2)
    if args.device == "cuda": torch.cuda.set_per_process_memory_fraction(.055)
    data = Data(args.development)
    if set(data.split) - {"train", "val"}: raise ValueError("Development file includes test states")
    out = Path(args.output); out.mkdir(parents=True, exist_ok=True)
    if args.mode == "search":
        if args.confirmation: raise ValueError("Search never accepts confirmation data")
        results = []
        for ci, candidate in enumerate(CANDIDATES):
            old = Path(args.old_search) / args.arm / f"candidate_{ci}" / "model.pt"
            od = out / args.arm / f"candidate_{ci}"
            model, weights, report, checkpoints = fit(args.arm, candidate, 101, 6000, [2000, 4000, 6000],
                  data, od, args.device, old_checkpoint=old)
            val = None if report["failure"] else checkpoints[-1]["validation"]
            score = np.inf if val is None else val["pure_latent_rollout"]["macro_rmse"]
            if not np.isfinite(score): score = np.inf
            results.append({"candidate_index": ci, **candidate, "validation_macro_rmse": float(score),
                            "finite": bool(np.isfinite(score)), "training_failure": report["failure"]})
            save(out / args.arm / "search_progress.json", results)
            print(json.dumps({"arm": args.arm, "candidate": ci, "score": clean(float(score))}), flush=True)
            del model
        finite = [x for x in results if x["finite"]]
        if not finite: raise RuntimeError("Every prespecified long-budget candidate failed; retain failure, do not tune on test")
        save(out / args.arm / "arm_selection.json", {"name": args.arm, "development_sha256": data.hash,
             "source_hashes": source_hashes(), "updates_per_candidate": 6000,
             "milestones": [2000, 4000, 6000], "seed": 101, "all_candidates": results,
             "winner": min(finite, key=lambda x: x["validation_macro_rmse"])})
    else:
        if not args.selection or not args.confirmation: raise ValueError("Frozen selection and confirmation file required")
        selected = json.loads(Path(args.selection).read_text(encoding="utf-8"))
        if Path(args.selection).with_suffix(".sha256").read_text(encoding="utf-8").strip() != sha(args.selection):
            raise ValueError("Selected hyperparameter file does not match its stored hash")
        if selected["development_sha256"] != data.hash or selected["source_hashes"] != source_hashes():
            raise ValueError("Data/code changed after selection lock")
        test = Data(args.confirmation, data.scaler)
        if set(test.split) & {"train", "val"} or set(data.bg) & set(test.bg):
            raise ValueError("Development/confirmation contamination")
        candidate = {k: selected["arms"][args.arm]["winner"][k] for k in ["learning_rate", "weight_decay"]}
        seeds = [args.seed] if args.seed else FINAL_SEEDS
        if not set(seeds).issubset(FINAL_SEEDS): raise ValueError("Seed is not in the frozen confirmation list")
        for seed in seeds:
            od = out / args.arm / f"seed_{seed}"
            model, weights, report, checkpoints = fit(args.arm, candidate, seed, 12000, [4000, 8000, 12000], data, od, args.device)
            result = {"selection_sha256": sha(args.selection), "confirmation_sha256": test.hash,
                      "training": report, "splits": {}}
            if report["failure"] is None:
                for split in sorted(set(test.split)):
                    result["splits"][split] = evaluate(model, weights, test, split, args.device, od / f"{split}_predictions.npz")
            save(od / "metrics.json", result)
            print(json.dumps({"arm": args.arm, "seed": seed,
                              "scores": {k: clean(v["pure_latent_rollout"]["macro_rmse"]) for k, v in result["splits"].items()}}), flush=True)
            del model


if __name__ == "__main__": main()
```
