## Supplementary Data - Compact audit, summary tables, code, and figure sources for "Panvascular-JEPA: Unseen-Structure Evaluation of Factorized Predictive States for Hemodynamic Networks": build_numeric_tables.py

```
"""Format completed, locked outcomes for author review; no outcome selection."""
from pathlib import Path
import csv, json, math
import numpy as np

ROOT = Path(__file__).resolve().parent
ARMS = ['mono_jepa','opf1','opf4','supervised_graph']
METHODS = ARMS + ['rlc_midpoint_1','rlc_midpoint_4','rc_reduced_4','structured_physics','persistence']
NAMES = {'mono_jepa':'Monolithic JEPA','opf1':'OPF-1','opf4':'OPF-4','supervised_graph':'Supervised graph',
         'rlc_midpoint_1':'Known RLC, 1 substep','rlc_midpoint_4':'Known RLC, 4 substeps',
         'rc_reduced_4':'Reduced RC','structured_physics':'Fitted RLC','persistence':'Persistence'}


def read(path):
    return json.loads((ROOT/path).read_text(encoding='utf-8'))


def rows(name):
    with (ROOT/'analysis'/name).open(newline='',encoding='utf-8') as f:
        return list(csv.DictReader(f))


def num(value):
    if value is None or value == '':
        return r'\mathrm{NF}'
    value = float(value)
    if not math.isfinite(value):
        return r'\mathrm{NF}'
    if value == 0:
        return '0'
    if 0.001 <= abs(value) < 1000:
        return f'{value:.3g}'
    mantissa, exponent = f'{value:.2e}'.split('e')
    return mantissa + r'\times10^{' + str(int(exponent)) + '}'


def mathnum(value):
    return '$' + num(value) + '$'


def table(caption,label,header,body,spec,note,placement='!tp'):
    return '\n'.join([r'\begin{table}['+placement+']',r'\centering\small',
        r'\setlength{\tabcolsep}{4pt}',r'\renewcommand{\arraystretch}{1.15}',
        '\\caption{'+caption+'}', '\\label{'+label+'}',
        '\\begin{tabularx}{\\linewidth}{'+spec+'}',r'\toprule',header+r'\\',r'\midrule',
        *[r+r'\\' for r in body],r'\bottomrule',r'\end{tabularx}',
        '\\par\\vspace{4pt}\\begin{minipage}{\\linewidth}\\footnotesize '+note+r'\end{minipage}',
        r'\end{table}'])


def main():
    summary=read('analysis/summary.json')
    tab={(r['method'],r['split']):r for r in summary['table']}
    body=[]
    for method in METHODS:
        values=[]
        for split in ['id_test','ood_topology']:
            r=tab[method,split]
            v=num(r['rmse_mean'])
            if method in ARMS:
                v+=r'\pm '+num(r['rmse_seed_sd'])
            values.append('$'+v+'$')
        body.append(NAMES[method]+' & '+' & '.join(values))
    primary=table('Autonomous 96-step prediction after frozen model selection.', 'tab:primary',
        'Method & Seen structures & Unseen structures',body,'Yrr',
        r'Errors are standardized, equal-type RMSE, averaged over paired action programs within backgrounds, then equally over backgrounds within structures and over structures. Neural entries are mean $\pm$ sample standard deviation across all five fixed training seeds; deterministic entries are single results. Seen testing contains 24 structures and 96 new backgrounds; unseen testing contains 14 structures and 168 backgrounds. NF denotes a nonfinite aggregate; failed seeds are not omitted.')
    (ROOT/'manuscript_table_primary.txt').write_text(primary,encoding='utf-8')
    body=[]
    for method in METHODS:
        r=tab[method,'ood_topology']
        body.append(NAMES[method]+' & '+mathnum(r['pressure_mae_mmHg'])+' & '+mathnum(r['flow_mae_ml_s'])+' & '+str(r['nonfinite_active_predictions']))
    physical=table('Physical-coordinate errors on the 14 unseen structures.','tab:physical',
        'Method & Pressure MAE & Flow MAE & Nonfinite',body,'Yrrr',
        r'Pressure is in mmHg and flow in mL/s. MAEs use background-then-structure averaging and, for neural methods, the mean of all five seeds. The last column counts nonfinite active scalar predictions across those seeds; a zero count does not imply that a very large finite forecast is stable or useful.')
    (ROOT/'manuscript_table_physical.txt').write_text(physical,encoding='utf-8')
    selected=read('search_long/selected_hyperparameters.json')
    body=[]
    for ci in range(6):
        c=selected['arms'][ARMS[0]]['all_candidates'][ci]
        scores=[]
        for arm in ARMS:
            d=selected['arms'][arm]['all_candidates'][ci]
            value=mathnum(d['validation_macro_rmse'])
            if selected['arms'][arm]['winner']['candidate_index']==ci:
                value=r'\textbf{'+value+r'}$^{*}$'
            scores.append(value)
        body.append(str(ci+1)+' & '+mathnum(c['learning_rate'])+' & '+mathnum(c['weight_decay'])+' & '+' & '.join(scores))
    search=table('Complete six-candidate search at the fixed 6,000-update checkpoint.','tab:selection',
        'ID & LR & WD & Mono & OPF-1 & OPF-4 & Supervised',body,'rYYrrrr',
        r'LR: learning rate; WD: weight decay. Entries are validation 96-step structure-macro RMSE at seed 101; asterisks mark the prespecified minimum-score selection. No earlier checkpoint was substituted. Values are rounded for display; full precision and every checkpoint are in the archived selection files.',placement='!htbp')
    (ROOT/'manuscript_table_selection.txt').write_text(search,encoding='utf-8')
    run=rows('run_metrics.csv')
    body=[]
    for method in ARMS:
        for seed in [211,223,227,229,233]:
            d={r['diagnostic']:r for r in run if r['method']==method and r['split']=='ood_topology' and int(r['seed'])==seed}
            body.append(NAMES[method]+' & '+str(seed)+' & '+' & '.join(mathnum(d[k]['rmse']) for k in ['reconstruction','one_step','rollout','decode_reencode']))
    diag=table('Every fixed-seed diagnostic on unseen structures.','tab:all-seeds',
        'Method & Seed & Recon. & One step & Rollout & Re-encode',body,'Yrrrrr',
        r"Reconstruction and one-step predictions use true observations and are diagnostic lower-complexity tasks, not autonomous forecasts. Re-encoding uses the model's decoded state, with no future truth. All entries use the primary standardized aggregation. NF denotes a nonfinite aggregate; affected runs are retained. The supervised reconstruction column is an identity reference and its re-encoding column duplicates its physical-state rollout; these are not separate learned capabilities.")
    (ROOT/'manuscript_table_all_seeds.txt').write_text(diag,encoding='utf-8')
    residuals=rows('residual_metrics.csv')
    body=[]
    for method in METHODS+['reference']:
        rr=[r for r in residuals if r['method']==method and r['split']=='ood_topology']
        assert len(rr)==(70 if method in ARMS else 14)
        # Equal number of structures for each seed; never drop a nonfinite row.
        mass=np.mean([float(r['mass_rmse_ml_s']) if r['mass_rmse_ml_s'] else np.nan for r in rr])
        momentum=np.mean([float(r['momentum_rmse_mmHg']) if r['momentum_rmse_mmHg'] else np.nan for r in rr])
        body.append(NAMES.get(method,'DOP853 reference')+' & '+mathnum(mass)+' & '+mathnum(momentum))
    residual_table=table('Discrete balance residuals on unseen structures.','tab:balance',
        'Method & Mass balance & Momentum balance',body,'Yrr',
        r'Mass-balance residual is in mL/s and momentum-balance residual in mmHg. Trapezoidal differences at the observation interval have a discretization floor, including for the DOP853 reference. Entries average trajectory RMS residuals within each graph and then equally across graphs and fixed seeds. They do not certify continuous-time conservation or clinical plausibility.')
    (ROOT/'manuscript_table_balance.txt').write_text(residual_table,encoding='utf-8')
    print('Wrote five numerical tables from completed analysis; no data/model changes.')


if __name__=='__main__':
    main()
```
