## Supplementary Data - Compact audit, summary tables, code, and figure sources for "Panvascular-JEPA: Unseen-Structure Evaluation of Factorized Predictive States for Hemodynamic Networks": collect_review_metrics.py

```
"""Descriptive reporting only: never changes selection, predictions, or locked analysis."""
from pathlib import Path
import csv, json
import numpy as np

ROOT = Path(__file__).resolve().parent
ARMS = ['mono_jepa', 'opf1', 'opf4', 'supervised_graph']
SEEDS = [211, 223, 227, 229, 233]


def read(path):
    return json.loads(path.read_text(encoding='utf-8'))


def csvwrite(path, rows):
    with path.open('w', newline='', encoding='utf-8') as f:
        w = csv.DictWriter(f, fieldnames=list(rows[0]))
        w.writeheader(); w.writerows(rows)


def valrow(arm, stage, candidate, seed, update, validation, status='observed_checkpoint'):
    fields = {'arm': arm, 'stage': stage, 'candidate': candidate,
              'seed': seed, 'update': update, 'checkpoint_status': status}
    for key in ['pure_latent_rollout', 'one_step', 'truth_encoding_readout', 'decode_reencode_rollout']:
        val = validation.get(key, {})
        fields[key + '_rmse'] = val.get('macro_rmse')
        fields[key + '_nonfinite_count'] = val.get('nonfinite_count')
    return fields


def main():
    if not (ROOT / 'analysis/summary.json').exists():
        raise RuntimeError('Reporting waits for complete locked analysis')
    selected = read(ROOT / 'search_long/selected_hyperparameters.json')
    rows = []; diagnostics = []; runtimes = []
    for arm in ARMS:
        for ci in range(6):
            training = read(ROOT / 'search_long' / arm / f'candidate_{ci}' / 'training.json')
            for update in [2000, 4000, 6000]:
                p = ROOT / 'search_long' / arm / f'candidate_{ci}' / f'validation_at_{update}.json'
                if p.exists():
                    d = read(p); status = 'observed_checkpoint'
                elif training['failure'] is not None:
                    d = {}; status = 'missing_after_recorded_training_failure'
                else:
                    raise RuntimeError(f'Unexplained missing checkpoint: {p}')
                rows.append(valrow(arm, 'search_v2', ci, 101, update, d.get('validation', {}), status))
        for seed in SEEDS:
            folder = ROOT / 'results' / arm / f'seed_{seed}'
            tr = read(folder / 'training.json')
            runtimes.append({'arm': arm, 'seed': seed, 'training_seconds': tr['training_seconds'],
                             'completed_updates': tr['completed_updates'],
                             'failure': tr['failure'], 'peak_cuda_memory_bytes': tr['peak_cuda_memory_bytes']})
            for update in [4000, 8000, 12000]:
                p = folder / f'validation_at_{update}.json'
                if p.exists():
                    d = read(p); status = 'observed_checkpoint'
                elif tr['failure'] is not None:
                    d = {}; status = 'missing_after_recorded_training_failure'
                else:
                    raise RuntimeError(f'Unexplained missing checkpoint: {p}')
                rows.append(valrow(arm, 'final_fit_validation', selected['arms'][arm]['winner']['candidate_index'],
                                   seed, update, d.get('validation', {}), status))
            d = tr.get('diagnostics') or {}
            b = d.get('basis', {})
            row = {'arm': arm, 'seed': seed,
                   'permutation_encoder_max_abs': d.get('permutation_encoder_max_abs'),
                   'permutation_prediction_max_abs': d.get('permutation_prediction_max_abs'),
                   'basis_condition': b.get('condition'), 'basis_rank': b.get('rank_at_rtol_1e_5'),
                   'basis_gram_loss': b.get('normalized_full_gram_loss')}
            for ty in ['pressure', 'flow']:
                a = d.get(ty + '_ema_latents', {})
                values = a.get('coordinate_std', [])
                valid = [v for v in values if v is not None and np.isfinite(v)]
                row[ty + '_coordinate_std_nonfinite_count'] = len(values) - len(valid)
                row[ty + '_coordinate_std_min'] = min(values) if values and len(valid) == len(values) else None
                row[ty + '_coordinate_std_max'] = max(values) if values and len(valid) == len(values) else None
                row[ty + '_effective_rank'] = a.get('effective_rank')
            diagnostics.append(row)
    csvwrite(ROOT / 'analysis/validation_budget_trajectories.csv', rows)
    csvwrite(ROOT / 'analysis/final_representation_diagnostics.csv', diagnostics)
    csvwrite(ROOT / 'analysis/final_training_resources.csv', runtimes)
    output = {'scope': 'Descriptive report; not a model or checkpoint selection',
              'search_candidates': 24, 'final_fits': 20,
              'search_updates': 6000, 'final_updates': 12000,
              'all_final_completed': all(r['completed_updates'] == 12000 and r['failure'] is None for r in runtimes),
              'basis_diagnostics': diagnostics,
              'all_training_seconds': sum(r['training_seconds'] for r in runtimes)}
    (ROOT / 'analysis/reporting_supplement.json').write_text(json.dumps(output, indent=2, allow_nan=False), encoding='utf-8')
    print(json.dumps({k:v for k,v in output.items() if k != 'basis_diagnostics'}, indent=2))


if __name__ == '__main__':
    main()
```
