## Supplementary Data - Compact audit, summary tables, code, and figure sources for "Panvascular-JEPA: Unseen-Structure Evaluation of Factorized Predictive States for Hemodynamic Networks": budget_sensitivity_figure.tex

\begin{figure}[!tp]
\centering
\begin{checkedfigure}
\begin{tikzpicture}[x=1mm,y=1mm,font=\fontsize{8}{10}\selectfont,text=PVnavy,line cap=round,line join=round]
\path[use as bounding box] (0,0) rectangle (152,180);
\node[font=\fontsize{8.8}{10.5}\selectfont\bfseries] at (44,177) {Search: six candidates};
\node[font=\fontsize{8.8}{10.5}\selectfont\bfseries] at (123,177) {Final fits: five seeds};
\node[anchor=west,font=\fontsize{9}{11}\selectfont\bfseries] at (0.7,168) {A\quad Mono JEPA};
\begin{axis}[at={(18mm,140mm)},anchor=south west,width=54mm,height=24mm,scale only axis,font=\fontsize{8}{10}\selectfont,tick label style={font=\fontsize{8}{9.5}\selectfont},label style={font=\fontsize{8}{10}\selectfont},axis line style={PVgray!65,line width=.45pt},tick style={PVgray!65},grid=major,grid style={PVgray!15,line width=.3pt},enlarge x limits=false,enlarge y limits=false,clip=true,ymode=log,ymin=100,ymax=1e+33,xmin=1.75,xmax=6.25,xtick={2,4,6},xlabel={},ylabel={Validation RMSE}]
\addplot[PVgray!45,dashed,line width=.4pt,mark=none] coordinates {(6,100) (6,1e+33)};
\addplot[color=PVgray,solid,mark=*,line width=.55pt,opacity=.75,mark=none] coordinates {(2,2920995.555) (4,3.641775875e+14)};
\addplot[color=PVgray,solid,mark=*,line width=.55pt,opacity=.75,mark=none] coordinates {(4,3.641775875e+14) (6,3.119042121e+17)};
\addplot[color=PVgray,solid,mark=*,line width=.55pt,opacity=.75,only marks,mark size=1.2pt] coordinates {(2,2920995.555) (4,3.641775875e+14) (6,3.119042121e+17)};
\addplot[color=PVgray,dashed,mark=square*,line width=.55pt,opacity=.75,mark=none] coordinates {(2,2879939.897) (4,3.50786185e+14)};
\addplot[color=PVgray,dashed,mark=square*,line width=.55pt,opacity=.75,mark=none] coordinates {(4,3.50786185e+14) (6,3.054390822e+17)};
\addplot[color=PVgray,dashed,mark=square*,line width=.55pt,opacity=.75,only marks,mark size=1.2pt] coordinates {(2,2879939.897) (4,3.50786185e+14) (6,3.054390822e+17)};
\addplot[color=PVgray,densely dotted,mark=triangle*,line width=.55pt,opacity=.75,mark=none] coordinates {(2,2.768229021e+16) (4,2.90716491e+12)};
\addplot[color=PVgray,densely dotted,mark=triangle*,line width=.55pt,opacity=.75,mark=none] coordinates {(4,2.90716491e+12) (6,1.392204772e+18)};
\addplot[color=PVgray,densely dotted,mark=triangle*,line width=.55pt,opacity=.75,only marks,mark size=1.2pt] coordinates {(2,2.768229021e+16) (4,2.90716491e+12) (6,1.392204772e+18)};
\addplot[color=PVgray,dashdotted,mark=diamond*,line width=.55pt,opacity=.75,mark=none] coordinates {(2,2.76442068e+16) (4,2.78631559e+12)};
\addplot[color=PVgray,dashdotted,mark=diamond*,line width=.55pt,opacity=.75,mark=none] coordinates {(4,2.78631559e+12) (6,1.227001277e+18)};
\addplot[color=PVgray,dashdotted,mark=diamond*,line width=.55pt,opacity=.75,only marks,mark size=1.2pt] coordinates {(2,2.76442068e+16) (4,2.78631559e+12) (6,1.227001277e+18)};
\addplot[color=PVgray,densely dashed,mark=asterisk,line width=.55pt,opacity=.75,mark=none] coordinates {(2,4.266046392e+12) (4,1.040872893e+26)};
\addplot[color=PVgray,densely dashed,mark=asterisk,line width=.55pt,opacity=.75,mark=none] coordinates {(4,1.040872893e+26) (6,1.263253326e+32)};
\addplot[color=PVgray,densely dashed,mark=asterisk,line width=.55pt,opacity=.75,only marks,mark size=1.2pt] coordinates {(2,4.266046392e+12) (4,1.040872893e+26) (6,1.263253326e+32)};
\addplot[color=PVgray,loosely dotted,mark=x,line width=.55pt,opacity=.75,mark=none] coordinates {(2,4.382817622e+12) (4,9.36942579e+25)};
\addplot[color=PVgray,loosely dotted,mark=x,line width=.55pt,opacity=.75,mark=none] coordinates {(4,9.36942579e+25) (6,1.262984456e+32)};
\addplot[color=PVgray,loosely dotted,mark=x,line width=.55pt,opacity=.75,only marks,mark size=1.2pt] coordinates {(2,4.382817622e+12) (4,9.36942579e+25) (6,1.262984456e+32)};
\end{axis}
\begin{axis}[at={(96mm,140mm)},anchor=south west,width=54mm,height=24mm,scale only axis,font=\fontsize{8}{10}\selectfont,tick label style={font=\fontsize{8}{9.5}\selectfont},label style={font=\fontsize{8}{10}\selectfont},axis line style={PVgray!65,line width=.45pt},tick style={PVgray!65},grid=major,grid style={PVgray!15,line width=.3pt},enlarge x limits=false,enlarge y limits=false,clip=true,ymode=log,ymin=100,ymax=1e+33,xmin=3.5,xmax=12.5,xtick={4,8,12},xlabel={},ylabel={}]
\addplot[PVgray!45,dashed,line width=.4pt,mark=none] coordinates {(12,100) (12,1e+33)};
\addplot[color=PVblue,solid,mark=*,line width=.6pt,mark=none] coordinates {(4,22607.88498) (8,7867.584192)};
\addplot[color=PVblue,solid,mark=*,line width=.6pt,mark=none] coordinates {(8,7867.584192) (12,2338.690418)};
\addplot[color=PVblue,solid,mark=*,line width=.6pt,only marks,mark size=1.2pt] coordinates {(4,22607.88498) (8,7867.584192) (12,2338.690418)};
\addplot[color=PVteal,solid,mark=*,line width=.6pt,mark=none] coordinates {(4,19940.47258) (8,77735.351)};
\addplot[color=PVteal,solid,mark=*,line width=.6pt,mark=none] coordinates {(8,77735.351) (12,5280.411392)};
\addplot[color=PVteal,solid,mark=*,line width=.6pt,only marks,mark size=1.2pt] coordinates {(4,19940.47258) (8,77735.351) (12,5280.411392)};
\addplot[color=PVorange,solid,mark=*,line width=.6pt,mark=none] coordinates {(4,172823.2676) (8,1017334.525)};
\addplot[color=PVorange,solid,mark=*,line width=.6pt,mark=none] coordinates {(8,1017334.525) (12,374609.0189)};
\addplot[color=PVorange,solid,mark=*,line width=.6pt,only marks,mark size=1.2pt] coordinates {(4,172823.2676) (8,1017334.525) (12,374609.0189)};
\addplot[color=PVred,solid,mark=*,line width=.6pt,mark=none] coordinates {(4,151609.4337) (8,2.199447819e+11)};
\addplot[color=PVred,solid,mark=*,line width=.6pt,mark=none] coordinates {(8,2.199447819e+11) (12,9.248039608e+12)};
\addplot[color=PVred,solid,mark=*,line width=.6pt,only marks,mark size=1.2pt] coordinates {(4,151609.4337) (8,2.199447819e+11) (12,9.248039608e+12)};
\addplot[color=PVnavy,solid,mark=*,line width=.6pt,mark=none] coordinates {(4,4933.689177) (8,939.6379428)};
\addplot[color=PVnavy,solid,mark=*,line width=.6pt,mark=none] coordinates {(8,939.6379428) (12,608.9251227)};
\addplot[color=PVnavy,solid,mark=*,line width=.6pt,only marks,mark size=1.2pt] coordinates {(4,4933.689177) (8,939.6379428) (12,608.9251227)};
\end{axis}
\node[anchor=west,font=\fontsize{9}{11}\selectfont\bfseries] at (0.7,132) {B\quad OPF ($K=1$)};
\begin{axis}[at={(18mm,104mm)},anchor=south west,width=54mm,height=24mm,scale only axis,font=\fontsize{8}{10}\selectfont,tick label style={font=\fontsize{8}{9.5}\selectfont},label style={font=\fontsize{8}{10}\selectfont},axis line style={PVgray!65,line width=.45pt},tick style={PVgray!65},grid=major,grid style={PVgray!15,line width=.3pt},enlarge x limits=false,enlarge y limits=false,clip=true,ymode=log,ymin=10,ymax=1e+13,xmin=1.75,xmax=6.25,xtick={2,4,6},xlabel={},ylabel={Validation RMSE}]
\addplot[PVgray!45,dashed,line width=.4pt,mark=none] coordinates {(6,10) (6,1e+13)};
\addplot[color=PVgray,solid,mark=*,line width=.55pt,opacity=.75,mark=none] coordinates {(2,306624.7365) (4,792172900.9)};
\addplot[color=PVgray,solid,mark=*,line width=.55pt,opacity=.75,mark=none] coordinates {(4,792172900.9) (6,170157075.2)};
\addplot[color=PVgray,solid,mark=*,line width=.55pt,opacity=.75,only marks,mark size=1.2pt] coordinates {(2,306624.7365) (4,792172900.9) (6,170157075.2)};
\addplot[color=PVgray,dashed,mark=square*,line width=.55pt,opacity=.75,mark=none] coordinates {(2,298953.055) (4,761285191.2)};
\addplot[color=PVgray,dashed,mark=square*,line width=.55pt,opacity=.75,mark=none] coordinates {(4,761285191.2) (6,200201905.3)};
\addplot[color=PVgray,dashed,mark=square*,line width=.55pt,opacity=.75,only marks,mark size=1.2pt] coordinates {(2,298953.055) (4,761285191.2) (6,200201905.3)};
\addplot[color=PVgray,densely dotted,mark=triangle*,line width=.55pt,opacity=.75,mark=none] coordinates {(2,149725820.3) (4,6790368701)};
\addplot[color=PVgray,densely dotted,mark=triangle*,line width=.55pt,opacity=.75,mark=none] coordinates {(4,6790368701) (6,7.533648845e+11)};
\addplot[color=PVgray,densely dotted,mark=triangle*,line width=.55pt,opacity=.75,only marks,mark size=1.2pt] coordinates {(2,149725820.3) (4,6790368701) (6,7.533648845e+11)};
\addplot[color=PVgray,dashdotted,mark=diamond*,line width=.55pt,opacity=.75,mark=none] coordinates {(2,165286758.6) (4,6298359734)};
\addplot[color=PVgray,dashdotted,mark=diamond*,line width=.55pt,opacity=.75,mark=none] coordinates {(4,6298359734) (6,8.417041133e+11)};
\addplot[color=PVgray,dashdotted,mark=diamond*,line width=.55pt,opacity=.75,only marks,mark size=1.2pt] coordinates {(2,165286758.6) (4,6298359734) (6,8.417041133e+11)};
\addplot[color=PVgray,densely dashed,mark=asterisk,line width=.55pt,opacity=.75,mark=none] coordinates {(2,6157164501) (4,7.440463163e+10)};
\addplot[color=PVgray,densely dashed,mark=asterisk,line width=.55pt,opacity=.75,mark=none] coordinates {(4,7.440463163e+10) (6,222864991.4)};
\addplot[color=PVgray,densely dashed,mark=asterisk,line width=.55pt,opacity=.75,only marks,mark size=1.2pt] coordinates {(2,6157164501) (4,7.440463163e+10) (6,222864991.4)};
\addplot[color=PVgray,loosely dotted,mark=x,line width=.55pt,opacity=.75,mark=none] coordinates {(2,8916806062) (4,1.045116354e+11)};
\addplot[color=PVgray,loosely dotted,mark=x,line width=.55pt,opacity=.75,mark=none] coordinates {(4,1.045116354e+11) (6,138445639.8)};
\addplot[color=PVgray,loosely dotted,mark=x,line width=.55pt,opacity=.75,only marks,mark size=1.2pt] coordinates {(2,8916806062) (4,1.045116354e+11) (6,138445639.8)};
\end{axis}
\begin{axis}[at={(96mm,104mm)},anchor=south west,width=54mm,height=24mm,scale only axis,font=\fontsize{8}{10}\selectfont,tick label style={font=\fontsize{8}{9.5}\selectfont},label style={font=\fontsize{8}{10}\selectfont},axis line style={PVgray!65,line width=.45pt},tick style={PVgray!65},grid=major,grid style={PVgray!15,line width=.3pt},enlarge x limits=false,enlarge y limits=false,clip=true,ymode=log,ymin=10,ymax=1e+13,xmin=3.5,xmax=12.5,xtick={4,8,12},xlabel={},ylabel={}]
\addplot[PVgray!45,dashed,line width=.4pt,mark=none] coordinates {(12,10) (12,1e+13)};
\addplot[color=PVblue,solid,mark=*,line width=.6pt,mark=none] coordinates {(4,130.6613544) (8,134158.2715)};
\addplot[color=PVblue,solid,mark=*,line width=.6pt,mark=none] coordinates {(8,134158.2715) (12,381.3323801)};
\addplot[color=PVblue,solid,mark=*,line width=.6pt,only marks,mark size=1.2pt] coordinates {(4,130.6613544) (8,134158.2715) (12,381.3323801)};
\addplot[color=PVteal,solid,mark=*,line width=.6pt,mark=none] coordinates {(4,243.5688655) (8,41036.61446)};
\addplot[color=PVteal,solid,mark=*,line width=.6pt,mark=none] coordinates {(8,41036.61446) (12,75.83858873)};
\addplot[color=PVteal,solid,mark=*,line width=.6pt,only marks,mark size=1.2pt] coordinates {(4,243.5688655) (8,41036.61446) (12,75.83858873)};
\addplot[color=PVorange,solid,mark=*,line width=.6pt,mark=none] coordinates {(4,1230161.231) (8,16361.94441)};
\addplot[color=PVorange,solid,mark=*,line width=.6pt,mark=none] coordinates {(8,16361.94441) (12,797.6659818)};
\addplot[color=PVorange,solid,mark=*,line width=.6pt,only marks,mark size=1.2pt] coordinates {(4,1230161.231) (8,16361.94441) (12,797.6659818)};
\addplot[color=PVred,solid,mark=*,line width=.6pt,mark=none] coordinates {(4,1111882.439) (8,78.05869569)};
\addplot[color=PVred,solid,mark=*,line width=.6pt,mark=none] coordinates {(8,78.05869569) (12,17.63159913)};
\addplot[color=PVred,solid,mark=*,line width=.6pt,only marks,mark size=1.2pt] coordinates {(4,1111882.439) (8,78.05869569) (12,17.63159913)};
\addplot[color=PVnavy,solid,mark=*,line width=.6pt,mark=none] coordinates {(4,29606.88307) (8,1858.789793)};
\addplot[color=PVnavy,solid,mark=*,line width=.6pt,mark=none] coordinates {(8,1858.789793) (12,62.34817679)};
\addplot[color=PVnavy,solid,mark=*,line width=.6pt,only marks,mark size=1.2pt] coordinates {(4,29606.88307) (8,1858.789793) (12,62.34817679)};
\end{axis}
\node[anchor=west,font=\fontsize{9}{11}\selectfont\bfseries] at (0.7,96) {C\quad OPF ($K=4$)};
\begin{axis}[at={(18mm,68mm)},anchor=south west,width=54mm,height=24mm,scale only axis,font=\fontsize{8}{10}\selectfont,tick label style={font=\fontsize{8}{9.5}\selectfont},label style={font=\fontsize{8}{10}\selectfont},axis line style={PVgray!65,line width=.45pt},tick style={PVgray!65},grid=major,grid style={PVgray!15,line width=.3pt},enlarge x limits=false,enlarge y limits=false,clip=true,ymode=log,ymin=10000,ymax=1e+16,xmin=1.75,xmax=6.25,xtick={2,4,6},xlabel={},ylabel={Validation RMSE}]
\addplot[PVgray!45,dashed,line width=.4pt,mark=none] coordinates {(6,10000) (6,1e+16)};
\addplot[color=PVgray,solid,mark=*,line width=.55pt,opacity=.75,mark=none] coordinates {(2,1.705308285e+10) (4,1.135950215e+10)};
\addplot[color=PVgray,solid,mark=*,line width=.55pt,opacity=.75,mark=none] coordinates {(4,1.135950215e+10) (6,5.371133589e+11)};
\addplot[color=PVgray,solid,mark=*,line width=.55pt,opacity=.75,only marks,mark size=1.2pt] coordinates {(2,1.705308285e+10) (4,1.135950215e+10) (6,5.371133589e+11)};
\addplot[color=PVgray,dashed,mark=square*,line width=.55pt,opacity=.75,mark=none] coordinates {(2,1.706806516e+10) (4,1.082496557e+10)};
\addplot[color=PVgray,dashed,mark=square*,line width=.55pt,opacity=.75,mark=none] coordinates {(4,1.082496557e+10) (6,5.283575136e+11)};
\addplot[color=PVgray,dashed,mark=square*,line width=.55pt,opacity=.75,only marks,mark size=1.2pt] coordinates {(2,1.706806516e+10) (4,1.082496557e+10) (6,5.283575136e+11)};
\addplot[color=PVgray,densely dotted,mark=triangle*,line width=.55pt,opacity=.75,mark=none] coordinates {(2,614802956.1) (4,102634630.7)};
\addplot[color=PVgray,densely dotted,mark=triangle*,line width=.55pt,opacity=.75,mark=none] coordinates {(4,102634630.7) (6,1.428122319e+11)};
\addplot[color=PVgray,densely dotted,mark=triangle*,line width=.55pt,opacity=.75,only marks,mark size=1.2pt] coordinates {(2,614802956.1) (4,102634630.7) (6,1.428122319e+11)};
\addplot[color=PVgray,dashdotted,mark=diamond*,line width=.55pt,opacity=.75,mark=none] coordinates {(2,595645131.6) (4,93071123.4)};
\addplot[color=PVgray,dashdotted,mark=diamond*,line width=.55pt,opacity=.75,mark=none] coordinates {(4,93071123.4) (6,3.827437136e+10)};
\addplot[color=PVgray,dashdotted,mark=diamond*,line width=.55pt,opacity=.75,only marks,mark size=1.2pt] coordinates {(2,595645131.6) (4,93071123.4) (6,3.827437136e+10)};
\addplot[color=PVgray,densely dashed,mark=asterisk,line width=.55pt,opacity=.75,mark=none] coordinates {(2,790349359.2) (4,1.187832251e+14)};
\addplot[color=PVgray,densely dashed,mark=asterisk,line width=.55pt,opacity=.75,mark=none] coordinates {(4,1.187832251e+14) (6,7.335326466e+14)};
\addplot[color=PVgray,densely dashed,mark=asterisk,line width=.55pt,opacity=.75,only marks,mark size=1.2pt] coordinates {(2,790349359.2) (4,1.187832251e+14) (6,7.335326466e+14)};
\addplot[color=PVgray,loosely dotted,mark=x,line width=.55pt,opacity=.75,mark=none] coordinates {(2,56170171.78) (4,5.691580984e+13)};
\addplot[color=PVgray,loosely dotted,mark=x,line width=.55pt,opacity=.75,mark=none] coordinates {(4,5.691580984e+13) (6,7.915623314e+14)};
\addplot[color=PVgray,loosely dotted,mark=x,line width=.55pt,opacity=.75,only marks,mark size=1.2pt] coordinates {(2,56170171.78) (4,5.691580984e+13) (6,7.915623314e+14)};
\end{axis}
\begin{axis}[at={(96mm,68mm)},anchor=south west,width=54mm,height=24mm,scale only axis,font=\fontsize{8}{10}\selectfont,tick label style={font=\fontsize{8}{9.5}\selectfont},label style={font=\fontsize{8}{10}\selectfont},axis line style={PVgray!65,line width=.45pt},tick style={PVgray!65},grid=major,grid style={PVgray!15,line width=.3pt},enlarge x limits=false,enlarge y limits=false,clip=true,ymode=log,ymin=10000,ymax=1e+16,xmin=3.5,xmax=12.5,xtick={4,8,12},xlabel={},ylabel={}]
\addplot[PVgray!45,dashed,line width=.4pt,mark=none] coordinates {(12,10000) (12,1e+16)};
\addplot[color=PVblue,solid,mark=*,line width=.6pt,mark=none] coordinates {(4,3682726.074) (8,62231.18874)};
\addplot[color=PVblue,solid,mark=*,line width=.6pt,mark=none] coordinates {(8,62231.18874) (12,54769.95333)};
\addplot[color=PVblue,solid,mark=*,line width=.6pt,only marks,mark size=1.2pt] coordinates {(4,3682726.074) (8,62231.18874) (12,54769.95333)};
\addplot[color=PVteal,solid,mark=*,line width=.6pt,mark=none] coordinates {(4,152544.0096) (8,48791.4123)};
\addplot[color=PVteal,solid,mark=*,line width=.6pt,mark=none] coordinates {(8,48791.4123) (12,138564.3683)};
\addplot[color=PVteal,solid,mark=*,line width=.6pt,only marks,mark size=1.2pt] coordinates {(4,152544.0096) (8,48791.4123) (12,138564.3683)};
\addplot[color=PVorange,solid,mark=*,line width=.6pt,mark=none] coordinates {(4,278963653.3) (8,5701887.59)};
\addplot[color=PVorange,solid,mark=*,line width=.6pt,mark=none] coordinates {(8,5701887.59) (12,983895.0381)};
\addplot[color=PVorange,solid,mark=*,line width=.6pt,only marks,mark size=1.2pt] coordinates {(4,278963653.3) (8,5701887.59) (12,983895.0381)};
\addplot[color=PVred,solid,mark=*,line width=.6pt,mark=none] coordinates {(4,88875465.85) (8,2.422519741e+13)};
\addplot[color=PVred,solid,mark=*,line width=.6pt,mark=none] coordinates {(8,2.422519741e+13) (12,1.303413193e+10)};
\addplot[color=PVred,solid,mark=*,line width=.6pt,only marks,mark size=1.2pt] coordinates {(4,88875465.85) (8,2.422519741e+13) (12,1.303413193e+10)};
\addplot[color=PVnavy,solid,mark=*,line width=.6pt,mark=none] coordinates {(4,1948145.927) (8,1026037.919)};
\addplot[color=PVnavy,solid,mark=*,line width=.6pt,mark=none] coordinates {(8,1026037.919) (12,460780.4459)};
\addplot[color=PVnavy,solid,mark=*,line width=.6pt,only marks,mark size=1.2pt] coordinates {(4,1948145.927) (8,1026037.919) (12,460780.4459)};
\end{axis}
\node[anchor=west,font=\fontsize{9}{11}\selectfont\bfseries] at (0.7,60) {D\quad Supervised GNN};
\begin{axis}[at={(18mm,32mm)},anchor=south west,width=54mm,height=24mm,scale only axis,font=\fontsize{8}{10}\selectfont,tick label style={font=\fontsize{8}{9.5}\selectfont},label style={font=\fontsize{8}{10}\selectfont},axis line style={PVgray!65,line width=.45pt},tick style={PVgray!65},grid=major,grid style={PVgray!15,line width=.3pt},enlarge x limits=false,enlarge y limits=false,clip=true,ymode=log,ymin=0.1,ymax=1e+33,xmin=1.75,xmax=6.25,xtick={2,4,6},xlabel={Updates (thousands)},ylabel={Validation RMSE}]
\addplot[PVgray!45,dashed,line width=.4pt,mark=none] coordinates {(6,0.1) (6,1e+33)};
\addplot[color=PVgray,solid,mark=*,line width=.55pt,opacity=.75,mark=none] coordinates {(2,0.7115439042) (4,200.9451964)};
\addplot[color=PVgray,solid,mark=*,line width=.55pt,opacity=.75,mark=none] coordinates {(4,200.9451964) (6,746154.6066)};
\addplot[color=PVgray,solid,mark=*,line width=.55pt,opacity=.75,only marks,mark size=1.2pt] coordinates {(2,0.7115439042) (4,200.9451964) (6,746154.6066)};
\addplot[color=PVgray,dashed,mark=square*,line width=.55pt,opacity=.75,mark=none] coordinates {(2,0.7109964325) (4,199.8200497)};
\addplot[color=PVgray,dashed,mark=square*,line width=.55pt,opacity=.75,mark=none] coordinates {(4,199.8200497) (6,729814.4271)};
\addplot[color=PVgray,dashed,mark=square*,line width=.55pt,opacity=.75,only marks,mark size=1.2pt] coordinates {(2,0.7109964325) (4,199.8200497) (6,729814.4271)};
\addplot[color=PVgray,densely dotted,mark=triangle*,line width=.55pt,opacity=.75,mark=none] coordinates {(2,249431.5975) (4,5886685238)};
\addplot[color=PVgray,densely dotted,mark=triangle*,line width=.55pt,opacity=.75,mark=none] coordinates {(4,5886685238) (6,248731793.6)};
\addplot[color=PVgray,densely dotted,mark=triangle*,line width=.55pt,opacity=.75,only marks,mark size=1.2pt] coordinates {(2,249431.5975) (4,5886685238) (6,248731793.6)};
\addplot[color=PVgray,dashdotted,mark=diamond*,line width=.55pt,opacity=.75,mark=none] coordinates {(2,247883.6554) (4,5755396003)};
\addplot[color=PVgray,dashdotted,mark=diamond*,line width=.55pt,opacity=.75,mark=none] coordinates {(4,5755396003) (6,241725097.6)};
\addplot[color=PVgray,dashdotted,mark=diamond*,line width=.55pt,opacity=.75,only marks,mark size=1.2pt] coordinates {(2,247883.6554) (4,5755396003) (6,241725097.6)};
\addplot[color=PVgray,densely dashed,mark=asterisk,line width=.55pt,opacity=.75,mark=none] coordinates {(2,5608757073) (4,1189334324)};
\addplot[color=PVgray,densely dashed,mark=asterisk,line width=.55pt,opacity=.75,mark=none] coordinates {(4,1189334324) (6,2.546461451e+10)};
\addplot[color=PVgray,densely dashed,mark=asterisk,line width=.55pt,opacity=.75,only marks,mark size=1.2pt] coordinates {(2,5608757073) (4,1189334324) (6,2.546461451e+10)};
\addplot[color=PVgray,loosely dotted,mark=x,line width=.55pt,opacity=.75,mark=none] coordinates {(2,6525322808) (4,808046328)};
\addplot[color=PVgray,loosely dotted,mark=x,line width=.55pt,opacity=.75,mark=none] coordinates {(4,808046328) (6,1.298547572e+10)};
\addplot[color=PVgray,loosely dotted,mark=x,line width=.55pt,opacity=.75,only marks,mark size=1.2pt] coordinates {(2,6525322808) (4,808046328) (6,1.298547572e+10)};
\end{axis}
\begin{axis}[at={(96mm,32mm)},anchor=south west,width=54mm,height=24mm,scale only axis,font=\fontsize{8}{10}\selectfont,tick label style={font=\fontsize{8}{9.5}\selectfont},label style={font=\fontsize{8}{10}\selectfont},axis line style={PVgray!65,line width=.45pt},tick style={PVgray!65},grid=major,grid style={PVgray!15,line width=.3pt},enlarge x limits=false,enlarge y limits=false,clip=true,ymode=log,ymin=0.1,ymax=1e+33,xmin=3.5,xmax=12.5,xtick={4,8,12},xlabel={Updates (thousands)},ylabel={}]
\addplot[PVgray!45,dashed,line width=.4pt,mark=none] coordinates {(12,0.1) (12,1e+33)};
\addplot[color=PVblue,solid,mark=*,line width=.6pt,mark=none] coordinates {(4,133.1761437) (8,3.773238953e+23)};
\addplot[color=PVblue,solid,mark=*,line width=.6pt,mark=none] coordinates {(8,3.773238953e+23) (12,1.858295099e+21)};
\addplot[color=PVblue,solid,mark=*,line width=.6pt,only marks,mark size=1.2pt] coordinates {(4,133.1761437) (8,3.773238953e+23) (12,1.858295099e+21)};
\addplot[color=PVteal,solid,mark=*,line width=.6pt,mark=none] coordinates {(4,6358543932) (8,1.439336309e+11)};
\addplot[color=PVteal,solid,mark=*,line width=.6pt,mark=none] coordinates {(8,1.439336309e+11) (12,2.03093063e+10)};
\addplot[color=PVteal,solid,mark=*,line width=.6pt,only marks,mark size=1.2pt] coordinates {(4,6358543932) (8,1.439336309e+11) (12,2.03093063e+10)};
\addplot[color=PVorange,solid,mark=*,line width=.6pt,mark=none] coordinates {(4,3.002522385e+15) (8,6.003626409e+30)};
\addplot[color=PVorange,solid,mark=*,line width=.6pt,mark=none] coordinates {(8,6.003626409e+30) (12,7.96637524e+31)};
\addplot[color=PVorange,solid,mark=*,line width=.6pt,only marks,mark size=1.2pt] coordinates {(4,3.002522385e+15) (8,6.003626409e+30) (12,7.96637524e+31)};
\addplot[color=PVred,solid,mark=*,line width=.6pt,mark=none] coordinates {(4,1.451634817) (8,1.752570329e+22)};
\addplot[color=PVred,solid,mark=*,line width=.6pt,mark=none] coordinates {(8,1.752570329e+22) (12,3.753573397e+21)};
\addplot[color=PVred,solid,mark=*,line width=.6pt,only marks,mark size=1.2pt] coordinates {(4,1.451634817) (8,1.752570329e+22) (12,3.753573397e+21)};
\addplot[color=PVnavy,solid,mark=*,line width=.6pt,mark=none] coordinates {(4,1738971.65) (8,2.380338017e+17)};
\addplot[color=PVnavy,solid,mark=*,line width=.6pt,mark=none] coordinates {(8,2.380338017e+17) (12,2.407424694e+14)};
\addplot[color=PVnavy,solid,mark=*,line width=.6pt,only marks,mark size=1.2pt] coordinates {(4,1738971.65) (8,2.380338017e+17) (12,2.407424694e+14)};
\end{axis}
\draw[PVgray,solid,line width=.55pt] (2.0,9)--(6.0,9);
\node[anchor=west,font=\fontsize{8}{9}\selectfont] at (6.5,9) {1};
\draw[PVgray,dashed,line width=.55pt] (13.4,9)--(17.4,9);
\node[anchor=west,font=\fontsize{8}{9}\selectfont] at (17.9,9) {2};
\draw[PVgray,densely dotted,line width=.55pt] (24.8,9)--(28.8,9);
\node[anchor=west,font=\fontsize{8}{9}\selectfont] at (29.3,9) {3};
\draw[PVgray,dashdotted,line width=.55pt] (36.2,9)--(40.2,9);
\node[anchor=west,font=\fontsize{8}{9}\selectfont] at (40.7,9) {4};
\draw[PVgray,densely dashed,line width=.55pt] (47.6,9)--(51.6,9);
\node[anchor=west,font=\fontsize{8}{9}\selectfont] at (52.1,9) {5};
\draw[PVgray,loosely dotted,line width=.55pt] (59.0,9)--(63.0,9);
\node[anchor=west,font=\fontsize{8}{9}\selectfont] at (63.5,9) {6};
\draw[PVblue,line width=.6pt] (80,9)--(84,9);
\node[anchor=west,font=\fontsize{8}{9}\selectfont] at (84.5,9) {211};
\draw[PVteal,line width=.6pt] (94,9)--(98,9);
\node[anchor=west,font=\fontsize{8}{9}\selectfont] at (98.5,9) {223};
\draw[PVorange,line width=.6pt] (108,9)--(112,9);
\node[anchor=west,font=\fontsize{8}{9}\selectfont] at (112.5,9) {227};
\draw[PVred,line width=.6pt] (122,9)--(126,9);
\node[anchor=west,font=\fontsize{8}{9}\selectfont] at (126.5,9) {229};
\draw[PVnavy,line width=.6pt] (136,9)--(140,9);
\node[anchor=west,font=\fontsize{8}{9}\selectfont] at (140.5,9) {233};
\node[align=center,text width=150mm,font=\fontsize{8}{9}\selectfont] at (76,2.5) {Search legend: candidates 1--6.\qquad Final-fit legend: fixed training seeds.};
\end{tikzpicture}
\end{checkedfigure}
\caption{Development error as training budget increases. Each row is one architecture, with all six search candidates (left: 2,000, 4,000 and 6,000 updates) and all five final-fit training seeds (right: 4,000, 8,000 and 12,000 updates). Both columns share the same complete finite logarithmic range within each row; ranges may differ between rows. The displayed values are 96-step validation rollout errors on seen structures. The fixed 6,000-update candidate scores determined hyperparameter selection; the fixed 12,000-update models generated test forecasts. Intermediate checkpoints are descriptive and were not selected by early stopping. Crosses indicate observed checkpoints with nonfinite rollout error; F marks a checkpoint unavailable after a recorded training failure. Adjacent line segments are omitted rather than interpolated through failures. Candidate numbers match the search-comparison figure; no run is removed from this display.}
\label{fig:budget-sensitivity}
\end{figure}
