## Supplementary Data - Compact audit, summary tables, code, and figure sources for "Panvascular-JEPA: Unseen-Structure Evaluation of Factorized Predictive States for Hemodynamic Networks": design_figures.tex

% Native vector figure; all coordinates in millimetres; minimum text 8 pt.
\begin{figure}[!tp]
\centering
\begin{checkedfigure}
\begin{tikzpicture}[x=1mm,y=1mm,>=Latex,
  font=\fontsize{8}{10}\selectfont,text=PVnavy,line cap=round,line join=round,
  frame/.style={draw=PVgray!30,fill=PVpale,rounded corners=1.4mm,line width=.45pt},
  box/.style={draw=PVblue!65,fill=white,rounded corners=1mm,line width=.6pt},
  title/.style={anchor=west,font=\fontsize{9.3}{11}\selectfont\bfseries},
  arrow/.style={->,draw=PVgray,line width=.8pt}]
\path[use as bounding box] (0,0) rectangle (152,150);
\node[title] at (.8,146.5) {A\quad Freeze the structures before simulating outcomes};
\path[box] (.3,111) rectangle (44,139);
\node[align=center,text width=40mm] at (22.15,125)
  {\textbf{38 nonisomorphic graphs}\\[2pt]6 nodes; 5, 6, or 7 edges\\Fixed finite structural catalogue};
\path[box] (50,111) rectangle (96,139);
\node[align=center,text width=42mm] at (73,125)
  {\textbf{24 seen structures}\\[2pt]576 training; 192 validation\\192 new-background test episodes};
\path[draw=PVteal,fill=PVteal!5,rounded corners=1mm,line width=.7pt]
  (102,111) rectangle (151.7,139);
\node[align=center,text width=45mm] at (126.85,125)
  {\textbf{14 unseen structures}\\[2pt]336 held-out test episodes\\No outcome access during tuning};
\draw[draw=PVgray,line width=.8pt] (44.5,125)--(47,125);
\draw[arrow] (47,125)--(49.4,125);
\draw[arrow] (47,125)--(47,108.5)--(126.85,108.5)--(126.85,110.6);
\node[align=center,text width=149mm,font=\fontsize{8}{10}\selectfont] at (76,104.8)
  {Independent parameter backgrounds; paired action programs; graph labels permuted consistently.};

\path[frame] (.3,47) rectangle (74.7,98);
\path[frame] (79,47) rectangle (151.7,98);
\node[title] at (3,92.5) {B\quad An explicit passive mechanism};
\node[anchor=west,font=\fontsize{9}{11}\selectfont] at (4,82)
  {$C\dot p = e_0 Q_{\mathrm{in}}-Bq-g(t)\odot p$};
\node[anchor=west,font=\fontsize{9}{11}\selectfont] at (4,73.5)
  {$L\dot q = B^{\mathsf T}p-R(t)\odot q$};
\node[anchor=north west,align=left,text width=67mm] at (4,65)
  {$p$: pressure above fixed distal reference;\\$q$: oriented edge flow; $B$: incidence matrix.\\[3pt]
   Three cycles; 96 forecast steps; repeated\\resistance changes with known future inputs.};

\node[title] at (82,92.5) {C\quad Match information and search budget};
\node[anchor=north west,align=left,text width=66mm] at (82,84)
  {\textbf{Shared inputs}\\Full initial state, network parameters,\\connectivity, and future control sequence.\\[4pt]
   \textbf{Learned comparisons}\\Monolithic JEPA; OPF with $K=1$ or $4$;\\directly supervised graph predictor.\\[4pt]
   Validation-only selection; five final seeds.};

\path[frame] (.3,.3) rectangle (151.7,41.5);
\node[title] at (3,36) {D\quad Separate prediction error from its possible sources};
\path[box] (3,7) rectangle (37,27);
\path[box] (41,7) rectangle (75,27);
\path[box] (79,7) rectangle (113,27);
\path[draw=PVteal,fill=white,rounded corners=1mm,line width=.6pt] (117,7) rectangle (149,27);
\node[align=center,text width=30mm] at (20,17)
  {\textbf{Readout}\\True-state latent\\reconstruction};
\node[align=center,text width=30mm] at (58,17)
  {\textbf{Local prediction}\\Refreshed one-step\\forecast};
\node[align=center,text width=30mm] at (96,17)
  {\textbf{Accumulation}\\Free recursive\\96-step rollout};
\node[align=center,text width=28mm] at (133,17)
  {\textbf{Physics reference}\\RLC / reduced RC\\error and runtime};
\draw[arrow] (37.4,17)--(40.5,17);
\draw[arrow] (75.4,17)--(78.5,17);
\draw[arrow] (113.4,17)--(116.5,17);
\end{tikzpicture}
\end{checkedfigure}
\caption{Design of the independent graph-structure experiment. Every episode is newly simulated; parameter backgrounds and action programs are separated from structural split units. Nodes carry compliance and edges carry resistance and inertance in a custom nodal RLC network. The pressure reference is 5~mmHg, and stored pressures are relative to that reference. Known-parameter mechanistic predictors receive the same information supplied to the learned models. The diagram describes the fixed experimental design, not an observed performance ranking or a patient-specific anatomy.}
\label{fig:independent-design}
\end{figure}


% Catalogue read exclusively from data/design_manifest.json; no outcomes loaded.
\begin{figure}[!tp]
\centering
\begin{checkedfigure}
\begin{tikzpicture}[x=1mm,y=1mm,font=\fontsize{7.5}{9}\selectfont,text=PVnavy,line cap=round,line join=round]
\path[use as bounding box] (0,0) rectangle (152,165);
\node[anchor=west,font=\fontsize{9.3}{11}\selectfont\bfseries] at (.8,161) {Six-node graph catalogue};
\fill[PVblue] (60.5,161) circle (.8);
\node[anchor=west] at (62,161) {S: seen (24)};
\fill[PVteal] (93,161) circle (.8);
\node[anchor=west] at (94.5,161) {H: held out (14)};
\draw[PVnavy,line width=.65pt,fill=white] (130.5,161) circle (1.0);
\node[anchor=west] at (132,161) {inlet};
\node[anchor=west,font=\fontsize{8.5}{10}\selectfont\bfseries] at (.8,153) {$E=5$: 6 structures (4 seen, 2 held out)};
\draw[PVgray!30,line width=.45pt] (.8,150) -- (151.2,150);
% atlas_81, seen, E=5; inlet is canonical node 0.
\path[draw=PVblue!25,fill=PVblue!3,rounded corners=.8mm,line width=.35pt] (1.000,131.200) rectangle (18.000,148.500);
\draw[PVblue,line width=.55pt] (6.263,137.755) -- (8.906,140.411);
\draw[PVblue,line width=.55pt] (6.263,137.755) -- (4.021,135.500);
\draw[PVblue,line width=.55pt] (14.979,143.259) -- (11.707,143.228);
\draw[PVblue,line width=.55pt] (11.720,146.500) -- (11.707,143.228);
\draw[PVblue,line width=.55pt] (11.707,143.228) -- (8.906,140.411);
\draw[draw=PVnavy,fill=white,line width=.7pt] (6.263,137.755) circle (.90);
\fill[PVblue] (14.979,143.259) circle (.65);
\fill[PVblue] (11.720,146.500) circle (.65);
\fill[PVblue] (11.707,143.228) circle (.65);
\fill[PVblue] (8.906,140.411) circle (.65);
\fill[PVblue] (4.021,135.500) circle (.65);
\node[text=PVblue,font=\fontsize{7.5}{9}\selectfont\bfseries] at (9.500,133.000) {81 / S};
% atlas_78, seen, E=5; inlet is canonical node 0.
\path[draw=PVblue!25,fill=PVblue!3,rounded corners=.8mm,line width=.35pt] (20.000,131.200) rectangle (37.000,148.500);
\draw[PVblue,line width=.55pt] (23.333,145.840) -- (26.777,142.533);
\draw[PVblue,line width=.55pt] (26.777,142.533) -- (30.794,138.669);
\draw[PVblue,line width=.55pt] (26.777,142.533) -- (22.915,139.901);
\draw[PVblue,line width=.55pt] (26.777,142.533) -- (29.251,146.500);
\draw[PVblue,line width=.55pt] (30.794,138.669) -- (34.085,135.500);
\draw[draw=PVnavy,fill=white,line width=.7pt] (23.333,145.840) circle (.90);
\fill[PVblue] (26.777,142.533) circle (.65);
\fill[PVblue] (30.794,138.669) circle (.65);
\fill[PVblue] (34.085,135.500) circle (.65);
\fill[PVblue] (22.915,139.901) circle (.65);
\fill[PVblue] (29.251,146.500) circle (.65);
\node[text=PVblue,font=\fontsize{7.5}{9}\selectfont\bfseries] at (28.500,133.000) {78 / S};
% atlas_80, seen, E=5; inlet is canonical node 0.
\path[draw=PVblue!25,fill=PVblue!3,rounded corners=.8mm,line width=.35pt] (39.000,131.200) rectangle (56.000,148.500);
\draw[PVblue,line width=.55pt] (51.917,136.101) -- (49.316,138.987);
\draw[PVblue,line width=.55pt] (49.316,138.987) -- (45.043,137.168);
\draw[PVblue,line width=.55pt] (49.316,138.987) -- (51.564,143.049);
\draw[PVblue,line width=.55pt] (45.043,137.168) -- (41.500,135.589);
\draw[PVblue,line width=.55pt] (51.564,143.049) -- (53.500,146.411);
\draw[draw=PVnavy,fill=white,line width=.7pt] (51.917,136.101) circle (.90);
\fill[PVblue] (49.316,138.987) circle (.65);
\fill[PVblue] (45.043,137.168) circle (.65);
\fill[PVblue] (51.564,143.049) circle (.65);
\fill[PVblue] (41.500,135.589) circle (.65);
\fill[PVblue] (53.500,146.411) circle (.65);
\node[text=PVblue,font=\fontsize{7.5}{9}\selectfont\bfseries] at (47.500,133.000) {80 / S};
% atlas_77, seen, E=5; inlet is canonical node 0.
\path[draw=PVblue!25,fill=PVblue!3,rounded corners=.8mm,line width=.35pt] (58.000,131.200) rectangle (75.000,148.500);
\draw[PVblue,line width=.55pt] (69.947,146.500) -- (66.470,141.549);
\draw[PVblue,line width=.55pt] (60.747,139.587) -- (66.470,141.549);
\draw[PVblue,line width=.55pt] (72.253,139.772) -- (66.470,141.549);
\draw[PVblue,line width=.55pt] (62.836,146.385) -- (66.470,141.549);
\draw[PVblue,line width=.55pt] (66.567,135.500) -- (66.470,141.549);
\draw[draw=PVnavy,fill=white,line width=.7pt] (69.947,146.500) circle (.90);
\fill[PVblue] (60.747,139.587) circle (.65);
\fill[PVblue] (72.253,139.772) circle (.65);
\fill[PVblue] (62.836,146.385) circle (.65);
\fill[PVblue] (66.567,135.500) circle (.65);
\fill[PVblue] (66.470,141.549) circle (.65);
\node[text=PVblue,font=\fontsize{7.5}{9}\selectfont\bfseries] at (66.500,133.000) {77 / S};
% atlas_79, heldout, E=5; inlet is canonical node 0.
\path[draw=PVteal!25,fill=PVteal!3,rounded corners=.8mm,line width=.35pt] (77.000,131.200) rectangle (94.000,148.500);
\draw[PVteal,line width=.55pt] (88.228,140.931) -- (91.500,143.830);
\draw[PVteal,line width=.55pt] (88.228,140.931) -- (91.348,137.869);
\draw[PVteal,line width=.55pt] (88.228,140.931) -- (82.772,141.069);
\draw[PVteal,line width=.55pt] (82.772,141.069) -- (79.500,138.170);
\draw[PVteal,line width=.55pt] (82.772,141.069) -- (79.652,144.131);
\draw[draw=PVnavy,fill=white,line width=.7pt] (88.228,140.931) circle (.90);
\fill[PVteal] (91.500,143.830) circle (.65);
\fill[PVteal] (91.348,137.869) circle (.65);
\fill[PVteal] (82.772,141.069) circle (.65);
\fill[PVteal] (79.500,138.170) circle (.65);
\fill[PVteal] (79.652,144.131) circle (.65);
\node[text=PVteal,font=\fontsize{7.5}{9}\selectfont\bfseries] at (85.500,133.000) {79 / H};
% atlas_83, heldout, E=5; inlet is canonical node 0.
\path[draw=PVteal!25,fill=PVteal!3,rounded corners=.8mm,line width=.35pt] (96.000,131.200) rectangle (113.000,148.500);
\draw[PVteal,line width=.55pt] (100.761,137.484) -- (98.651,135.500);
\draw[PVteal,line width=.55pt] (100.761,137.484) -- (103.223,139.799);
\draw[PVteal,line width=.55pt] (110.349,146.500) -- (108.239,144.516);
\draw[PVteal,line width=.55pt] (108.239,144.516) -- (105.777,142.201);
\draw[PVteal,line width=.55pt] (105.777,142.201) -- (103.223,139.799);
\draw[draw=PVnavy,fill=white,line width=.7pt] (100.761,137.484) circle (.90);
\fill[PVteal] (98.651,135.500) circle (.65);
\fill[PVteal] (110.349,146.500) circle (.65);
\fill[PVteal] (108.239,144.516) circle (.65);
\fill[PVteal] (105.777,142.201) circle (.65);
\fill[PVteal] (103.223,139.799) circle (.65);
\node[text=PVteal,font=\fontsize{7.5}{9}\selectfont\bfseries] at (104.500,133.000) {83 / H};
\node[anchor=west,font=\fontsize{8.5}{10}\selectfont\bfseries] at (.8,122) {$E=6$: 13 structures (8 seen, 5 held out)};
\draw[PVgray!30,line width=.45pt] (.8,119) -- (151.2,119);
% atlas_100, seen, E=6; inlet is canonical node 0.
\path[draw=PVblue!25,fill=PVblue!3,rounded corners=.8mm,line width=.35pt] (1.000,100.200) rectangle (18.000,117.500);
\draw[PVblue,line width=.55pt] (11.028,108.519) -- (15.050,108.436);
\draw[PVblue,line width=.55pt] (11.028,108.519) -- (11.188,104.500);
\draw[PVblue,line width=.55pt] (11.028,108.519) -- (7.444,112.036);
\draw[PVblue,line width=.55pt] (7.444,112.036) -- (3.950,113.538);
\draw[PVblue,line width=.55pt] (7.444,112.036) -- (5.875,115.500);
\draw[PVblue,line width=.55pt] (3.950,113.538) -- (5.875,115.500);
\draw[draw=PVnavy,fill=white,line width=.7pt] (11.028,108.519) circle (.90);
\fill[PVblue] (15.050,108.436) circle (.65);
\fill[PVblue] (11.188,104.500) circle (.65);
\fill[PVblue] (7.444,112.036) circle (.65);
\fill[PVblue] (3.950,113.538) circle (.65);
\fill[PVblue] (5.875,115.500) circle (.65);
\node[text=PVblue,font=\fontsize{7.5}{9}\selectfont\bfseries] at (9.500,102.000) {100 / S};
% atlas_95, seen, E=6; inlet is canonical node 0.
\path[draw=PVblue!25,fill=PVblue!3,rounded corners=.8mm,line width=.35pt] (20.000,100.200) rectangle (37.000,117.500);
\draw[PVblue,line width=.55pt] (30.499,112.352) -- (25.605,110.061);
\draw[PVblue,line width=.55pt] (30.499,112.352) -- (34.500,114.238);
\draw[PVblue,line width=.55pt] (25.243,105.762) -- (22.500,106.999);
\draw[PVblue,line width=.55pt] (25.243,105.762) -- (25.605,110.061);
\draw[PVblue,line width=.55pt] (22.500,106.999) -- (25.605,110.061);
\draw[PVblue,line width=.55pt] (22.612,113.518) -- (25.605,110.061);
\draw[draw=PVnavy,fill=white,line width=.7pt] (30.499,112.352) circle (.90);
\fill[PVblue] (25.243,105.762) circle (.65);
\fill[PVblue] (22.500,106.999) circle (.65);
\fill[PVblue] (22.612,113.518) circle (.65);
\fill[PVblue] (25.605,110.061) circle (.65);
\fill[PVblue] (34.500,114.238) circle (.65);
\node[text=PVblue,font=\fontsize{7.5}{9}\selectfont\bfseries] at (28.500,102.000) {95 / S};
% atlas_102, seen, E=6; inlet is canonical node 0.
\path[draw=PVblue!25,fill=PVblue!3,rounded corners=.8mm,line width=.35pt] (39.000,100.200) rectangle (56.000,117.500);
\draw[PVblue,line width=.55pt] (50.602,113.151) -- (47.894,110.384);
\draw[PVblue,line width=.55pt] (50.602,113.151) -- (52.903,115.500);
\draw[PVblue,line width=.55pt] (43.782,104.500) -- (42.097,106.150);
\draw[PVblue,line width=.55pt] (43.782,104.500) -- (45.015,107.445);
\draw[PVblue,line width=.55pt] (42.097,106.150) -- (45.015,107.445);
\draw[PVblue,line width=.55pt] (45.015,107.445) -- (47.894,110.384);
\draw[draw=PVnavy,fill=white,line width=.7pt] (50.602,113.151) circle (.90);
\fill[PVblue] (43.782,104.500) circle (.65);
\fill[PVblue] (42.097,106.150) circle (.65);
\fill[PVblue] (45.015,107.445) circle (.65);
\fill[PVblue] (47.894,110.384) circle (.65);
\fill[PVblue] (52.903,115.500) circle (.65);
\node[text=PVblue,font=\fontsize{7.5}{9}\selectfont\bfseries] at (47.500,102.000) {102 / S};
% atlas_103, seen, E=6; inlet is canonical node 0.
\path[draw=PVblue!25,fill=PVblue!3,rounded corners=.8mm,line width=.35pt] (58.000,100.200) rectangle (75.000,117.500);
\draw[PVblue,line width=.55pt] (69.388,112.302) -- (65.631,109.307);
\draw[PVblue,line width=.55pt] (69.388,112.302) -- (72.500,114.782);
\draw[PVblue,line width=.55pt] (64.035,105.494) -- (60.500,105.218);
\draw[PVblue,line width=.55pt] (64.035,105.494) -- (65.631,109.307);
\draw[PVblue,line width=.55pt] (61.558,108.603) -- (60.500,105.218);
\draw[PVblue,line width=.55pt] (61.558,108.603) -- (65.631,109.307);
\draw[draw=PVnavy,fill=white,line width=.7pt] (69.388,112.302) circle (.90);
\fill[PVblue] (64.035,105.494) circle (.65);
\fill[PVblue] (61.558,108.603) circle (.65);
\fill[PVblue] (60.500,105.218) circle (.65);
\fill[PVblue] (65.631,109.307) circle (.65);
\fill[PVblue] (72.500,114.782) circle (.65);
\node[text=PVblue,font=\fontsize{7.5}{9}\selectfont\bfseries] at (66.500,102.000) {103 / S};
% atlas_104, seen, E=6; inlet is canonical node 0.
\path[draw=PVblue!25,fill=PVblue!3,rounded corners=.8mm,line width=.35pt] (77.000,100.200) rectangle (94.000,117.500);
\draw[PVblue,line width=.55pt] (82.458,107.930) -- (87.593,108.001);
\draw[PVblue,line width=.55pt] (82.458,107.930) -- (79.500,111.693);
\draw[PVblue,line width=.55pt] (87.593,108.001) -- (86.626,113.045);
\draw[PVblue,line width=.55pt] (87.593,108.001) -- (91.500,104.818);
\draw[PVblue,line width=.55pt] (86.626,113.045) -- (82.343,115.182);
\draw[PVblue,line width=.55pt] (82.343,115.182) -- (79.500,111.693);
\draw[draw=PVnavy,fill=white,line width=.7pt] (82.458,107.930) circle (.90);
\fill[PVblue] (87.593,108.001) circle (.65);
\fill[PVblue] (86.626,113.045) circle (.65);
\fill[PVblue] (82.343,115.182) circle (.65);
\fill[PVblue] (79.500,111.693) circle (.65);
\fill[PVblue] (91.500,104.818) circle (.65);
\node[text=PVblue,font=\fontsize{7.5}{9}\selectfont\bfseries] at (85.500,102.000) {104 / S};
% atlas_98, seen, E=6; inlet is canonical node 0.
\path[draw=PVblue!25,fill=PVblue!3,rounded corners=.8mm,line width=.35pt] (96.000,100.200) rectangle (113.000,117.500);
\draw[PVblue,line width=.55pt] (101.198,109.051) -- (98.853,112.916);
\draw[PVblue,line width=.55pt] (101.198,109.051) -- (105.335,112.423);
\draw[PVblue,line width=.55pt] (101.198,109.051) -- (99.439,104.500);
\draw[PVblue,line width=.55pt] (102.023,115.500) -- (98.853,112.916);
\draw[PVblue,line width=.55pt] (102.023,115.500) -- (105.335,112.423);
\draw[PVblue,line width=.55pt] (105.335,112.423) -- (110.147,113.229);
\draw[draw=PVnavy,fill=white,line width=.7pt] (101.198,109.051) circle (.90);
\fill[PVblue] (102.023,115.500) circle (.65);
\fill[PVblue] (98.853,112.916) circle (.65);
\fill[PVblue] (105.335,112.423) circle (.65);
\fill[PVblue] (99.439,104.500) circle (.65);
\fill[PVblue] (110.147,113.229) circle (.65);
\node[text=PVblue,font=\fontsize{7.5}{9}\selectfont\bfseries] at (104.500,102.000) {98 / S};
% atlas_97, seen, E=6; inlet is canonical node 0.
\path[draw=PVblue!25,fill=PVblue!3,rounded corners=.8mm,line width=.35pt] (115.000,100.200) rectangle (132.000,117.500);
\draw[PVblue,line width=.55pt] (129.451,104.500) -- (125.998,106.324);
\draw[PVblue,line width=.55pt] (122.691,108.991) -- (125.998,106.324);
\draw[PVblue,line width=.55pt] (122.691,108.991) -- (119.893,112.555);
\draw[PVblue,line width=.55pt] (122.691,108.991) -- (123.025,105.737);
\draw[PVblue,line width=.55pt] (125.998,106.324) -- (123.025,105.737);
\draw[PVblue,line width=.55pt] (119.893,112.555) -- (117.549,115.500);
\draw[draw=PVnavy,fill=white,line width=.7pt] (129.451,104.500) circle (.90);
\fill[PVblue] (122.691,108.991) circle (.65);
\fill[PVblue] (125.998,106.324) circle (.65);
\fill[PVblue] (119.893,112.555) circle (.65);
\fill[PVblue] (123.025,105.737) circle (.65);
\fill[PVblue] (117.549,115.500) circle (.65);
\node[text=PVblue,font=\fontsize{7.5}{9}\selectfont\bfseries] at (123.500,102.000) {97 / S};
% atlas_105, seen, E=6; inlet is canonical node 0.
\path[draw=PVblue!25,fill=PVblue!3,rounded corners=.8mm,line width=.35pt] (134.000,100.200) rectangle (151.000,117.500);
\draw[PVblue,line width=.55pt] (138.972,105.115) -- (136.500,110.611);
\draw[PVblue,line width=.55pt] (138.972,105.115) -- (144.971,104.504);
\draw[PVblue,line width=.55pt] (136.500,110.611) -- (140.029,115.496);
\draw[PVblue,line width=.55pt] (140.029,115.496) -- (146.028,114.885);
\draw[PVblue,line width=.55pt] (146.028,114.885) -- (148.500,109.389);
\draw[PVblue,line width=.55pt] (148.500,109.389) -- (144.971,104.504);
\draw[draw=PVnavy,fill=white,line width=.7pt] (138.972,105.115) circle (.90);
\fill[PVblue] (136.500,110.611) circle (.65);
\fill[PVblue] (140.029,115.496) circle (.65);
\fill[PVblue] (146.028,114.885) circle (.65);
\fill[PVblue] (148.500,109.389) circle (.65);
\fill[PVblue] (144.971,104.504) circle (.65);
\node[text=PVblue,font=\fontsize{7.5}{9}\selectfont\bfseries] at (142.500,102.000) {105 / S};
% atlas_96, heldout, E=6; inlet is canonical node 0.
\path[draw=PVteal!25,fill=PVteal!3,rounded corners=.8mm,line width=.35pt] (1.000,78.200) rectangle (18.000,95.500);
\draw[PVteal,line width=.55pt] (8.530,87.755) -- (13.647,87.286);
\draw[PVteal,line width=.55pt] (8.530,87.755) -- (11.216,83.375);
\draw[PVteal,line width=.55pt] (8.530,87.755) -- (3.500,86.944);
\draw[PVteal,line width=.55pt] (8.530,87.755) -- (7.031,92.625);
\draw[PVteal,line width=.55pt] (15.500,83.423) -- (13.647,87.286);
\draw[PVteal,line width=.55pt] (15.500,83.423) -- (11.216,83.375);
\draw[draw=PVnavy,fill=white,line width=.7pt] (8.530,87.755) circle (.90);
\fill[PVteal] (15.500,83.423) circle (.65);
\fill[PVteal] (13.647,87.286) circle (.65);
\fill[PVteal] (11.216,83.375) circle (.65);
\fill[PVteal] (3.500,86.944) circle (.65);
\fill[PVteal] (7.031,92.625) circle (.65);
\node[text=PVteal,font=\fontsize{7.5}{9}\selectfont\bfseries] at (9.500,80.000) {96 / H};
% atlas_94, heldout, E=6; inlet is canonical node 0.
\path[draw=PVteal!25,fill=PVteal!3,rounded corners=.8mm,line width=.35pt] (20.000,78.200) rectangle (37.000,95.500);
\draw[PVteal,line width=.55pt] (34.500,92.505) -- (30.463,90.570);
\draw[PVteal,line width=.55pt] (27.702,82.573) -- (28.045,87.037);
\draw[PVteal,line width=.55pt] (22.500,93.427) -- (26.195,90.898);
\draw[PVteal,line width=.55pt] (26.195,90.898) -- (30.463,90.570);
\draw[PVteal,line width=.55pt] (26.195,90.898) -- (28.045,87.037);
\draw[PVteal,line width=.55pt] (30.463,90.570) -- (28.045,87.037);
\draw[draw=PVnavy,fill=white,line width=.7pt] (34.500,92.505) circle (.90);
\fill[PVteal] (27.702,82.573) circle (.65);
\fill[PVteal] (22.500,93.427) circle (.65);
\fill[PVteal] (26.195,90.898) circle (.65);
\fill[PVteal] (30.463,90.570) circle (.65);
\fill[PVteal] (28.045,87.037) circle (.65);
\node[text=PVteal,font=\fontsize{7.5}{9}\selectfont\bfseries] at (28.500,80.000) {94 / H};
% atlas_92, heldout, E=6; inlet is canonical node 0.
\path[draw=PVteal!25,fill=PVteal!3,rounded corners=.8mm,line width=.35pt] (39.000,78.200) rectangle (56.000,95.500);
\draw[PVteal,line width=.55pt] (49.242,93.500) -- (47.230,87.364);
\draw[PVteal,line width=.55pt] (42.348,83.136) -- (47.230,87.364);
\draw[PVteal,line width=.55pt] (41.868,90.931) -- (47.230,87.364);
\draw[PVteal,line width=.55pt] (50.894,82.500) -- (53.132,85.864);
\draw[PVteal,line width=.55pt] (50.894,82.500) -- (47.230,87.364);
\draw[PVteal,line width=.55pt] (53.132,85.864) -- (47.230,87.364);
\draw[draw=PVnavy,fill=white,line width=.7pt] (49.242,93.500) circle (.90);
\fill[PVteal] (42.348,83.136) circle (.65);
\fill[PVteal] (41.868,90.931) circle (.65);
\fill[PVteal] (50.894,82.500) circle (.65);
\fill[PVteal] (53.132,85.864) circle (.65);
\fill[PVteal] (47.230,87.364) circle (.65);
\node[text=PVteal,font=\fontsize{7.5}{9}\selectfont\bfseries] at (47.500,80.000) {92 / H};
% atlas_93, heldout, E=6; inlet is canonical node 0.
\path[draw=PVteal!25,fill=PVteal!3,rounded corners=.8mm,line width=.35pt] (58.000,78.200) rectangle (75.000,95.500);
\draw[PVteal,line width=.55pt] (68.760,86.424) -- (69.146,89.907);
\draw[PVteal,line width=.55pt] (68.760,86.424) -- (64.837,87.043);
\draw[PVteal,line width=.55pt] (60.500,88.783) -- (64.837,87.043);
\draw[PVteal,line width=.55pt] (62.579,82.895) -- (64.837,87.043);
\draw[PVteal,line width=.55pt] (69.146,89.907) -- (64.837,87.043);
\draw[PVteal,line width=.55pt] (69.146,89.907) -- (72.500,93.105);
\draw[draw=PVnavy,fill=white,line width=.7pt] (68.760,86.424) circle (.90);
\fill[PVteal] (60.500,88.783) circle (.65);
\fill[PVteal] (62.579,82.895) circle (.65);
\fill[PVteal] (69.146,89.907) circle (.65);
\fill[PVteal] (64.837,87.043) circle (.65);
\fill[PVteal] (72.500,93.105) circle (.65);
\node[text=PVteal,font=\fontsize{7.5}{9}\selectfont\bfseries] at (66.500,80.000) {93 / H};
% atlas_99, heldout, E=6; inlet is canonical node 0.
\path[draw=PVteal!25,fill=PVteal!3,rounded corners=.8mm,line width=.35pt] (77.000,78.200) rectangle (94.000,95.500);
\draw[PVteal,line width=.55pt] (91.500,93.437) -- (88.273,90.513);
\draw[PVteal,line width=.55pt] (79.500,82.563) -- (82.726,85.486);
\draw[PVteal,line width=.55pt] (88.273,90.513) -- (86.818,86.546);
\draw[PVteal,line width=.55pt] (88.273,90.513) -- (84.182,89.454);
\draw[PVteal,line width=.55pt] (82.726,85.486) -- (86.818,86.546);
\draw[PVteal,line width=.55pt] (82.726,85.486) -- (84.182,89.454);
\draw[draw=PVnavy,fill=white,line width=.7pt] (91.500,93.437) circle (.90);
\fill[PVteal] (79.500,82.563) circle (.65);
\fill[PVteal] (88.273,90.513) circle (.65);
\fill[PVteal] (82.726,85.486) circle (.65);
\fill[PVteal] (86.818,86.546) circle (.65);
\fill[PVteal] (84.182,89.454) circle (.65);
\node[text=PVteal,font=\fontsize{7.5}{9}\selectfont\bfseries] at (85.500,80.000) {99 / H};
\node[anchor=west,font=\fontsize{8.5}{10}\selectfont\bfseries] at (.8,69) {$E=7$: 19 structures (12 seen, 7 held out)};
\draw[PVgray!30,line width=.45pt] (.8,66) -- (151.2,66);
% atlas_128, seen, E=7; inlet is canonical node 0.
\path[draw=PVblue!25,fill=PVblue!3,rounded corners=.8mm,line width=.35pt] (1.000,47.200) rectangle (18.000,64.500);
\draw[PVblue,line width=.55pt] (9.715,59.625) -- (3.912,59.981);
\draw[PVblue,line width=.55pt] (9.715,59.625) -- (9.285,54.375);
\draw[PVblue,line width=.55pt] (9.715,59.625) -- (15.500,59.039);
\draw[PVblue,line width=.55pt] (3.912,59.981) -- (3.500,54.961);
\draw[PVblue,line width=.55pt] (3.500,54.961) -- (9.285,54.375);
\draw[PVblue,line width=.55pt] (9.285,54.375) -- (15.088,54.019);
\draw[PVblue,line width=.55pt] (15.088,54.019) -- (15.500,59.039);
\draw[draw=PVnavy,fill=white,line width=.7pt] (9.715,59.625) circle (.90);
\fill[PVblue] (3.912,59.981) circle (.65);
\fill[PVblue] (3.500,54.961) circle (.65);
\fill[PVblue] (9.285,54.375) circle (.65);
\fill[PVblue] (15.088,54.019) circle (.65);
\fill[PVblue] (15.500,59.039) circle (.65);
\node[text=PVblue,font=\fontsize{7.5}{9}\selectfont\bfseries] at (9.500,49.000) {128 / S};
% atlas_130, seen, E=7; inlet is canonical node 0.
\path[draw=PVblue!25,fill=PVblue!3,rounded corners=.8mm,line width=.35pt] (20.000,47.200) rectangle (37.000,64.500);
\draw[PVblue,line width=.55pt] (30.593,55.253) -- (34.500,53.937);
\draw[PVblue,line width=.55pt] (30.593,55.253) -- (32.585,51.643);
\draw[PVblue,line width=.55pt] (30.593,55.253) -- (26.407,58.747);
\draw[PVblue,line width=.55pt] (34.500,53.937) -- (32.585,51.643);
\draw[PVblue,line width=.55pt] (26.407,58.747) -- (22.500,60.063);
\draw[PVblue,line width=.55pt] (26.407,58.747) -- (24.415,62.357);
\draw[PVblue,line width=.55pt] (22.500,60.063) -- (24.415,62.357);
\draw[draw=PVnavy,fill=white,line width=.7pt] (30.593,55.253) circle (.90);
\fill[PVblue] (34.500,53.937) circle (.65);
\fill[PVblue] (32.585,51.643) circle (.65);
\fill[PVblue] (26.407,58.747) circle (.65);
\fill[PVblue] (22.500,60.063) circle (.65);
\fill[PVblue] (24.415,62.357) circle (.65);
\node[text=PVblue,font=\fontsize{7.5}{9}\selectfont\bfseries] at (28.500,49.000) {130 / S};
% atlas_123, seen, E=7; inlet is canonical node 0.
\path[draw=PVblue!25,fill=PVblue!3,rounded corners=.8mm,line width=.35pt] (39.000,47.200) rectangle (56.000,64.500);
\draw[PVblue,line width=.55pt] (50.484,54.539) -- (46.843,57.542);
\draw[PVblue,line width=.55pt] (50.484,54.539) -- (53.500,52.053);
\draw[PVblue,line width=.55pt] (43.139,58.961) -- (44.744,60.907);
\draw[PVblue,line width=.55pt] (43.139,58.961) -- (41.500,61.947);
\draw[PVblue,line width=.55pt] (43.139,58.961) -- (46.843,57.542);
\draw[PVblue,line width=.55pt] (44.744,60.907) -- (41.500,61.947);
\draw[PVblue,line width=.55pt] (44.744,60.907) -- (46.843,57.542);
\draw[draw=PVnavy,fill=white,line width=.7pt] (50.484,54.539) circle (.90);
\fill[PVblue] (43.139,58.961) circle (.65);
\fill[PVblue] (44.744,60.907) circle (.65);
\fill[PVblue] (41.500,61.947) circle (.65);
\fill[PVblue] (46.843,57.542) circle (.65);
\fill[PVblue] (53.500,52.053) circle (.65);
\node[text=PVblue,font=\fontsize{7.5}{9}\selectfont\bfseries] at (47.500,49.000) {123 / S};
% atlas_113, seen, E=7; inlet is canonical node 0.
\path[draw=PVblue!25,fill=PVblue!3,rounded corners=.8mm,line width=.35pt] (58.000,47.200) rectangle (75.000,64.500);
\draw[PVblue,line width=.55pt] (68.037,54.021) -- (64.027,53.295);
\draw[PVblue,line width=.55pt] (68.037,54.021) -- (63.924,56.884);
\draw[PVblue,line width=.55pt] (68.037,54.021) -- (72.500,52.401);
\draw[PVblue,line width=.55pt] (60.500,54.400) -- (64.027,53.295);
\draw[PVblue,line width=.55pt] (60.500,54.400) -- (63.924,56.884);
\draw[PVblue,line width=.55pt] (64.027,53.295) -- (63.924,56.884);
\draw[PVblue,line width=.55pt] (63.924,56.884) -- (62.779,61.599);
\draw[draw=PVnavy,fill=white,line width=.7pt] (68.037,54.021) circle (.90);
\fill[PVblue] (60.500,54.400) circle (.65);
\fill[PVblue] (64.027,53.295) circle (.65);
\fill[PVblue] (63.924,56.884) circle (.65);
\fill[PVblue] (72.500,52.401) circle (.65);
\fill[PVblue] (62.779,61.599) circle (.65);
\node[text=PVblue,font=\fontsize{7.5}{9}\selectfont\bfseries] at (66.500,49.000) {113 / S};
% atlas_120, seen, E=7; inlet is canonical node 0.
\path[draw=PVblue!25,fill=PVblue!3,rounded corners=.8mm,line width=.35pt] (77.000,47.200) rectangle (94.000,64.500);
\draw[PVblue,line width=.55pt] (82.983,54.537) -- (87.254,57.963);
\draw[PVblue,line width=.55pt] (82.983,54.537) -- (79.500,51.741);
\draw[PVblue,line width=.55pt] (87.498,62.259) -- (90.329,60.431);
\draw[PVblue,line width=.55pt] (87.498,62.259) -- (87.254,57.963);
\draw[PVblue,line width=.55pt] (91.500,57.271) -- (90.329,60.431);
\draw[PVblue,line width=.55pt] (91.500,57.271) -- (87.254,57.963);
\draw[PVblue,line width=.55pt] (90.329,60.431) -- (87.254,57.963);
\draw[draw=PVnavy,fill=white,line width=.7pt] (82.983,54.537) circle (.90);
\fill[PVblue] (87.498,62.259) circle (.65);
\fill[PVblue] (91.500,57.271) circle (.65);
\fill[PVblue] (90.329,60.431) circle (.65);
\fill[PVblue] (87.254,57.963) circle (.65);
\fill[PVblue] (79.500,51.741) circle (.65);
\node[text=PVblue,font=\fontsize{7.5}{9}\selectfont\bfseries] at (85.500,49.000) {120 / S};
% atlas_125, seen, E=7; inlet is canonical node 0.
\path[draw=PVblue!25,fill=PVblue!3,rounded corners=.8mm,line width=.35pt] (96.000,47.200) rectangle (113.000,64.500);
\draw[PVblue,line width=.55pt] (98.500,54.000) -- (110.500,54.000);
\draw[PVblue,line width=.55pt] (100.000,55.500) -- (110.500,54.000);
\draw[PVblue,line width=.55pt] (100.000,55.500) -- (104.500,60.000);
\draw[PVblue,line width=.55pt] (100.000,55.500) -- (106.000,57.000);
\draw[PVblue,line width=.55pt] (110.500,54.000) -- (106.000,58.500);
\draw[PVblue,line width=.55pt] (104.500,60.000) -- (106.000,58.500);
\draw[PVblue,line width=.55pt] (106.000,57.000) -- (106.000,58.500);
\draw[draw=PVnavy,fill=white,line width=.7pt] (98.500,54.000) circle (.90);
\fill[PVblue] (100.000,55.500) circle (.65);
\fill[PVblue] (110.500,54.000) circle (.65);
\fill[PVblue] (104.500,60.000) circle (.65);
\fill[PVblue] (106.000,57.000) circle (.65);
\fill[PVblue] (106.000,58.500) circle (.65);
\node[text=PVblue,font=\fontsize{7.5}{9}\selectfont\bfseries] at (104.500,49.000) {125 / S};
% atlas_111, seen, E=7; inlet is canonical node 0.
\path[draw=PVblue!25,fill=PVblue!3,rounded corners=.8mm,line width=.35pt] (115.000,47.200) rectangle (132.000,64.500);
\draw[PVblue,line width=.55pt] (129.500,53.798) -- (123.291,56.982);
\draw[PVblue,line width=.55pt] (119.853,51.539) -- (117.500,55.721);
\draw[PVblue,line width=.55pt] (119.853,51.539) -- (123.291,56.982);
\draw[PVblue,line width=.55pt] (117.901,60.502) -- (117.500,55.721);
\draw[PVblue,line width=.55pt] (117.901,60.502) -- (123.291,56.982);
\draw[PVblue,line width=.55pt] (117.500,55.721) -- (123.291,56.982);
\draw[PVblue,line width=.55pt] (123.291,56.982) -- (127.612,62.461);
\draw[draw=PVnavy,fill=white,line width=.7pt] (129.500,53.798) circle (.90);
\fill[PVblue] (119.853,51.539) circle (.65);
\fill[PVblue] (117.901,60.502) circle (.65);
\fill[PVblue] (117.500,55.721) circle (.65);
\fill[PVblue] (123.291,56.982) circle (.65);
\fill[PVblue] (127.612,62.461) circle (.65);
\node[text=PVblue,font=\fontsize{7.5}{9}\selectfont\bfseries] at (123.500,49.000) {111 / S};
% atlas_121, seen, E=7; inlet is canonical node 0.
\path[draw=PVblue!25,fill=PVblue!3,rounded corners=.8mm,line width=.35pt] (134.000,47.200) rectangle (151.000,64.500);
\draw[PVblue,line width=.55pt] (146.149,51.500) -- (143.815,56.294);
\draw[PVblue,line width=.55pt] (142.221,59.569) -- (143.815,56.294);
\draw[PVblue,line width=.55pt] (142.221,59.569) -- (140.793,62.500);
\draw[PVblue,line width=.55pt] (143.815,56.294) -- (138.851,58.127);
\draw[PVblue,line width=.55pt] (143.815,56.294) -- (145.433,61.332);
\draw[PVblue,line width=.55pt] (138.851,58.127) -- (140.793,62.500);
\draw[PVblue,line width=.55pt] (140.793,62.500) -- (145.433,61.332);
\draw[draw=PVnavy,fill=white,line width=.7pt] (146.149,51.500) circle (.90);
\fill[PVblue] (142.221,59.569) circle (.65);
\fill[PVblue] (143.815,56.294) circle (.65);
\fill[PVblue] (138.851,58.127) circle (.65);
\fill[PVblue] (140.793,62.500) circle (.65);
\fill[PVblue] (145.433,61.332) circle (.65);
\node[text=PVblue,font=\fontsize{7.5}{9}\selectfont\bfseries] at (142.500,49.000) {121 / S};
% atlas_127, seen, E=7; inlet is canonical node 0.
\path[draw=PVblue!25,fill=PVblue!3,rounded corners=.8mm,line width=.35pt] (1.000,25.200) rectangle (18.000,42.500);
\draw[PVblue,line width=.55pt] (3.692,36.808) -- (9.890,39.470);
\draw[PVblue,line width=.55pt] (3.692,36.808) -- (3.500,30.676);
\draw[PVblue,line width=.55pt] (15.500,40.404) -- (9.890,39.470);
\draw[PVblue,line width=.55pt] (15.500,40.404) -- (13.425,35.109);
\draw[PVblue,line width=.55pt] (9.890,39.470) -- (13.425,35.109);
\draw[PVblue,line width=.55pt] (13.425,35.109) -- (9.539,29.596);
\draw[PVblue,line width=.55pt] (9.539,29.596) -- (3.500,30.676);
\draw[draw=PVnavy,fill=white,line width=.7pt] (3.692,36.808) circle (.90);
\fill[PVblue] (15.500,40.404) circle (.65);
\fill[PVblue] (9.890,39.470) circle (.65);
\fill[PVblue] (13.425,35.109) circle (.65);
\fill[PVblue] (9.539,29.596) circle (.65);
\fill[PVblue] (3.500,30.676) circle (.65);
\node[text=PVblue,font=\fontsize{7.5}{9}\selectfont\bfseries] at (9.500,27.000) {127 / S};
% atlas_122, seen, E=7; inlet is canonical node 0.
\path[draw=PVblue!25,fill=PVblue!3,rounded corners=.8mm,line width=.35pt] (20.000,25.200) rectangle (37.000,42.500);
\draw[PVblue,line width=.55pt] (26.110,34.152) -- (31.098,36.031);
\draw[PVblue,line width=.55pt] (26.110,34.152) -- (26.348,30.271);
\draw[PVblue,line width=.55pt] (26.110,34.152) -- (22.500,31.559);
\draw[PVblue,line width=.55pt] (31.098,36.031) -- (30.867,31.372);
\draw[PVblue,line width=.55pt] (31.098,36.031) -- (34.500,39.729);
\draw[PVblue,line width=.55pt] (30.867,31.372) -- (26.348,30.271);
\draw[PVblue,line width=.55pt] (26.348,30.271) -- (22.500,31.559);
\draw[draw=PVnavy,fill=white,line width=.7pt] (26.110,34.152) circle (.90);
\fill[PVblue] (31.098,36.031) circle (.65);
\fill[PVblue] (30.867,31.372) circle (.65);
\fill[PVblue] (26.348,30.271) circle (.65);
\fill[PVblue] (22.500,31.559) circle (.65);
\fill[PVblue] (34.500,39.729) circle (.65);
\node[text=PVblue,font=\fontsize{7.5}{9}\selectfont\bfseries] at (28.500,27.000) {122 / S};
% atlas_117, seen, E=7; inlet is canonical node 0.
\path[draw=PVblue!25,fill=PVblue!3,rounded corners=.8mm,line width=.35pt] (39.000,25.200) rectangle (56.000,42.500);
\draw[PVblue,line width=.55pt] (48.129,35.800) -- (53.096,40.280);
\draw[PVblue,line width=.55pt] (48.129,35.800) -- (43.938,40.500);
\draw[PVblue,line width=.55pt] (48.129,35.800) -- (41.904,36.827);
\draw[PVblue,line width=.55pt] (48.129,35.800) -- (48.514,29.500);
\draw[PVblue,line width=.55pt] (48.129,35.800) -- (52.376,31.147);
\draw[PVblue,line width=.55pt] (43.938,40.500) -- (41.904,36.827);
\draw[PVblue,line width=.55pt] (48.514,29.500) -- (52.376,31.147);
\draw[draw=PVnavy,fill=white,line width=.7pt] (48.129,35.800) circle (.90);
\fill[PVblue] (53.096,40.280) circle (.65);
\fill[PVblue] (43.938,40.500) circle (.65);
\fill[PVblue] (41.904,36.827) circle (.65);
\fill[PVblue] (48.514,29.500) circle (.65);
\fill[PVblue] (52.376,31.147) circle (.65);
\node[text=PVblue,font=\fontsize{7.5}{9}\selectfont\bfseries] at (47.500,27.000) {117 / S};
% atlas_112, seen, E=7; inlet is canonical node 0.
\path[draw=PVblue!25,fill=PVblue!3,rounded corners=.8mm,line width=.35pt] (58.000,25.200) rectangle (75.000,42.500);
\draw[PVblue,line width=.55pt] (60.500,40.226) -- (64.576,36.676);
\draw[PVblue,line width=.55pt] (64.576,36.676) -- (68.424,33.324);
\draw[PVblue,line width=.55pt] (64.576,36.676) -- (64.129,32.278);
\draw[PVblue,line width=.55pt] (64.576,36.676) -- (68.871,37.722);
\draw[PVblue,line width=.55pt] (68.424,33.324) -- (72.500,29.774);
\draw[PVblue,line width=.55pt] (68.424,33.324) -- (64.129,32.278);
\draw[PVblue,line width=.55pt] (68.424,33.324) -- (68.871,37.722);
\draw[draw=PVnavy,fill=white,line width=.7pt] (60.500,40.226) circle (.90);
\fill[PVblue] (64.576,36.676) circle (.65);
\fill[PVblue] (68.424,33.324) circle (.65);
\fill[PVblue] (72.500,29.774) circle (.65);
\fill[PVblue] (64.129,32.278) circle (.65);
\fill[PVblue] (68.871,37.722) circle (.65);
\node[text=PVblue,font=\fontsize{7.5}{9}\selectfont\bfseries] at (66.500,27.000) {112 / S};
% atlas_114, heldout, E=7; inlet is canonical node 0.
\path[draw=PVteal!25,fill=PVteal!3,rounded corners=.8mm,line width=.35pt] (77.000,25.200) rectangle (94.000,42.500);
\draw[PVteal,line width=.55pt] (83.797,40.309) -- (84.651,35.224);
\draw[PVteal,line width=.55pt] (87.458,30.982) -- (89.389,33.371);
\draw[PVteal,line width=.55pt] (87.458,30.982) -- (91.500,29.691);
\draw[PVteal,line width=.55pt] (87.458,30.982) -- (84.651,35.224);
\draw[PVteal,line width=.55pt] (89.389,33.371) -- (91.500,29.691);
\draw[PVteal,line width=.55pt] (89.389,33.371) -- (84.651,35.224);
\draw[PVteal,line width=.55pt] (84.651,35.224) -- (79.500,34.991);
\draw[draw=PVnavy,fill=white,line width=.7pt] (83.797,40.309) circle (.90);
\fill[PVteal] (87.458,30.982) circle (.65);
\fill[PVteal] (89.389,33.371) circle (.65);
\fill[PVteal] (91.500,29.691) circle (.65);
\fill[PVteal] (84.651,35.224) circle (.65);
\fill[PVteal] (79.500,34.991) circle (.65);
\node[text=PVteal,font=\fontsize{7.5}{9}\selectfont\bfseries] at (85.500,27.000) {114 / H};
% atlas_118, heldout, E=7; inlet is canonical node 0.
\path[draw=PVteal!25,fill=PVteal!3,rounded corners=.8mm,line width=.35pt] (96.000,25.200) rectangle (113.000,42.500);
\draw[PVteal,line width=.55pt] (99.269,39.974) -- (102.316,35.046);
\draw[PVteal,line width=.55pt] (99.269,39.974) -- (104.409,40.500);
\draw[PVteal,line width=.55pt] (102.316,35.046) -- (108.836,35.012);
\draw[PVteal,line width=.55pt] (102.316,35.046) -- (104.409,40.500);
\draw[PVteal,line width=.55pt] (102.316,35.046) -- (98.646,29.500);
\draw[PVteal,line width=.55pt] (108.836,35.012) -- (110.354,40.217);
\draw[PVteal,line width=.55pt] (110.354,40.217) -- (104.409,40.500);
\draw[draw=PVnavy,fill=white,line width=.7pt] (99.269,39.974) circle (.90);
\fill[PVteal] (102.316,35.046) circle (.65);
\fill[PVteal] (108.836,35.012) circle (.65);
\fill[PVteal] (110.354,40.217) circle (.65);
\fill[PVteal] (104.409,40.500) circle (.65);
\fill[PVteal] (98.646,29.500) circle (.65);
\node[text=PVteal,font=\fontsize{7.5}{9}\selectfont\bfseries] at (104.500,27.000) {118 / H};
% atlas_126, heldout, E=7; inlet is canonical node 0.
\path[draw=PVteal!25,fill=PVteal!3,rounded corners=.8mm,line width=.35pt] (115.000,25.200) rectangle (132.000,42.500);
\draw[PVteal,line width=.55pt] (129.500,29.889) -- (128.053,34.374);
\draw[PVteal,line width=.55pt] (129.500,29.889) -- (124.813,30.380);
\draw[PVteal,line width=.55pt] (117.500,37.267) -- (122.517,35.555);
\draw[PVteal,line width=.55pt] (117.500,37.267) -- (119.807,40.111);
\draw[PVteal,line width=.55pt] (128.053,34.374) -- (122.517,35.555);
\draw[PVteal,line width=.55pt] (122.517,35.555) -- (119.807,40.111);
\draw[PVteal,line width=.55pt] (122.517,35.555) -- (124.813,30.380);
\draw[draw=PVnavy,fill=white,line width=.7pt] (129.500,29.889) circle (.90);
\fill[PVteal] (117.500,37.267) circle (.65);
\fill[PVteal] (128.053,34.374) circle (.65);
\fill[PVteal] (122.517,35.555) circle (.65);
\fill[PVteal] (119.807,40.111) circle (.65);
\fill[PVteal] (124.813,30.380) circle (.65);
\node[text=PVteal,font=\fontsize{7.5}{9}\selectfont\bfseries] at (123.500,27.000) {126 / H};
% atlas_129, heldout, E=7; inlet is canonical node 0.
\path[draw=PVteal!25,fill=PVteal!3,rounded corners=.8mm,line width=.35pt] (134.000,25.200) rectangle (151.000,42.500);
\draw[PVteal,line width=.55pt] (141.685,30.489) -- (148.453,31.766);
\draw[PVteal,line width=.55pt] (141.685,30.489) -- (140.736,35.005);
\draw[PVteal,line width=.55pt] (141.685,30.489) -- (136.500,35.044);
\draw[PVteal,line width=.55pt] (148.453,31.766) -- (148.500,38.146);
\draw[PVteal,line width=.55pt] (148.500,38.146) -- (141.751,39.511);
\draw[PVteal,line width=.55pt] (141.751,39.511) -- (140.736,35.005);
\draw[PVteal,line width=.55pt] (141.751,39.511) -- (136.500,35.044);
\draw[draw=PVnavy,fill=white,line width=.7pt] (141.685,30.489) circle (.90);
\fill[PVteal] (148.453,31.766) circle (.65);
\fill[PVteal] (148.500,38.146) circle (.65);
\fill[PVteal] (141.751,39.511) circle (.65);
\fill[PVteal] (140.736,35.005) circle (.65);
\fill[PVteal] (136.500,35.044) circle (.65);
\node[text=PVteal,font=\fontsize{7.5}{9}\selectfont\bfseries] at (142.500,27.000) {129 / H};
% atlas_115, heldout, E=7; inlet is canonical node 0.
\path[draw=PVteal!25,fill=PVteal!3,rounded corners=.8mm,line width=.35pt] (1.000,3.200) rectangle (18.000,20.500);
\draw[PVteal,line width=.55pt] (15.500,17.669) -- (12.347,15.215);
\draw[PVteal,line width=.55pt] (12.347,15.215) -- (8.758,13.953);
\draw[PVteal,line width=.55pt] (12.347,15.215) -- (10.242,12.047);
\draw[PVteal,line width=.55pt] (6.653,10.785) -- (3.500,8.331);
\draw[PVteal,line width=.55pt] (6.653,10.785) -- (8.758,13.953);
\draw[PVteal,line width=.55pt] (6.653,10.785) -- (10.242,12.047);
\draw[PVteal,line width=.55pt] (8.758,13.953) -- (10.242,12.047);
\draw[draw=PVnavy,fill=white,line width=.7pt] (15.500,17.669) circle (.90);
\fill[PVteal] (12.347,15.215) circle (.65);
\fill[PVteal] (6.653,10.785) circle (.65);
\fill[PVteal] (3.500,8.331) circle (.65);
\fill[PVteal] (8.758,13.953) circle (.65);
\fill[PVteal] (10.242,12.047) circle (.65);
\node[text=PVteal,font=\fontsize{7.5}{9}\selectfont\bfseries] at (9.500,5.000) {115 / H};
% atlas_124, heldout, E=7; inlet is canonical node 0.
\path[draw=PVteal!25,fill=PVteal!3,rounded corners=.8mm,line width=.35pt] (20.000,3.200) rectangle (37.000,20.500);
\draw[PVteal,line width=.55pt] (29.480,11.651) -- (26.485,14.464);
\draw[PVteal,line width=.55pt] (29.480,11.651) -- (30.592,14.586);
\draw[PVteal,line width=.55pt] (29.480,11.651) -- (33.260,10.026);
\draw[PVteal,line width=.55pt] (26.485,14.464) -- (30.592,14.586);
\draw[PVteal,line width=.55pt] (26.485,14.464) -- (22.500,15.974);
\draw[PVteal,line width=.55pt] (30.592,14.586) -- (34.500,13.298);
\draw[PVteal,line width=.55pt] (34.500,13.298) -- (33.260,10.026);
\draw[draw=PVnavy,fill=white,line width=.7pt] (29.480,11.651) circle (.90);
\fill[PVteal] (26.485,14.464) circle (.65);
\fill[PVteal] (30.592,14.586) circle (.65);
\fill[PVteal] (34.500,13.298) circle (.65);
\fill[PVteal] (33.260,10.026) circle (.65);
\fill[PVteal] (22.500,15.974) circle (.65);
\node[text=PVteal,font=\fontsize{7.5}{9}\selectfont\bfseries] at (28.500,5.000) {124 / H};
% atlas_119, heldout, E=7; inlet is canonical node 0.
\path[draw=PVteal!25,fill=PVteal!3,rounded corners=.8mm,line width=.35pt] (39.000,3.200) rectangle (56.000,20.500);
\draw[PVteal,line width=.55pt] (45.655,14.162) -- (42.420,17.131);
\draw[PVteal,line width=.55pt] (45.655,14.162) -- (45.514,10.209);
\draw[PVteal,line width=.55pt] (45.655,14.162) -- (48.996,10.320);
\draw[PVteal,line width=.55pt] (45.655,14.162) -- (45.142,18.500);
\draw[PVteal,line width=.55pt] (42.420,17.131) -- (45.142,18.500);
\draw[PVteal,line width=.55pt] (45.514,10.209) -- (48.996,10.320);
\draw[PVteal,line width=.55pt] (48.996,10.320) -- (52.580,7.500);
\draw[draw=PVnavy,fill=white,line width=.7pt] (45.655,14.162) circle (.90);
\fill[PVteal] (42.420,17.131) circle (.65);
\fill[PVteal] (45.514,10.209) circle (.65);
\fill[PVteal] (48.996,10.320) circle (.65);
\fill[PVteal] (45.142,18.500) circle (.65);
\fill[PVteal] (52.580,7.500) circle (.65);
\node[text=PVteal,font=\fontsize{7.5}{9}\selectfont\bfseries] at (47.500,5.000) {119 / H};
\end{tikzpicture}
\end{checkedfigure}
\caption{The complete fixed catalogue of 38 connected, pairwise nonisomorphic, six-node graphs with five, six, or seven edges. Colours and letter tags identify the 24 seen structures (S) and 14 held-out structures (H); numbers are NetworkX graph-atlas indices. An outlined node marks the inlet. Edge orientation is only a sign convention and is omitted here. The held-out structures are not used in model fitting or hyperparameter selection. These are abstract circuit topologies, not reconstructed vascular anatomies.}
\label{fig:structure-catalogue}
\end{figure}
