## Supplementary Data - Compact audit, summary tables, code, and figure sources for "Panvascular-JEPA: Unseen-Structure Evaluation of Factorized Predictive States for Hemodynamic Networks": result_figures.tex

\begin{figure}[!tp]
\centering
\begin{checkedfigure}
\begin{tikzpicture}[x=1mm,y=1mm,font=\fontsize{8}{10}\selectfont,text=PVnavy,line cap=round,line join=round]
\path[use as bounding box] (0,0) rectangle (152,143);
\node[anchor=west,font=\fontsize{9}{11}\selectfont\bfseries] at (1,140) {A\quad Mono JEPA};
\begin{axis}[at={(17mm,94mm)},anchor=south west,width=55mm,height=37mm,scale only axis,font=\fontsize{8}{10}\selectfont,tick label style={font=\fontsize{8}{9.5}\selectfont},label style={font=\fontsize{8}{10}\selectfont},axis line style={PVgray!65,line width=.45pt},tick style={PVgray!65},grid=major,grid style={PVgray!15,line width=.3pt},enlarge x limits=false,enlarge y limits=false,clip=true,ymode=log,ymin=100000,ymax=1e+33,xmin=.5,xmax=6.5,xtick={1,2,3,4,5,6},xlabel={Candidate},ylabel={Validation RMSE}]
\addplot[only marks,mark=*,mark size=2pt,color=PVblue] coordinates {(1,3.119042121e+17)};
\addplot[only marks,mark=*,mark size=2pt,color=PVblue] coordinates {(2,3.054390822e+17)};
\addplot[only marks,mark=*,mark size=2pt,color=PVblue] coordinates {(3,1.392204772e+18)};
\addplot[only marks,mark=*,mark size=2pt,color=PVblue] coordinates {(4,1.227001277e+18)};
\addplot[only marks,mark=*,mark size=2pt,color=PVblue] coordinates {(5,1.263253326e+32)};
\addplot[only marks,mark=*,mark size=2pt,color=PVblue] coordinates {(6,1.262984456e+32)};
\addplot[only marks,mark=o,mark size=3.7pt,color=PVnavy,line width=.8pt] coordinates {(2,3.054390822e+17)};
\end{axis}
\node[anchor=west,font=\fontsize{9}{11}\selectfont\bfseries] at (79,140) {B\quad OPF ($K=1$)};
\begin{axis}[at={(95mm,94mm)},anchor=south west,width=55mm,height=37mm,scale only axis,font=\fontsize{8}{10}\selectfont,tick label style={font=\fontsize{8}{9.5}\selectfont},label style={font=\fontsize{8}{10}\selectfont},axis line style={PVgray!65,line width=.45pt},tick style={PVgray!65},grid=major,grid style={PVgray!15,line width=.3pt},enlarge x limits=false,enlarge y limits=false,clip=true,ymode=log,ymin=100000,ymax=1e+33,xmin=.5,xmax=6.5,xtick={1,2,3,4,5,6},xlabel={Candidate},ylabel={Validation RMSE}]
\addplot[only marks,mark=*,mark size=2pt,color=PVorange] coordinates {(1,170157075.2)};
\addplot[only marks,mark=*,mark size=2pt,color=PVorange] coordinates {(2,200201905.3)};
\addplot[only marks,mark=*,mark size=2pt,color=PVorange] coordinates {(3,7.533648845e+11)};
\addplot[only marks,mark=*,mark size=2pt,color=PVorange] coordinates {(4,8.417041133e+11)};
\addplot[only marks,mark=*,mark size=2pt,color=PVorange] coordinates {(5,222864991.4)};
\addplot[only marks,mark=*,mark size=2pt,color=PVorange] coordinates {(6,138445639.8)};
\addplot[only marks,mark=o,mark size=3.7pt,color=PVnavy,line width=.8pt] coordinates {(6,138445639.8)};
\end{axis}
\node[anchor=west,font=\fontsize{9}{11}\selectfont\bfseries] at (1,87) {C\quad OPF ($K=4$)};
\begin{axis}[at={(17mm,41mm)},anchor=south west,width=55mm,height=37mm,scale only axis,font=\fontsize{8}{10}\selectfont,tick label style={font=\fontsize{8}{9.5}\selectfont},label style={font=\fontsize{8}{10}\selectfont},axis line style={PVgray!65,line width=.45pt},tick style={PVgray!65},grid=major,grid style={PVgray!15,line width=.3pt},enlarge x limits=false,enlarge y limits=false,clip=true,ymode=log,ymin=100000,ymax=1e+33,xmin=.5,xmax=6.5,xtick={1,2,3,4,5,6},xlabel={Candidate},ylabel={Validation RMSE}]
\addplot[only marks,mark=*,mark size=2pt,color=PVred] coordinates {(1,5.371133589e+11)};
\addplot[only marks,mark=*,mark size=2pt,color=PVred] coordinates {(2,5.283575136e+11)};
\addplot[only marks,mark=*,mark size=2pt,color=PVred] coordinates {(3,1.428122319e+11)};
\addplot[only marks,mark=*,mark size=2pt,color=PVred] coordinates {(4,3.827437136e+10)};
\addplot[only marks,mark=*,mark size=2pt,color=PVred] coordinates {(5,7.335326466e+14)};
\addplot[only marks,mark=*,mark size=2pt,color=PVred] coordinates {(6,7.915623314e+14)};
\addplot[only marks,mark=o,mark size=3.7pt,color=PVnavy,line width=.8pt] coordinates {(4,3.827437136e+10)};
\end{axis}
\node[anchor=west,font=\fontsize{9}{11}\selectfont\bfseries] at (79,87) {D\quad Supervised GNN};
\begin{axis}[at={(95mm,41mm)},anchor=south west,width=55mm,height=37mm,scale only axis,font=\fontsize{8}{10}\selectfont,tick label style={font=\fontsize{8}{9.5}\selectfont},label style={font=\fontsize{8}{10}\selectfont},axis line style={PVgray!65,line width=.45pt},tick style={PVgray!65},grid=major,grid style={PVgray!15,line width=.3pt},enlarge x limits=false,enlarge y limits=false,clip=true,ymode=log,ymin=100000,ymax=1e+33,xmin=.5,xmax=6.5,xtick={1,2,3,4,5,6},xlabel={Candidate},ylabel={Validation RMSE}]
\addplot[only marks,mark=*,mark size=2pt,color=PVteal] coordinates {(1,746154.6066)};
\addplot[only marks,mark=*,mark size=2pt,color=PVteal] coordinates {(2,729814.4271)};
\addplot[only marks,mark=*,mark size=2pt,color=PVteal] coordinates {(3,248731793.6)};
\addplot[only marks,mark=*,mark size=2pt,color=PVteal] coordinates {(4,241725097.6)};
\addplot[only marks,mark=*,mark size=2pt,color=PVteal] coordinates {(5,2.546461451e+10)};
\addplot[only marks,mark=*,mark size=2pt,color=PVteal] coordinates {(6,1.298547572e+10)};
\addplot[only marks,mark=o,mark size=3.7pt,color=PVnavy,line width=.8pt] coordinates {(2,729814.4271)};
\end{axis}
\node[align=center,text width=150mm,font=\fontsize{8}{10}\selectfont] at (76,18) {Candidates 1/2: learning rate $3\!\times\!10^{-4}$; 3/4: $10^{-3}$; 5/6: $3\!\times\!10^{-3}$.\\Odd candidates: weight decay 0; even candidates: $10^{-4}$.\\Outlined points were selected using validation data only.};
\end{tikzpicture}
\end{checkedfigure}
\caption{Equal-budget development search. Each architecture used the same six learning-rate/weight-decay combinations, 6,000 updates per candidate and development seed 101. The vertical log scale is identical across panels and displays the complete finite range. Crosses, if present, denote nonfinite or failed candidates rather than omitted trials. Selection uses the 96-step equal-structure macro-RMSE on new parameter backgrounds from seen structures; these are not held-out-structure test results.}
\label{fig:validation-search}
\end{figure}

\begin{figure}[!tp]
\centering
\begin{checkedfigure}
\begin{tikzpicture}[x=1mm,y=1mm,font=\fontsize{8}{10}\selectfont,text=PVnavy,line cap=round,line join=round]
\path[use as bounding box] (0,0) rectangle (152,189);
\node[anchor=west,font=\fontsize{9}{11}\selectfont\bfseries] at (0.5,185) {A\quad Forecast error on seen and unseen structures};
\begin{axis}[at={(35mm,112mm)},anchor=south west,width=111mm,height=64mm,scale only axis,font=\fontsize{8}{10}\selectfont,tick label style={font=\fontsize{8}{9.5}\selectfont},label style={font=\fontsize{8}{10}\selectfont},axis line style={PVgray!65,line width=.45pt},tick style={PVgray!65},grid=major,grid style={PVgray!15,line width=.3pt},enlarge x limits=false,enlarge y limits=false,clip=true,xmode=log,xmin=1e-05,xmax=1e+33,ymin=.45,ymax=9.55,ytick={1,2,3,4,5,6,7,8,9},yticklabels={{Persistence},{Fitted RLC},{Reduced RC},{RLC, 4 substeps},{RLC, 1 substep},{Supervised GNN},{OPF ($K=4$)},{OPF ($K=1$)},{Mono JEPA}},xlabel={96-step standardized RMSE},ylabel={},yticklabel style={align=right},]
\addplot[only marks,mark=*,mark size=1pt,color=PVblue,opacity=.35] coordinates {(2523.324543,9.14)};
\addplot[only marks,mark=*,mark size=1pt,color=PVblue,opacity=.35] coordinates {(4550.798941,9.14)};
\addplot[only marks,mark=*,mark size=1pt,color=PVblue,opacity=.35] coordinates {(328217.8863,9.14)};
\addplot[only marks,mark=*,mark size=1pt,color=PVblue,opacity=.35] coordinates {(4.438128778e+12,9.14)};
\addplot[only marks,mark=*,mark size=1pt,color=PVblue,opacity=.35] coordinates {(525.5671867,9.14)};
\addplot[only marks,mark=*,mark size=2.3pt,color=PVblue] coordinates {(8.876258228e+11,9.14)};
\addplot[only marks,mark=square*,mark size=1pt,color=PVteal,opacity=.35] coordinates {(2583.477159,8.86)};
\addplot[only marks,mark=square*,mark size=1pt,color=PVteal,opacity=.35] coordinates {(4280.728721,8.86)};
\addplot[only marks,mark=square*,mark size=1pt,color=PVteal,opacity=.35] coordinates {(406084.3488,8.86)};
\addplot[only marks,mark=square*,mark size=1pt,color=PVteal,opacity=.35] coordinates {(7.449460288e+12,8.86)};
\addplot[only marks,mark=square*,mark size=1pt,color=PVteal,opacity=.35] coordinates {(540.2915151,8.86)};
\addplot[only marks,mark=square*,mark size=2.3pt,color=PVteal] coordinates {(1.48989214e+12,8.86)};
\addplot[only marks,mark=*,mark size=1pt,color=PVblue,opacity=.35] coordinates {(424.423425,8.14)};
\addplot[only marks,mark=*,mark size=1pt,color=PVblue,opacity=.35] coordinates {(249.836235,8.14)};
\addplot[only marks,mark=*,mark size=1pt,color=PVblue,opacity=.35] coordinates {(784.024433,8.14)};
\addplot[only marks,mark=*,mark size=1pt,color=PVblue,opacity=.35] coordinates {(15.79991692,8.14)};
\addplot[only marks,mark=*,mark size=1pt,color=PVblue,opacity=.35] coordinates {(61.58474568,8.14)};
\addplot[only marks,mark=*,mark size=2.3pt,color=PVblue] coordinates {(307.1337511,8.14)};
\addplot[only marks,mark=square*,mark size=1pt,color=PVteal,opacity=.35] coordinates {(462.383624,7.86)};
\addplot[only marks,mark=square*,mark size=1pt,color=PVteal,opacity=.35] coordinates {(122.4983545,7.86)};
\addplot[only marks,mark=square*,mark size=1pt,color=PVteal,opacity=.35] coordinates {(974.8022446,7.86)};
\addplot[only marks,mark=square*,mark size=1pt,color=PVteal,opacity=.35] coordinates {(16.31660116,7.86)};
\addplot[only marks,mark=square*,mark size=1pt,color=PVteal,opacity=.35] coordinates {(71.31534225,7.86)};
\addplot[only marks,mark=square*,mark size=2.3pt,color=PVteal] coordinates {(329.4632333,7.86)};
\addplot[only marks,mark=*,mark size=1pt,color=PVblue,opacity=.35] coordinates {(49511.49121,7.14)};
\addplot[only marks,mark=*,mark size=1pt,color=PVblue,opacity=.35] coordinates {(52975.44567,7.14)};
\addplot[only marks,mark=*,mark size=1pt,color=PVblue,opacity=.35] coordinates {(477447.2433,7.14)};
\addplot[only marks,mark=*,mark size=1pt,color=PVblue,opacity=.35] coordinates {(1715420013,7.14)};
\addplot[only marks,mark=*,mark size=1pt,color=PVblue,opacity=.35] coordinates {(1185333.115,7.14)};
\addplot[only marks,mark=*,mark size=2.3pt,color=PVblue] coordinates {(343437056.1,7.14)};
\addplot[only marks,mark=square*,mark size=1pt,color=PVteal,opacity=.35] coordinates {(44727.84261,6.86)};
\addplot[only marks,mark=square*,mark size=1pt,color=PVteal,opacity=.35] coordinates {(127701.2032,6.86)};
\addplot[only marks,mark=square*,mark size=1pt,color=PVteal,opacity=.35] coordinates {(633656.6517,6.86)};
\addplot[only marks,mark=square*,mark size=1pt,color=PVteal,opacity=.35] coordinates {(4.227795057e+10,6.86)};
\addplot[only marks,mark=square*,mark size=1pt,color=PVteal,opacity=.35] coordinates {(462784.8946,6.86)};
\addplot[only marks,mark=square*,mark size=2.3pt,color=PVteal] coordinates {(8455843888,6.86)};
\addplot[only marks,mark=*,mark size=1pt,color=PVblue,opacity=.35] coordinates {(4.275598159e+22,6.14)};
\addplot[only marks,mark=*,mark size=1pt,color=PVblue,opacity=.35] coordinates {(2.548655598e+10,6.14)};
\addplot[only marks,mark=*,mark size=1pt,color=PVblue,opacity=.35] coordinates {(3.448023142e+31,6.14)};
\addplot[only marks,mark=*,mark size=1pt,color=PVblue,opacity=.35] coordinates {(1.510503594e+21,6.14)};
\addplot[only marks,mark=*,mark size=1pt,color=PVblue,opacity=.35] coordinates {(1.270151428e+14,6.14)};
\addplot[only marks,mark=*,mark size=2.3pt,color=PVblue] coordinates {(6.896046292e+30,6.14)};
\addplot[only marks,mark=square*,mark size=1pt,color=PVteal,opacity=.35] coordinates {(1.122494355e+22,5.86)};
\addplot[only marks,mark=square*,mark size=1pt,color=PVteal,opacity=.35] coordinates {(4.005651432e+10,5.86)};
\addplot[only marks,mark=square*,mark size=1pt,color=PVteal,opacity=.35] coordinates {(9.142451731e+31,5.86)};
\addplot[only marks,mark=square*,mark size=1pt,color=PVteal,opacity=.35] coordinates {(3.129657166e+21,5.86)};
\addplot[only marks,mark=square*,mark size=1pt,color=PVteal,opacity=.35] coordinates {(2.629926534e+14,5.86)};
\addplot[only marks,mark=square*,mark size=2.3pt,color=PVteal] coordinates {(1.828490347e+31,5.86)};
\addplot[only marks,mark=*,mark size=1pt,color=PVblue,opacity=.35] coordinates {(0.001797032968,5.14)};
\addplot[only marks,mark=*,mark size=2.3pt,color=PVblue] coordinates {(0.001797032968,5.14)};
\addplot[only marks,mark=square*,mark size=1pt,color=PVteal,opacity=.35] coordinates {(0.001729890833,4.86)};
\addplot[only marks,mark=square*,mark size=2.3pt,color=PVteal] coordinates {(0.001729890833,4.86)};
\addplot[only marks,mark=*,mark size=1pt,color=PVblue,opacity=.35] coordinates {(0.0001131492736,4.14)};
\addplot[only marks,mark=*,mark size=2.3pt,color=PVblue] coordinates {(0.0001131492736,4.14)};
\addplot[only marks,mark=square*,mark size=1pt,color=PVteal,opacity=.35] coordinates {(0.0001089433117,3.86)};
\addplot[only marks,mark=square*,mark size=2.3pt,color=PVteal] coordinates {(0.0001089433117,3.86)};
\addplot[only marks,mark=*,mark size=1pt,color=PVblue,opacity=.35] coordinates {(0.03380480491,3.14)};
\addplot[only marks,mark=*,mark size=2.3pt,color=PVblue] coordinates {(0.03380480491,3.14)};
\addplot[only marks,mark=square*,mark size=1pt,color=PVteal,opacity=.35] coordinates {(0.03031837496,2.86)};
\addplot[only marks,mark=square*,mark size=2.3pt,color=PVteal] coordinates {(0.03031837496,2.86)};
\addplot[only marks,mark=*,mark size=1pt,color=PVblue,opacity=.35] coordinates {(0.0007034757244,2.14)};
\addplot[only marks,mark=*,mark size=2.3pt,color=PVblue] coordinates {(0.0007034757244,2.14)};
\addplot[only marks,mark=square*,mark size=1pt,color=PVteal,opacity=.35] coordinates {(0.0006773440697,1.86)};
\addplot[only marks,mark=square*,mark size=2.3pt,color=PVteal] coordinates {(0.0006773440697,1.86)};
\addplot[only marks,mark=*,mark size=1pt,color=PVblue,opacity=.35] coordinates {(0.2935819154,1.14)};
\addplot[only marks,mark=*,mark size=2.3pt,color=PVblue] coordinates {(0.2935819154,1.14)};
\addplot[only marks,mark=square*,mark size=1pt,color=PVteal,opacity=.35] coordinates {(0.2845136609,0.86)};
\addplot[only marks,mark=square*,mark size=2.3pt,color=PVteal] coordinates {(0.2845136609,0.86)};
\end{axis}
\fill[PVblue] (43,181.5) circle (.75);\node[anchor=west] at (45,181.5) {Seen structures};
\path[fill=PVteal] (101,180.8) rectangle (102.4,182.2);\node[anchor=west] at (104,181.5) {Unseen structures};
\node[anchor=west,font=\fontsize{9}{11}\selectfont\bfseries] at (0.5,94) {B\quad Paired OPF difference for each held-out structure};
\begin{axis}[at={(22mm,18mm)},anchor=south west,width=124mm,height=68mm,scale only axis,font=\fontsize{8}{10}\selectfont,tick label style={font=\fontsize{8}{9.5}\selectfont},label style={font=\fontsize{8}{10}\selectfont},axis line style={PVgray!65,line width=.45pt},tick style={PVgray!65},grid=major,grid style={PVgray!15,line width=.3pt},enlarge x limits=false,enlarge y limits=false,clip=true,xmin=-14.150000000000004,xmax=14.150000000000004,ymin=.5,ymax=14.5,ytick={1,2,3,4,5,6,7,8,9,10,11,12,13,14},yticklabels={129,126,124,119,118,115,114,99,96,94,93,92,83,79},xtick={-14,-12,-8.000000004,-4.000043427,0,4.000043427,8.000000004,12,14},xticklabels={$-10^{12}$,$-10^{10}$,$-10^{6}$,$-10^{2}$,0,$10^{2}$,$10^{6}$,$10^{10}$,$10^{12}$},xlabel={OPF ($K=4$) minus OPF ($K=1$): standardized RMSE},ylabel={Atlas ID}]
\addplot[color=PVgray,dashed,mark=none,line width=.7pt] coordinates {(0,0.5) (0,14.5)};
\addplot[color=PVred,mark=none,line width=.85pt] coordinates {(0,14) (6.699337625,14)};
\addplot[only marks,color=PVred,mark=*,mark size=2pt] coordinates {(6.699337625,14)};
\addplot[color=PVred,mark=none,line width=.85pt] coordinates {(0,13) (6.669875229,13)};
\addplot[only marks,color=PVred,mark=*,mark size=2pt] coordinates {(6.669875229,13)};
\addplot[color=PVred,mark=none,line width=.85pt] coordinates {(0,12) (13.07326786,12)};
\addplot[only marks,color=PVred,mark=*,mark size=2pt] coordinates {(13.07326786,12)};
\addplot[color=PVred,mark=none,line width=.85pt] coordinates {(0,11) (7.484013489,11)};
\addplot[only marks,color=PVred,mark=*,mark size=2pt] coordinates {(7.484013489,11)};
\addplot[color=PVred,mark=none,line width=.85pt] coordinates {(0,10) (8.054009218,10)};
\addplot[only marks,color=PVred,mark=*,mark size=2pt] coordinates {(8.054009218,10)};
\addplot[color=PVred,mark=none,line width=.85pt] coordinates {(0,9) (7.409605754,9)};
\addplot[only marks,color=PVred,mark=*,mark size=2pt] coordinates {(7.409605754,9)};
\addplot[color=PVred,mark=none,line width=.85pt] coordinates {(0,8) (6.942768001,8)};
\addplot[only marks,color=PVred,mark=*,mark size=2pt] coordinates {(6.942768001,8)};
\addplot[color=PVred,mark=none,line width=.85pt] coordinates {(0,7) (7.836213281,7)};
\addplot[only marks,color=PVred,mark=*,mark size=2pt] coordinates {(7.836213281,7)};
\addplot[color=PVred,mark=none,line width=.85pt] coordinates {(0,6) (8.114078283,6)};
\addplot[only marks,color=PVred,mark=*,mark size=2pt] coordinates {(8.114078283,6)};
\addplot[color=PVred,mark=none,line width=.85pt] coordinates {(0,5) (6.993011184,5)};
\addplot[only marks,color=PVred,mark=*,mark size=2pt] coordinates {(6.993011184,5)};
\addplot[color=PVred,mark=none,line width=.85pt] coordinates {(0,4) (7.627278551,4)};
\addplot[only marks,color=PVred,mark=*,mark size=2pt] coordinates {(7.627278551,4)};
\addplot[color=PVred,mark=none,line width=.85pt] coordinates {(0,3) (6.910670817,3)};
\addplot[only marks,color=PVred,mark=*,mark size=2pt] coordinates {(6.910670817,3)};
\addplot[color=PVred,mark=none,line width=.85pt] coordinates {(0,2) (7.133540682,2)};
\addplot[only marks,color=PVred,mark=*,mark size=2pt] coordinates {(7.133540682,2)};
\addplot[color=PVred,mark=none,line width=.85pt] coordinates {(0,1) (6.792060018,1)};
\addplot[only marks,color=PVred,mark=*,mark size=2pt] coordinates {(6.792060018,1)};
\end{axis}
\end{tikzpicture}
\end{checkedfigure}
\caption{Primary and structure-specific comparisons. (A) Large points are means across the five fixed neural training seeds; small points show the individual seeds. Deterministic comparators have one result. Both axes include the complete finite range without clipping divergent runs. (B) Differences are computed after averaging seeds within each held-out structure; positive values favour $K=1$. To retain both sign and large magnitudes, horizontal coordinates use $\operatorname{sign}(d)\log_{10}(1+|d|/0.01)$, with ticks labelled in original RMSE units. These are paired differences for the fixed catalogue, not fourteen independent patient cohorts.}
\label{fig:primary-structure-results}
\end{figure}

\begin{figure}[!tp]
\centering
\begin{checkedfigure}
\begin{tikzpicture}[x=1mm,y=1mm,font=\fontsize{8}{10}\selectfont,text=PVnavy,line cap=round,line join=round]
\path[use as bounding box] (0,0) rectangle (152,182);
\node[anchor=west,font=\fontsize{9}{11}\selectfont\bfseries] at (1,178) {A\quad Seen structures};
\begin{axis}[at={(17mm,129mm)},anchor=south west,width=55mm,height=40mm,scale only axis,font=\fontsize{8}{10}\selectfont,tick label style={font=\fontsize{8}{9.5}\selectfont},label style={font=\fontsize{8}{10}\selectfont},axis line style={PVgray!65,line width=.45pt},tick style={PVgray!65},grid=major,grid style={PVgray!15,line width=.3pt},enlarge x limits=false,enlarge y limits=false,clip=true,ymode=log,ymin=1e-05,ymax=1e+33,xmin=0,xmax=3,xtick={0,1,2,3},xlabel={Forecast cycles},ylabel={Horizon RMSE}]
\addplot[color=PVblue,solid,line width=.75pt,mark=none] coordinates {(0.03125,0.06111036558) (0.0625,0.1233491235) (0.09375,0.1855899355) (0.125,0.2458736872) (0.15625,0.302802619) (0.1875,0.3556723366) (0.21875,0.4045775119) (0.25,0.4502767713) (0.28125,0.4939911221) (0.3125,0.5373134953) (0.34375,0.5823566228) (0.375,0.6317863154) (0.40625,0.6886129504) (0.4375,0.7558616434) (0.46875,0.8364145371) (0.5,0.9331637038) (0.53125,1.049811382) (0.5625,1.191456129) (0.59375,1.364945627) (0.625,1.5796913) (0.65625,1.848662782) (0.6875,2.190190997) (0.71875,2.630503463) (0.75,3.207002994) (0.78125,3.972806756) (0.8125,5.002573233) (0.84375,6.400169333) (0.875,8.309870502) (0.90625,10.93391825) (0.9375,14.55930833) (0.96875,19.59647483) (1,26.63616382) (1.03125,36.53337603) (1.0625,50.53159343) (1.09375,70.44497368) (1.125,98.92674342) (1.15625,139.8660576) (1.1875,198.9910477) (1.21875,284.7567845) (1.25,409.6774203) (1.28125,592.3247907) (1.3125,860.3158573) (1.34375,1254.827205) (1.375,1837.400016) (1.40625,2700.219868) (1.4375,3981.624823) (1.46875,5889.668687) (1.5,8737.801267) (1.53125,12999.04895) (1.5625,19388.40036) (1.59375,28988.22607) (1.625,43439.45042) (1.65625,65233.15403) (1.6875,98155.94696) (1.71875,147970.941) (1.75,223459.2895) (1.78125,338015.3671) (1.8125,512091.7938) (1.84375,776948.8088) (1.875,1180407.174) (1.90625,1795690.579) (1.9375,2735007.362) (1.96875,4170432.631) (2,6366044.855) (2.03125,9727392.304) (2.0625,14877688.34) (2.09375,22775230.01) (2.125,34894412.72) (2.15625,53504949.66) (2.1875,82102714.74) (2.21875,126074850.2) (2.25,193726840.7) (2.28125,297869286) (2.3125,458270050.3) (2.34375,705445069.2) (2.375,1086521223) (2.40625,1674304124) (2.4375,2581310879) (2.46875,3981489321) (2.5,6143846416) (2.53125,9484522092) (2.5625,1.464745797e+10) (2.59375,2.262937709e+10) (2.625,3.497348533e+10) (2.65625,5.406972575e+10) (2.6875,8.362026358e+10) (2.71875,1.293614301e+11) (2.75,2.001833839e+11) (2.78125,3.098672067e+11) (2.8125,4.797802185e+11) (2.84375,7.430588625e+11) (2.875,1.151101189e+12) (2.90625,1.783646732e+12) (2.9375,2.764426145e+12) (2.96875,4.285465658e+12) (3,6.644833651e+12)};
\addplot[color=PVorange,solid,line width=.75pt,mark=none] coordinates {(0.03125,0.04533580864) (0.0625,0.07710930758) (0.09375,0.114426801) (0.125,0.1548238614) (0.15625,0.1969663861) (0.1875,0.2392038161) (0.21875,0.2822785891) (0.25,0.3292704293) (0.28125,0.3787774859) (0.3125,0.4311950175) (0.34375,0.4942679889) (0.375,0.5684164381) (0.40625,0.6474803415) (0.4375,0.7338179186) (0.46875,0.8301254679) (0.5,0.9302787619) (0.53125,1.025523462) (0.5625,1.112991108) (0.59375,1.227056351) (0.625,1.393002702) (0.65625,1.582047818) (0.6875,1.753319159) (0.71875,1.939935196) (0.75,2.091804669) (0.78125,2.221881332) (0.8125,2.321134002) (0.84375,2.465139221) (0.875,2.632517756) (0.90625,2.846845311) (0.9375,3.120801459) (0.96875,3.422637382) (1,3.708244936) (1.03125,4.014737537) (1.0625,4.2755579) (1.09375,4.610078553) (1.125,4.973919864) (1.15625,5.48123067) (1.1875,6.065044717) (1.21875,6.450113686) (1.25,7.011674065) (1.28125,7.533078454) (1.3125,7.928000163) (1.34375,8.435983898) (1.375,8.999847452) (1.40625,9.492916191) (1.4375,10.23426909) (1.46875,11.30913364) (1.5,12.75169846) (1.53125,14.04243783) (1.5625,15.49041799) (1.59375,16.69188358) (1.625,18.56245392) (1.65625,21.01952861) (1.6875,22.56066656) (1.71875,24.36540944) (1.75,25.73765081) (1.78125,27.2524769) (1.8125,29.10698914) (1.84375,31.17974631) (1.875,34.13844686) (1.90625,38.0076294) (1.9375,42.74390461) (1.96875,49.24594663) (2,55.37683005) (2.03125,61.39439641) (2.0625,64.63482935) (2.09375,69.25672488) (2.125,77.71328209) (2.15625,82.5557436) (2.1875,90.72448791) (2.21875,98.13176725) (2.25,106.79768) (2.28125,116.4930291) (2.3125,126.3993798) (2.34375,138.533115) (2.375,151.2403674) (2.40625,166.8014407) (2.4375,192.5080088) (2.46875,219.1766286) (2.5,254.6540923) (2.53125,285.4824636) (2.5625,312.2343853) (2.59375,347.9297939) (2.625,373.821899) (2.65625,404.7413809) (2.6875,439.1885422) (2.71875,483.6302457) (2.75,530.2120527) (2.78125,581.3191566) (2.8125,637.6267785) (2.84375,702.5720266) (2.875,776.8983498) (2.90625,881.9135935) (2.9375,1028.605163) (2.96875,1156.519326) (3,1334.430576)};
\addplot[color=PVred,solid,line width=.75pt,mark=none] coordinates {(0.03125,0.04467488212) (0.0625,0.09150637223) (0.09375,0.1397331577) (0.125,0.187945141) (0.15625,0.2363618511) (0.1875,0.2861142948) (0.21875,0.3388546852) (0.25,0.3962245563) (0.28125,0.4594496228) (0.3125,0.5293724331) (0.34375,0.6066547592) (0.375,0.6919975016) (0.40625,0.7863154845) (0.4375,0.8908238246) (0.46875,1.006967346) (0.5,1.13621667) (0.53125,1.280087334) (0.5625,1.440221843) (0.59375,1.618650208) (0.625,1.817994195) (0.65625,2.041633821) (0.6875,2.29384778) (0.71875,2.579910938) (0.75,2.906343999) (0.78125,3.280901207) (0.8125,3.710221924) (0.84375,4.200644113) (0.875,4.760655183) (0.90625,5.399672206) (0.9375,6.127756933) (0.96875,6.965349504) (1,7.941402567) (1.03125,9.082672002) (1.0625,10.41198917) (1.09375,11.95821534) (1.125,13.75901312) (1.15625,15.85031931) (1.1875,18.27056238) (1.21875,21.07252906) (1.25,24.33892897) (1.28125,28.17118155) (1.3125,32.70867436) (1.34375,38.11381903) (1.375,44.60425428) (1.40625,52.4509716) (1.4375,61.99252166) (1.46875,73.70094151) (1.5,88.16260107) (1.53125,106.2045931) (1.5625,129.0236024) (1.59375,158.262475) (1.625,196.1326897) (1.65625,245.585515) (1.6875,311.0028074) (1.71875,398.3871341) (1.75,516.1075723) (1.78125,676.5130604) (1.8125,897.7419756) (1.84375,1206.494642) (1.875,1642.447514) (1.90625,2264.588562) (1.9375,3159.774475) (1.96875,4454.076165) (2,6335.531998) (2.03125,9083.989122) (2.0625,13115.73477) (2.09375,19053.70352) (2.125,27833.41581) (2.15625,40859.71838) (2.1875,60245.51495) (2.21875,89171.95017) (2.25,132432.1962) (2.28125,197249.5835) (2.3125,294525.0082) (2.34375,440712.2519) (2.375,660664.7575) (2.40625,991923.2413) (2.4375,1491199.666) (2.46875,2244225.57) (2.5,3380625.473) (2.53125,5096469.255) (2.5625,7688386.36) (2.59375,11605247.5) (2.625,17526462.63) (2.65625,26480592.05) (2.6875,40025069.73) (2.71875,60518582.88) (2.75,91533880.35) (2.78125,138483033) (2.8125,209565337) (2.84375,317204825.1) (2.875,480229499.1) (2.90625,727175197.4) (2.9375,1101295067) (2.96875,1668157669) (3,2527168609)};
\addplot[color=PVteal,solid,line width=.75pt,mark=none] coordinates {(0.03125,0.01993558517) (0.0625,0.03735434488) (0.09375,0.05234216328) (0.125,0.08932006535) (0.15625,0.1263141327) (0.1875,0.2077782159) (0.21875,0.443242457) (0.25,0.6285493149) (0.28125,1.045186867) (0.3125,1.837496576) (0.34375,2.55455204) (0.375,3.581354306) (0.40625,5.585944617) (0.4375,9.254595595) (0.46875,16.12059509) (0.5,30.24185828) (0.53125,60.80092551) (0.5625,128.1945038) (0.59375,279.2326572) (0.625,621.9035933) (0.65625,1407.002504) (0.6875,3219.465129) (0.71875,7429.945886) (0.75,17261.52421) (0.78125,40315.65981) (0.8125,94567.62211) (0.84375,222619.857) (0.875,525640.7935) (0.90625,1244288.57) (0.9375,2951915.378) (0.96875,7016270.794) (1,16703894.39) (1.03125,39824853.79) (1.0625,95070714.95) (1.09375,227215556.6) (1.125,543600965.8) (1.15625,1301766037) (1.1875,3120046190) (1.21875,7484003540) (1.25,1.796483226e+10) (1.28125,4.315216571e+10) (1.3125,1.037180415e+11) (1.34375,2.494381295e+11) (1.375,6.002257052e+11) (1.40625,1.44509867e+12) (1.4375,3.48096956e+12) (1.46875,8.389046755e+12) (1.5,2.022671589e+13) (1.53125,4.878991123e+13) (1.5625,1.177385668e+14) (1.59375,2.842393232e+14) (1.625,6.864666219e+14) (1.65625,1.658509956e+15) (1.6875,4.008427593e+15) (1.71875,9.691289539e+15) (1.75,2.343878886e+16) (1.78125,5.670608335e+16) (1.8125,1.372334866e+17) (1.84375,3.32216953e+17) (1.875,8.044706153e+17) (1.90625,1.948592297e+18) (1.9375,4.721173825e+18) (1.96875,1.144177038e+19) (2,2.773619399e+19) (2.03125,6.725230477e+19) (2.0625,1.63106238e+20) (2.09375,3.956704417e+20) (2.125,9.600480834e+20) (2.15625,2.329943866e+21) (2.1875,5.655720243e+21) (2.21875,1.373148025e+22) (2.25,3.334501406e+22) (2.28125,8.098893918e+22) (2.3125,1.967429008e+23) (2.34375,4.780225101e+23) (2.375,1.161638535e+24) (2.40625,2.82334921e+24) (2.4375,6.863200643e+24) (2.46875,1.668610693e+25) (2.5,4.057395312e+25) (2.53125,9.867371101e+25) (2.5625,2.400022348e+26) (2.59375,5.838305583e+26) (2.625,1.420411163e+27) (2.65625,3.456170479e+27) (2.6875,8.410624298e+27) (2.71875,2.046970168e+28) (2.75,4.982453668e+28) (2.78125,1.212891194e+29) (2.8125,2.952878784e+29) (2.84375,7.189737554e+29) (2.875,1.750743707e+30) (2.90625,4.263563907e+30) (2.9375,1.038394146e+31) (2.96875,2.529237366e+31) (3,6.161032779e+31)};
\addplot[color=PVgray,dashed,line width=.75pt,mark=none] coordinates {(0.03125,0.0003185730294) (0.0625,0.0006173650842) (0.09375,0.0009317562073) (0.125,0.001237861462) (0.15625,0.001466334621) (0.1875,0.001575230061) (0.21875,0.001554246355) (0.25,0.001421171146) (0.28125,0.001237220715) (0.3125,0.001167864046) (0.34375,0.001233903613) (0.375,0.001267960852) (0.40625,0.001289378092) (0.4375,0.001252936629) (0.46875,0.001220835558) (0.5,0.001227492555) (0.53125,0.001312642154) (0.5625,0.001399382369) (0.59375,0.001446861474) (0.625,0.001447322542) (0.65625,0.001381791944) (0.6875,0.001342478449) (0.71875,0.001317795186) (0.75,0.001343823231) (0.78125,0.001435768604) (0.8125,0.001590230968) (0.84375,0.001723008605) (0.875,0.001774501662) (0.90625,0.001721453652) (0.9375,0.001603013546) (0.96875,0.001458211464) (1,0.001314670086) (1.03125,0.001302768624) (1.0625,0.001573953873) (1.09375,0.001896143613) (1.125,0.002112675247) (1.15625,0.00221724836) (1.1875,0.002174879894) (1.21875,0.00210629485) (1.25,0.002149831304) (1.28125,0.002397504732) (1.3125,0.002519654924) (1.34375,0.002520266796) (1.375,0.002574310897) (1.40625,0.002464046124) (1.4375,0.002466482285) (1.46875,0.002391211373) (1.5,0.002319836299) (1.53125,0.002093511783) (1.5625,0.002045671317) (1.59375,0.002020696348) (1.625,0.001948903334) (1.65625,0.001723481731) (1.6875,0.001589284649) (1.71875,0.001553405072) (1.75,0.001539952288) (1.78125,0.001621031908) (1.8125,0.001814823571) (1.84375,0.002000465661) (1.875,0.002116824839) (1.90625,0.00225068121) (1.9375,0.002193105445) (1.96875,0.002166047689) (2,0.002125002315) (2.03125,0.002124744375) (2.0625,0.002154121455) (2.09375,0.002223420855) (2.125,0.002306540451) (2.15625,0.002342257785) (2.1875,0.002321213504) (2.21875,0.002155509127) (2.25,0.001937337567) (2.28125,0.001740116807) (2.3125,0.00165127348) (2.34375,0.001542892248) (2.375,0.001450856851) (2.40625,0.001334928149) (2.4375,0.001218711117) (2.46875,0.001150949306) (2.5,0.001110527931) (2.53125,0.001090104536) (2.5625,0.001147541468) (2.59375,0.00127331564) (2.625,0.001317096934) (2.65625,0.001300475442) (2.6875,0.001228623421) (2.71875,0.001204798993) (2.75,0.001321513079) (2.78125,0.001567174251) (2.8125,0.001752943657) (2.84375,0.001880775917) (2.875,0.001963726289) (2.90625,0.00195508402) (2.9375,0.001912135598) (2.96875,0.001772134738) (3,0.001598787086)};
\addplot[color=PVnavy,densely dotted,line width=.75pt,mark=none] coordinates {(0.03125,2.429711778e-05) (0.0625,4.41229996e-05) (0.09375,6.441901418e-05) (0.125,8.371586379e-05) (0.15625,9.755885086e-05) (0.1875,0.0001015131535) (0.21875,9.889682811e-05) (0.25,8.893929654e-05) (0.28125,7.707460575e-05) (0.3125,7.387645086e-05) (0.34375,7.763778954e-05) (0.375,7.895375619e-05) (0.40625,7.984429844e-05) (0.4375,7.718383213e-05) (0.46875,7.539563714e-05) (0.5,7.715952283e-05) (0.53125,8.277523462e-05) (0.5625,8.662729951e-05) (0.59375,9.064878501e-05) (0.625,8.902793277e-05) (0.65625,8.586374976e-05) (0.6875,8.286819367e-05) (0.71875,8.234996475e-05) (0.75,8.500098551e-05) (0.78125,9.271810382e-05) (0.8125,0.000104356482) (0.84375,0.0001128259454) (0.875,0.0001141313324) (0.90625,0.0001099522927) (0.9375,0.0001019318209) (0.96875,9.193810205e-05) (1,8.089378922e-05) (1.03125,8.439607377e-05) (1.0625,0.0001032012395) (1.09375,0.0001227059448) (1.125,0.0001356629893) (1.15625,0.0001403953797) (1.1875,0.00013691956) (1.21875,0.0001328681475) (1.25,0.0001396477919) (1.28125,0.0001523208208) (1.3125,0.0001555338869) (1.34375,0.00016151141) (1.375,0.0001559163049) (1.40625,0.000156056713) (1.4375,0.0001510865011) (1.46875,0.0001515287909) (1.5,0.0001396164697) (1.53125,0.0001317891962) (1.5625,0.0001264883768) (1.59375,0.0001274010897) (1.625,0.0001168206463) (1.65625,0.0001046185394) (1.6875,9.94992438e-05) (1.71875,9.61418044e-05) (1.75,9.839950881e-05) (1.78125,0.0001067238087) (1.8125,0.0001194249831) (1.84375,0.000128705538) (1.875,0.0001374629977) (1.90625,0.0001418198717) (1.9375,0.000138638648) (1.96875,0.0001354552747) (2,0.0001328958064) (2.03125,0.0001330766836) (2.0625,0.0001340550002) (2.09375,0.0001411056926) (2.125,0.0001450254693) (2.15625,0.0001490989276) (2.1875,0.000143663817) (2.21875,0.0001338013545) (2.25,0.0001195638815) (2.28125,0.0001095876873) (2.3125,0.0001012996681) (2.34375,9.506175143e-05) (2.375,8.935147185e-05) (2.40625,8.071613937e-05) (2.4375,7.544633767e-05) (2.46875,7.147837125e-05) (2.5,6.799045386e-05) (2.53125,6.675408447e-05) (2.5625,7.277350996e-05) (2.59375,7.937478178e-05) (2.625,8.146687357e-05) (2.65625,7.928466658e-05) (2.6875,7.488948272e-05) (2.71875,7.59924302e-05) (2.75,8.679226958e-05) (2.78125,0.0001017529692) (2.8125,0.0001121019947) (2.84375,0.0001222743352) (2.875,0.0001247788086) (2.90625,0.0001251210786) (2.9375,0.0001191747584) (2.96875,0.0001100271509) (3,9.63142904e-05)};
\addplot[color=PVorange,dashed,line width=.75pt,mark=none] coordinates {(0.03125,0.06015502139) (0.0625,0.04577326949) (0.09375,0.04184209158) (0.125,0.03943819677) (0.15625,0.03749735369) (0.1875,0.03570694207) (0.21875,0.03367223178) (0.25,0.03116240424) (0.28125,0.02851585285) (0.3125,0.02717735289) (0.34375,0.02721340371) (0.375,0.02754802787) (0.40625,0.02762219131) (0.4375,0.02708421444) (0.46875,0.02562428621) (0.5,0.02359705333) (0.53125,0.02217165259) (0.5625,0.02166299548) (0.59375,0.02225871606) (0.625,0.02376365629) (0.65625,0.02548824964) (0.6875,0.02663744031) (0.71875,0.0268838975) (0.75,0.02642470978) (0.78125,0.02579915001) (0.8125,0.02576720828) (0.84375,0.02714680241) (0.875,0.03021214467) (0.90625,0.03437210851) (0.9375,0.03857728398) (0.96875,0.04189589459) (1,0.04380091966) (1.03125,0.0438417001) (1.0625,0.0439830462) (1.09375,0.04548307988) (1.125,0.04736675787) (1.15625,0.04847984222) (1.1875,0.04824823009) (1.21875,0.04632670145) (1.25,0.04313980645) (1.28125,0.03986538185) (1.3125,0.03564018345) (1.34375,0.0310022503) (1.375,0.02699918994) (1.40625,0.02395269743) (1.4375,0.02151083359) (1.46875,0.01958268569) (1.5,0.01828791724) (1.53125,0.01824320054) (1.5625,0.01923159858) (1.59375,0.02082241922) (1.625,0.02285793657) (1.65625,0.0249842853) (1.6875,0.02650049594) (1.71875,0.02697732406) (1.75,0.02656314922) (1.78125,0.02625445167) (1.8125,0.0279144789) (1.84375,0.03138377693) (1.875,0.03561202404) (1.90625,0.04029449084) (1.9375,0.04474167686) (1.96875,0.04782568147) (2,0.04888512944) (2.03125,0.0483135053) (2.0625,0.04648379828) (2.09375,0.04396142525) (2.125,0.04128751771) (2.15625,0.038771282) (2.1875,0.03642846154) (2.21875,0.03404983332) (2.25,0.0314426889) (2.28125,0.02870177554) (2.3125,0.02594672781) (2.34375,0.02338705933) (2.375,0.02123736575) (2.40625,0.01958558343) (2.4375,0.01831074299) (2.46875,0.01718133525) (2.5,0.01628579001) (2.53125,0.01665119385) (2.5625,0.01924244102) (2.59375,0.02265075058) (2.625,0.02584721522) (2.65625,0.02812828316) (2.6875,0.02912799691) (2.71875,0.0289137861) (2.75,0.02809581249) (2.78125,0.02732852627) (2.8125,0.02679837094) (2.84375,0.0275198222) (2.875,0.03020154697) (2.90625,0.03421230867) (2.9375,0.03835499645) (2.96875,0.04170979278) (3,0.04373852854)};
\addplot[color=PVteal,densely dotted,line width=.75pt,mark=none] coordinates {(0.03125,0.0002343125902) (0.0625,0.0003381877851) (0.09375,0.0003599946594) (0.125,0.0003766121742) (0.15625,0.0004015149052) (0.1875,0.0004170925345) (0.21875,0.0004183289478) (0.25,0.0004139755378) (0.28125,0.0004031404299) (0.3125,0.0003866721867) (0.34375,0.0003658504609) (0.375,0.0003402995246) (0.40625,0.0003134557341) (0.4375,0.0002866619051) (0.46875,0.000265925376) (0.5,0.0002558711561) (0.53125,0.00025737382) (0.5625,0.0002703057369) (0.59375,0.0002905472653) (0.625,0.0003161947669) (0.65625,0.0003472927913) (0.6875,0.0003812622069) (0.71875,0.0004177024797) (0.75,0.0004547373623) (0.78125,0.000492291101) (0.8125,0.0005294010888) (0.84375,0.0005650342536) (0.875,0.0005990952375) (0.90625,0.0006314948694) (0.9375,0.0006622406538) (0.96875,0.00069067288) (1,0.0007129127031) (1.03125,0.0007248438902) (1.0625,0.000724831075) (1.09375,0.0007157206943) (1.125,0.0007016768765) (1.15625,0.0006876320357) (1.1875,0.0006758634946) (1.21875,0.0006654209365) (1.25,0.0006559505955) (1.28125,0.0006449924268) (1.3125,0.0006324811171) (1.34375,0.0006207089115) (1.375,0.0006052383984) (1.40625,0.0005898342288) (1.4375,0.0005742336505) (1.46875,0.0005657221662) (1.5,0.0005641533074) (1.53125,0.0005717855606) (1.5625,0.0005867047667) (1.59375,0.0006083877315) (1.625,0.0006355965702) (1.65625,0.0006678704355) (1.6875,0.0007037313282) (1.71875,0.0007419892872) (1.75,0.000781794443) (1.78125,0.0008204673832) (1.8125,0.0008547271623) (1.84375,0.0008831314982) (1.875,0.0009073149269) (1.90625,0.0009282854579) (1.9375,0.0009451050314) (1.96875,0.0009572656607) (2,0.0009636779655) (2.03125,0.0009627092314) (2.0625,0.0009532325681) (2.09375,0.000936001703) (2.125,0.0009132524126) (2.15625,0.0008893984873) (2.1875,0.0008666884494) (2.21875,0.0008463877465) (2.25,0.0008275545111) (2.28125,0.0008093566626) (2.3125,0.0007895498148) (2.34375,0.0007666971306) (2.375,0.000741348739) (2.40625,0.0007159224151) (2.4375,0.0006937828449) (2.46875,0.0006787747169) (2.5,0.0006733280086) (2.53125,0.0006770923543) (2.5625,0.0006883715112) (2.59375,0.0007055322482) (2.625,0.0007282676913) (2.65625,0.0007564783627) (2.6875,0.0007891066988) (2.71875,0.0008247872186) (2.75,0.0008620810245) (2.78125,0.0008986445316) (2.8125,0.0009326097429) (2.84375,0.0009635671587) (2.875,0.0009903575607) (2.90625,0.001012829548) (2.9375,0.001030804526) (2.96875,0.00104439047) (3,0.001051345362)};
\addplot[color=PVgray,dashdotted,line width=.75pt,mark=none] coordinates {(0.03125,0.06116827837) (0.0625,0.1225964637) (0.09375,0.1811051357) (0.125,0.2338790144) (0.15625,0.2788868869) (0.1875,0.3150642877) (0.21875,0.3423321268) (0.25,0.3614778205) (0.28125,0.3739040207) (0.3125,0.3812747766) (0.34375,0.385213351) (0.375,0.3870473537) (0.40625,0.3875948177) (0.4375,0.3870995001) (0.46875,0.3852984814) (0.5,0.3815322642) (0.53125,0.3750490361) (0.5625,0.3655292524) (0.59375,0.3531743259) (0.625,0.3385854003) (0.65625,0.3228440412) (0.6875,0.3073741478) (0.71875,0.2934656426) (0.75,0.2816254647) (0.78125,0.2711831095) (0.8125,0.2604250438) (0.84375,0.2471942769) (0.875,0.2297241644) (0.90625,0.2075771684) (0.9375,0.1827878766) (0.96875,0.1614803984) (1,0.1550549374) (1.03125,0.1716114284) (1.0625,0.203949221) (1.09375,0.2418858815) (1.125,0.2796585267) (1.15625,0.3141439911) (1.1875,0.3434388478) (1.21875,0.3662892756) (1.25,0.3817855679) (1.28125,0.3894468602) (1.3125,0.3903343249) (1.34375,0.3867912734) (1.375,0.3808453786) (1.40625,0.3737197802) (1.4375,0.365930092) (1.46875,0.3575001791) (1.5,0.3480947323) (1.53125,0.3371797938) (1.5625,0.3242918068) (1.59375,0.3093249835) (1.625,0.2927822883) (1.65625,0.2759128308) (1.6875,0.2605387864) (1.71875,0.2483673395) (1.75,0.2400540091) (1.78125,0.2345021892) (1.8125,0.2288023779) (1.84375,0.2197430973) (1.875,0.204829328) (1.90625,0.1823813061) (1.9375,0.1524690428) (1.96875,0.1189456017) (2,0.09476938472) (2.03125,0.102178006) (2.0625,0.1356222146) (2.09375,0.1790101196) (2.125,0.222903242) (2.15625,0.2621977487) (2.1875,0.2944667402) (2.21875,0.3190144107) (2.25,0.3362925089) (2.28125,0.3474657782) (2.3125,0.3540076921) (2.34375,0.3573329587) (2.375,0.3585438139) (2.40625,0.3583265705) (2.4375,0.3569653807) (2.46875,0.3544051396) (2.5,0.350325461) (2.53125,0.3441702765) (2.5625,0.3352424892) (2.59375,0.3231773909) (2.625,0.3083645698) (2.65625,0.2919255248) (2.6875,0.2753869817) (2.71875,0.260071595) (2.75,0.2464018001) (2.78125,0.2335728878) (2.8125,0.2199026005) (2.84375,0.2031647779) (2.875,0.1810015742) (2.90625,0.1517684767) (2.9375,0.1154635043) (2.96875,0.07563758171) (3,0.05045866279)};
\end{axis}
\node[anchor=west,font=\fontsize{9}{11}\selectfont\bfseries] at (79,178) {B\quad Unseen structures};
\begin{axis}[at={(95mm,129mm)},anchor=south west,width=55mm,height=40mm,scale only axis,font=\fontsize{8}{10}\selectfont,tick label style={font=\fontsize{8}{9.5}\selectfont},label style={font=\fontsize{8}{10}\selectfont},axis line style={PVgray!65,line width=.45pt},tick style={PVgray!65},grid=major,grid style={PVgray!15,line width=.3pt},enlarge x limits=false,enlarge y limits=false,clip=true,ymode=log,ymin=1e-05,ymax=1e+33,xmin=0,xmax=3,xtick={0,1,2,3},xlabel={Forecast cycles},ylabel={Horizon RMSE}]
\addplot[color=PVblue,solid,line width=.75pt,mark=none] coordinates {(0.03125,0.06066499152) (0.0625,0.1208971302) (0.09375,0.1807543992) (0.125,0.2385771819) (0.15625,0.2932566737) (0.1875,0.3443628379) (0.21875,0.3922502364) (0.25,0.4378955869) (0.28125,0.4825832337) (0.3125,0.5278729999) (0.34375,0.5759600129) (0.375,0.6296588421) (0.40625,0.6921161144) (0.4375,0.7665313867) (0.46875,0.855981752) (0.5,0.9635330745) (0.53125,1.092944218) (0.5625,1.249311409) (0.59375,1.439616398) (0.625,1.673513618) (0.65625,1.964032864) (0.6875,2.32871941) (0.71875,2.791720707) (0.75,3.387265624) (0.78125,4.164404646) (0.8125,5.192297632) (0.84375,6.566991822) (0.875,8.422398098) (0.90625,10.94836378) (0.9375,14.41683957) (0.96875,19.21858226) (1,25.91954628) (1.03125,35.34722945) (1.0625,48.71579495) (1.09375,67.81133615) (1.125,95.27322592) (1.15625,135.0158066) (1.1875,192.8698483) (1.21875,277.5499788) (1.25,402.1197729) (1.28125,586.2199955) (1.3125,859.4633653) (1.34375,1266.614704) (1.375,1875.512414) (1.40625,2789.185897) (1.4375,4164.405821) (1.46875,6240.181018) (1.5,9381.499353) (1.53125,14146.66516) (1.5625,21390.99659) (1.59375,32426.58043) (1.625,49268.35393) (1.65625,75014.52469) (1.6875,114434.0049) (1.71875,174874.1962) (1.75,267664.8139) (1.78125,410290.8497) (1.8125,629757.6862) (1.84375,967802.8231) (1.875,1488974.968) (1.90625,2293159.885) (1.9375,3535010.72) (1.96875,5454096.867) (2,8421700.362) (2.03125,13013480.07) (2.0625,20122327.29) (2.09375,31133687.55) (2.125,48198005.97) (2.15625,74654208.15) (2.1875,115688009) (2.21875,179355569) (2.25,278175552.5) (2.28125,431605643.5) (2.3125,669895047.3) (2.34375,1040079409) (2.375,1615310095) (2.40625,2509374303) (2.4375,3899297195) (2.46875,6060529144) (2.5,9421730431) (2.53125,1.465008016e+10) (2.5625,2.278412111e+10) (2.59375,3.544066129e+10) (2.625,5.513701098e+10) (2.65625,8.579300113e+10) (2.6875,1.335128946e+11) (2.71875,2.078035834e+11) (2.75,3.234726981e+11) (2.78125,5.035857753e+11) (2.8125,7.84074466e+11) (2.84375,1.220916929e+12) (2.875,1.901328398e+12) (2.90625,2.961200399e+12) (2.9375,4.612280417e+12) (2.96875,7.184535425e+12) (3,1.119217866e+13)};
\addplot[color=PVorange,solid,line width=.75pt,mark=none] coordinates {(0.03125,0.04805417345) (0.0625,0.07979978851) (0.09375,0.116321825) (0.125,0.1549588734) (0.15625,0.1949921298) (0.1875,0.2355656269) (0.21875,0.277288019) (0.25,0.3240270401) (0.28125,0.3757172918) (0.3125,0.4322482244) (0.34375,0.4964092025) (0.375,0.5696611285) (0.40625,0.6534444167) (0.4375,0.7509481213) (0.46875,0.8576280507) (0.5,0.9611732266) (0.53125,1.063320954) (0.5625,1.169126232) (0.59375,1.266760572) (0.625,1.42470705) (0.65625,1.655313519) (0.6875,1.886709005) (0.71875,2.057728284) (0.75,2.208246278) (0.78125,2.35687342) (0.8125,2.474752077) (0.84375,2.594917599) (0.875,2.764614856) (0.90625,2.9906539) (0.9375,3.281279638) (0.96875,3.594444479) (1,3.891857251) (1.03125,4.138918025) (1.0625,4.473866714) (1.09375,5.047784032) (1.125,5.612641586) (1.15625,6.068877758) (1.1875,6.523558563) (1.21875,6.965216532) (1.25,7.378699099) (1.28125,7.802007232) (1.3125,8.272660838) (1.34375,8.882539895) (1.375,9.556595392) (1.40625,10.2204471) (1.4375,11.0544486) (1.46875,12.00947391) (1.5,12.70495887) (1.53125,14.40573963) (1.5625,17.32144344) (1.59375,19.45273063) (1.625,20.63341506) (1.65625,21.75492301) (1.6875,23.27351153) (1.71875,25.24936619) (1.75,27.09374276) (1.78125,29.06585262) (1.8125,31.75038939) (1.84375,34.84622443) (1.875,38.04639507) (1.90625,41.33888242) (1.9375,45.32631948) (1.96875,47.71286416) (2,53.29143227) (2.03125,65.16367814) (2.0625,75.97889659) (2.09375,80.84308027) (2.125,82.82559361) (2.15625,87.23619995) (2.1875,94.76982895) (2.21875,103.3550998) (2.25,112.4327187) (2.28125,123.3654574) (2.3125,136.720011) (2.34375,150.9080384) (2.375,163.4782634) (2.40625,179.543294) (2.4375,190.0524353) (2.46875,212.8424713) (2.5,264.4258236) (2.53125,321.6653061) (2.5625,351.7756201) (2.59375,367.2200526) (2.625,388.0326543) (2.65625,419.1150051) (2.6875,454.6241255) (2.71875,500.9217187) (2.75,557.5741507) (2.78125,632.677158) (2.8125,719.8884624) (2.84375,789.9063458) (2.875,860.6460374) (2.90625,924.4043958) (2.9375,1004.096212) (2.96875,1208.458768) (3,1519.031468)};
\addplot[color=PVred,solid,line width=.75pt,mark=none] coordinates {(0.03125,0.04558417209) (0.0625,0.09232375688) (0.09375,0.1399292917) (0.125,0.1874689473) (0.15625,0.2357381056) (0.1875,0.286305673) (0.21875,0.3409669304) (0.25,0.4012261312) (0.28125,0.4680220753) (0.3125,0.5417522632) (0.34375,0.6226260357) (0.375,0.7111139605) (0.40625,0.8081470835) (0.4375,0.915182069) (0.46875,1.034126973) (0.5,1.167264939) (0.53125,1.317418403) (0.5625,1.487686443) (0.59375,1.681155427) (0.625,1.901096684) (0.65625,2.150163112) (0.6875,2.430587219) (0.71875,2.745275728) (0.75,3.098198418) (0.78125,3.494430125) (0.8125,3.940384893) (0.84375,4.446662487) (0.875,5.031389038) (0.90625,5.718956933) (0.9375,6.542091244) (0.96875,7.551836303) (1,8.805608926) (1.03125,10.34288011) (1.0625,12.18996369) (1.09375,14.36780758) (1.125,16.91218264) (1.15625,19.88184479) (1.1875,23.37445625) (1.21875,27.54885184) (1.25,32.6263137) (1.28125,38.95209955) (1.3125,47.03930281) (1.34375,57.65999599) (1.375,71.94518122) (1.40625,91.58515075) (1.4375,119.0341883) (1.46875,157.8555439) (1.5,213.3404591) (1.53125,293.3218213) (1.5625,409.5596575) (1.59375,579.8259566) (1.625,830.8941268) (1.65625,1203.119412) (1.6875,1757.364075) (1.71875,2585.627141) (1.75,3827.106764) (1.78125,5692.609371) (1.8125,8501.501288) (1.84375,12737.78715) (1.875,19136.4672) (1.90625,28813.33346) (1.9375,43459.41001) (1.96875,65641.97426) (2,99257.57296) (2.03125,150212.6766) (2.0625,227471.9174) (2.09375,344643.3466) (2.125,522387.1972) (2.15625,792073.0295) (2.1875,1201327.758) (2.21875,1822471.053) (2.25,2765319.45) (2.28125,4196627.827) (2.3125,6369610.892) (2.34375,9668779.852) (2.375,14678013.82) (2.40625,22283922.58) (2.4375,33832840.88) (2.46875,51369220.93) (2.5,77997634.61) (2.53125,118432506.3) (2.5625,179832952.9) (2.59375,273070557.7) (2.625,414654511.5) (2.65625,629655113.1) (2.6875,956143398.5) (2.71875,1451933111) (2.75,2204818660) (2.78125,3348122787) (2.8125,5084306649) (2.84375,7720824256) (2.875,1.172456808e+10) (2.90625,1.780455531e+10) (2.9375,2.703748048e+10) (2.96875,4.105840729e+10) (3,6.235030287e+10)};
\addplot[color=PVteal,solid,line width=.75pt,mark=none] coordinates {(0.03125,0.01997570492) (0.0625,0.03732434541) (0.09375,0.05297698954) (0.125,0.08586145325) (0.15625,0.1272022176) (0.1875,0.2072550554) (0.21875,0.4084655114) (0.25,0.658781967) (0.28125,1.124413028) (0.3125,1.862149079) (0.34375,2.656273536) (0.375,3.799848333) (0.40625,6.005658967) (0.4375,10.09292089) (0.46875,18.18563471) (0.5,35.62329626) (0.53125,74.70467632) (0.5625,163.6840641) (0.59375,368.8461401) (0.625,846.9161499) (0.65625,1969.37785) (0.6875,4620.320756) (0.71875,10910.34593) (0.75,25891.19921) (0.78125,61680.68387) (0.8125,147399.7182) (0.84375,353141.4067) (0.875,847846.6242) (0.90625,2039188.593) (0.9375,4911952.455) (0.96875,11847205.56) (1,28606747.83) (1.03125,69143436.87) (1.0625,167267910.3) (1.09375,404959078.9) (1.125,981094086.9) (1.15625,2378375376) (1.1875,5768911071) (1.21875,1.399999839e+10) (1.25,3.399090713e+10) (1.28125,8.256205887e+10) (1.3125,2.006160493e+11) (1.34375,4.876456802e+11) (1.375,1.18572579e+12) (1.40625,2.883993047e+12) (1.4375,7.016559387e+12) (1.46875,1.707517628e+13) (1.5,4.156319782e+13) (1.53125,1.01192426e+14) (1.5625,2.46419766e+14) (1.59375,6.001852357e+14) (1.625,1.462081648e+15) (1.65625,3.562289938e+15) (1.6875,8.680673206e+15) (1.71875,2.11562971e+16) (1.75,5.156843302e+16) (1.78125,1.257136604e+17) (1.8125,3.065009407e+17) (1.84375,7.473580574e+17) (1.875,1.822511253e+18) (1.90625,4.444813535e+18) (1.9375,1.084116957e+19) (1.96875,2.644451518e+19) (2,6.451039838e+19) (2.03125,1.57382496e+20) (2.0625,3.839845816e+20) (2.09375,9.369146297e+20) (2.125,2.286195854e+21) (2.15625,5.578949965e+21) (2.1875,1.361493758e+22) (2.21875,3.32278038e+22) (2.25,8.10978088e+22) (2.28125,1.979415243e+23) (2.3125,4.831521104e+23) (2.34375,1.179367033e+24) (2.375,2.878930844e+24) (2.40625,7.027966866e+24) (2.4375,1.715708559e+25) (2.46875,4.188628593e+25) (2.5,1.022619215e+26) (2.53125,2.49671562e+26) (2.5625,6.095882094e+26) (2.59375,1.488386626e+27) (2.625,3.634176888e+27) (2.65625,8.873744309e+27) (2.6875,2.166794686e+28) (2.71875,5.291004257e+28) (2.75,1.292014752e+29) (2.78125,3.155043913e+29) (2.8125,7.70462483e+29) (2.84375,1.881504787e+30) (2.875,4.594799067e+30) (2.90625,1.122108193e+31) (2.9375,2.740372524e+31) (2.96875,6.692537514e+31) (3,1.6344742e+32)};
\addplot[color=PVgray,dashed,line width=.75pt,mark=none] coordinates {(0.03125,0.0003006845346) (0.0625,0.0006054788909) (0.09375,0.0009461866138) (0.125,0.001260548505) (0.15625,0.001471750508) (0.1875,0.001540301023) (0.21875,0.001454167788) (0.25,0.001239301315) (0.28125,0.001008629521) (0.3125,0.0009478825595) (0.34375,0.001021913845) (0.375,0.00111543859) (0.40625,0.001211443146) (0.4375,0.00122704329) (0.46875,0.001194963523) (0.5,0.001199564667) (0.53125,0.001269505586) (0.5625,0.001357206171) (0.59375,0.001399311578) (0.625,0.001391000646) (0.65625,0.001397644114) (0.6875,0.001374152405) (0.71875,0.001332618528) (0.75,0.001325229271) (0.78125,0.001403948771) (0.8125,0.001560551648) (0.84375,0.001691438851) (0.875,0.001740802762) (0.90625,0.001697452482) (0.9375,0.001555864798) (0.96875,0.001337136729) (1,0.001113658927) (1.03125,0.001083457479) (1.0625,0.00138085772) (1.09375,0.001780783953) (1.125,0.002047144687) (1.15625,0.002196137829) (1.1875,0.002188198356) (1.21875,0.002110135207) (1.25,0.002111720914) (1.28125,0.002281192082) (1.3125,0.002405532963) (1.34375,0.002481476727) (1.375,0.002494104562) (1.40625,0.002496093151) (1.4375,0.002442838999) (1.46875,0.002389444944) (1.5,0.002289751083) (1.53125,0.002135889688) (1.5625,0.001976208114) (1.59375,0.001931331702) (1.625,0.00188065913) (1.65625,0.001772472874) (1.6875,0.001643541992) (1.71875,0.001544517819) (1.75,0.001522460126) (1.78125,0.001572524325) (1.8125,0.001784952667) (1.84375,0.001939682146) (1.875,0.00210199345) (1.90625,0.002225376123) (1.9375,0.002180994501) (1.96875,0.002058822129) (2,0.001984175924) (2.03125,0.001975921795) (2.0625,0.00203678473) (2.09375,0.002154007159) (2.125,0.002229502113) (2.15625,0.002288953005) (2.1875,0.00224097699) (2.21875,0.002091994731) (2.25,0.001843319613) (2.28125,0.00158234628) (2.3125,0.001402914713) (2.34375,0.00130527115) (2.375,0.001247470152) (2.40625,0.001221481816) (2.4375,0.001190494337) (2.46875,0.001091546734) (2.5,0.0009744294076) (2.53125,0.0009086581679) (2.5625,0.0009788562692) (2.59375,0.001125692539) (2.625,0.001215688474) (2.65625,0.001242214357) (2.6875,0.001207244836) (2.71875,0.001204866925) (2.75,0.001308535808) (2.78125,0.001499279371) (2.8125,0.001678184776) (2.84375,0.001840089154) (2.875,0.001950408871) (2.90625,0.00196138317) (2.9375,0.00184770022) (2.96875,0.001679344079) (3,0.001488192301)};
\addplot[color=PVnavy,densely dotted,line width=.75pt,mark=none] coordinates {(0.03125,2.332020765e-05) (0.0625,4.432474379e-05) (0.09375,6.580851886e-05) (0.125,8.528695815e-05) (0.15625,9.756380663e-05) (0.1875,9.984309696e-05) (0.21875,9.270523133e-05) (0.25,7.757131098e-05) (0.28125,6.32261764e-05) (0.3125,6.038557969e-05) (0.34375,6.403241025e-05) (0.375,7.016715464e-05) (0.40625,7.531082572e-05) (0.4375,7.554159397e-05) (0.46875,7.449767696e-05) (0.5,7.460186757e-05) (0.53125,7.915979372e-05) (0.5625,8.439842794e-05) (0.59375,8.689282657e-05) (0.625,8.643705589e-05) (0.65625,8.627137898e-05) (0.6875,8.453367656e-05) (0.71875,8.228190467e-05) (0.75,8.353118875e-05) (0.78125,9.103167402e-05) (0.8125,0.0001022342423) (0.84375,0.0001101470754) (0.875,0.0001120375402) (0.90625,0.0001082661853) (0.9375,9.823477992e-05) (0.96875,8.327514628e-05) (1,6.82533006e-05) (1.03125,6.998504964e-05) (1.0625,9.267085983e-05) (1.09375,0.0001155796422) (1.125,0.000132207392) (1.15625,0.0001398910508) (1.1875,0.0001381125138) (1.21875,0.0001329283142) (1.25,0.0001349439665) (1.28125,0.0001423674337) (1.3125,0.0001502649308) (1.34375,0.0001538050144) (1.375,0.0001564353809) (1.40625,0.0001532038062) (1.4375,0.0001517749429) (1.46875,0.0001469079233) (1.5,0.0001411919359) (1.53125,0.0001308622566) (1.5625,0.0001235772233) (1.59375,0.0001200875797) (1.625,0.0001157234864) (1.65625,0.0001077682885) (1.6875,0.0001004820592) (1.71875,9.639668776e-05) (1.75,9.576436628e-05) (1.78125,0.0001040857992) (1.8125,0.0001176011936) (1.84375,0.0001251181875) (1.875,0.0001357142637) (1.90625,0.000141904826) (1.9375,0.0001374757125) (1.96875,0.0001296611554) (2,0.0001241216152) (2.03125,0.0001226401636) (2.0625,0.0001288060035) (2.09375,0.0001353278992) (2.125,0.0001427182823) (2.15625,0.0001437768548) (2.1875,0.0001417049752) (2.21875,0.0001299251314) (2.25,0.0001126688322) (2.28125,9.738967439e-05) (2.3125,8.734724345e-05) (2.34375,7.933433373e-05) (2.375,7.601010797e-05) (2.40625,7.561543361e-05) (2.4375,7.186087587e-05) (2.46875,6.735610039e-05) (2.5,5.936600931e-05) (2.53125,5.695266107e-05) (2.5625,6.382868546e-05) (2.59375,7.076631111e-05) (2.625,7.508532496e-05) (2.65625,7.59817635e-05) (2.6875,7.423407545e-05) (2.71875,7.591883285e-05) (2.75,8.428039092e-05) (2.78125,9.621869944e-05) (2.8125,0.0001084204191) (2.84375,0.0001191722424) (2.875,0.0001262706385) (2.90625,0.0001237304565) (2.9375,0.0001158218725) (2.96875,0.0001033263505) (3,9.066816309e-05)};
\addplot[color=PVorange,dashed,line width=.75pt,mark=none] coordinates {(0.03125,0.04983898322) (0.0625,0.03936769561) (0.09375,0.03676566452) (0.125,0.03459919808) (0.15625,0.03205252577) (0.1875,0.02911636175) (0.21875,0.02599425666) (0.25,0.02300275349) (0.28125,0.02078062262) (0.3125,0.02100270393) (0.34375,0.02303544785) (0.375,0.02519145218) (0.40625,0.02648913371) (0.4375,0.0264166995) (0.46875,0.02481857691) (0.5,0.02224151834) (0.53125,0.02032455227) (0.5625,0.01981768839) (0.59375,0.0208017132) (0.625,0.02265975226) (0.65625,0.02434001824) (0.6875,0.02519314083) (0.71875,0.02506980188) (0.75,0.02423986246) (0.78125,0.02332053994) (0.8125,0.02305987769) (0.84375,0.02395161158) (0.875,0.02612344623) (0.90625,0.02940346058) (0.9375,0.03307948824) (0.96875,0.03617987812) (1,0.0381584715) (1.03125,0.03863691398) (1.0625,0.03955286956) (1.09375,0.04189122595) (1.125,0.04439906549) (1.15625,0.04589165236) (1.1875,0.045785971) (1.21875,0.0437576596) (1.25,0.04024286537) (1.28125,0.03658197802) (1.3125,0.03202293205) (1.34375,0.02719922502) (1.375,0.02347364227) (1.40625,0.02107528605) (1.4375,0.01922125618) (1.46875,0.01723140338) (1.5,0.01526397803) (1.53125,0.01444686709) (1.5625,0.01558613481) (1.59375,0.018229519) (1.625,0.02124504234) (1.65625,0.02362352198) (1.6875,0.02489296907) (1.71875,0.02498656677) (1.75,0.02424531855) (1.78125,0.0237673406) (1.8125,0.02547621122) (1.84375,0.02910476126) (1.875,0.03325358498) (1.90625,0.03750208202) (1.9375,0.04142661515) (1.96875,0.0441234882) (2,0.0449386102) (2.03125,0.04420454099) (2.0625,0.04223306198) (2.09375,0.03963300495) (2.125,0.03683233474) (2.15625,0.03380022673) (2.1875,0.03042563418) (2.21875,0.02685481793) (2.25,0.02347222098) (2.28125,0.02071382386) (2.3125,0.01884625479) (2.34375,0.01781899145) (2.375,0.01723978395) (2.40625,0.01661597007) (2.4375,0.01556028021) (2.46875,0.01405314847) (2.5,0.01261997186) (2.53125,0.01281720926) (2.5625,0.01613460298) (2.59375,0.02087718506) (2.625,0.02504006728) (2.65625,0.02766738206) (2.6875,0.02861165152) (2.71875,0.02815832899) (2.75,0.02703889978) (2.78125,0.02597373634) (2.8125,0.02511773495) (2.84375,0.02530535726) (2.875,0.02715711854) (2.90625,0.03040679338) (2.9375,0.03401605643) (2.96875,0.03692103575) (3,0.03862611948)};
\addplot[color=PVteal,densely dotted,line width=.75pt,mark=none] coordinates {(0.03125,0.0002174363026) (0.0625,0.0003025822147) (0.09375,0.0003198824909) (0.125,0.000333001928) (0.15625,0.0003514199509) (0.1875,0.0003636533498) (0.21875,0.000363084084) (0.25,0.0003510161835) (0.28125,0.0003298850675) (0.3125,0.0003033001794) (0.34375,0.0002737404448) (0.375,0.0002493006757) (0.40625,0.0002339031351) (0.4375,0.0002240663639) (0.46875,0.0002198702067) (0.5,0.000221141526) (0.53125,0.0002286883406) (0.5625,0.0002450597928) (0.59375,0.0002705534083) (0.625,0.0003024946653) (0.65625,0.0003386762309) (0.6875,0.0003752626468) (0.71875,0.0004101717023) (0.75,0.0004431405228) (0.78125,0.0004754719161) (0.8125,0.0005090977417) (0.84375,0.0005450976499) (0.875,0.0005827209695) (0.90625,0.000620020723) (0.9375,0.0006531449197) (0.96875,0.0006780643297) (1,0.0006927306188) (1.03125,0.000697417128) (1.0625,0.0006937490549) (1.09375,0.0006842422602) (1.125,0.0006729564362) (1.15625,0.0006615916045) (1.1875,0.0006488383338) (1.21875,0.0006326571598) (1.25,0.0006142843179) (1.28125,0.000595855761) (1.3125,0.0005800874863) (1.34375,0.0005676587555) (1.375,0.0005565741437) (1.40625,0.0005457774341) (1.4375,0.0005381830559) (1.46875,0.0005366738171) (1.5,0.0005428823263) (1.53125,0.0005559843847) (1.5625,0.0005746987076) (1.59375,0.0005980800353) (1.625,0.0006280195989) (1.65625,0.0006631081488) (1.6875,0.0007003053321) (1.71875,0.000736363691) (1.75,0.0007703704724) (1.78125,0.0008032164011) (1.8125,0.0008336221222) (1.84375,0.0008615716175) (1.875,0.0008889213973) (1.90625,0.0009141195336) (1.9375,0.0009340243121) (1.96875,0.0009459587233) (2,0.0009490096273) (2.03125,0.0009433694841) (2.0625,0.0009309152001) (2.09375,0.0009125534671) (2.125,0.0008907592325) (2.15625,0.0008668354924) (2.1875,0.0008419131795) (2.21875,0.000815941124) (2.25,0.0007883614083) (2.28125,0.000759205339) (2.3125,0.0007308874252) (2.34375,0.0007053729688) (2.375,0.0006837785475) (2.40625,0.0006679893296) (2.4375,0.0006592358802) (2.46875,0.0006569276703) (2.5,0.0006594700732) (2.53125,0.0006667568235) (2.5625,0.0006796275305) (2.59375,0.0006984767979) (2.625,0.0007232218401) (2.65625,0.0007527848629) (2.6875,0.0007851522815) (2.71875,0.0008181337979) (2.75,0.0008505386897) (2.78125,0.0008821590047) (2.8125,0.000913475281) (2.84375,0.0009441549743) (2.875,0.0009734098792) (2.90625,0.0009992120525) (2.9375,0.00101946379) (2.96875,0.001032155288) (3,0.001035901927)};
\addplot[color=PVgray,dashdotted,line width=.75pt,mark=none] coordinates {(0.03125,0.05901585131) (0.0625,0.1176565207) (0.09375,0.1729490145) (0.125,0.2223565017) (0.15625,0.264160765) (0.1875,0.2976138965) (0.21875,0.322916372) (0.25,0.3410382954) (0.28125,0.3533815009) (0.3125,0.3614774824) (0.34375,0.36679459) (0.375,0.3704820861) (0.40625,0.3731659083) (0.4375,0.374919644) (0.46875,0.3753300017) (0.5,0.3735936431) (0.53125,0.3687830961) (0.5625,0.3604335911) (0.59375,0.3487348949) (0.625,0.3343650119) (0.65625,0.3185398503) (0.6875,0.3028790799) (0.71875,0.2888754566) (0.75,0.277157385) (0.78125,0.267025861) (0.8125,0.2566498719) (0.84375,0.2438083144) (0.875,0.2268016625) (0.90625,0.2053554498) (0.9375,0.1816290241) (0.96875,0.1616088297) (1,0.1560388271) (1.03125,0.172169248) (1.0625,0.2026712865) (1.09375,0.2377349495) (1.125,0.2721418948) (1.15625,0.3033160314) (1.1875,0.3297217305) (1.21875,0.3502331137) (1.25,0.3639047251) (1.28125,0.3702928302) (1.3125,0.3706567993) (1.34375,0.3674769972) (1.375,0.3626814064) (1.40625,0.3572495084) (1.4375,0.3514449773) (1.46875,0.3450738915) (1.5,0.3376749194) (1.53125,0.3286279006) (1.5625,0.3173145486) (1.59375,0.3034451383) (1.625,0.2874625996) (1.65625,0.2707601652) (1.6875,0.2554623673) (1.71875,0.2436235968) (1.75,0.2360818668) (1.78125,0.2315819787) (1.8125,0.2268562965) (1.84375,0.2184008256) (1.875,0.2035937713) (1.90625,0.1809174547) (1.9375,0.1509677798) (1.96875,0.1181109649) (2,0.09493304656) (2.03125,0.1017621886) (2.0625,0.1327139076) (2.09375,0.172600178) (2.125,0.2127848008) (2.15625,0.248627845) (2.1875,0.2779856958) (2.21875,0.3003961337) (2.25,0.3165121134) (2.28125,0.3275868496) (2.3125,0.3350168056) (2.34375,0.3399826964) (2.375,0.3432506519) (2.40625,0.3451713416) (2.4375,0.3457996762) (2.46875,0.3449928712) (2.5,0.3424283932) (2.53125,0.3375545961) (2.5625,0.3296199558) (2.59375,0.318167507) (2.625,0.303561278) (2.65625,0.2870370053) (2.6875,0.2703750333) (2.71875,0.2551961696) (2.75,0.2421109342) (2.78125,0.2302374897) (2.8125,0.2175707658) (2.84375,0.2015661129) (2.875,0.1797723515) (2.90625,0.1507449478) (2.9375,0.1147859875) (2.96875,0.07553346618) (3,0.05016556759)};
\end{axis}
\node[anchor=west,font=\fontsize{9}{11}\selectfont\bfseries] at (0.5,114) {C\quad Prediction diagnostics on unseen structures};
\begin{axis}[at={(22mm,59mm)},anchor=south west,width=128mm,height=46mm,scale only axis,font=\fontsize{8}{10}\selectfont,tick label style={font=\fontsize{8}{9.5}\selectfont},label style={font=\fontsize{8}{10}\selectfont},axis line style={PVgray!65,line width=.45pt},tick style={PVgray!65},grid=major,grid style={PVgray!15,line width=.3pt},enlarge x limits=false,enlarge y limits=false,clip=true,ymode=log,ymin=1e-08,ymax=1e+32,xmin=.5,xmax=4.5,xtick={1,2,3,4},xticklabels={Readout,{One step},Rollout,{Re-encode}},ylabel={Standardized RMSE},xlabel={Diagnostic}]
\addplot[only marks,mark=*,mark size=2.4pt,color=PVblue] coordinates {(0.85,0.01122492079) (1.85,0.04303858501) (2.85,1.48989214e+12) (3.85,9.143944182e+29)};
\addplot[only marks,mark=square*,mark size=2.4pt,color=PVorange] coordinates {(0.95,0.02674050527) (1.95,0.04911928844) (2.95,329.4632333)};
\node[text=PVorange,anchor=north] at (axis cs:3.95,1e+32) {$\times$};
\addplot[only marks,mark=triangle*,mark size=2.4pt,color=PVred] coordinates {(1.05,0.01199468704) (2.05,0.03951267565) (3.05,8455843888)};
\node[text=PVred,anchor=north] at (axis cs:4.05,1e+32) {$\times$};
\addplot[only marks,mark=diamond*,mark size=2.4pt,color=PVteal] coordinates {(1.15,2.526324245e-08) (2.15,0.01454091612) (3.15,1.828490347e+31) (4.15,1.828490347e+31)};
\end{axis}
\node[align=center,text width=150mm] at (76,36) {Readout uses true-state target encodings; one-step forecasts refresh the observed state.\\Rollout propagates latent predictions; re-encode is a secondary diagnostic.};
\draw[PVblue,solid,line width=.8pt] (1,17)--(8,17);
\node[anchor=west,font=\fontsize{8}{10}\selectfont] at (9,17) {Mono JEPA};
\draw[PVorange,solid,line width=.8pt] (53,17)--(60,17);
\node[anchor=west,font=\fontsize{8}{10}\selectfont] at (61,17) {OPF ($K=1$)};
\draw[PVred,solid,line width=.8pt] (105,17)--(112,17);
\node[anchor=west,font=\fontsize{8}{10}\selectfont] at (113,17) {OPF ($K=4$)};
\draw[PVteal,solid,line width=.8pt] (1,11)--(8,11);
\node[anchor=west,font=\fontsize{8}{10}\selectfont] at (9,11) {Supervised GNN};
\draw[PVgray,dashed,line width=.8pt] (53,11)--(60,11);
\node[anchor=west,font=\fontsize{8}{10}\selectfont] at (61,11) {RLC, 1 substep};
\draw[PVnavy,densely dotted,line width=.8pt] (105,11)--(112,11);
\node[anchor=west,font=\fontsize{8}{10}\selectfont] at (113,11) {RLC, 4 substeps};
\draw[PVorange,dashed,line width=.8pt] (1,5)--(8,5);
\node[anchor=west,font=\fontsize{8}{10}\selectfont] at (9,5) {Reduced RC};
\draw[PVteal,densely dotted,line width=.8pt] (53,5)--(60,5);
\node[anchor=west,font=\fontsize{8}{10}\selectfont] at (61,5) {Fitted RLC};
\draw[PVgray,dashdotted,line width=.8pt] (105,5)--(112,5);
\node[anchor=west,font=\fontsize{8}{10}\selectfont] at (113,5) {Persistence};
\end{tikzpicture}
\end{checkedfigure}
\caption{Forecast horizon and representation/rollout diagnostics. (A,B) Equal-structure mean errors at each of 96 forecast steps use identical logarithmic axes; every planned model is included. (C) Means across the five neural seeds separate readout reconstruction, observation-refreshed one-step prediction, primary free rollout, and secondary decoded-state re-encoding. A label of zero denotes an exact identity/zero error displayed at the log-axis floor; crosses denote nonfinite or unavailable diagnostics. Large rollout errors are retained, not replaced by a more favourable diagnostic.}
\label{fig:horizon-diagnostics}
\end{figure}

\begin{figure}[!tp]
\centering
\begin{checkedfigure}
\begin{tikzpicture}[x=1mm,y=1mm,font=\fontsize{8}{10}\selectfont,text=PVnavy,line cap=round,line join=round]
\path[use as bounding box] (0,0) rectangle (152,157);
\node[anchor=west,font=\fontsize{9}{11}\selectfont\bfseries] at (1,153) {A\quad Inlet pressure};
\begin{axis}[at={(17mm,106mm)},anchor=south west,width=55mm,height=38mm,scale only axis,font=\fontsize{8}{10}\selectfont,tick label style={font=\fontsize{8}{9.5}\selectfont},label style={font=\fontsize{8}{10}\selectfont},axis line style={PVgray!65,line width=.45pt},tick style={PVgray!65},grid=major,grid style={PVgray!15,line width=.3pt},enlarge x limits=false,enlarge y limits=false,clip=true,xmin=0,xmax=3,ymin=78.41906456,ymax=93.11305255,xtick={0,1,2,3},xlabel={Cycles},ylabel={mmHg},scaled y ticks=false]
\addplot[color=PVgray,dashed,line width=.75pt,mark=none] coordinates {(0.03125,85.05534521) (0.0625,85.9176164) (0.09375,86.84053037) (0.125,87.76209501) (0.15625,88.63321738) (0.1875,89.41496556) (0.21875,90.08166098) (0.25,90.62679722) (0.28125,91.06092685) (0.3125,91.40000629) (0.34375,91.65380271) (0.375,91.8231888) (0.40625,91.90417772) (0.4375,91.88908496) (0.46875,91.76264409) (0.5,91.5028301) (0.53125,91.09161878) (0.5625,90.52385895) (0.59375,89.80192571) (0.625,88.92662869) (0.65625,87.90389828) (0.6875,86.76122718) (0.71875,85.54897873) (0.75,84.3243912) (0.78125,83.14271921) (0.8125,82.06475507) (0.84375,81.15988146) (0.875,80.49030004) (0.90625,80.09274542) (0.9375,79.97727551) (0.96875,80.13604615) (1,80.54408105) (1.03125,81.15473276) (1.0625,81.90864944) (1.09375,82.75669192) (1.125,83.66953604) (1.15625,84.61985186) (1.1875,85.56600444) (1.21875,86.4665678) (1.25,87.30348226) (1.28125,88.07486542) (1.3125,88.76995176) (1.34375,89.37009051) (1.375,89.87326302) (1.40625,90.29659276) (1.4375,90.64882388) (1.46875,90.91554276) (1.5,91.07612955) (1.53125,91.11926889) (1.5625,91.03309278) (1.59375,90.79393438) (1.625,90.37986463) (1.65625,89.7930011) (1.6875,89.06194087) (1.71875,88.22876064) (1.75,87.34452991) (1.78125,86.47478072) (1.8125,85.69484761) (1.84375,85.07054019) (1.875,84.6442047) (1.90625,84.43804175) (1.9375,84.46046583) (1.96875,84.70203854) (2,85.1316392) (2.03125,85.70676244) (2.0625,86.38712858) (2.09375,87.13244403) (2.125,87.89238417) (2.15625,88.6133726) (2.1875,89.25899171) (2.21875,89.81531403) (2.25,90.27591024) (2.28125,90.63367996) (2.3125,90.8917251) (2.34375,91.07047596) (2.375,91.19360793) (2.40625,91.26985848) (2.4375,91.292901) (2.46875,91.25192057) (2.5,91.13269287) (2.53125,90.91105567) (2.5625,90.55767236) (2.59375,90.05600704) (2.625,89.41236339) (2.65625,88.64708073) (2.6875,87.78460932) (2.71875,86.85937595) (2.75,85.92489586) (2.78125,85.04602116) (2.8125,84.28177952) (2.84375,83.68109633) (2.875,83.29003409) (2.90625,83.14899585) (2.9375,83.27699138) (2.96875,83.66387081) (3,84.27932675)};
\addplot[color=PVnavy,densely dotted,line width=.75pt,mark=none] coordinates {(0.03125,85.04972762) (0.0625,85.90750715) (0.09375,86.83157777) (0.125,87.7534631) (0.15625,88.62229029) (0.1875,89.40326898) (0.21875,90.07283873) (0.25,90.62341222) (0.28125,91.06394844) (0.3125,91.40903561) (0.34375,91.66632519) (0.375,91.8359027) (0.40625,91.91634322) (0.4375,91.90004564) (0.46875,91.76708946) (0.5,91.49571049) (0.53125,91.0762682) (0.5625,90.50838597) (0.59375,89.78796145) (0.625,88.91004053) (0.65625,87.88863508) (0.6875,86.75885096) (0.71875,85.56017266) (0.75,84.33702894) (0.78125,83.15179468) (0.8125,82.0772529) (0.84375,81.17747416) (0.875,80.50380737) (0.90625,80.09662804) (0.9375,79.9761628) (0.96875,80.13424714) (1,80.53884512) (1.03125,81.14188651) (1.0625,81.88830978) (1.09375,82.73536177) (1.125,83.65531771) (1.15625,84.6098358) (1.1875,85.55167852) (1.21875,86.45572632) (1.25,87.31270287) (1.28125,88.09880441) (1.3125,88.78650032) (1.34375,89.37613055) (1.375,89.88434768) (1.40625,90.31169579) (1.4375,90.64976528) (1.46875,90.90104911) (1.5,91.06394672) (1.53125,91.11551122) (1.5625,91.02703059) (1.59375,90.78155135) (1.625,90.37028303) (1.65625,89.79187356) (1.6875,89.0645595) (1.71875,88.23055672) (1.75,87.34787148) (1.78125,86.48370409) (1.8125,85.70892527) (1.84375,85.08496187) (1.875,84.65503469) (1.90625,84.44653778) (1.9375,84.46814589) (1.96875,84.70375291) (2,85.1211546) (2.03125,85.68692049) (2.0625,86.36670895) (2.09375,87.11347983) (2.125,87.87002) (2.15625,88.59120902) (2.1875,89.24982667) (2.21875,89.82101302) (2.25,90.2839469) (2.28125,90.63846004) (2.3125,90.90090582) (2.34375,91.08650619) (2.375,91.20690973) (2.40625,91.27423206) (2.4375,91.29317782) (2.46875,91.25286905) (2.5,91.13135124) (2.53125,90.90314006) (2.5625,90.54432265) (2.59375,90.04287375) (2.625,89.40386711) (2.65625,88.6412726) (2.6875,87.77816929) (2.71875,86.85709133) (2.75,85.93435993) (2.78125,85.06412438) (2.8125,84.298038) (2.84375,83.69312866) (2.875,83.30317672) (2.90625,83.161404) (2.9375,83.27951742) (2.96875,83.6551727) (3,84.26746935)};
\addplot[color=PVorange,dashed,line width=.75pt,mark=none] coordinates {(0.03125,84.15460998) (0.0625,84.88384979) (0.09375,85.82967139) (0.125,86.87607284) (0.15625,87.93725217) (0.1875,88.94493369) (0.21875,89.84821783) (0.25,90.61493821) (0.28125,91.22721636) (0.3125,91.67652396) (0.34375,91.96521631) (0.375,92.10528966) (0.40625,92.11305255) (0.4375,92.00519517) (0.46875,91.79668348) (0.5,91.50045798) (0.53125,91.12060443) (0.5625,90.64436019) (0.59375,90.04978161) (0.625,89.31745805) (0.65625,88.43697423) (0.6875,87.41225999) (0.71875,86.26490002) (0.75,85.03488268) (0.78125,83.77856709) (0.8125,82.56403923) (0.84375,81.46441808) (0.875,80.54999107) (0.90625,79.88025029) (0.9375,79.49693596) (0.96875,79.41906456) (1,79.64064433) (1.03125,80.14214531) (1.0625,80.90014056) (1.09375,81.87373025) (1.125,82.99914454) (1.15625,84.20087579) (1.1875,85.40343156) (1.21875,86.54048272) (1.25,87.55991017) (1.28125,88.43131711) (1.3125,89.15125507) (1.34375,89.73421979) (1.375,90.20142499) (1.40625,90.5747335) (1.4375,90.8709499) (1.46875,91.09772216) (1.5,91.25165706) (1.53125,91.31893127) (1.5625,91.27828944) (1.59375,91.10593458) (1.625,90.78150181) (1.65625,90.29411593) (1.6875,89.64749748) (1.71875,88.86320832) (1.75,87.98139675) (1.78125,87.05518773) (1.8125,86.14282637) (1.84375,85.30535979) (1.875,84.60330031) (1.90625,84.08980963) (1.9375,83.80562213) (1.96875,83.77654293) (2,84.01348803) (2.03125,84.50624057) (2.0625,85.21418828) (2.09375,86.07411983) (2.125,87.01446137) (2.15625,87.96399216) (2.1875,88.85961484) (2.21875,89.65228108) (2.25,90.31047743) (2.28125,90.82095521) (2.3125,91.1867717) (2.34375,91.42310145) (2.375,91.55160104) (2.40625,91.59431339) (2.4375,91.5681471) (2.46875,91.48085411) (2.5,91.32917312) (2.53125,91.10405513) (2.5625,90.79664411) (2.59375,90.39539018) (2.625,89.88582573) (2.65625,89.25693964) (2.6875,88.50747855) (2.71875,87.65049029) (2.75,86.71495012) (2.78125,85.74723743) (2.8125,84.81094154) (2.84375,83.97866519) (2.875,83.3209859) (2.90625,82.89809725) (2.9375,82.75262804) (2.96875,82.90455591) (3,83.34885061)};
\addplot[color=PVteal,densely dotted,line width=.75pt,mark=none] coordinates {(0.03125,85.04449107) (0.0625,85.90034992) (0.09375,86.82633548) (0.125,87.74749974) (0.15625,88.61431962) (0.1875,89.39646863) (0.21875,90.06893869) (0.25,90.62138765) (0.28125,91.06357321) (0.3125,91.41085897) (0.34375,91.66961587) (0.375,91.83924983) (0.40625,91.91892056) (0.4375,91.90122995) (0.46875,91.76617106) (0.5,91.49246807) (0.53125,91.07104381) (0.5625,90.50164248) (0.59375,89.78019673) (0.625,88.90187329) (0.65625,87.88060682) (0.6875,86.75107254) (0.71875,85.55225734) (0.75,84.3284325) (0.78125,83.14203567) (0.8125,82.06574831) (0.84375,81.16355833) (0.875,80.4871146) (0.90625,80.07718916) (0.9375,79.95412484) (0.96875,80.10984118) (1,80.51259624) (1.03125,81.11455478) (1.0625,81.86071681) (1.09375,82.70847987) (1.125,83.63015857) (1.15625,84.58706971) (1.1875,85.53181005) (1.21875,86.43930478) (1.25,87.29965982) (1.28125,88.08816165) (1.3125,88.77726917) (1.34375,89.36775447) (1.375,89.87605512) (1.40625,90.30250439) (1.4375,90.63941887) (1.46875,90.89000443) (1.5,91.05242082) (1.53125,91.10344544) (1.5625,91.01473606) (1.59375,90.76948305) (1.625,90.35843323) (1.65625,89.77999309) (1.6875,89.052587) (1.71875,88.21840066) (1.75,87.33515528) (1.78125,86.46999445) (1.8125,85.69389347) (1.84375,85.06821859) (1.875,84.63614215) (1.90625,84.42522143) (1.9375,84.44427224) (1.96875,84.67730196) (2,85.09247371) (2.03125,85.65680836) (2.0625,86.33612978) (2.09375,87.08336678) (2.125,87.84136529) (2.15625,88.56501545) (2.1875,89.22664976) (2.21875,89.80072728) (2.25,90.26617724) (2.28125,90.62287368) (2.3125,90.88691044) (2.34375,91.07321977) (2.375,91.1936657) (2.40625,91.26070833) (2.4375,91.27904928) (2.46875,91.2378034) (2.5,91.1153013) (2.53125,90.88628622) (2.5625,90.52684647) (2.59375,90.02502277) (2.625,89.38597869) (2.65625,88.62355436) (2.6875,87.76070195) (2.71875,86.83992438) (2.75,85.91725789) (2.78125,85.04644671) (2.8125,84.27911559) (2.84375,83.67248403) (2.875,83.28032946) (2.90625,83.13592824) (2.9375,83.25145157) (2.96875,83.62504168) (3,84.23589061)};
\addplot[color=PVgray,dashdotted,line width=.75pt,mark=none] coordinates {(0.03125,84.32542123) (0.0625,84.32542123) (0.09375,84.32542123) (0.125,84.32542123) (0.15625,84.32542123) (0.1875,84.32542123) (0.21875,84.32542123) (0.25,84.32542123) (0.28125,84.32542123) (0.3125,84.32542123) (0.34375,84.32542123) (0.375,84.32542123) (0.40625,84.32542123) (0.4375,84.32542123) (0.46875,84.32542123) (0.5,84.32542123) (0.53125,84.32542123) (0.5625,84.32542123) (0.59375,84.32542123) (0.625,84.32542123) (0.65625,84.32542123) (0.6875,84.32542123) (0.71875,84.32542123) (0.75,84.32542123) (0.78125,84.32542123) (0.8125,84.32542123) (0.84375,84.32542123) (0.875,84.32542123) (0.90625,84.32542123) (0.9375,84.32542123) (0.96875,84.32542123) (1,84.32542123) (1.03125,84.32542123) (1.0625,84.32542123) (1.09375,84.32542123) (1.125,84.32542123) (1.15625,84.32542123) (1.1875,84.32542123) (1.21875,84.32542123) (1.25,84.32542123) (1.28125,84.32542123) (1.3125,84.32542123) (1.34375,84.32542123) (1.375,84.32542123) (1.40625,84.32542123) (1.4375,84.32542123) (1.46875,84.32542123) (1.5,84.32542123) (1.53125,84.32542123) (1.5625,84.32542123) (1.59375,84.32542123) (1.625,84.32542123) (1.65625,84.32542123) (1.6875,84.32542123) (1.71875,84.32542123) (1.75,84.32542123) (1.78125,84.32542123) (1.8125,84.32542123) (1.84375,84.32542123) (1.875,84.32542123) (1.90625,84.32542123) (1.9375,84.32542123) (1.96875,84.32542123) (2,84.32542123) (2.03125,84.32542123) (2.0625,84.32542123) (2.09375,84.32542123) (2.125,84.32542123) (2.15625,84.32542123) (2.1875,84.32542123) (2.21875,84.32542123) (2.25,84.32542123) (2.28125,84.32542123) (2.3125,84.32542123) (2.34375,84.32542123) (2.375,84.32542123) (2.40625,84.32542123) (2.4375,84.32542123) (2.46875,84.32542123) (2.5,84.32542123) (2.53125,84.32542123) (2.5625,84.32542123) (2.59375,84.32542123) (2.625,84.32542123) (2.65625,84.32542123) (2.6875,84.32542123) (2.71875,84.32542123) (2.75,84.32542123) (2.78125,84.32542123) (2.8125,84.32542123) (2.84375,84.32542123) (2.875,84.32542123) (2.90625,84.32542123) (2.9375,84.32542123) (2.96875,84.32542123) (3,84.32542123)};
\addplot[color=black,line width=.95pt,mark=none] coordinates {(0.03125,85.04923371) (0.0625,85.90691902) (0.09375,86.83114043) (0.125,87.75274862) (0.15625,88.62140617) (0.1875,89.40260455) (0.21875,90.07237639) (0.25,90.62316412) (0.28125,91.06414939) (0.3125,91.4097116) (0.34375,91.66716555) (0.375,91.83669631) (0.40625,91.91720264) (0.4375,91.90080357) (0.46875,91.76718169) (0.5,91.49507681) (0.53125,91.07541587) (0.5625,90.50751749) (0.59375,89.78682505) (0.625,88.90872414) (0.65625,87.88794516) (0.6875,86.7591507) (0.71875,85.56070432) (0.75,84.33740044) (0.78125,83.15262641) (0.8125,82.07856061) (0.84375,81.17837286) (0.875,80.50415699) (0.90625,80.09701899) (0.9375,79.97649087) (0.96875,80.13398911) (1,80.53814841) (1.03125,81.1410874) (1.0625,81.88715366) (1.09375,82.73399602) (1.125,83.65442717) (1.15625,84.60899506) (1.1875,85.55046165) (1.21875,86.45544266) (1.25,87.31389984) (1.28125,88.09987972) (1.3125,88.78687766) (1.34375,89.37711966) (1.375,89.88584856) (1.40625,90.31197842) (1.4375,90.64892957) (1.46875,90.90071512) (1.5,91.06396722) (1.53125,91.11472506) (1.5625,91.02595391) (1.59375,90.78121866) (1.625,90.37023964) (1.65625,89.79148221) (1.6875,89.0643233) (1.71875,88.23089551) (1.75,87.34838333) (1.78125,86.48418413) (1.8125,85.70968204) (1.84375,85.0859165) (1.875,84.65579285) (1.90625,84.44716628) (1.9375,84.46866955) (1.96875,84.70366814) (2,85.12030971) (2.03125,85.68577963) (2.0625,86.36550806) (2.09375,87.11198792) (2.125,87.86828973) (2.15625,88.59005308) (2.1875,89.249696) (2.21875,89.82118517) (2.25,90.28399019) (2.28125,90.63901203) (2.3125,90.90203981) (2.34375,91.08732304) (2.375,91.2072288) (2.40625,91.27466093) (2.4375,91.29363422) (2.46875,91.25280613) (2.5,91.13089718) (2.53125,90.90267352) (2.5625,90.54368251) (2.59375,90.04203431) (2.625,89.40324402) (2.65625,88.64081348) (2.6875,87.77770103) (2.71875,86.85715303) (2.75,85.93520035) (2.78125,85.06509153) (2.8125,84.2988405) (2.84375,83.69417801) (2.875,83.30433105) (2.90625,83.16189186) (2.9375,83.27927911) (2.96875,83.65482248) (3,84.26701905)};
\end{axis}
\node[anchor=west,font=\fontsize{9}{11}\selectfont\bfseries] at (1,93) {C\quad Pressure absolute error};
\begin{axis}[at={(17mm,46mm)},anchor=south west,width=55mm,height=38mm,scale only axis,font=\fontsize{8}{10}\selectfont,tick label style={font=\fontsize{8}{9.5}\selectfont},label style={font=\fontsize{8}{10}\selectfont},axis line style={PVgray!65,line width=.45pt},tick style={PVgray!65},grid=major,grid style={PVgray!15,line width=.3pt},enlarge x limits=false,enlarge y limits=false,clip=true,ymode=log,ymin=1e-05,ymax=1e+23,xmin=0,xmax=3,xtick={0,1,2,3},xlabel={Cycles},ylabel={mmHg}]
\addplot[color=PVblue,solid,line width=.75pt,mark=none] coordinates {(0.03125,0.4367018978) (0.0625,0.9551565779) (0.09375,2.031530091) (0.125,3.634310729) (0.15625,5.706949011) (0.1875,8.176237807) (0.21875,10.96019753) (0.25,13.96654542) (0.28125,17.1015472) (0.3125,20.29274503) (0.34375,23.50974792) (0.375,26.76478011) (0.40625,30.09918865) (0.4375,33.574302) (0.46875,37.2613927) (0.5,41.21654538) (0.53125,45.45850343) (0.5625,49.96964253) (0.59375,54.71046116) (0.625,59.62451978) (0.65625,64.63246017) (0.6875,69.65029794) (0.71875,74.62577852) (0.75,79.55454334) (0.78125,84.4799747) (0.8125,89.50543145) (0.84375,94.80223276) (0.875,100.5891264) (0.90625,107.0984595) (0.9375,114.5445746) (0.96875,123.0834629) (1,132.7615449) (1.03125,143.4707964) (1.0625,154.9184152) (1.09375,166.6159118) (1.125,177.9382288) (1.15625,188.2771011) (1.1875,197.2181304) (1.21875,204.6977006) (1.25,211.1388919) (1.28125,217.4888606) (1.3125,225.0841802) (1.34375,235.4200034) (1.375,249.948582) (1.40625,269.9230165) (1.4375,296.2469296) (1.46875,329.3909461) (1.5,369.4488626) (1.53125,416.2966744) (1.5625,469.7323364) (1.59375,529.5647116) (1.625,595.7709612) (1.65625,668.731541) (1.6875,749.1514098) (1.71875,837.3605642) (1.75,932.9366764) (1.78125,1035.666162) (1.8125,1146.761558) (1.84375,1268.818994) (1.875,1404.514508) (1.90625,1555.129068) (1.9375,1719.825953) (1.96875,1896.835862) (2,2085.669936) (2.03125,2288.649913) (2.0625,2505.951374) (2.09375,2734.199763) (2.125,2968.663297) (2.15625,3204.583909) (2.1875,3440.635982) (2.21875,3680.891171) (2.25,3928.934114) (2.28125,4181.383507) (2.3125,4434.49722) (2.34375,4659.380982) (2.375,4842.751001) (2.40625,4961.59623) (2.4375,4982.005102) (2.46875,4876.673521) (2.5,4626.035301) (2.53125,4200.931119) (2.5625,3564.84289) (2.59375,2731.138537) (2.625,1674.37774) (2.65625,358.6046677) (2.6875,1283.842478) (2.71875,3417.482572) (2.75,6195.522718) (2.78125,9678.036933) (2.8125,13880.8478) (2.84375,18768.96239) (2.875,24092.04122) (2.90625,29674.85221) (2.9375,35634.36129) (2.96875,41731.95904) (3,48572.53158)};
\addplot[color=PVorange,solid,line width=.75pt,mark=none] coordinates {(0.03125,1.357764116) (0.0625,2.717804503) (0.09375,4.034935013) (0.125,5.261133867) (0.15625,6.379473128) (0.1875,7.397804448) (0.21875,8.339202176) (0.25,9.239536211) (0.28125,10.14143004) (0.3125,11.07750405) (0.34375,12.05646376) (0.375,13.06592121) (0.40625,14.08262076) (0.4375,15.07243575) (0.46875,15.98919927) (0.5,16.79034573) (0.53125,17.45310513) (0.5625,17.97294739) (0.59375,18.35212392) (0.625,18.60304036) (0.65625,18.76569448) (0.6875,18.90126135) (0.71875,19.06134869) (0.75,19.2725099) (0.78125,19.54392683) (0.8125,19.87263077) (0.84375,20.24014369) (0.875,20.6108474) (0.90625,20.92986839) (0.9375,21.13677665) (0.96875,21.212247) (1,21.19299043) (1.03125,21.1174307) (1.0625,20.99302307) (1.09375,20.83243853) (1.125,20.67621433) (1.15625,20.5679855) (1.1875,20.55281053) (1.21875,20.69795219) (1.25,21.06758951) (1.28125,21.675655) (1.3125,22.48570689) (1.34375,23.43439738) (1.375,24.43140442) (1.40625,25.35906463) (1.4375,26.11305215) (1.46875,26.63814538) (1.5,26.91977798) (1.53125,26.96764929) (1.5625,26.78050127) (1.59375,26.20012686) (1.625,24.76993222) (1.65625,21.99787297) (1.6875,17.77795962) (1.71875,12.38150962) (1.75,6.273849736) (1.78125,0.01855505789) (1.8125,5.937554063) (1.84375,11.49284064) (1.875,16.77571592) (1.90625,21.8302861) (1.9375,26.47057808) (1.96875,30.36270174) (2,33.15822186) (2.03125,34.59166911) (2.0625,34.56997488) (2.09375,33.1218878) (2.125,30.19269209) (2.15625,25.83099308) (2.1875,20.44084392) (2.21875,14.92906897) (2.25,10.11137991) (2.28125,5.960573484) (2.3125,2.113812678) (2.34375,1.531923226) (2.375,4.826829483) (2.40625,7.782467742) (2.4375,10.88256833) (2.46875,13.88674939) (2.5,16.22612156) (2.53125,18.24841307) (2.5625,20.56173597) (2.59375,22.50411434) (2.625,18.19915888) (2.65625,6.529874391) (2.6875,20.08085593) (2.71875,51.84045881) (2.75,81.60930063) (2.78125,116.1324936) (2.8125,153.969323) (2.84375,190.6681936) (2.875,225.4419289) (2.90625,259.4513958) (2.9375,291.3822086) (2.96875,321.1134282) (3,348.0788354)};
\addplot[color=PVred,solid,line width=.75pt,mark=none] coordinates {(0.03125,0.4699881676) (0.0625,0.6819942416) (0.09375,0.5142210041) (0.125,0.08641875675) (0.15625,0.4937992506) (0.1875,1.165054285) (0.21875,1.914054497) (0.25,2.749451037) (0.28125,3.68016982) (0.3125,4.714646863) (0.34375,5.871173265) (0.375,7.179922169) (0.40625,8.672911917) (0.4375,10.37404804) (0.46875,12.28373727) (0.5,14.35075893) (0.53125,16.45722884) (0.5625,18.43953005) (0.59375,20.13679695) (0.625,21.43567178) (0.65625,22.29852384) (0.6875,22.7918165) (0.71875,23.09572033) (0.75,23.45329856) (0.78125,24.07816257) (0.8125,25.07793634) (0.84375,26.40799829) (0.875,27.84618319) (0.90625,28.99727445) (0.9375,29.33746669) (0.96875,28.28250905) (1,25.23744414) (1.03125,19.59065765) (1.0625,10.68862572) (1.09375,2.162118942) (1.125,19.65720893) (1.15625,42.45345666) (1.1875,71.19059013) (1.21875,106.5208656) (1.25,149.0366948) (1.28125,199.2478) (1.3125,257.7487275) (1.34375,325.3221202) (1.375,402.7214316) (1.40625,490.5051765) (1.4375,589.4361958) (1.46875,700.9930479) (1.5,827.4120616) (1.53125,971.3494087) (1.5625,1135.507153) (1.59375,1322.319557) (1.625,1533.470102) (1.65625,1770.169437) (1.6875,2034.477215) (1.71875,2331.084434) (1.75,2669.828041) (1.78125,3061.369311) (1.8125,3507.926114) (1.84375,4016.955503) (1.875,4596.65967) (1.90625,5257.105746) (1.9375,6008.103081) (1.96875,6845.457983) (2,7779.113448) (2.03125,8813.408291) (2.0625,9953.908575) (2.09375,11198.0648) (2.125,12568.93151) (2.15625,14090.54452) (2.1875,15807.98353) (2.21875,17744.31429) (2.25,19941.08196) (2.28125,22439.6699) (2.3125,25269.78249) (2.34375,28475.44005) (2.375,32115.81477) (2.40625,36255.41143) (2.4375,41142.46476) (2.46875,46931.49182) (2.5,53768.97057) (2.53125,61703.00097) (2.5625,71134.61001) (2.59375,81962.00467) (2.625,93896.72502) (2.65625,107209.2377) (2.6875,122721.283) (2.71875,141537.0882) (2.75,164413.1834) (2.78125,192142.7331) (2.8125,225157.2227) (2.84375,264382.3031) (2.875,310236.6109) (2.90625,364030.3263) (2.9375,426298.1749) (2.96875,493018.5541) (3,561101.628)};
\addplot[color=PVteal,solid,line width=.75pt,mark=none] coordinates {(0.03125,0.077543985) (0.0625,0.1061822197) (0.09375,0.3040763578) (0.125,0.7391377283) (0.15625,0.9138703354) (0.1875,0.04834384834) (0.21875,1.396282305) (0.25,0.3014039301) (0.28125,5.613948873) (0.3125,9.44295046) (0.34375,2.053475828) (0.375,32.24805084) (0.40625,50.78079857) (0.4375,37.36684025) (0.46875,57.87749352) (0.5,102.1001126) (0.53125,184.0985183) (0.5625,341.0507076) (0.59375,620.2929656) (0.625,1120.380022) (0.65625,2035.352295) (0.6875,3699.076672) (0.71875,6723.662417) (0.75,12220.84901) (0.78125,22200.97087) (0.8125,40308.11602) (0.84375,73154.71409) (0.875,132729.3348) (0.90625,240767.5036) (0.9375,436679.2727) (0.96875,791922.7388) (1,1436064.942) (1.03125,2604034.237) (1.0625,4721796.401) (1.09375,8561684.161) (1.125,15524051.2) (1.15625,28147927.07) (1.1875,51036937.32) (1.21875,92538083.18) (1.25,167785569.8) (1.28125,304219807.6) (1.3125,551594002.1) (1.34375,1000117181) (1.375,1813350686) (1.40625,3287853260) (1.4375,5961324419) (1.46875,1.080868829e+10) (1.5,1.95976079e+10) (1.53125,3.553309761e+10) (1.5625,6.442627491e+10) (1.59375,1.168134586e+11) (1.625,2.117984299e+11) (1.65625,3.840188682e+11) (1.6875,6.962775275e+11) (1.71875,1.262444045e+12) (1.75,2.288979806e+12) (1.78125,4.150225983e+12) (1.8125,7.524914413e+12) (1.84375,1.364367434e+13) (1.875,2.473780374e+13) (1.90625,4.485294096e+13) (1.9375,8.132437842e+13) (1.96875,1.474519638e+14) (2,2.673500698e+14) (2.03125,4.847413879e+14) (2.0625,8.789009293e+14) (2.09375,1.593564449e+15) (2.125,2.889345165e+15) (2.15625,5.238769253e+15) (2.1875,9.498587218e+15) (2.21875,1.722220704e+16) (2.25,3.12261614e+16) (2.28125,5.661719578e+16) (2.3125,1.026545386e+17) (2.34375,1.861264004e+17) (2.375,3.374720118e+17) (2.40625,6.1188192e+17) (2.4375,1.109423589e+18) (2.46875,2.011532997e+18) (2.5,3.647177832e+18) (2.53125,6.61282062e+18) (2.5625,1.198992601e+19) (2.59375,2.173933454e+19) (2.625,3.941631318e+19) (2.65625,7.146703779e+19) (2.6875,1.29579292e+20) (2.71875,2.349445353e+20) (2.75,4.259857983e+20) (2.78125,7.723691593e+20) (2.8125,1.400408376e+21) (2.84375,2.539127729e+21) (2.875,4.6037778e+21) (2.90625,8.347263408e+21) (2.9375,1.513470326e+22) (2.96875,2.74412335e+22) (3,4.975462028e+22)};
\addplot[color=PVgray,dashed,line width=.75pt,mark=none] coordinates {(0.03125,0.006111499843) (0.0625,0.0106973733) (0.09375,0.009389934731) (0.125,0.009346392343) (0.15625,0.01181121453) (0.1875,0.01236101247) (0.21875,0.00928458578) (0.25,0.003633103923) (0.28125,0.00322253844) (0.3125,0.009705303206) (0.34375,0.01336283923) (0.375,0.01350751342) (0.40625,0.0130249235) (0.4375,0.01171861239) (0.46875,0.004537598335) (0.5,0.007753289791) (0.53125,0.01620291375) (0.5625,0.01634146547) (0.59375,0.01510065428) (0.625,0.01790455624) (0.65625,0.01595311519) (0.6875,0.002076486865) (0.71875,0.01172558745) (0.75,0.01300923956) (0.78125,0.009907197009) (0.8125,0.01380553935) (0.84375,0.01849140142) (0.875,0.01385695429) (0.90625,0.004273567162) (0.9375,0.0007846458087) (0.96875,0.002057044466) (1,0.005932640288) (1.03125,0.01364535752) (1.0625,0.02149577324) (1.09375,0.02269590548) (1.125,0.01510887649) (1.15625,0.01085680247) (1.1875,0.01554278143) (1.21875,0.01112514004) (1.25,0.01041757504) (1.28125,0.02501430284) (1.3125,0.01692589987) (1.34375,0.007029144312) (1.375,0.01258554028) (1.40625,0.01538566596) (1.4375,0.0001056833234) (1.46875,0.01482764474) (1.5,0.01216233062) (1.53125,0.004543831352) (1.5625,0.007138871669) (1.59375,0.01271572007) (1.625,0.00962498587) (1.65625,0.00151888708) (1.6875,0.002382431568) (1.71875,0.002134868784) (1.75,0.003853413774) (1.78125,0.009403412356) (1.8125,0.01483443064) (1.84375,0.01537631072) (1.875,0.01158814818) (1.90625,0.009124532529) (1.9375,0.008203727318) (1.96875,0.00162959892) (2,0.01132948991) (2.03125,0.02098281198) (2.0625,0.02162051611) (2.09375,0.02045610886) (2.125,0.02409443894) (2.15625,0.02331952147) (2.1875,0.009295706499) (2.21875,0.005871143379) (2.25,0.008079955201) (2.28125,0.00533206436) (2.3125,0.01031471105) (2.34375,0.01684707957) (2.375,0.01362086514) (2.40625,0.004802451278) (2.4375,0.0007332178222) (2.46875,0.0008855542312) (2.5,0.001795696803) (2.53125,0.00838214863) (2.5625,0.01398985194) (2.59375,0.01397273486) (2.625,0.009119370619) (2.65625,0.00626725769) (2.6875,0.006908292382) (2.71875,0.002222924361) (2.75,0.01030449229) (2.78125,0.01907036813) (2.8125,0.01706098956) (2.84375,0.01308168283) (2.875,0.01429696445) (2.90625,0.01289600509) (2.9375,0.002287727919) (2.96875,0.009048326213) (3,0.01230769951)};
\addplot[color=PVnavy,densely dotted,line width=.75pt,mark=none] coordinates {(0.03125,0.0004939088407) (0.0625,0.0005881242807) (0.09375,0.0004373428495) (0.125,0.0007144755522) (0.15625,0.0008841197734) (0.1875,0.000664428429) (0.21875,0.000462339383) (0.25,0.000248105662) (0.28125,0.0002009514227) (0.3125,0.0006759855602) (0.34375,0.0008403626749) (0.375,0.0007936138297) (0.40625,0.0008594261983) (0.4375,0.0007579339119) (0.46875,9.223074406e-05) (0.5,0.0006336736293) (0.53125,0.0008523300933) (0.5625,0.0008684798796) (0.59375,0.001136400528) (0.625,0.001316393746) (0.65625,0.0006899142988) (0.6875,0.0002997380475) (0.71875,0.0005316577857) (0.75,0.0003714990402) (0.78125,0.000831729259) (0.8125,0.001307712265) (0.84375,0.0008986949865) (0.875,0.0003496154763) (0.90625,0.0003909523601) (0.9375,0.0003280719318) (0.96875,0.000258030414) (1,0.0006967116596) (1.03125,0.0007991024495) (1.0625,0.001156112799) (1.09375,0.001365755659) (1.125,0.0008905447647) (1.15625,0.0008407394816) (1.1875,0.001216867465) (1.21875,0.0002836568008) (1.25,0.00119697393) (1.28125,0.001075309518) (1.3125,0.0003773442308) (1.34375,0.0009891118757) (1.375,0.001500879394) (1.40625,0.0002826383513) (1.4375,0.0008357193598) (1.46875,0.0003339925983) (1.5,2.050059823e-05) (1.53125,0.0007861552215) (1.5625,0.001076679876) (1.59375,0.0003326881579) (1.625,4.33866111e-05) (1.65625,0.0003913489876) (1.6875,0.0002362020381) (1.71875,0.000338788646) (1.75,0.0005118485464) (1.78125,0.0004800356597) (1.8125,0.0007567665647) (1.84375,0.0009546278835) (1.875,0.0007581600726) (1.90625,0.0006285046271) (1.9375,0.0005236652769) (1.96875,8.476190294e-05) (2,0.0008448983049) (2.03125,0.001140857737) (2.0625,0.001200885162) (2.09375,0.001491916307) (2.125,0.001730271186) (2.15625,0.001155942636) (2.1875,0.0001306657526) (2.21875,0.0001721570672) (2.25,4.329509379e-05) (2.28125,0.0005519846929) (2.3125,0.001133988326) (2.34375,0.0008168473414) (2.375,0.0003190701918) (2.40625,0.0004288672667) (2.4375,0.0004564065685) (2.46875,6.292286929e-05) (2.5,0.0004540641131) (2.53125,0.0004665328326) (2.5625,0.0006401378649) (2.59375,0.0008394364989) (2.625,0.0006230877641) (2.65625,0.0004591278777) (2.6875,0.0004682649594) (2.71875,6.170038328e-05) (2.75,0.0008404139799) (2.78125,0.000967146498) (2.8125,0.0008025003441) (2.84375,0.001049355314) (2.875,0.001154335981) (2.90625,0.0004878504016) (2.9375,0.0002383101988) (2.96875,0.0003502214218) (3,0.0004502995774)};
\addplot[color=PVorange,dashed,line width=.75pt,mark=none] coordinates {(0.03125,0.8946237318) (0.0625,1.023069233) (0.09375,1.001469046) (0.125,0.8766757754) (0.15625,0.6841540025) (0.1875,0.4576708594) (0.21875,0.2241585592) (0.25,0.008225909491) (0.28125,0.1630669714) (0.3125,0.2668123602) (0.34375,0.298050759) (0.375,0.26859335) (0.40625,0.1958499074) (0.4375,0.1043915961) (0.46875,0.02950179063) (0.5,0.005381164618) (0.53125,0.04518856077) (0.5625,0.136842708) (0.59375,0.262956561) (0.625,0.4087339177) (0.65625,0.5490290707) (0.6875,0.653109292) (0.71875,0.7041957029) (0.75,0.6974822389) (0.78125,0.6259406821) (0.8125,0.4854786213) (0.84375,0.2860452255) (0.875,0.04583408026) (0.90625,0.2167687037) (0.9375,0.4795549048) (0.96875,0.7149245414) (1,0.8975040711) (1.03125,0.998942094) (1.0625,0.9870131069) (1.09375,0.860265764) (1.125,0.6552826258) (1.15625,0.408119267) (1.1875,0.1470300899) (1.21875,0.08504005991) (1.25,0.2460103301) (1.28125,0.3314373865) (1.3125,0.3643774043) (1.34375,0.3571001363) (1.375,0.3155764354) (1.40625,0.2627550736) (1.4375,0.2220203345) (1.46875,0.1970070422) (1.5,0.1876898414) (1.53125,0.2042062064) (1.5625,0.2523355326) (1.59375,0.324715916) (1.625,0.4112621737) (1.65625,0.5026337197) (1.6875,0.5831741796) (1.71875,0.6323128097) (1.75,0.63301342) (1.78125,0.5710036025) (1.8125,0.4331443325) (1.84375,0.2194432941) (1.875,0.0524925432) (1.90625,0.357356647) (1.9375,0.6630474201) (1.96875,0.9271252152) (2,1.10682168) (2.03125,1.179539057) (2.0625,1.151319783) (2.09375,1.037868087) (2.125,0.8538283565) (2.15625,0.6260609192) (2.1875,0.3900811596) (2.21875,0.1689040965) (2.25,0.02648723452) (2.28125,0.1819431822) (2.3125,0.2847318841) (2.34375,0.3357784096) (2.375,0.3443722464) (2.40625,0.3196524611) (2.4375,0.2745128783) (2.46875,0.2280479836) (2.5,0.1982759412) (2.53125,0.2013816081) (2.5625,0.2529615946) (2.59375,0.3533558685) (2.625,0.4825817064) (2.65625,0.6161261593) (2.6875,0.7297775189) (2.71875,0.7933372619) (2.75,0.7797497687) (2.78125,0.6821459017) (2.8125,0.5121010351) (2.84375,0.2844871822) (2.875,0.01665484351) (2.90625,0.263794604) (2.9375,0.5266510681) (2.96875,0.7502665691) (3,0.9181684398)};
\addplot[color=PVteal,densely dotted,line width=.75pt,mark=none] coordinates {(0.03125,0.004742637915) (0.0625,0.006569100198) (0.09375,0.004804954196) (0.125,0.005248875196) (0.15625,0.007086550527) (0.1875,0.006135915626) (0.21875,0.003437702571) (0.25,0.001776461559) (0.28125,0.000576186086) (0.3125,0.001147375346) (0.34375,0.002450317502) (0.375,0.002553524448) (0.40625,0.00171791935) (0.4375,0.0004263797878) (0.46875,0.001010633186) (0.5,0.002608744509) (0.53125,0.004372058467) (0.5625,0.005875010395) (0.59375,0.006628324725) (0.625,0.006850850771) (0.65625,0.007338345902) (0.6875,0.008078153561) (0.71875,0.00844697967) (0.75,0.008967941349) (0.78125,0.01059074335) (0.8125,0.01281230192) (0.84375,0.01481452616) (0.875,0.01704238603) (0.90625,0.01982982744) (0.9375,0.0223660249) (0.96875,0.02414792186) (1,0.02555216916) (1.03125,0.02653262329) (1.0625,0.02643685633) (1.09375,0.02551614722) (1.125,0.02426860246) (1.15625,0.0219253476) (1.1875,0.01865160032) (1.21875,0.01613787465) (1.25,0.01424002231) (1.28125,0.0117180748) (1.3125,0.009608493059) (1.34375,0.009365186661) (1.375,0.009793442558) (1.40625,0.009474030588) (1.4375,0.009510699091) (1.46875,0.01071069177) (1.5,0.01154639479) (1.53125,0.01127962672) (1.5625,0.01121784167) (1.59375,0.01173561192) (1.625,0.01180641292) (1.65625,0.01148912607) (1.6875,0.01173630563) (1.71875,0.01249485166) (1.75,0.0132280408) (1.78125,0.01418968288) (1.8125,0.01578856414) (1.84375,0.01769790689) (1.875,0.01965069922) (1.90625,0.02194484755) (1.9375,0.02439731571) (1.96875,0.02636618334) (2,0.02783599997) (2.03125,0.02897126757) (2.0625,0.02937828335) (2.09375,0.02862114311) (2.125,0.02692443413) (2.15625,0.02503763076) (2.1875,0.02304624272) (2.21875,0.02045788784) (2.25,0.01781295435) (2.28125,0.0161383457) (2.3125,0.0151293721) (2.34375,0.01410327645) (2.375,0.01356310117) (2.40625,0.01395260238) (2.4375,0.01458494668) (2.46875,0.01500272276) (2.5,0.01559587419) (2.53125,0.01638730872) (2.5625,0.01683604441) (2.59375,0.0170115396) (2.625,0.01726533616) (2.65625,0.01725911584) (2.6875,0.01699907717) (2.71875,0.01722864821) (2.75,0.01794245418) (2.78125,0.01864481956) (2.8125,0.01972491586) (2.84375,0.02169398185) (2.875,0.02400158947) (2.90625,0.02596361582) (2.9375,0.02782753515) (2.96875,0.02978080682) (3,0.03112843542)};
\addplot[color=PVgray,dashdotted,line width=.75pt,mark=none] coordinates {(0.03125,0.7238124814) (0.0625,1.581497794) (0.09375,2.505719203) (0.125,3.427327392) (0.15625,4.295984942) (0.1875,5.077183321) (0.21875,5.746955162) (0.25,6.297742887) (0.28125,6.738728164) (0.3125,7.084290369) (0.34375,7.341744323) (0.375,7.511275081) (0.40625,7.591781416) (0.4375,7.575382343) (0.46875,7.441760462) (0.5,7.169655586) (0.53125,6.749994639) (0.5625,6.182096258) (0.59375,5.461403824) (0.625,4.583302908) (0.65625,3.562523935) (0.6875,2.433729469) (0.71875,1.235283092) (0.75,0.01197921565) (0.78125,1.172794817) (0.8125,2.246860616) (0.84375,3.14704837) (0.875,3.821264238) (0.90625,4.228402236) (0.9375,4.348930359) (0.96875,4.191432122) (1,3.787272822) (1.03125,3.184333824) (1.0625,2.438267564) (1.09375,1.59142521) (1.125,0.6709940597) (1.15625,0.2835738331) (1.1875,1.225040427) (1.21875,2.130021431) (1.25,2.988478611) (1.28125,3.774458494) (1.3125,4.461456434) (1.34375,5.05169843) (1.375,5.560427331) (1.40625,5.986557196) (1.4375,6.323508337) (1.46875,6.57529389) (1.5,6.738545989) (1.53125,6.789303835) (1.5625,6.700532677) (1.59375,6.455797433) (1.625,6.044818413) (1.65625,5.466060983) (1.6875,4.738902074) (1.71875,3.905474283) (1.75,3.022962097) (1.78125,2.158762901) (1.8125,1.384260809) (1.84375,0.760495271) (1.875,0.3303716244) (1.90625,0.1217450523) (1.9375,0.1432483246) (1.96875,0.3782469154) (2,0.7948884782) (2.03125,1.3603584) (2.0625,2.040086833) (2.09375,2.786566691) (2.125,3.542868499) (2.15625,4.26463185) (2.1875,4.924274774) (2.21875,5.495763944) (2.25,5.958568963) (2.28125,6.313590799) (2.3125,6.576618585) (2.34375,6.761901814) (2.375,6.881807569) (2.40625,6.949239701) (2.4375,6.968212994) (2.46875,6.927384899) (2.5,6.805475947) (2.53125,6.577252296) (2.5625,6.218261283) (2.59375,5.716613081) (2.625,5.077822796) (2.65625,4.315392248) (2.6875,3.452279801) (2.71875,2.531731799) (2.75,1.60977912) (2.78125,0.7396702979) (2.8125,0.02658072314) (2.84375,0.631243217) (2.875,1.021090175) (2.90625,1.163529373) (2.9375,1.046142121) (2.96875,0.6705987448) (3,0.05840218219)};
\end{axis}
\node[anchor=west,font=\fontsize{9}{11}\selectfont\bfseries] at (79,153) {B\quad Edge flow};
\begin{axis}[at={(95mm,106mm)},anchor=south west,width=55mm,height=38mm,scale only axis,font=\fontsize{8}{10}\selectfont,tick label style={font=\fontsize{8}{9.5}\selectfont},label style={font=\fontsize{8}{10}\selectfont},axis line style={PVgray!65,line width=.45pt},tick style={PVgray!65},grid=major,grid style={PVgray!15,line width=.3pt},enlarge x limits=false,enlarge y limits=false,clip=true,xmin=0,xmax=3,ymin=37.57247993,ymax=58.7516345,xtick={0,1,2,3},xlabel={Cycles},ylabel={mL/s},scaled y ticks=false]
\addplot[color=PVgray,dashed,line width=.75pt,mark=none] coordinates {(0.03125,47.42280742) (0.0625,49.55595075) (0.09375,51.66822142) (0.125,53.58169294) (0.15625,55.14492138) (0.1875,56.25876943) (0.21875,56.88537584) (0.25,57.03866496) (0.28125,56.76331931) (0.3125,56.09673933) (0.34375,55.05683408) (0.375,53.71518504) (0.40625,52.27429309) (0.4375,51.00819547) (0.46875,50.1100746) (0.5,49.62399812) (0.53125,49.49024863) (0.5625,49.54883192) (0.59375,49.48439203) (0.625,48.92895717) (0.65625,47.74071244) (0.6875,46.12645077) (0.71875,44.43298571) (0.75,42.87376725) (0.78125,41.49203014) (0.8125,40.30563578) (0.84375,39.39781563) (0.875,38.88653708) (0.90625,38.87295433) (0.9375,39.42223098) (0.96875,40.54184225) (1,42.15181499) (1.03125,44.11990902) (1.0625,46.39764528) (1.09375,49.06392656) (1.125,52.05724716) (1.15625,54.87938585) (1.1875,56.80115608) (1.21875,57.4511601) (1.25,57.07700808) (1.28125,56.21810765) (1.3125,55.30272546) (1.34375,54.58011481) (1.375,54.17402605) (1.40625,54.03040103) (1.4375,53.93540043) (1.46875,53.71586936) (1.5,53.38454236) (1.53125,53.04539497) (1.5625,52.72105858) (1.59375,52.32650764) (1.625,51.76095451) (1.65625,50.96564919) (1.6875,49.92284346) (1.71875,48.66477088) (1.75,47.2927021) (1.78125,45.94365333) (1.8125,44.70789519) (1.84375,43.6071439) (1.875,42.69317905) (1.90625,42.13612017) (1.9375,42.15963097) (1.96875,42.90080279) (2,44.36710663) (2.03125,46.47909245) (2.0625,49.03286532) (2.09375,51.60782125) (2.125,53.6835767) (2.15625,54.97414789) (2.1875,55.58732718) (2.21875,55.80643631) (2.25,55.79292703) (2.28125,55.54340998) (2.3125,55.03906894) (2.34375,54.33601527) (2.375,53.53495641) (2.40625,52.73908717) (2.4375,52.04383915) (2.46875,51.51155897) (2.5,51.13106949) (2.53125,50.83216426) (2.5625,50.57402272) (2.59375,50.38630887) (2.625,50.25427362) (2.65625,49.98864105) (2.6875,49.31145709) (2.71875,48.09918159) (2.75,46.49540949) (2.78125,44.79641705) (2.8125,43.30251324) (2.84375,42.27330399) (2.875,41.89750266) (2.90625,42.21868212) (2.9375,43.12819411) (2.96875,44.47547856) (3,46.16436206)};
\addplot[color=PVnavy,densely dotted,line width=.75pt,mark=none] coordinates {(0.03125,47.43552001) (0.0625,49.57238845) (0.09375,51.67620105) (0.125,53.58064145) (0.15625,55.13437389) (0.1875,56.22995276) (0.21875,56.83526933) (0.25,56.97574292) (0.28125,56.70390668) (0.3125,56.04642573) (0.34375,54.99155588) (0.375,53.63171087) (0.40625,52.23027311) (0.4375,51.05041437) (0.46875,50.21219413) (0.5,49.73415445) (0.53125,49.5985324) (0.5625,49.65497416) (0.59375,49.51600788) (0.625,48.81454257) (0.65625,47.55430083) (0.6875,46.01429724) (0.71875,44.43244736) (0.75,42.91488436) (0.78125,41.52076757) (0.8125,40.31726785) (0.84375,39.39803991) (0.875,38.88690828) (0.90625,38.89972285) (0.9375,39.49196496) (0.96875,40.63766089) (1,42.23662601) (1.03125,44.1532146) (1.0625,46.38510958) (1.09375,49.11131496) (1.125,52.20524261) (1.15625,54.94453377) (1.1875,56.61750596) (1.21875,57.13408441) (1.25,56.85503082) (1.28125,56.14988316) (1.3125,55.32041082) (1.34375,54.67094785) (1.375,54.32846802) (1.40625,54.14229174) (1.4375,53.92622623) (1.46875,53.64652992) (1.5,53.35267013) (1.53125,53.06789853) (1.5625,52.76012887) (1.59375,52.35193731) (1.625,51.75809569) (1.65625,50.93013411) (1.6875,49.86821943) (1.71875,48.61737969) (1.75,47.27046595) (1.78125,45.94933926) (1.8125,44.72538749) (1.84375,43.60042821) (1.875,42.6566185) (1.90625,42.12179158) (1.9375,42.20982227) (1.96875,43.00068191) (2,44.4855656) (2.03125,46.61310776) (2.0625,49.17040216) (2.09375,51.65878269) (2.125,53.56573213) (2.15625,54.76408938) (2.1875,55.44088946) (2.21875,55.76860186) (2.25,55.79898164) (2.28125,55.54252887) (2.3125,55.02364533) (2.34375,54.30452473) (2.375,53.49634975) (2.40625,52.72465415) (2.4375,52.0766259) (2.46875,51.57786339) (2.5,51.19494211) (2.53125,50.86094673) (2.5625,50.56806294) (2.59375,50.39756558) (2.625,50.30858792) (2.65625,50.01575739) (2.6875,49.24079573) (2.71875,47.96606618) (2.75,46.38484819) (2.78125,44.74229683) (2.8125,43.29781129) (2.84375,42.3277498) (2.875,42.01005398) (2.90625,42.33369244) (2.9375,43.1915986) (2.96875,44.49753965) (3,46.18546744)};
\addplot[color=PVorange,dashed,line width=.75pt,mark=none] coordinates {(0.03125,50.19555268) (0.0625,50.95219716) (0.09375,52.1385368) (0.125,53.36191561) (0.15625,54.39006182) (0.1875,55.12012799) (0.21875,55.52551937) (0.25,55.62922204) (0.28125,55.39824375) (0.3125,54.85466838) (0.34375,54.11756737) (0.375,53.28127942) (0.40625,52.42609035) (0.4375,51.62281201) (0.46875,50.93367273) (0.5,50.40812653) (0.53125,49.92101887) (0.5625,49.30001049) (0.59375,48.51945951) (0.625,47.56552688) (0.65625,46.44666215) (0.6875,45.20160924) (0.71875,43.89960695) (0.75,42.63452921) (0.78125,41.51431885) (0.8125,40.64740848) (0.84375,40.12816833) (0.875,40.02355124) (0.90625,40.36293939) (0.9375,41.13274063) (0.96875,42.27659384) (1,43.70122366) (1.03125,45.50504064) (1.0625,47.65461113) (1.09375,49.83129459) (1.125,51.79378482) (1.15625,53.39364102) (1.1875,54.55361962) (1.21875,55.24686939) (1.25,55.47721845) (1.28125,55.39930941) (1.3125,55.18874496) (1.34375,54.89126379) (1.375,54.56241527) (1.40625,54.24481925) (1.4375,53.95615381) (1.46875,53.68693462) (1.5,53.40424109) (1.53125,53.06003927) (1.5625,52.60253562) (1.59375,51.98862285) (1.625,51.19532837) (1.65625,50.22832793) (1.6875,49.12603296) (1.71875,47.95843872) (1.75,46.82072796) (1.78125,45.74667119) (1.8125,44.79240209) (1.84375,44.07883176) (1.875,43.69849216) (1.90625,43.71376439) (1.9375,44.1642194) (1.96875,45.07186298) (2,46.43957929) (2.03125,48.07960759) (2.0625,49.72498498) (2.09375,51.25650326) (2.125,52.57412615) (2.15625,53.60102994) (2.1875,54.29511193) (2.21875,54.65254851) (2.25,54.70484491) (2.28125,54.5106951) (2.3125,54.14429293) (2.34375,53.68208007) (2.375,53.19006011) (2.40625,52.71366391) (2.4375,52.27171696) (2.46875,51.85539491) (2.5,51.43225669) (2.53125,51.04952817) (2.5625,50.70425747) (2.59375,50.28202698) (2.625,49.70165397) (2.65625,48.92939171) (2.6875,47.97678401) (2.71875,46.89347241) (2.75,45.75473743) (2.78125,44.71354102) (2.8125,43.93575242) (2.84375,43.51206803) (2.875,43.5053825) (2.90625,43.94418363) (2.9375,44.81495043) (2.96875,46.06153579) (3,47.5907239)};
\addplot[color=PVteal,densely dotted,line width=.75pt,mark=none] coordinates {(0.03125,47.44511723) (0.0625,49.5842078) (0.09375,51.68396943) (0.125,53.58766539) (0.15625,55.14005454) (0.1875,56.22425886) (0.21875,56.81713649) (0.25,56.95497721) (0.28125,56.68338855) (0.3125,56.02254875) (0.34375,54.96756694) (0.375,53.61503858) (0.40625,52.22246376) (0.4375,51.04928583) (0.46875,50.21728658) (0.5,49.7438101) (0.53125,49.60750191) (0.5625,49.65779504) (0.59375,49.5090924) (0.625,48.7971964) (0.65625,47.53054016) (0.6875,45.99053854) (0.71875,44.41258013) (0.75,42.89919411) (0.78125,41.50817568) (0.8125,40.30704672) (0.84375,39.39048173) (0.875,38.88316718) (0.90625,38.90063929) (0.9375,39.49692491) (0.96875,40.64447953) (1,42.24239937) (1.03125,44.15587133) (1.0625,46.38545058) (1.09375,49.1115269) (1.125,52.20197442) (1.15625,54.928855) (1.1875,56.58605014) (1.21875,57.09397286) (1.25,56.81648868) (1.28125,56.11964277) (1.3125,55.30140655) (1.34375,54.66253209) (1.375,54.32581743) (1.40625,54.13941096) (1.4375,53.92049724) (1.46875,53.63889868) (1.5,53.34482574) (1.53125,53.06036582) (1.5625,52.7517725) (1.59375,52.34098377) (1.625,51.74393283) (1.65625,50.91345561) (1.6875,49.84994391) (1.71875,48.59868096) (1.75,47.25296747) (1.78125,45.93399327) (1.8125,44.71169476) (1.84375,43.58797545) (1.875,42.64715889) (1.90625,42.11808893) (1.9375,42.21275648) (1.96875,43.00890184) (2,44.49666107) (2.03125,46.62372444) (2.0625,49.17564655) (2.09375,51.6530633) (2.125,53.54700995) (2.15625,54.73663563) (2.1875,55.41128555) (2.21875,55.7406755) (2.25,55.77348477) (2.28125,55.51888442) (2.3125,55.00117811) (2.34375,54.28362791) (2.375,53.47860508) (2.40625,52.71140571) (2.4375,52.06780917) (2.46875,51.57190981) (2.5,51.18936822) (2.53125,50.85372805) (2.5625,50.55921394) (2.59375,50.38817887) (2.625,50.29724602) (2.65625,49.99861805) (2.6875,49.21643743) (2.71875,47.93744385) (2.75,46.3566302) (2.78125,44.71861425) (2.8125,43.2816453) (2.84375,42.3200814) (2.875,42.00907602) (2.90625,42.33595017) (2.9375,43.19435886) (2.96875,44.49966946) (3,46.18657531)};
\addplot[color=PVgray,dashdotted,line width=.75pt,mark=none] coordinates {(0.03125,45.44931039) (0.0625,45.44931039) (0.09375,45.44931039) (0.125,45.44931039) (0.15625,45.44931039) (0.1875,45.44931039) (0.21875,45.44931039) (0.25,45.44931039) (0.28125,45.44931039) (0.3125,45.44931039) (0.34375,45.44931039) (0.375,45.44931039) (0.40625,45.44931039) (0.4375,45.44931039) (0.46875,45.44931039) (0.5,45.44931039) (0.53125,45.44931039) (0.5625,45.44931039) (0.59375,45.44931039) (0.625,45.44931039) (0.65625,45.44931039) (0.6875,45.44931039) (0.71875,45.44931039) (0.75,45.44931039) (0.78125,45.44931039) (0.8125,45.44931039) (0.84375,45.44931039) (0.875,45.44931039) (0.90625,45.44931039) (0.9375,45.44931039) (0.96875,45.44931039) (1,45.44931039) (1.03125,45.44931039) (1.0625,45.44931039) (1.09375,45.44931039) (1.125,45.44931039) (1.15625,45.44931039) (1.1875,45.44931039) (1.21875,45.44931039) (1.25,45.44931039) (1.28125,45.44931039) (1.3125,45.44931039) (1.34375,45.44931039) (1.375,45.44931039) (1.40625,45.44931039) (1.4375,45.44931039) (1.46875,45.44931039) (1.5,45.44931039) (1.53125,45.44931039) (1.5625,45.44931039) (1.59375,45.44931039) (1.625,45.44931039) (1.65625,45.44931039) (1.6875,45.44931039) (1.71875,45.44931039) (1.75,45.44931039) (1.78125,45.44931039) (1.8125,45.44931039) (1.84375,45.44931039) (1.875,45.44931039) (1.90625,45.44931039) (1.9375,45.44931039) (1.96875,45.44931039) (2,45.44931039) (2.03125,45.44931039) (2.0625,45.44931039) (2.09375,45.44931039) (2.125,45.44931039) (2.15625,45.44931039) (2.1875,45.44931039) (2.21875,45.44931039) (2.25,45.44931039) (2.28125,45.44931039) (2.3125,45.44931039) (2.34375,45.44931039) (2.375,45.44931039) (2.40625,45.44931039) (2.4375,45.44931039) (2.46875,45.44931039) (2.5,45.44931039) (2.53125,45.44931039) (2.5625,45.44931039) (2.59375,45.44931039) (2.625,45.44931039) (2.65625,45.44931039) (2.6875,45.44931039) (2.71875,45.44931039) (2.75,45.44931039) (2.78125,45.44931039) (2.8125,45.44931039) (2.84375,45.44931039) (2.875,45.44931039) (2.90625,45.44931039) (2.9375,45.44931039) (2.96875,45.44931039) (3,45.44931039)};
\addplot[color=black,line width=.95pt,mark=none] coordinates {(0.03125,47.43650613) (0.0625,49.57336962) (0.09375,51.6764821) (0.125,53.58100518) (0.15625,55.13395678) (0.1875,56.22731317) (0.21875,56.8315481) (0.25,56.97212382) (0.28125,56.70032505) (0.3125,56.04269592) (0.34375,54.98650563) (0.375,53.62631155) (0.40625,52.22869734) (0.4375,51.05397916) (0.46875,50.21838398) (0.5,49.74102127) (0.53125,49.60608972) (0.5625,49.66185251) (0.59375,49.51610342) (0.625,48.8053275) (0.65625,47.54375712) (0.6875,46.00964964) (0.71875,44.43237948) (0.75,42.91629018) (0.78125,41.52240825) (0.8125,40.31776235) (0.84375,39.39739517) (0.875,38.88733668) (0.90625,38.90268618) (0.9375,39.49719144) (0.96875,40.64384318) (1,42.24137236) (1.03125,44.15362278) (1.0625,46.38369901) (1.09375,49.11785636) (1.125,52.21731093) (1.15625,54.94446099) (1.1875,56.60151816) (1.21875,57.11611902) (1.25,56.84366139) (1.28125,56.14444335) (1.3125,55.321405) (1.34375,54.67913885) (1.375,54.33793274) (1.40625,54.14590069) (1.4375,53.9245807) (1.46875,53.64410014) (1.5,53.35207206) (1.53125,53.06983354) (1.5625,52.76274338) (1.59375,52.35227158) (1.625,51.75636548) (1.65625,50.92821843) (1.6875,49.8657865) (1.71875,48.61436307) (1.75,47.26906319) (1.78125,45.95047813) (1.8125,44.72630026) (1.84375,43.59820571) (1.875,42.65364246) (1.90625,42.12249058) (1.9375,42.21416633) (1.96875,43.00681596) (2,44.49335034) (2.03125,46.62251233) (2.0625,49.17904182) (2.09375,51.65994383) (2.125,53.55636801) (2.15625,54.75203936) (2.1875,55.43393512) (2.21875,55.76617412) (2.25,55.79834409) (2.28125,55.54229453) (2.3125,55.02208355) (2.34375,54.3015319) (2.375,53.49424859) (2.40625,52.72501937) (2.4375,52.07941998) (2.46875,51.58215621) (2.5,51.19847662) (2.53125,50.86133343) (2.5625,50.56697812) (2.59375,50.40020958) (2.625,50.3137706) (2.65625,50.01583162) (2.6875,49.23431158) (2.71875,47.95817583) (2.75,46.37857927) (2.78125,44.73837931) (2.8125,43.29780482) (2.84375,42.33259501) (2.875,42.01740063) (2.90625,42.33964659) (2.9375,43.19492247) (2.96875,44.49956426) (3,46.18756608)};
\end{axis}
\node[anchor=west,font=\fontsize{9}{11}\selectfont\bfseries] at (79,93) {D\quad Flow absolute error};
\begin{axis}[at={(95mm,46mm)},anchor=south west,width=55mm,height=38mm,scale only axis,font=\fontsize{8}{10}\selectfont,tick label style={font=\fontsize{8}{9.5}\selectfont},label style={font=\fontsize{8}{10}\selectfont},axis line style={PVgray!65,line width=.45pt},tick style={PVgray!65},grid=major,grid style={PVgray!15,line width=.3pt},enlarge x limits=false,enlarge y limits=false,clip=true,ymode=log,ymin=1e-06,ymax=1e+23,xmin=0,xmax=3,xtick={0,1,2,3},xlabel={Cycles},ylabel={mL/s}]
\addplot[color=PVblue,solid,line width=.75pt,mark=none] coordinates {(0.03125,1.979686048) (0.0625,4.299534637) (0.09375,6.57083167) (0.125,8.628349287) (0.15625,10.3181224) (0.1875,11.53011493) (0.21875,12.23191656) (0.25,12.44503903) (0.28125,12.21579644) (0.3125,11.56455677) (0.34375,10.47023376) (0.375,9.016506953) (0.40625,7.458117734) (0.4375,6.043660513) (0.46875,4.878975932) (0.5,3.974614001) (0.53125,3.308337584) (0.5625,2.724144041) (0.59375,1.826535049) (0.625,0.2484902751) (0.65625,1.999281852) (0.6875,4.642270471) (0.71875,7.454912588) (0.75,10.33697283) (0.78125,13.23233964) (0.8125,16.07960411) (0.84375,18.79066944) (0.875,21.24835258) (0.90625,23.34913686) (0.9375,25.05417705) (0.96875,26.40991974) (1,27.54353491) (1.03125,28.62526821) (1.0625,29.69754805) (1.09375,30.63518527) (1.125,31.65970209) (1.15625,33.62241216) (1.1875,37.37641211) (1.21875,43.20521417) (1.25,51.03568325) (1.28125,60.87616225) (1.3125,72.89411964) (1.34375,87.37084534) (1.375,104.8985731) (1.40625,126.472764) (1.4375,153.2462488) (1.46875,186.3488136) (1.5,226.9531851) (1.53125,276.3700931) (1.5625,336.0678093) (1.59375,407.6426793) (1.625,492.7624905) (1.65625,593.1187641) (1.6875,710.4197974) (1.71875,846.3988468) (1.75,1002.789981) (1.78125,1181.253717) (1.8125,1383.178677) (1.84375,1609.207137) (1.875,1858.432316) (1.90625,2127.784411) (1.9375,2412.306261) (1.96875,2706.456205) (2,3005.887824) (2.03125,3309.437915) (2.0625,3619.80259) (2.09375,3939.252302) (2.125,4262.790251) (2.15625,4572.753073) (2.1875,4843.782936) (2.21875,5040.223809) (2.25,5145.065802) (2.28125,5207.322973) (2.3125,5294.639176) (2.34375,5323.0877) (2.375,5227.605875) (2.40625,4956.271929) (2.4375,4478.240391) (2.46875,3747.301458) (2.5,2680.983451) (2.53125,1200.591855) (2.5625,792.960704) (2.59375,3412.661606) (2.625,6770.832171) (2.65625,10906.90514) (2.6875,15887.576) (2.71875,21848.02591) (2.75,28886.81241) (2.78125,37138.16217) (2.8125,46756.09838) (2.84375,57908.95375) (2.875,70794.40247) (2.90625,85624.3846) (2.9375,102612.5193) (2.96875,121906.5058) (3,143692.6767)};
\addplot[color=PVorange,solid,line width=.75pt,mark=none] coordinates {(0.03125,0.2473322526) (0.0625,1.217846602) (0.09375,2.279718991) (0.125,3.258142011) (0.15625,3.993679199) (0.1875,4.374476867) (0.21875,4.369515724) (0.25,4.0039891) (0.28125,3.330053413) (0.3125,2.375347073) (0.34375,1.129170182) (0.375,0.3105536139) (0.40625,1.672353484) (0.4375,2.689523839) (0.46875,3.237908835) (0.5,3.289283132) (0.53125,2.849692301) (0.5625,2.061126693) (0.59375,1.30672303) (0.625,0.9414924351) (0.65625,0.9435333427) (0.6875,1.028032009) (0.71875,0.9600591871) (0.75,0.6310240573) (0.78125,0.02297468711) (0.8125,1.070634934) (0.84375,2.607518693) (0.875,4.759034601) (0.90625,7.638844565) (0.9375,11.29873038) (0.96875,15.709318) (1,20.76669544) (1.03125,26.3320821) (1.0625,32.40587933) (1.09375,39.17093438) (1.125,46.48298738) (1.15625,53.59623878) (1.1875,59.80171172) (1.21875,65.01300728) (1.25,69.56891907) (1.28125,73.81122593) (1.3125,78.02283718) (1.34375,82.48640416) (1.375,87.29698148) (1.40625,92.27417634) (1.4375,97.20814517) (1.46875,102.0342489) (1.5,106.7628396) (1.53125,111.3760876) (1.5625,115.7989748) (1.59375,119.9117133) (1.625,123.5919571) (1.65625,126.753269) (1.6875,129.3568542) (1.71875,131.4139816) (1.75,132.9905731) (1.78125,134.1828399) (1.8125,135.0408084) (1.84375,135.5545071) (1.875,135.8041135) (1.90625,136.0151956) (1.9375,136.3937755) (1.96875,137.0153936) (2,137.8713658) (2.03125,138.9111674) (2.0625,139.9246759) (2.09375,140.4191714) (2.125,139.8992687) (2.15625,138.2660806) (2.1875,135.7280711) (2.21875,132.4750282) (2.25,128.5866507) (2.28125,124.1096102) (2.3125,119.1086292) (2.34375,113.6944802) (2.375,108.034248) (2.40625,102.3125279) (2.4375,96.67895256) (2.46875,91.22381467) (2.5,85.97511207) (2.53125,80.92139639) (2.5625,76.10286397) (2.59375,71.63213662) (2.625,67.47082026) (2.65625,63.31709544) (2.6875,58.8714083) (2.71875,54.07973681) (2.75,49.07558236) (2.78125,44.03933595) (2.8125,39.18193305) (2.84375,34.74281168) (2.875,30.85357491) (2.90625,27.44672059) (2.9375,24.34575524) (2.96875,21.36845122) (3,18.31702595)};
\addplot[color=PVred,solid,line width=.75pt,mark=none] coordinates {(0.03125,1.059456121) (0.0625,2.896664367) (0.09375,4.837443877) (0.125,6.70207228) (0.15625,8.328433063) (0.1875,9.601941487) (0.21875,10.48683022) (0.25,11.00096651) (0.28125,11.18593033) (0.3125,11.05607576) (0.34375,10.58366746) (0.375,9.845717281) (0.40625,9.089914117) (0.4375,8.558073425) (0.46875,8.350123982) (0.5,8.473750278) (0.53125,8.909752258) (0.5625,9.51511948) (0.59375,9.923008798) (0.625,9.817649989) (0.65625,9.293342988) (0.6875,8.753695328) (0.71875,8.615758914) (0.75,9.254302407) (0.78125,11.10725715) (0.8125,14.75076341) (0.84375,20.94973542) (0.875,30.69578074) (0.90625,45.2111677) (0.9375,65.94674853) (0.96875,94.55463889) (1,132.7041455) (1.03125,181.6208359) (1.0625,241.5023173) (1.09375,310.9689835) (1.125,386.9535936) (1.15625,466.2404386) (1.1875,547.8807888) (1.21875,633.6442986) (1.25,726.243064) (1.28125,827.5213316) (1.3125,938.4213581) (1.34375,1059.958013) (1.375,1193.649513) (1.40625,1341.523247) (1.4375,1505.717726) (1.46875,1688.092054) (1.5,1890.460711) (1.53125,2115.578867) (1.5625,2371.264993) (1.59375,2667.505075) (1.625,3008.137014) (1.65625,3397.215718) (1.6875,3840.137369) (1.71875,4344.45362) (1.75,4921.608593) (1.78125,5580.860784) (1.8125,6329.102009) (1.84375,7173.91894) (1.875,8122.210687) (1.90625,9163.747785) (1.9375,10285.14601) (1.96875,11484.37062) (2,12755.87416) (2.03125,14119.14357) (2.0625,15594.92619) (2.09375,17152.06686) (2.125,18765.35621) (2.15625,20364.30962) (2.1875,21897.75687) (2.21875,23366.15052) (2.25,24757.21898) (2.28125,25995.77225) (2.3125,27130.98373) (2.34375,28154.26869) (2.375,29017.65969) (2.40625,29606.14623) (2.4375,29772.92135) (2.46875,29475.98482) (2.5,28763.38769) (2.53125,27731.93009) (2.5625,26365.6786) (2.59375,24631.42474) (2.625,22660.97439) (2.65625,20614.71896) (2.6875,18478.60636) (2.71875,16268.34633) (2.75,13968.26206) (2.78125,11580.1787) (2.8125,9092.99026) (2.84375,6451.348566) (2.875,3938.038566) (2.90625,1924.537326) (2.9375,1182.882636) (2.96875,1734.973081) (3,2808.371212)};
\addplot[color=PVteal,solid,line width=.75pt,mark=none] coordinates {(0.03125,0.9896450632) (0.0625,2.119895793) (0.09375,1.871635429) (0.125,0.9279944225) (0.15625,4.431907134) (0.1875,2.255973942) (0.21875,10.90618343) (0.25,25.38707994) (0.28125,10.26027539) (0.3125,58.10468573) (0.34375,128.8514244) (0.375,52.89630786) (0.40625,119.0953051) (0.4375,252.3341281) (0.46875,294.6472751) (0.5,372.5127198) (0.53125,486.8041583) (0.5625,638.9089614) (0.59375,848.711029) (0.625,1163.706184) (0.65625,1657.096094) (0.6875,2421.326082) (0.71875,3670.156109) (0.75,5787.932805) (0.78125,9524.654733) (0.8125,16352.42385) (0.84375,28703.5744) (0.875,51000.05089) (0.90625,91277.62266) (0.9375,164146.8195) (0.96875,295887.5815) (1,533901.8107) (1.03125,964409.202) (1.0625,1743647.353) (1.09375,3154808.641) (1.125,5711249.425) (1.15625,10343604.38) (1.1875,18739058.68) (1.21875,33956463.61) (1.25,61541605.72) (1.28125,111549364.9) (1.3125,202210016.2) (1.34375,366576900.8) (1.375,664578932.6) (1.40625,1204874723) (1.4375,2184475194) (1.46875,3960583511) (1.5,7180858587) (1.53125,1.30195849e+10) (1.5625,2.360589566e+10) (1.59375,4.280020012e+10) (1.625,7.760186701e+10) (1.65625,1.40701806e+11) (1.6875,2.55110131e+11) (1.71875,4.625475393e+11) (1.75,8.386589269e+11) (1.78125,1.520598469e+12) (1.8125,2.757044616e+12) (1.84375,4.998885865e+12) (1.875,9.063642691e+12) (1.90625,1.643358742e+13) (1.9375,2.979628361e+13) (1.96875,5.402464002e+13) (2,9.795390126e+13) (2.03125,1.776035112e+14) (2.0625,3.220187825e+14) (2.09375,5.838630313e+14) (2.125,1.058621522e+15) (2.15625,1.919421919e+15) (2.1875,3.480168142e+15) (2.21875,6.31000994e+15) (2.25,1.144088933e+16) (2.28125,2.074385776e+16) (2.3125,3.761138242e+16) (2.34375,6.819445729e+16) (2.375,1.236456694e+17) (2.40625,2.241860771e+17) (2.4375,4.064792579e+17) (2.46875,7.370010087e+17) (2.5,1.336281576e+18) (2.53125,2.422857047e+18) (2.5625,4.392963881e+18) (2.59375,7.965028949e+18) (2.625,1.444165992e+19) (2.65625,2.618466424e+19) (2.6875,4.74762992e+19) (2.71875,8.608087825e+19) (2.75,1.560761085e+20) (2.78125,2.829867474e+20) (2.8125,5.130927692e+20) (2.84375,9.303056433e+20) (2.875,1.686768291e+21) (2.90625,3.058336414e+21) (2.9375,5.545171968e+21) (2.96875,1.005413751e+22) (3,1.822948999e+22)};
\addplot[color=PVgray,dashed,line width=.75pt,mark=none] coordinates {(0.03125,0.01369871149) (0.0625,0.017418871) (0.09375,0.008260679979) (0.125,0.0006877517093) (0.15625,0.01096460038) (0.1875,0.03145626042) (0.21875,0.05382774042) (0.25,0.06654113637) (0.28125,0.06299425253) (0.3125,0.0540434021) (0.34375,0.07032844199) (0.375,0.08887349913) (0.40625,0.04559574903) (0.4375,0.04578368476) (0.46875,0.10830938) (0.5,0.1170231484) (0.53125,0.1158410875) (0.5625,0.1130205958) (0.59375,0.03171139047) (0.625,0.1236296644) (0.65625,0.1969553222) (0.6875,0.1168011287) (0.71875,0.0006062247485) (0.75,0.04252292745) (0.78125,0.03037810848) (0.8125,0.01212656616) (0.84375,0.0004204517075) (0.875,0.0007996067747) (0.90625,0.02973184867) (0.9375,0.07496046257) (0.96875,0.1020009261) (1,0.08955737243) (1.03125,0.03371375439) (1.0625,0.01394626559) (1.09375,0.05392979411) (1.125,0.1600637756) (1.15625,0.06507514329) (1.1875,0.1996379133) (1.21875,0.3350410739) (1.25,0.2333466915) (1.28125,0.07366430845) (1.3125,0.01867953851) (1.34375,0.09902404177) (1.375,0.1639066862) (1.40625,0.1154996559) (1.4375,0.01081972905) (1.46875,0.07176921654) (1.5,0.03247029285) (1.53125,0.0244385673) (1.5625,0.04168480052) (1.59375,0.02576393505) (1.625,0.004589030241) (1.65625,0.03743075925) (1.6875,0.05705695104) (1.71875,0.05040780829) (1.75,0.02363891581) (1.78125,0.00682480487) (1.8125,0.01840506781) (1.84375,0.008938183318) (1.875,0.03953659244) (1.90625,0.01362958721) (1.9375,0.0545353505) (1.96875,0.1060131685) (2,0.126243717) (2.03125,0.1434198818) (2.0625,0.1461765035) (2.09375,0.05212258582) (2.125,0.1272086984) (2.15625,0.2221085303) (2.1875,0.1533920586) (2.21875,0.04026219205) (2.25,0.005417052481) (2.28125,0.001115452001) (2.3125,0.01698539335) (2.34375,0.03448336386) (2.375,0.04070782335) (2.40625,0.01406780647) (2.4375,0.03558082568) (2.46875,0.07059724109) (2.5,0.06740712831) (2.53125,0.02916916921) (2.5625,0.007044601735) (2.59375,0.01390070695) (2.625,0.05949697744) (2.65625,0.02719056644) (2.6875,0.07714551162) (2.71875,0.1410057595) (2.75,0.1168302179) (2.78125,0.05803773385) (2.8125,0.004708416775) (2.84375,0.05929102006) (2.875,0.1198979646) (2.90625,0.1209644636) (2.9375,0.06672836404) (2.96875,0.0240856954) (3,0.02320401672)};
\addplot[color=PVnavy,densely dotted,line width=.75pt,mark=none] coordinates {(0.03125,0.0009861126743) (0.0625,0.0009811700932) (0.09375,0.0002810444523) (0.125,0.0003637296755) (0.15625,0.0004171105342) (0.1875,0.002639585554) (0.21875,0.00372123101) (0.25,0.003619100655) (0.28125,0.003581630144) (0.3125,0.003729806741) (0.34375,0.005050242768) (0.375,0.005399326026) (0.40625,0.001575765154) (0.4375,0.003564789286) (0.46875,0.006189846857) (0.5,0.006866814291) (0.53125,0.007557321203) (0.5625,0.006878352941) (0.59375,9.554065127e-05) (0.625,0.009215066249) (0.65625,0.01054370478) (0.6875,0.004647598152) (0.71875,6.787892877e-05) (0.75,0.001405815731) (0.78125,0.001640682988) (0.8125,0.0004944966975) (0.84375,0.00064474113) (0.875,0.0004284055975) (0.90625,0.002963332648) (0.9375,0.005226478792) (0.96875,0.006182287813) (1,0.004746344607) (1.03125,0.0004081745732) (1.0625,0.001410568474) (1.09375,0.006541399314) (1.125,0.01206832002) (1.15625,7.2777703e-05) (1.1875,0.01598779083) (1.21875,0.01796539312) (1.25,0.01136942699) (1.28125,0.005439814823) (1.3125,0.000994180179) (1.34375,0.008190997367) (1.375,0.009464722504) (1.40625,0.003608944101) (1.4375,0.001645529037) (1.46875,0.002429776751) (1.5,0.0005980690052) (1.53125,0.001935001541) (1.5625,0.002614516074) (1.59375,0.000334271339) (1.625,0.001730212144) (1.65625,0.001915681387) (1.6875,0.002432923147) (1.71875,0.003016616018) (1.75,0.001402765475) (1.78125,0.001138873107) (1.8125,0.0009127695247) (1.84375,0.002222500448) (1.875,0.002976043014) (1.90625,0.0006990016908) (1.9375,0.004344052812) (1.96875,0.006134048002) (2,0.007784746898) (2.03125,0.009404566792) (2.0625,0.008639668658) (2.09375,0.001161138513) (2.125,0.009364119271) (2.15625,0.01205001518) (2.1875,0.006954345319) (2.21875,0.002427744637) (2.25,0.0006375500837) (2.28125,0.0002343350243) (2.3125,0.001561777588) (2.34375,0.002992828116) (2.375,0.002101160647) (2.40625,0.0003652143502) (2.4375,0.002794081847) (2.46875,0.004292826638) (2.5,0.003534505402) (2.53125,0.0003867022486) (2.5625,0.001084820736) (2.59375,0.002643996793) (2.625,0.005182682455) (2.65625,7.423096977e-05) (2.6875,0.006484155142) (2.71875,0.007890351557) (2.75,0.006268924164) (2.78125,0.003917521622) (2.8125,6.470597384e-06) (2.84375,0.00484520772) (2.875,0.007346646618) (2.90625,0.005954153099) (2.9375,0.0033238789) (2.96875,0.002024607912) (3,0.002098635895)};
\addplot[color=PVorange,dashed,line width=.75pt,mark=none] coordinates {(0.03125,2.759046551) (0.0625,1.378827541) (0.09375,0.4620547049) (0.125,0.2190895702) (0.15625,0.7438949614) (0.1875,1.107185187) (0.21875,1.306028734) (0.25,1.342901788) (0.28125,1.302081304) (0.3125,1.18802754) (0.34375,0.8689382647) (0.375,0.3450321302) (0.40625,0.1973930068) (0.4375,0.5688328528) (0.46875,0.7152887549) (0.5,0.6671052622) (0.53125,0.3149291506) (0.5625,0.3618420196) (0.59375,0.9966439105) (0.625,1.239800627) (0.65625,1.09709497) (0.6875,0.8080404062) (0.71875,0.5327725332) (0.75,0.2817609719) (0.78125,0.008089395623) (0.8125,0.3296461306) (0.84375,0.730773159) (0.875,1.136214559) (0.90625,1.460253208) (0.9375,1.635549192) (0.96875,1.632750662) (1,1.459851299) (1.03125,1.351417868) (1.0625,1.270912118) (1.09375,0.7134382305) (1.125,0.4235261137) (1.15625,1.550819974) (1.1875,2.047898541) (1.21875,1.869249633) (1.25,1.36644294) (1.28125,0.7451339328) (1.3125,0.132660039) (1.34375,0.2121249417) (1.375,0.2244825306) (1.40625,0.09891855933) (1.4375,0.03157311463) (1.46875,0.04283447937) (1.5,0.05216902379) (1.53125,0.009794269511) (1.5625,0.1602077614) (1.59375,0.3636487318) (1.625,0.5610371115) (1.65625,0.6998905015) (1.6875,0.7397535442) (1.71875,0.6559243481) (1.75,0.4483352294) (1.78125,0.2038069426) (1.8125,0.06610182994) (1.84375,0.4806260413) (1.875,1.044849695) (1.90625,1.591273802) (1.9375,1.950053077) (1.96875,2.065047021) (2,1.946228946) (2.03125,1.457095261) (2.0625,0.5459431549) (2.09375,0.4034405676) (2.125,0.9822418531) (2.15625,1.151009419) (2.1875,1.138823191) (2.21875,1.113625603) (2.25,1.093499172) (2.28125,1.031599433) (2.3125,0.8777906197) (2.34375,0.6194518372) (2.375,0.3041884774) (2.40625,0.01135545295) (2.4375,0.1922969802) (2.46875,0.2732386978) (2.5,0.2337800753) (2.53125,0.1881947419) (2.5625,0.1372793554) (2.59375,0.1181825989) (2.625,0.6121166317) (2.65625,1.086439913) (2.6875,1.25752757) (2.71875,1.064703426) (2.75,0.6238418407) (2.78125,0.02483828891) (2.8125,0.6379476014) (2.84375,1.17947302) (2.875,1.48798187) (2.90625,1.604537043) (2.9375,1.62002796) (2.96875,1.561971533) (3,1.403157816)};
\addplot[color=PVteal,densely dotted,line width=.75pt,mark=none] coordinates {(0.03125,0.00861110124) (0.0625,0.01083817564) (0.09375,0.007487336809) (0.125,0.006660206692) (0.15625,0.006097755316) (0.1875,0.003054310321) (0.21875,0.01441160758) (0.25,0.01714661836) (0.28125,0.01693650856) (0.3125,0.02014717733) (0.34375,0.01893869317) (0.375,0.011272962) (0.40625,0.006233582522) (0.4375,0.004693329162) (0.46875,0.001097394118) (0.5,0.002788831232) (0.53125,0.001412184916) (0.5625,0.004057469402) (0.59375,0.007011024161) (0.625,0.008131096986) (0.65625,0.01321696514) (0.6875,0.0191111031) (0.71875,0.01979935368) (0.75,0.01709607118) (0.78125,0.014232572) (0.8125,0.0107156338) (0.84375,0.006913447071) (0.875,0.00416950453) (0.90625,0.002046890099) (0.9375,0.0002665265617) (0.96875,0.0006363521062) (1,0.001027011351) (1.03125,0.002248557155) (1.0625,0.001751563433) (1.09375,0.006329460711) (1.125,0.01533651691) (1.15625,0.01560599497) (1.1875,0.01546802311) (1.21875,0.02214616178) (1.25,0.02717271741) (1.28125,0.02480057297) (1.3125,0.01999844086) (1.34375,0.0166067623) (1.375,0.0121153063) (1.40625,0.00648973038) (1.4375,0.004083460072) (1.46875,0.005201468496) (1.5,0.007246321664) (1.53125,0.009467716334) (1.5625,0.01097088061) (1.59375,0.01128780544) (1.625,0.01243265314) (1.65625,0.01476282096) (1.6875,0.01584259498) (1.71875,0.01568210801) (1.75,0.01609571703) (1.78125,0.01648485622) (1.8125,0.01460549404) (1.84375,0.01023026416) (1.875,0.00648357149) (1.90625,0.004401649194) (1.9375,0.001409849286) (1.96875,0.00208588387) (2,0.003310726384) (2.03125,0.001212114306) (2.0625,0.003395277519) (2.09375,0.006880536427) (2.125,0.009358058841) (2.15625,0.01540372973) (2.1875,0.02264956268) (2.21875,0.02549861877) (2.25,0.02485932011) (2.28125,0.02341010987) (2.3125,0.02090544078) (2.34375,0.01790399301) (2.375,0.01564350883) (2.40625,0.0136136579) (2.4375,0.01161080954) (2.46875,0.01024640482) (2.5,0.009108394883) (2.53125,0.007605377599) (2.5625,0.007764174491) (2.59375,0.01203070824) (2.625,0.01652458559) (2.65625,0.0172135739) (2.6875,0.01787414234) (2.71875,0.02073197891) (2.75,0.02194906901) (2.78125,0.0197650594) (2.8125,0.01615951665) (2.84375,0.01251360425) (2.875,0.00832460533) (2.90625,0.003696421042) (2.9375,0.0005636103999) (2.96875,0.0001051965176) (3,0.0009907711871)};
\addplot[color=PVgray,dashdotted,line width=.75pt,mark=none] coordinates {(0.03125,1.987195739) (0.0625,4.124059233) (0.09375,6.227171709) (0.125,8.131694795) (0.15625,9.684646395) (0.1875,10.77800278) (0.21875,11.38223771) (0.25,11.52281343) (0.28125,11.25101467) (0.3125,10.59338553) (0.34375,9.537195246) (0.375,8.177001157) (0.40625,6.779386953) (0.4375,5.604668768) (0.46875,4.769073588) (0.5,4.29171088) (0.53125,4.156779333) (0.5625,4.212542125) (0.59375,4.066793032) (0.625,3.356017113) (0.65625,2.094446733) (0.6875,0.5603392561) (0.71875,1.016930908) (0.75,2.533020212) (0.78125,3.92690214) (0.8125,5.131548039) (0.84375,6.051915215) (0.875,6.561973707) (0.90625,6.546624211) (0.9375,5.952118949) (0.96875,4.805467208) (1,3.20793803) (1.03125,1.295687614) (1.0625,0.934388625) (1.09375,3.668545967) (1.125,6.768000546) (1.15625,9.495150603) (1.1875,11.15220778) (1.21875,11.66680863) (1.25,11.394351) (1.28125,10.69513296) (1.3125,9.872094606) (1.34375,9.229828459) (1.375,8.888622352) (1.40625,8.6965903) (1.4375,8.475270309) (1.46875,8.194789756) (1.5,7.902761675) (1.53125,7.620523147) (1.5625,7.313432993) (1.59375,6.902961189) (1.625,6.307055094) (1.65625,5.47890804) (1.6875,4.416476115) (1.71875,3.165052681) (1.75,1.819752797) (1.78125,0.5011677417) (1.8125,0.7230101297) (1.84375,1.851104675) (1.875,2.795667928) (1.90625,3.326819805) (1.9375,3.235144063) (1.96875,2.442494429) (2,0.9559600439) (2.03125,1.173201941) (2.0625,3.729731436) (2.09375,6.210633444) (2.125,8.107057617) (2.15625,9.302728975) (2.1875,9.984624728) (2.21875,10.31686373) (2.25,10.3490337) (2.28125,10.09298414) (2.3125,9.572773162) (2.34375,8.852221515) (2.375,8.044938199) (2.40625,7.275708977) (2.4375,6.630109592) (2.46875,6.132845825) (2.5,5.74916623) (2.53125,5.412023043) (2.5625,5.117667727) (2.59375,4.950899193) (2.625,4.864460213) (2.65625,4.566521231) (2.6875,3.785001187) (2.71875,2.508865444) (2.75,0.9292688801) (2.78125,0.7109310763) (2.8125,2.151505568) (2.84375,3.116715382) (2.875,3.431909761) (2.90625,3.1096638) (2.9375,2.254387914) (2.96875,0.9497461298) (3,0.7382556921)};
\end{axis}
\node[align=center,text width=150mm] at (76,29) {Prespecified case: atlas 79, background 80, program A, neural seed 211.};
\draw[PVblue,solid,line width=.8pt] (1,17)--(8,17);
\node[anchor=west,font=\fontsize{8}{10}\selectfont] at (9,17) {Mono JEPA};
\draw[PVorange,solid,line width=.8pt] (53,17)--(60,17);
\node[anchor=west,font=\fontsize{8}{10}\selectfont] at (61,17) {OPF ($K=1$)};
\draw[PVred,solid,line width=.8pt] (105,17)--(112,17);
\node[anchor=west,font=\fontsize{8}{10}\selectfont] at (113,17) {OPF ($K=4$)};
\draw[PVteal,solid,line width=.8pt] (1,11)--(8,11);
\node[anchor=west,font=\fontsize{8}{10}\selectfont] at (9,11) {Supervised GNN};
\draw[PVgray,dashed,line width=.8pt] (53,11)--(60,11);
\node[anchor=west,font=\fontsize{8}{10}\selectfont] at (61,11) {RLC, 1 substep};
\draw[PVnavy,densely dotted,line width=.8pt] (105,11)--(112,11);
\node[anchor=west,font=\fontsize{8}{10}\selectfont] at (113,11) {RLC, 4 substeps};
\draw[PVorange,dashed,line width=.8pt] (1,5)--(8,5);
\node[anchor=west,font=\fontsize{8}{10}\selectfont] at (9,5) {Reduced RC};
\draw[PVteal,densely dotted,line width=.8pt] (53,5)--(60,5);
\node[anchor=west,font=\fontsize{8}{10}\selectfont] at (61,5) {Fitted RLC};
\draw[PVgray,dashdotted,line width=.8pt] (105,5)--(112,5);
\node[anchor=west,font=\fontsize{8}{10}\selectfont] at (113,5) {Persistence};
\end{tikzpicture}
\end{checkedfigure}
\caption{A case selected before viewing test outputs. (A,B) The black line is the reference; known-parameter RLC, reduced RC, fitted RLC and persistence predictions are shown on physical axes. Pressure is shown on the specified reference scale, including the 5~mmHg distal datum. (C,D) Absolute errors include every method, including all neural predictions, over the complete finite logarithmic range. This separates readable reference dynamics from potentially very large neural errors without truncating unstable predictions. No representative or favourable example was selected retrospectively. Zero errors, if any, are displayed at the recorded positive log floor; nonfinite values are flagged explicitly.}
\label{fig:prespecified-example}
\end{figure}
