## Supplementary Data - Compact audit, summary tables, code, and figure sources for "Panvascular-JEPA: Unseen-Structure Evaluation of Factorized Predictive States for Hemodynamic Networks": validation_figure.tex

\begin{figure}[!tp]
\centering
\begin{checkedfigure}
\begin{tikzpicture}[x=1mm,y=1mm,font=\fontsize{8}{10}\selectfont,text=PVnavy,line cap=round,line join=round]
\path[use as bounding box] (0,0) rectangle (152,143);
\node[anchor=west,font=\fontsize{9}{11}\selectfont\bfseries] at (1,140) {A\quad Mono JEPA};
\begin{axis}[at={(17mm,94mm)},anchor=south west,width=55mm,height=37mm,scale only axis,font=\fontsize{8}{10}\selectfont,tick label style={font=\fontsize{8}{9.5}\selectfont},label style={font=\fontsize{8}{10}\selectfont},axis line style={PVgray!65,line width=.45pt},tick style={PVgray!65},grid=major,grid style={PVgray!15,line width=.3pt},enlarge x limits=false,enlarge y limits=false,clip=true,ymode=log,ymin=10,ymax=1e+11,xmin=.5,xmax=6.5,xtick={1,2,3,4,5,6},xlabel={Candidate},ylabel={Validation RMSE}]
\addplot+[only marks,mark=*,mark size=2pt,color=PVblue] coordinates {(1,41185.97264)};
\addplot+[only marks,mark=*,mark size=2pt,color=PVblue] coordinates {(2,41049.8105)};
\addplot+[only marks,mark=*,mark size=2pt,color=PVblue] coordinates {(3,137116.1389)};
\addplot+[only marks,mark=*,mark size=2pt,color=PVblue] coordinates {(4,136834.5585)};
\addplot+[only marks,mark=*,mark size=2pt,color=PVblue] coordinates {(5,7.215732641e+10)};
\addplot+[only marks,mark=*,mark size=2pt,color=PVblue] coordinates {(6,7.239629828e+10)};
\addplot+[only marks,mark=o,mark size=3.7pt,color=PVnavy,line width=.8pt] coordinates {(2,41049.8105)};
\end{axis}
\node[anchor=west,font=\fontsize{9}{11}\selectfont\bfseries] at (79,140) {B\quad OPF ($K=1$)};
\begin{axis}[at={(95mm,94mm)},anchor=south west,width=55mm,height=37mm,scale only axis,font=\fontsize{8}{10}\selectfont,tick label style={font=\fontsize{8}{9.5}\selectfont},label style={font=\fontsize{8}{10}\selectfont},axis line style={PVgray!65,line width=.45pt},tick style={PVgray!65},grid=major,grid style={PVgray!15,line width=.3pt},enlarge x limits=false,enlarge y limits=false,clip=true,ymode=log,ymin=10,ymax=1e+11,xmin=.5,xmax=6.5,xtick={1,2,3,4,5,6},xlabel={Candidate},ylabel={Validation RMSE}]
\addplot+[only marks,mark=*,mark size=2pt,color=PVorange] coordinates {(1,555477.0206)};
\addplot+[only marks,mark=*,mark size=2pt,color=PVorange] coordinates {(2,554104.9077)};
\addplot+[only marks,mark=*,mark size=2pt,color=PVorange] coordinates {(3,112756.1258)};
\addplot+[only marks,mark=*,mark size=2pt,color=PVorange] coordinates {(4,116135.7655)};
\addplot+[only marks,mark=*,mark size=2pt,color=PVorange] coordinates {(5,251017.0612)};
\addplot+[only marks,mark=*,mark size=2pt,color=PVorange] coordinates {(6,271366.7338)};
\addplot+[only marks,mark=o,mark size=3.7pt,color=PVnavy,line width=.8pt] coordinates {(3,112756.1258)};
\end{axis}
\node[anchor=west,font=\fontsize{9}{11}\selectfont\bfseries] at (1,87) {C\quad OPF ($K=4$)};
\begin{axis}[at={(17mm,41mm)},anchor=south west,width=55mm,height=37mm,scale only axis,font=\fontsize{8}{10}\selectfont,tick label style={font=\fontsize{8}{9.5}\selectfont},label style={font=\fontsize{8}{10}\selectfont},axis line style={PVgray!65,line width=.45pt},tick style={PVgray!65},grid=major,grid style={PVgray!15,line width=.3pt},enlarge x limits=false,enlarge y limits=false,clip=true,ymode=log,ymin=10,ymax=1e+11,xmin=.5,xmax=6.5,xtick={1,2,3,4,5,6},xlabel={Candidate},ylabel={Validation RMSE}]
\addplot+[only marks,mark=*,mark size=2pt,color=PVred] coordinates {(1,1.522993455e+10)};
\addplot+[only marks,mark=*,mark size=2pt,color=PVred] coordinates {(2,1.528919894e+10)};
\addplot+[only marks,mark=*,mark size=2pt,color=PVred] coordinates {(3,1253234308)};
\addplot+[only marks,mark=*,mark size=2pt,color=PVred] coordinates {(4,1290650781)};
\addplot+[only marks,mark=*,mark size=2pt,color=PVred] coordinates {(5,3303367.232)};
\addplot+[only marks,mark=*,mark size=2pt,color=PVred] coordinates {(6,3186307.599)};
\addplot+[only marks,mark=o,mark size=3.7pt,color=PVnavy,line width=.8pt] coordinates {(6,3186307.599)};
\end{axis}
\node[anchor=west,font=\fontsize{9}{11}\selectfont\bfseries] at (79,87) {D\quad Supervised GNN};
\begin{axis}[at={(95mm,41mm)},anchor=south west,width=55mm,height=37mm,scale only axis,font=\fontsize{8}{10}\selectfont,tick label style={font=\fontsize{8}{9.5}\selectfont},label style={font=\fontsize{8}{10}\selectfont},axis line style={PVgray!65,line width=.45pt},tick style={PVgray!65},grid=major,grid style={PVgray!15,line width=.3pt},enlarge x limits=false,enlarge y limits=false,clip=true,ymode=log,ymin=10,ymax=1e+11,xmin=.5,xmax=6.5,xtick={1,2,3,4,5,6},xlabel={Candidate},ylabel={Validation RMSE}]
\addplot+[only marks,mark=*,mark size=2pt,color=PVteal] coordinates {(1,29159920.68)};
\addplot+[only marks,mark=*,mark size=2pt,color=PVteal] coordinates {(2,29128478.68)};
\addplot+[only marks,mark=*,mark size=2pt,color=PVteal] coordinates {(3,45.42623979)};
\addplot+[only marks,mark=*,mark size=2pt,color=PVteal] coordinates {(4,45.40187519)};
\addplot+[only marks,mark=*,mark size=2pt,color=PVteal] coordinates {(5,30857.20375)};
\addplot+[only marks,mark=*,mark size=2pt,color=PVteal] coordinates {(6,28103.2001)};
\addplot+[only marks,mark=o,mark size=3.7pt,color=PVnavy,line width=.8pt] coordinates {(4,45.40187519)};
\end{axis}
\node[align=center,text width=150mm,font=\fontsize{8}{10}\selectfont] at (76,18) {Candidates 1/2: learning rate $3\!\times\!10^{-4}$; 3/4: $10^{-3}$; 5/6: $3\!\times\!10^{-3}$.\\Odd candidates: weight decay 0; even candidates: $10^{-4}$.\\Outlined points were selected using validation data only.};
\end{tikzpicture}
\end{checkedfigure}
\caption{Equal-budget development search. Each architecture used the same six learning-rate/weight-decay combinations, 1,000 updates per candidate and development seed 101. The vertical log scale is identical across panels and displays the complete finite range. Crosses, if present, denote nonfinite or failed candidates rather than omitted trials. Selection uses the 96-step equal-structure macro-RMSE on new parameter backgrounds from seen structures; these are not held-out-structure test results.}
\label{fig:validation-search}
\end{figure}
