## Supplementary Data - Compact audit, summary tables, code, and figure sources for "Panvascular-JEPA: Unseen-Structure Evaluation of Factorized Predictive States for Hemodynamic Networks": generate_network_data.py

```
"""Fresh, custom nodal RLC graph benchmark; no patient or prior-manuscript data.

This is not a replication of the internal circuit layout of svZeroDSolver's
BloodVessel block. For B positive at an edge's source and negative at its target,
C p' = u - B q - g p / m_distal and L q' = B.T p - R m_edge q.
The stored p is gauge pressure above fixed distal pressure Pv=5 mmHg; absolute
pressure is p+5. Units: pressure mmHg, flow mL/s, time s. Ranges are dimensioned
engineering ranges, not calibration to any patient's anatomy or vascular bed.
"""
from __future__ import annotations
import os
for _name in ("OMP_NUM_THREADS", "OPENBLAS_NUM_THREADS", "MKL_NUM_THREADS"):
    os.environ.setdefault(_name, "1")
import argparse
import hashlib
import json
import time
from concurrent.futures import ProcessPoolExecutor, as_completed
from pathlib import Path
import networkx as nx
import numpy as np
from scipy.integrate import solve_ivp

ROOT = Path(__file__).resolve().parent
DATA = ROOT / "data"
SEED = 2026101101
N, EMAX, STEPS = 6, 7, 96
CONTROL_WIDTH = 1 + EMAX + N
SPLIT_COUNTS = {"train": 12, "val": 4, "id_test": 4, "ood_topology": 12}
EDGE_TARGETS = [[.65, 1.30, .85, 1.15], [1.40, .80, 1.25, .90]]
RUNOFF_TARGETS = [[.80, 1.20, .90, 1.10], [1.25, .75, 1.15, .85]]

def write_json(path, obj):
    Path(path).write_text(json.dumps(obj, indent=2, allow_nan=False), encoding="utf-8")

def sha256(path):
    return hashlib.sha256(Path(path).read_bytes()).hexdigest()

def incidence(edge_index, edge_mask):
    out = np.zeros((N, EMAX), float)
    for j in np.flatnonzero(edge_mask):
        out[int(edge_index[0, j]), j] = 1.
        out[int(edge_index[1, j]), j] = -1.
    return out

def build_system(node_features, edge_features, edge_index, edge_mask, control,
                 coefficients=None):
    """Return A,b for x'=Ax+b. Five optional coefficients support local fitting.

    Coefficients multiply inflow, incidence, runoff, pressure gradient and edge
    resistance, respectively; all ones give the known-parameter mechanism.
    """
    nf, ef, ctrl = np.asarray(node_features), np.asarray(edge_features), np.asarray(control)
    coef = np.ones(5) if coefficients is None else np.asarray(coefficients)
    C, g, inlet = nf.T
    R, L = ef.T
    B = incidence(edge_index, edge_mask)
    A = np.zeros((N + EMAX, N + EMAX), float)
    A[:N, :N] = -np.diag(coef[2] * g / ctrl[1+EMAX:] / C)
    A[:N, N:] = -coef[1] * B / C[:, None]
    A[N:, :N] = coef[3] * B.T / L[:, None]
    A[N:, N:] = -np.diag(coef[4] * R * ctrl[1:1+EMAX] / L)
    # Inactive dummy entries are identically zero and excluded from every metric.
    for j in np.flatnonzero(~np.asarray(edge_mask, bool)):
        A[N+j, N+j] = -1.
    b = np.r_[coef[0] * inlet * ctrl[0] / C, np.zeros(EMAX)]
    return A, b

def integrate_midpoint(initial, node_features, edge_features, edge_index,
                       edge_mask, controls, dt, substeps=1, coefficients=None):
    """Known information only; controls are linear within each observed interval."""
    x = np.asarray(initial, float).copy()
    out = [x.copy()]
    h = float(dt) / int(substeps)
    eye = np.eye(N+EMAX)
    for c0, c1 in zip(controls[:-1], controls[1:]):
        for s in range(substeps):
            ctrl = c0 + (c1-c0) * ((s+.5)/substeps)
            A, b = build_system(node_features, edge_features, edge_index, edge_mask, ctrl, coefficients)
            x = np.linalg.solve(eye-.5*h*A, (eye+.5*h*A) @ x + h*b)
        out.append(x.copy())
    return np.asarray(out)

def integrate_rc(initial, node_features, edge_features, edge_index, edge_mask,
                 controls, dt, substeps=4):
    """Prespecified mechanism reduction L=0; retains known C,R,runoff and graph."""
    C, g, inlet = np.asarray(node_features).T
    R = np.asarray(edge_features)[:, 0]
    B = incidence(edge_index, edge_mask)
    p = np.asarray(initial[:N], float).copy()
    out = [np.asarray(initial, float).copy()]
    h, eye = float(dt)/substeps, np.eye(N)
    for c0, c1 in zip(controls[:-1], controls[1:]):
        for s in range(substeps):
            ctrl = c0 + (c1-c0)*((s+.5)/substeps)
            invR = np.divide(edge_mask, R*ctrl[1:1+EMAX], out=np.zeros(EMAX), where=R>0)
            A = -(B @ np.diag(invR) @ B.T + np.diag(g/ctrl[1+EMAX:]))/C[:, None]
            b = inlet*ctrl[0]/C
            p = np.linalg.solve(eye-.5*h*A, (eye+.5*h*A)@p+h*b)
        invR = np.divide(edge_mask, R*c1[1:1+EMAX], out=np.zeros(EMAX), where=R>0)
        q = invR*(B.T@p)
        out.append(np.r_[p,q])
    return np.asarray(out)

def freeze_design():
    DATA.mkdir(parents=True, exist_ok=True)
    manifest = DATA / "design_manifest.json"
    if manifest.exists():
        return json.loads(manifest.read_text(encoding="utf-8"))
    rng = np.random.default_rng(SEED)
    atlas = [(i,g) for i,g in enumerate(nx.graph_atlas_g())
             if len(g)==N and nx.is_connected(g) and g.number_of_edges() in (5,6,7)]
    structures = []
    for ecount, ntrain in [(5,4),(6,8),(7,12)]:
        group = [(i,g) for i,g in atlas if g.number_of_edges()==ecount]
        order = rng.permutation(len(group))
        assert len(group) == {5:6,6:13,7:19}[ecount]
        for rank, idx in enumerate(order):
            atlas_id, graph = group[int(idx)]
            d = dict(nx.shortest_path_length(graph, source=0))
            outlets = sorted(range(1,N), key=lambda j:(-d[j],j))[:2]
            structures.append({"graph_id": f"atlas_{atlas_id}", "atlas_id":atlas_id,
                "edges":sorted([list(e) for e in graph.edges()]), "num_edges":ecount,
                "structure_split":"seen" if rank<ntrain else "heldout",
                "root":0,"outlets":outlets, "cycle_rank":ecount-N+1})
    rows = []
    bid = 0
    for structure in structures:
        splits = ("train","val","id_test") if structure["structure_split"]=="seen" else ("ood_topology",)
        for split in splits:
            for local in range(SPLIT_COUNTS[split]):
                # Each background has its own independently seeded parameters;
                # all seeds and IDs are fixed before generating any outcomes.
                rows.append({"background_id":bid,"graph_id":structure["graph_id"],
                    "split":split,"parameter_seed":SEED+10000+bid,
                    "replicate":local,"arms":[0,1]})
                bid += 1
    result = {"version":"fresh-nodal-rlc-v1", "seed":SEED,"networkx_version":nx.__version__,
        "n_nodes":N,"max_edges":EMAX,"steps":STEPS,"cycles":3,"steps_per_cycle":32,
        "input_interpolation":"piecewise linear between every observed control sample",
        "structural_sampling":"all connected, nonisomorphic six-node graph-atlas graphs with 5,6,7 edges",
        "control_columns":["Qin"]+[f"edge_R_multiplier_{i}" for i in range(EMAX)]+[f"runoff_R_multiplier_{i}" for i in range(N)],
        "state_columns":[f"gauge_pressure_{i}" for i in range(N)]+[f"flow_{i}" for i in range(EMAX)],
        "pressure_datum":{"stored":"p = absolute pressure - Pv","Pv_mmHg":5.,"absolute_pressure":"stored p + 5 mmHg"},
        "initialization":"Exact constant-parameter periodic solution for continuous two-harmonic baseline forcing; cloned across arms. It is not the exact periodic solution for piecewise-linear forecast input interpolation.",
        "node_feature_columns":["C","g","inlet_indicator"],"edge_feature_columns":["R","L"],
        "units":{"pressure":"mmHg","flow":"mL/s","time":"s","C":"mL/mmHg","R":"mmHg s/mL","L":"mmHg s^2/mL"},
        "parameter_ranges":{"Qmean":[60,100],"period":[.75,1.05],"total_C":[.8,1.5],
            "edge_R":[.03,.10],"edge_L":[.002,.010],"systemic_runoff_R":[.7,1.2],
            "runoff_per_outlet":"2 * systemic_R * U(.85,1.15)","C_weights":"U(.75,1.25), renormalized to total_C",
            "inflow_harmonic1_fraction":[.25,.55],"inflow_harmonic2_fraction":[.05,.15]},
        "actions":{"begin_cycles":[.25,1.,1.75,2.5],"ramp_width_cycles":.25,
            "edge_targets":EDGE_TARGETS,"runoff_targets":RUNOFF_TARGETS,
            "selection":"one seeded edge and one distinct preselected runoff site per arm; no outcome-based selection"},
        "reference":{"method":"DOP853","rtol":1e-10,"atol":1e-11,"restart_each_observation_interval":True},
        "structures":structures,"backgrounds":rows,
        "counts":{"structures":38,"seen_structures":24,"heldout_structures":14,"backgrounds":len(rows),
            "episodes":2*len(rows),"train":576,"val":192,"id_test":192,"ood_topology":336},
        "scope":"custom dimensioned engineering networks; not anatomy-derived or patient-calibrated; no clinical efficacy inference"}
    assert len(rows)==648
    write_json(manifest,result)
    write_json(DATA/"design_manifest_sha256.json",{"sha256":sha256(manifest),"created_before_any_trajectory":True})
    return result

def background_parameters(row, structure):
    rng = np.random.default_rng(row["parameter_seed"])
    perm = np.r_[0,rng.permutation(np.arange(1,N))]
    edges = np.asarray(structure["edges"],int)
    ecount = len(edges)
    order = rng.permutation(ecount)
    newedges = perm[edges][order]
    edge_index = np.zeros((2,EMAX), np.int64)
    edge_index[:,:ecount] = newedges.T
    edge_mask = np.arange(EMAX)<ecount
    totalC = rng.uniform(.8,1.5)
    weights = rng.uniform(.75,1.25,N)
    C = totalC*weights/weights.sum()
    R = np.r_[rng.uniform(.03,.10,ecount),np.zeros(EMAX-ecount)]
    L = np.r_[rng.uniform(.002,.010,ecount),np.ones(EMAX-ecount)]
    outlets = perm[np.asarray(structure["outlets"])]
    Rsys = rng.uniform(.7,1.2)
    g = np.zeros(N)
    g[outlets] = 1/(2*Rsys*rng.uniform(.85,1.15,2))
    inlet = np.zeros(N); inlet[0] = 1.
    return {"node_features":np.stack([C,g,inlet],-1),"edge_features":np.stack([R,L],-1),
        "edge_index":edge_index,"edge_mask":edge_mask,"state_mask":np.r_[np.ones(N,bool),edge_mask],
        "T":float(rng.uniform(.75,1.05)),"Qmean":float(rng.uniform(60,100)),
        "harmonics":[float(rng.uniform(.25,.55)),float(rng.uniform(.05,.15))],
        "action_edges":rng.choice(ecount,size=2,replace=False),"action_outlets":outlets,
        "node_permutation":perm,"edge_permutation":order}

def controls_for(p, arm):
    phase = np.arange(STEPS+1)/32.
    omega = 2*np.pi
    ctrl = np.ones((STEPS+1,CONTROL_WIDTH),float)
    ctrl[:,0] = p["Qmean"]*(1+p["harmonics"][0]*np.sin(omega*phase)+p["harmonics"][1]*np.sin(2*omega*phase+.3))
    for column, targets in [(1+int(p["action_edges"][arm]),EDGE_TARGETS[arm]),
                (1+EMAX+int(p["action_outlets"][arm]),RUNOFF_TARGETS[arm])]:
        previous = 1.
        for start, target in zip([.25,1.,1.75,2.5],targets):
            u = np.clip((phase-start)/.25,0,1)
            # Smooth target sample generation; interpolation between these samples
            # is piecewise linear, explicitly shared by all forecasting methods.
            ctrl[:,column] += (target-previous)*u*u*(3-2*u)
            previous = target
    return ctrl

def initial_state(p):
    ctrl = np.ones(CONTROL_WIDTH); ctrl[0] = 1.
    A,b = build_system(p["node_features"],p["edge_features"],p["edge_index"],p["edge_mask"],ctrl)
    x = np.linalg.solve(-A,b*p["Qmean"])
    omega = 2*np.pi/p["T"]
    for harmonic, amplitude, phase in [(1,p["harmonics"][0],0.),(2,p["harmonics"][1],.3)]:
        z = np.linalg.solve(1j*harmonic*omega*np.eye(N+EMAX)-A,b*p["Qmean"]*amplitude)
        x += np.imag(z*np.exp(1j*phase))
    return x

def reference_trajectory(initial, p, controls, dt, method="DOP853", rtol=1e-10, atol=1e-11):
    C,g,inlet = p["node_features"].T
    R,L = p["edge_features"].T
    B = incidence(p["edge_index"],p["edge_mask"])
    y = np.asarray(initial,float).copy()
    out, nfev = [y.copy()], 0
    for c0,c1 in zip(controls[:-1],controls[1:]):
        slope = (c1-c0)/dt
        def rhs(t,x):
            ctrl = c0+slope*t
            pressure,q = x[:N],x[N:]
            dp = (inlet*ctrl[0]-B@q-g*pressure/ctrl[1+EMAX:])/C
            dq = (B.T@pressure-R*ctrl[1:1+EMAX]*q)/L
            return np.r_[dp,dq]
        sol = solve_ivp(rhs,(0,dt),y,method=method,rtol=rtol,atol=atol,t_eval=[dt])
        if not sol.success or not np.all(np.isfinite(sol.y)):
            raise RuntimeError(f"{method} failure: {sol.message}")
        y = sol.y[:,-1]
        out.append(y.copy()); nfev += sol.nfev
    return np.asarray(out), nfev

def one_background(payload):
    row, structure = payload
    p = background_parameters(row,structure)
    x0 = initial_state(p)
    dt = p["T"]/32.
    records = []
    for arm in (0,1):
        ctrl = controls_for(p,arm)
        start = time.perf_counter()
        states,nfev = reference_trajectory(x0,p,ctrl,dt)
        runtime = time.perf_counter()-start
        records.append({**p,"states":states,"controls":ctrl,"dt":dt,
            "background_id":row["background_id"],"graph_id":row["graph_id"],"split":row["split"],"arm":arm,
            "reference_seconds":runtime,"reference_nfev":nfev})
    return records

def pack(records,path):
    records.sort(key=lambda r:(r["background_id"],r["arm"]))
    keys = ["states","controls","node_features","edge_features","edge_index","edge_mask","state_mask",
        "dt","background_id","graph_id","split","arm","T","Qmean","harmonics","reference_seconds","reference_nfev"]
    np.savez_compressed(path,**{key:np.asarray([r[key] for r in records]) for key in keys})

def generate(part="development",workers=4):
    design = freeze_design()
    allowed = {"development":{"train","val"},"test":{"id_test","ood_topology"}}[part]
    if part=="test" and not (ROOT/"analysis"/"model_selection_frozen.json").exists():
        raise RuntimeError("Final test generation requires root-created analysis/model_selection_frozen.json")
    target = DATA/f"{part}.npz"
    if target.exists():
        raise RuntimeError(f"Refuse to overwrite existing outcomes: {target}")
    structures = {s["graph_id"]:s for s in design["structures"]}
    rows = [r for r in design["backgrounds"] if r["split"] in allowed]
    records,failures = [],[]
    started = time.perf_counter()
    with ProcessPoolExecutor(max_workers=workers) as pool:
        jobs = {pool.submit(one_background,(r,structures[r["graph_id"]])):r for r in rows}
        for i,future in enumerate(as_completed(jobs)):
            row = jobs[future]
            try:
                records.extend(future.result())
            except Exception as error:
                failures.append({"background_id":row["background_id"],"error":repr(error)})
            if (i+1)%24==0:
                print(json.dumps({"part":part,"complete_backgrounds":i+1,"total":len(rows),"failures":len(failures)}),flush=True)
    write_json(DATA/f"{part}_generation_log.json",{"part":part,"wall_seconds":time.perf_counter()-started,
        "attempted_backgrounds":len(rows),"saved_episodes":len(records),"failures":failures,
        "generator_sha256":sha256(__file__),"design_sha256":sha256(DATA/"design_manifest.json")})
    if failures:
        raise RuntimeError("Simulation failures recorded; none silently excluded; dataset not finalized.")
    pack(records,target)
    write_json(DATA/f"{part}_sha256.json",{"sha256":sha256(target),"episodes":len(records),
        "backgrounds":len(rows),"state_shape":list(np.asarray([r["states"] for r in records]).shape)})
    print(json.dumps({"written":str(target),"episodes":len(records)}),flush=True)

def main():
    parser = argparse.ArgumentParser()
    parser.add_argument("--part",choices=["freeze","development","test"],default="freeze")
    parser.add_argument("--workers",type=int,default=4)
    args = parser.parse_args()
    if args.part=="freeze":
        print(json.dumps(freeze_design()["counts"]))
    else:
        generate(args.part,args.workers)

if __name__=="__main__":
    main()
```
