## Supplementary Data - Compact audit, summary tables, code, and figure sources for "Panvascular-JEPA: Unseen-Structure Evaluation of Factorized Predictive States for Hemodynamic Networks": graph_models.py

```
"""Shared, permutation-equivariant entity models for the locked graph benchmark.

Node pressure and oriented-edge flow are entity states.  All learnable maps are
shared across entities; graph connectivity is supplied as oriented incidence
aggregation.  There are no graph identifiers or trainable node/edge embeddings.
"""
from __future__ import annotations

import copy
import torch
from torch import nn


def mlp(inp: int, hidden: int, out: int):
    return nn.Sequential(nn.Linear(inp, hidden), nn.SiLU(), nn.Linear(hidden, out))


def aggregate(z, adjacency):
    """Two oriented SUM aggregates; degree is part of the known graph."""
    if z.ndim == 2:
        z = z.unsqueeze(-1)
    return torch.einsum("bcij,bjd->bcid", adjacency, z).permute(0, 2, 1, 3).flatten(2)


class EntityEncoder(nn.Module):
    def __init__(self, latent=16):
        super().__init__()
        # Scalar entity state, two oriented neighboring sums, node/edge flags.
        self.net = mlp(5, 32, latent)

    def forward(self, state, adjacency, entity_type, mask):
        x = torch.cat([state.unsqueeze(-1), aggregate(state, adjacency), entity_type], dim=-1)
        return self.net(x) * mask.unsqueeze(-1)


class GraphModel(nn.Module):
    ARMS = ("mono_jepa", "opf1", "opf4", "supervised_graph")

    def __init__(self, name, static_dim=6, control_dim=7, latent=16):
        super().__init__()
        if name not in self.ARMS:
            raise ValueError(name)
        self.name, self.latent = name, latent
        self.k = 4 if name == "opf4" else 1
        self.encoder = EntityEncoder(latent)
        self.inp = 3 * latent + static_dim + control_dim
        # Target budget: OPF4 with 32 hidden units in each factor head.
        target_predictor = 4 * ((self.inp + 1) * 32 + (32 + 1) * (latent // 4)) + latent ** 2
        if name == "supervised_graph":
            h = min(range(1, 400), key=lambda x: abs((self.inp + 1) * x + x + 1 - target_predictor))
            self.head = mlp(self.inp, h, 1)
            self.head_hidden = h
        else:
            self.target = copy.deepcopy(self.encoder).requires_grad_(False)
            if name == "mono_jepa":
                self.register_buffer("P", torch.eye(latent))
            else:
                q, _ = torch.linalg.qr(torch.randn(latent, latent))
                self.P = nn.Parameter(q)
            pcount = 0 if name == "mono_jepa" else latent ** 2
            h = 32 if self.k == 4 else min(range(1, 400), key=lambda x: abs(
                (self.inp + 1) * x + (x + 1) * latent + pcount - target_predictor))
            self.heads = nn.ModuleList([mlp(self.inp, h, latent // self.k) for _ in range(self.k)])
            self.head_hidden = h

    def context(self, z, adjacency, static, controls):
        return torch.cat([z, aggregate(z, adjacency), static, controls], dim=-1)

    def encode(self, state, adjacency, entity_type, mask, target=False):
        enc = self.target if target and self.name != "supervised_graph" else self.encoder
        return enc(state, adjacency, entity_type, mask)

    def factors(self, z, adjacency, static, controls, mask):
        x = self.context(z, adjacency, static, controls)
        # Residual transition, in the same basis as the target factor coordinates.
        return (z @ self.P + torch.cat([h(x) for h in self.heads], dim=-1)) * mask.unsqueeze(-1)

    def supervised_step(self, state, adjacency, entity_type, mask, static, controls):
        z = self.encode(state, adjacency, entity_type, mask)
        delta = self.head(self.context(z, adjacency, static, controls)).squeeze(-1)
        return (state + delta) * mask

    def orthogonality(self):
        # One complete Gram loss, independent of the number of factor blocks.
        eye = torch.eye(self.latent, device=self.P.device)
        return ((self.P.T @ self.P - eye) ** 2).mean()

    @staticmethod
    def activity(z, mask, entity_type):
        terms = []
        for j in (0, 1):
            vals = z[(mask > 0) & (entity_type[..., j] > 0)]
            terms.append(torch.relu(1.0 - torch.sqrt(vals.var(0, unbiased=False) + 1e-4)).mean())
        return torch.stack(terms).mean()

    @staticmethod
    def equal_type_mse(pred, truth, mask, entity_type):
        terms = []
        err = (pred - truth) ** 2
        if err.ndim == 3:
            err = err.mean(-1)
        for j in (0, 1):
            active = mask * entity_type[..., j]
            terms.append((err * active).sum() / active.sum())
        return torch.stack(terms).mean()

    def objective(self, state, next_state, adjacency, entity_type, mask, static, controls,
                  orth_weight=0.1, target_activity_weight=0.05, online_activity_weight=0.02):
        if self.name == "supervised_graph":
            pred = self.supervised_step(state, adjacency, entity_type, mask, static, controls)
            loss = self.equal_type_mse(pred, next_state, mask, entity_type)
            return loss, {"physical_prediction": loss.detach()}
        z = self.encode(state, adjacency, entity_type, mask)
        with torch.no_grad():
            z_next = self.encode(next_state, adjacency, entity_type, mask, target=True)
        target_factors = z_next @ self.P
        prediction = self.factors(z, adjacency, static, controls, mask)
        pred_loss = self.equal_type_mse(prediction, target_factors, mask, entity_type)
        orth = self.orthogonality()
        target_activity = self.activity(target_factors, mask, entity_type)
        online_activity = self.activity(z, mask, entity_type)
        ow = 0.0 if self.name == "mono_jepa" else orth_weight
        loss = pred_loss + ow * orth + target_activity_weight * target_activity + online_activity_weight * online_activity
        return loss, {"latent_prediction": pred_loss.detach(), "orthogonality": orth.detach(),
                      "target_activity": target_activity.detach(), "online_activity": online_activity.detach()}

    @torch.no_grad()
    def update_target(self, momentum=0.99):
        if self.name != "supervised_graph":
            for target, source in zip(self.target.parameters(), self.encoder.parameters()):
                target.mul_(momentum).add_(source, alpha=1 - momentum)

    def synthesis(self):
        return torch.linalg.pinv(self.P, rtol=1e-5)

    def capacity(self):
        return {"trainable_parameters": sum(p.numel() for p in self.parameters() if p.requires_grad),
                "frozen_ema_parameters": sum(p.numel() for p in self.parameters() if not p.requires_grad),
                "latent_dimension": self.latent, "factor_count": self.k,
                "predictor_hidden_width_per_head": self.head_hidden,
                "readout_parameters": 0 if self.name == "supervised_graph" else 2 * (self.latent + 1)}
```
