## Supplementary Data - Compact audit, summary tables, code, and figure sources for "Panvascular-JEPA: Unseen-Structure Evaluation of Factorized Predictive States for Hemodynamic Networks": manuscript_table_physical.txt

\begin{table}[!tp]
\centering\small
\setlength{\tabcolsep}{4pt}
\renewcommand{\arraystretch}{1.15}
\caption{Physical-coordinate errors on the 14 unseen structures.}
\label{tab:physical}
\begin{tabularx}{\linewidth}{Yrrr}
\toprule
Method & Pressure MAE & Flow MAE & Nonfinite\\
\midrule
Monolithic JEPA & $1.89\times10^{12}$ & $1.52\times10^{13}$ & 0\\
OPF-1 & $1.69\times10^{3}$ & $4.52\times10^{3}$ & 0\\
OPF-4 & $1.65\times10^{10}$ & $8.34\times10^{10}$ & 0\\
Supervised graph & $3.28\times10^{31}$ & $1.03\times10^{32}$ & 0\\
Known RLC, 1 substep & $0.0112$ & $0.0655$ & 0\\
Known RLC, 4 substeps & $6.93\times10^{-4}$ & $0.00414$ & 0\\
Reduced RC & $0.287$ & $1.03$ & 0\\
Fitted RLC & $0.0129$ & $0.0106$ & 0\\
Persistence & $4.75$ & $6.66$ & 0\\
\bottomrule
\end{tabularx}
\par\vspace{4pt}\begin{minipage}{\linewidth}\footnotesize Pressure is in mmHg and flow in mL/s. MAEs use background-then-structure averaging and, for neural methods, the mean of all five seeds. The last column counts nonfinite active scalar predictions across those seeds; a zero count does not imply that a very large finite forecast is stable or useful.\end{minipage}
\end{table}
