## Supplementary Data - Compact audit, summary tables, code, and figure sources for "Panvascular-JEPA: Unseen-Structure Evaluation of Factorized Predictive States for Hemodynamic Networks": mechanistic_baselines.py

```
"""Deterministic mechanism comparisons; preserve per-episode predictions and cost."""
from __future__ import annotations
import os
for key in ("OMP_NUM_THREADS","OPENBLAS_NUM_THREADS","MKL_NUM_THREADS"):
    os.environ.setdefault(key,"1")
import argparse
import json
import time
import numpy as np
from generate_network_data import ROOT, DATA, N, EMAX, incidence, integrate_midpoint, integrate_rc, write_json, sha256

def balance_residuals(states,node_features,edge_features,edge_index,edge_mask,controls,dt):
    """Trapezoidal differential-equation residual, in mL/s and mmHg.

    These residuals have a time-discretization floor even for reference states;
    they are not exact continuous conservation checks or percentage errors.
    """
    C,g,inlet = node_features.T
    R,L = edge_features.T
    B = incidence(edge_index,edge_mask)
    p,q = states[:,:N],states[:,N:]
    mass_rhs = controls[:,0,None]*inlet-(q@B.T)-p*g/controls[:,1+EMAX:]
    momentum_rhs = p@B-q*R*controls[:,1:1+EMAX]
    mass = np.diff(p,axis=0)*C/dt-.5*(mass_rhs[:-1]+mass_rhs[1:])
    momentum = np.diff(q,axis=0)*L/dt-.5*(momentum_rhs[:-1]+momentum_rhs[1:])
    return {"mass_rmse_ml_s":float(np.sqrt(np.mean(mass**2))),
        "momentum_rmse_mmHg":float(np.sqrt(np.mean(momentum[:,edge_mask]**2)))}

def main():
    parser = argparse.ArgumentParser()
    parser.add_argument("--part",choices=["development","test"],default="test")
    args = parser.parse_args()
    with np.load(DATA/f"{args.part}.npz") as handle:
        d = {key:handle[key] for key in handle.files}
    dest = ROOT/"results"
    dest.mkdir(parents=True,exist_ok=True)
    methods = {"rlc_midpoint_1":lambda x,n,e,idx,m,c,dt:integrate_midpoint(x,n,e,idx,m,c,dt,1),
        "rlc_midpoint_4":lambda x,n,e,idx,m,c,dt:integrate_midpoint(x,n,e,idx,m,c,dt,4),
        "rc_reduced_4":lambda x,n,e,idx,m,c,dt:integrate_rc(x,n,e,idx,m,c,dt,4),
        "persistence":lambda x,n,e,idx,m,c,dt:np.repeat(x[None],len(c),axis=0)}
    runtime_rows = []
    for name,fn in methods.items():
        prediction,seconds,mass,momentum = [],[],[],[]
        for i in range(len(d["states"])):
            values = [d[k][i] for k in ("node_features","edge_features","edge_index","edge_mask","controls","dt")]
            start = time.perf_counter()
            y = fn(d["states"][i,0],*values)
            seconds.append(time.perf_counter()-start)
            prediction.append(y)
            residual = balance_residuals(y,*values)
            mass.append(residual["mass_rmse_ml_s"]);momentum.append(residual["momentum_rmse_mmHg"])
        prediction = np.asarray(prediction)
        method_dir = dest/name/"seed_0"
        method_dir.mkdir(parents=True,exist_ok=True)
        for split in np.unique(d["split"]):
            select = d["split"]==split
            np.savez_compressed(method_dir/f"{split}_predictions.npz",
                rollout_physical=prediction[select,1:],truth_physical=d["states"][select,1:],
                seconds=np.asarray(seconds)[select],mass_residual=np.asarray(mass)[select],
                momentum_residual=np.asarray(momentum)[select],state_mask=d["state_mask"][select],
                background_id=d["background_id"][select],graph_id=d["graph_id"][select],
                arm=d["arm"][select],split=d["split"][select])
        runtime_rows.append({"method":name,"episodes":len(seconds),"median_seconds_per_96step_episode":float(np.median(seconds)),"total_seconds":float(np.sum(seconds))})
        print(json.dumps(runtime_rows[-1]),flush=True)
    write_json(dest/f"mechanistic_{args.part}_runtime.json",{"scope":"single CPU process, BLAS thread environment variables fixed at1; no I/O in timer; constructor and linear solve included",
        "methods":runtime_rows,"dataset_sha256":sha256(DATA/f"{args.part}.npz"),"code_sha256":sha256(__file__)})

if __name__=="__main__":
    main()
```
