## Supplementary Data - Compact audit, summary tables, code, and figure sources for "Panvascular-JEPA: Unseen-Structure Evaluation of Factorized Predictive States for Hemodynamic Networks": prepare_manuscript_assets.py

```
"""Preserve the previous manuscript and extract its confirmed declarations."""
from pathlib import Path
import re,json,hashlib
root=Path(__file__).resolve().parent
source=root.parent/'Panvascular-JEPA_submission.tex'
old=source.read_text(encoding='utf-8')
archive=root/'provenance';archive.mkdir(exist_ok=True)
backup=archive/'Panvascular-JEPA_before_independent_experiments.tex'
if not backup.exists():backup.write_bytes(source.read_bytes())
else:old=backup.read_text(encoding='utf-8')
prefix=old[:old.index(r'\begin{abstract}')]
prefix=re.sub(r'\\title\{.*?\}\n\n',lambda m:r'\title{\textbf{Panvascular-JEPA: Unseen-Structure Evaluation of Factorized Predictive States for Hemodynamic Networks}}'+'\n\n',prefix,count=1,flags=re.S)
(root/'manuscript_prefix.txt').write_text(prefix,encoding='utf-8')
declarations=old[old.index(r'\section*{Author contributions}'):old.index(r'\clearpage',old.index(r'\section*{Author contributions}'))]
(root/'manuscript_declarations.txt').write_text(declarations,encoding='utf-8')
items={m.group(1):m.group(0).strip() for m in re.finditer(r'\\bibitem\{([^}]+)\}.*?(?=\n\\bibitem|\n\\end\{thebibliography\})',old,re.S)}
new={
'graphrom':r'\bibitem{graphrom} Pegolotti L, Pfaller MR, Rubio NL, Ding K, Brugarolas Brufau R, Darve E, Marsden AL. Learning Reduced-Order Models for Cardiovascular Simulations with Graph Neural Networks. Preprint, arXiv:2303.07310, 2023. \url{https://arxiv.org/abs/2303.07310}',
'metamodel':r'\bibitem{metamodel} Hanna JM, Varsos P, Kowalski J, Sala L, Meiburg R, Vignon-Clementel IE. A comparative analysis of metamodels for 0D cardiovascular models, and pipeline for sensitivity analysis, parameter estimation, and uncertainty quantification. Preprint, arXiv:2410.12654v2, 2025. \url{https://arxiv.org/abs/2410.12654v2}',
'selection':r'\bibitem{selection} Cawley GC, Talbot NLC. On Over-fitting in Model Selection and Subsequent Selection Bias in Performance Evaluation. \emph{Journal of Machine Learning Research}. 2010;11:2079--2107. \url{https://jmlr.org/papers/v11/cawley10a.html}',
'atlas':r'\bibitem{atlas} NetworkX Developers. Graph Atlas: graph\_atlas\_g. NetworkX documentation and software, version 3.6.1 used in this study. \url{https://networkx.org/documentation/networkx-3.6.1/reference/generated/networkx.generators.atlas.graph_atlas_g.html}.'}
items.update(new)
keys=['rom','svzerod','graphrom','metamodel','ijepa','jepa','vicreg','iabp','selection','atlas','svarchive']
refs='\\begin{thebibliography}{11}\n\\small\n'+'\n'.join(items[k] for k in keys)+'\n\\end{thebibliography}\n'
(root/'manuscript_references.txt').write_text(refs,encoding='utf-8')
(archive/'source_history.json').write_text(json.dumps({'old_source_sha256':hashlib.sha256(backup.read_bytes()).hexdigest(),'old_source_path':str(source),'action':'preserved before new experiment manuscript integration','new_study_directory':str(root),'submission_hold':True},indent=2),encoding='utf-8')
print('Prefix, confirmed declarations, 11 relevant references, and original source preserved.')
```
