## Supplementary Data - Compact audit, summary tables, code, and figure sources for "Panvascular-JEPA: Unseen-Structure Evaluation of Factorized Predictive States for Hemodynamic Networks": README.txt

Panvascular-JEPA independent revision — compact supplementary package

This archive accompanies the revised manuscript uploaded to bioRxiv draft BIORXIV/2026/758273. It contains the audit-ready summary tables, model-selection records, validation protocols, figure sources/metadata, and reproducibility scripts used for the reported analyses.

The complete raw prediction archives generated during the independent network-structure, fair-tuning, and mechanistic-baseline experiments are retained by the authors because their combined size exceeds the practical payload for the bioRxiv submission portal. The manuscript reports this limitation explicitly; the raw archives and checksums can be supplied by the corresponding author and will be deposited in a persistent data repository before public release. The compact tables in this package are the exact files used to populate the manuscript results.
