## Supplementary Data - Compact audit, summary tables, code, and figure sources for "Panvascular-JEPA: Unseen-Structure Evaluation of Factorized Predictive States for Hemodynamic Networks": train_graph_models.py

```
"""Development-only search, followed by locked fresh graph confirmation.

The search command never accepts or opens a held-out confirmation file.  Final
evaluation requires the saved winner file and verifies its development hash.
No checkpoint selection occurs during final confirmation runs.
"""
from __future__ import annotations

import argparse
import hashlib
import json
import os
import platform
import random
import time
from pathlib import Path

os.environ.setdefault("CUBLAS_WORKSPACE_CONFIG", ":4096:8")
import numpy as np
import torch
from graph_models import GraphModel

ROOT = Path(__file__).resolve().parent
ALPHAS = [1e-6, 1e-4, 1e-2, 1., 100.]
CANDIDATES = [{"learning_rate": lr, "weight_decay": wd}
              for lr in [3e-4, 1e-3, 3e-3] for wd in [0., 1e-4]]
FINAL_SEEDS = [211, 223, 227, 229, 233]


def sha(path):
    return hashlib.sha256(Path(path).read_bytes()).hexdigest()


def clean(x):
    if isinstance(x, dict):
        return {str(k): clean(v) for k, v in x.items()}
    if isinstance(x, (list, tuple)):
        return [clean(v) for v in x]
    if isinstance(x, np.ndarray):
        return clean(x.tolist())
    if isinstance(x, np.generic):
        return clean(x.item())
    if isinstance(x, float) and not np.isfinite(x):
        return None
    return x


def save(path, x):
    Path(path).parent.mkdir(parents=True, exist_ok=True)
    Path(path).write_text(json.dumps(clean(x), indent=2, allow_nan=False), encoding="utf-8")


def seed_all(seed):
    random.seed(seed)
    np.random.seed(seed)
    torch.manual_seed(seed)
    torch.cuda.manual_seed_all(seed)
    torch.use_deterministic_algorithms(True)
    torch.backends.cuda.matmul.allow_tf32 = False
    torch.backends.cudnn.allow_tf32 = False


class Data:
    def __init__(self, path, scaler=None):
        self.path = Path(path)
        self.hash = sha(path)
        with np.load(path, allow_pickle=False) as z:
            self.raw = {k: z[k] for k in z.files}
        r = self.raw
        self.split = r["split"].astype(str)
        self.bg = r["background_id"]
        self.graph = r["graph_id"]
        self.nn = r["node_features"].shape[1]
        self.ne = r["edge_features"].shape[1]
        self.nstate = self.nn + self.ne
        self.nt = r["states"].shape[1] - 1
        self.mask = r["state_mask"].astype(np.float32)
        self.etype = np.zeros((len(self.split), self.nstate, 2), np.float32)
        self.etype[:, :self.nn, 0] = 1
        self.etype[:, self.nn:, 1] = 1
        self.etype *= self.mask[..., None]
        assert r["controls"].shape[1:] == (self.nt + 1, 1 + self.ne + self.nn)
        assert r["states"].shape[2] == self.nstate
        assert np.isfinite(r["states"]).all()
        self.adj = np.zeros((len(self.split), 2, self.nstate, self.nstate), np.float32)
        for b in range(len(self.split)):
            for e in range(self.ne):
                if r["edge_mask"][b, e]:
                    src, dst = r["edge_index"][b, :, e].astype(int)
                    self.adj[b, 0, src, self.nn + e] = 1  # outgoing flow
                    self.adj[b, 1, dst, self.nn + e] = 1  # incoming flow
                    self.adj[b, 0, self.nn + e, src] = 1  # source pressure
                    self.adj[b, 1, self.nn + e, dst] = 1  # target pressure
        static, static_masks = self.make_static()
        control, control_masks = self.make_controls()
        if scaler is None:
            train = self.split == "train"
            if not train.any():
                raise ValueError("A new scaler requires training episodes")
            sm, sd = [], []
            for j in range(2):
                valid = np.broadcast_to((self.etype[train, :, j] > 0)[:, None, :], r["states"][train].shape)
                vals = r["states"][train][valid].astype(np.float64)
                sm.append(vals.mean()); sd.append(vals.std())
            gm, gs = [], []
            for j in range(4):
                vals = static[train, :, j][static_masks[train, :, j]].astype(np.float64)
                gm.append(vals.mean()); gs.append(max(vals.std(), 1e-8))
            cm, cs = [], []
            for j in range(7):
                vals = control[train, :, :, j][control_masks[train, :, :, j]].astype(np.float64)
                cm.append(vals.mean()); cs.append(max(vals.std(), 1e-8))
            scaler = {"state_mean": sm, "state_std": sd, "static_mean": gm,
                      "static_std": gs, "control_mean": cm, "control_std": cs}
        self.scaler = scaler
        type_index = (np.arange(self.nstate) >= self.nn).astype(int)
        sm = np.array(scaler["state_mean"])[type_index]
        sd = np.array(scaler["state_std"])[type_index]
        self.s = (((r["states"] - sm) / sd) * self.mask[:, None, :]).astype(np.float32)
        for j in range(4):
            static[..., j] = np.where(static_masks[..., j], (static[..., j] - scaler["static_mean"][j]) / scaler["static_std"][j], 0)
        for j in range(7):
            control[..., j] = np.where(control_masks[..., j], (control[..., j] - scaler["control_mean"][j]) / scaler["control_std"][j], 0)
        self.static, self.ctrl = static.astype(np.float32), control.astype(np.float32)
        self._tensors = None

    def make_static(self):
        r = self.raw
        x = np.zeros((len(self.split), self.nstate, 6), np.float64)
        m = np.zeros_like(x, bool)
        x[:, :self.nn, 0] = np.log(r["node_features"][..., 0]); m[:, :self.nn, 0] = True
        g = r["node_features"][..., 1]
        x[:, :self.nn, 1] = np.log(np.maximum(g, 1e-20)); m[:, :self.nn, 1] = g > 0
        for j in (0, 1):
            x[:, self.nn:, j + 2] = np.log(np.maximum(r["edge_features"][..., j], 1e-20))
            m[:, self.nn:, j + 2] = r["edge_mask"] > 0
        x[:, :self.nn, 4] = g > 0
        x[:, :self.nn, 5] = r["node_features"][..., 2]
        return x, m

    def make_controls(self):
        r = self.raw; c = r["controls"]
        x = np.zeros((len(self.split), self.nt, self.nstate, 7), np.float64)
        m = np.zeros_like(x, bool)
        root = r["node_features"][..., 2]
        for offset in (0, 1):
            tt = slice(offset, self.nt + offset)
            x[:, :, :self.nn, offset] = c[:, tt, 0, None] * root[:, None, :]
            m[:, :, :self.nn, offset] = root[:, None, :] > 0
            x[:, :, self.nn:, 2 + offset] = c[:, tt, 1:1 + self.ne]
            m[:, :, self.nn:, 2 + offset] = r["edge_mask"][:, None, :] > 0
            x[:, :, :self.nn, 4 + offset] = c[:, tt, 1 + self.ne:]
            m[:, :, :self.nn, 4 + offset] = True
        dt = np.asarray(r["dt"]).reshape(-1)
        if dt.size == 1:
            dt = np.repeat(dt, len(self.split))
        x[..., 6] = dt[:, None, None]
        m[..., 6] = self.mask[:, None, :] > 0
        return x, m

    def tensors(self, device):
        if self._tensors is None:
            self._tensors = {k: torch.as_tensor(v, dtype=torch.float32, device=device) for k, v in
                             dict(s=self.s, adj=self.adj, etype=self.etype, mask=self.mask,
                                  static=self.static, ctrl=self.ctrl).items()}
        return self._tensors


def metric(pred, truth, data, rows):
    """Both arms/time first; then equal backgrounds within/equal graph structures."""
    e = (np.asarray(pred, np.float64) - np.asarray(truth, np.float64)) ** 2
    mask = data.mask[rows]
    e = np.where(mask[:, None, :] > 0, e, 0.)
    finite = np.isfinite(e).all()
    # Equal pressure/flow types; inactive padded edge coordinates are excluded.
    pe = e[:, :, :data.nn].mean(2)
    fe = (e[:, :, data.nn:] * mask[:, None, data.nn:]).sum(2) / mask[:, None, data.nn:].sum(2)
    se = .5 * (pe + fe)
    backgrounds = []
    bg = data.bg[rows]; gg = data.graph[rows]
    for b in np.unique(bg):
        ix = bg == b
        backgrounds.append({"background_id": str(b), "graph_id": str(gg[ix][0]),
                            "rmse": float(np.sqrt(se[ix].mean())),
                            "horizon_rmse": np.sqrt(se[ix].mean(0))})
    structures = []
    for g in np.unique(gg):
        bb = [b for b in backgrounds if b["graph_id"] == str(g)]
        structures.append({"graph_id": str(g), "rmse": float(np.mean([b["rmse"] for b in bb])),
                           "horizon_rmse": np.mean([b["horizon_rmse"] for b in bb], axis=0)})
    return {"finite": bool(finite), "nonfinite_count": int((~np.isfinite(e)).sum()),
            "macro_rmse": float(np.mean([g["rmse"] for g in structures])),
            "horizon_rmse": np.mean([g["horizon_rmse"] for g in structures], axis=0),
            "structures": structures, "backgrounds": backgrounds}


@torch.no_grad()
def all_latents(model, data, rows, device, target=True):
    t = data.tensors(device)
    n = len(rows)
    zz = np.empty((n, data.nt + 1, data.nstate, model.latent), np.float32)
    for start in range(0, n, 16):
        rr = rows[start:start + 16]
        b = len(rr); horizon = data.nt + 1
        state = t["s"][rr].reshape(b * horizon, data.nstate)
        other = {k: t[k][rr].repeat_interleave(horizon, 0) for k in ["adj", "etype", "mask"]}
        out = model.encode(state, other["adj"], other["etype"], other["mask"], target=target)
        zz[start:start + b] = out.reshape(b, horizon, data.nstate, model.latent).cpu().numpy()
    return zz


def fit_readout(model, data, device):
    train = np.flatnonzero(data.split == "train"); val = np.flatnonzero(data.split == "val")
    tz = all_latents(model, data, train, device, target=True)
    vz = all_latents(model, data, val, device, target=True)
    weights = np.zeros((2, model.latent + 1), np.float64)
    report = []
    for j in range(2):
        tm = np.broadcast_to((data.etype[train, :, j] > 0)[:, None, :], tz.shape[:3])
        vm = np.broadcast_to((data.etype[val, :, j] > 0)[:, None, :], vz.shape[:3])
        tx = tz[tm].astype(np.float64); ty = data.s[train][tm].astype(np.float64)
        vx = vz[vm].astype(np.float64); vy = data.s[val][vm].astype(np.float64)
        tx = np.column_stack([tx, np.ones(len(tx))]); vx = np.column_stack([vx, np.ones(len(vx))])
        xx, xy = tx.T @ tx, tx.T @ ty
        candidates = []
        for alpha in ALPHAS:
            penalty = np.eye(model.latent + 1) * alpha; penalty[-1, -1] = 0
            w = np.linalg.solve(xx + penalty, xy)
            error = float(np.sqrt(np.mean((vx @ w - vy) ** 2)))
            candidates.append({"alpha": alpha, "validation_reconstruction_rmse": error, "weights": w})
        best = min(candidates, key=lambda x: x["validation_reconstruction_rmse"])
        weights[j] = best["weights"]
        report.append({"entity_type": ["pressure", "flow"][j], "selected_alpha": best["alpha"],
                       "candidates": [{k: v for k, v in c.items() if k != "weights"} for c in candidates]})
    return weights, report


def decode(z, weights, etype):
    return ((z @ weights[:, :-1].T + weights[:, -1]) * etype).sum(-1)


@torch.no_grad()
def diagnostics(model, data, device):
    t = data.tensors(device)
    rows = np.flatnonzero(data.split == "val")
    rr = rows[:min(len(rows), 8)]
    # Simultaneous, nontrivial node and edge relabeling, including padding.
    perm = torch.as_tensor([2, 5, 0, 4, 1, 3, 12, 8, 10, 6, 11, 9, 7], device=device)
    s = t["s"][rr, 0]; a = t["adj"][rr]; ty = t["etype"][rr]; m = t["mask"][rr]
    g = t["static"][rr]; c = t["ctrl"][rr, 0]
    pa = a[:, :, perm][:, :, :, perm]
    z = model.encode(s, a, ty, m)
    pz = model.encode(s[:, perm], pa, ty[:, perm], m[:, perm])
    encoder_error = float((pz - z[:, perm]).abs().max().cpu())
    if model.name == "supervised_graph":
        y = model.supervised_step(s, a, ty, m, g, c)
        py = model.supervised_step(s[:, perm], pa, ty[:, perm], m[:, perm], g[:, perm], c[:, perm])
    else:
        y = model.factors(z, a, g, c, m)
        py = model.factors(pz, pa, g[:, perm], c[:, perm], m[:, perm])
    prediction_error = float((py - y[:, perm]).abs().max().cpu())
    report = {"development_only": True, "permutation_encoder_max_abs": encoder_error,
              "permutation_prediction_max_abs": prediction_error,
              "permutation_tolerance": 1e-4}
    if max(encoder_error, prediction_error) > 1e-4:
        raise AssertionError("Entity permutation equivariance test failed")
    if model.name != "supervised_graph":
        p = model.P.detach().cpu().numpy().astype(np.float64)
        sv = np.linalg.svd(p, compute_uv=False)
        report["basis"] = {"singular_values": sv, "condition": float(sv.max() / max(sv.min(), 1e-30)),
                           "rank_at_rtol_1e_5": int((sv > sv.max() * 1e-5).sum()),
                           "normalized_full_gram_loss": float(np.mean((p.T @ p - np.eye(model.latent)) ** 2))}
        zz = all_latents(model, data, rows, device, target=True)
        for j, name in enumerate(["pressure", "flow"]):
            valid = np.broadcast_to((data.etype[rows, :, j] > 0)[:, None, :], zz.shape[:3])
            vals = zz[valid].astype(np.float64)
            sing = np.linalg.svd(vals - vals.mean(0), compute_uv=False)
            probs = sing / max(sing.sum(), 1e-30)
            report[name + "_ema_latents"] = {"coordinate_std": vals.std(0),
                "effective_rank": float(np.exp(-np.sum(probs[probs > 0] * np.log(probs[probs > 0]))))}
    return report


@torch.no_grad()
def evaluate(model, weights, data, split, device, save_path=None):
    rows = np.flatnonzero(data.split == split)
    if not len(rows):
        return None
    t = data.tensors(device)
    g = {k: t[k][rows] for k in ["adj", "etype", "mask", "static", "ctrl", "s"]}
    wt = None if weights is None else torch.as_tensor(weights, dtype=torch.float32, device=device)
    shape = (len(rows), data.nt, data.nstate)
    pure = np.empty(shape, np.float32); one = np.empty(shape, np.float32)
    reencode = np.empty(shape, np.float32); oracle = np.empty(shape, np.float32)
    is_sup = model.name == "supervised_graph"
    if device == "cuda": torch.cuda.synchronize()
    start = time.perf_counter()
    if not is_sup:
        inv = model.synthesis()
        z = model.encode(g["s"][:, 0], g["adj"], g["etype"], g["mask"])
    phys = g["s"][:, 0]
    for step in range(data.nt):
        cc = g["ctrl"][:, step]
        if is_sup:
            phys = model.supervised_step(phys, g["adj"], g["etype"], g["mask"], g["static"], cc)
            pure[:, step] = phys.cpu().numpy()
        else:
            z = model.factors(z, g["adj"], g["static"], cc, g["mask"]) @ inv
            pure[:, step] = (decode(z, wt, g["etype"]) * g["mask"]).cpu().numpy()
    if device == "cuda": torch.cuda.synchronize()
    elapsed = time.perf_counter() - start
    # One-step, decoder reconstruction and optional decode/re-encode mechanism diagnostics.
    phys = g["s"][:, 0]
    for step in range(data.nt):
        cc = g["ctrl"][:, step]
        if is_sup:
            pred = model.supervised_step(g["s"][:, step], g["adj"], g["etype"], g["mask"], g["static"], cc)
            one[:, step] = pred.cpu().numpy()
            reencode[:, step] = pure[:, step]
            oracle[:, step] = g["s"][:, step + 1].cpu().numpy()
        else:
            teacher = model.encode(g["s"][:, step], g["adj"], g["etype"], g["mask"])
            zp = model.factors(teacher, g["adj"], g["static"], cc, g["mask"]) @ inv
            one[:, step] = (decode(zp, wt, g["etype"]) * g["mask"]).cpu().numpy()
            zo = model.encode(g["s"][:, step + 1], g["adj"], g["etype"], g["mask"], target=True)
            oracle[:, step] = (decode(zo, wt, g["etype"]) * g["mask"]).cpu().numpy()
            zr = model.encode(phys, g["adj"], g["etype"], g["mask"])
            zr = model.factors(zr, g["adj"], g["static"], cc, g["mask"]) @ inv
            phys = decode(zr, wt, g["etype"]) * g["mask"]
            reencode[:, step] = phys.cpu().numpy()
    truth = data.s[rows, 1:]
    result = {"split": split, "episodes": len(rows), "pure_latent_rollout": metric(pure, truth, data, rows),
              "one_step": metric(one, truth, data, rows),
              "truth_encoding_readout": metric(oracle, truth, data, rows),
              "decode_reencode_rollout": metric(reencode, truth, data, rows),
              "timing": {"device": device, "total_rollout_seconds": elapsed,
                         "seconds_per_trajectory": elapsed / len(rows), "batch_episodes": len(rows),
                         "scope": "batched rollout including per-step device-to-CPU prediction transfer; excludes preprocessing, readout fitting, and diagnostic passes"}}
    if save_path is not None:
        type_index = (np.arange(data.nstate) >= data.nn).astype(int)
        sm = np.array(data.scaler["state_mean"])[type_index]
        sd = np.array(data.scaler["state_std"])[type_index]
        def physical(arr):
            return ((arr.astype(np.float64) * sd + sm) * data.mask[rows, None, :]).astype(np.float64)
        np.savez_compressed(save_path, predicted=pure, one_step=one, reconstruction=oracle,
                            decode_reencode=reencode, truth=truth, rows=rows,
                            rollout_physical=physical(pure), truth_physical=data.raw["states"][rows, 1:],
                            one_step_physical=physical(one), reconstruction_physical=physical(oracle),
                            decode_reencode_physical=physical(reencode),
                            background_id=data.bg[rows], graph_id=data.graph[rows],
                            arm=data.raw["arm"][rows], split=data.split[rows],
                            state_mask=data.mask[rows], mask=data.mask[rows],
                            state_mean=np.array(data.scaler["state_mean"]),
                            state_std=np.array(data.scaler["state_std"]))
    return result


def train(name, candidate, seed, steps, data, output, device, batch=128):
    output = Path(output); output.mkdir(parents=True, exist_ok=True)
    seed_all(seed)
    model = GraphModel(name).to(device)
    opt = torch.optim.AdamW(model.parameters(), lr=candidate["learning_rate"], weight_decay=candidate["weight_decay"])
    t = data.tensors(device)
    train_rows = torch.as_tensor(np.flatnonzero(data.split == "train"), device=device)
    curve = []; fail = None; completed = 0
    if device == "cuda": torch.cuda.synchronize(); torch.cuda.reset_peak_memory_stats()
    start = time.perf_counter()
    for step in range(steps):
        rr = train_rows[torch.randint(len(train_rows), (batch,), device=device)]
        tt = torch.randint(data.nt, (batch,), device=device)
        opt.zero_grad(set_to_none=True)
        loss, parts = model.objective(t["s"][rr, tt], t["s"][rr, tt + 1], t["adj"][rr],
                                      t["etype"][rr], t["mask"][rr], t["static"][rr], t["ctrl"][rr, tt])
        if not torch.isfinite(loss):
            fail = f"Nonfinite training loss at update {step + 1}"
            break
        loss.backward()
        norm = torch.nn.utils.clip_grad_norm_(model.parameters(), 5.0)
        if not torch.isfinite(norm):
            fail = f"Nonfinite gradient norm at update {step + 1}"
            break
        opt.step(); model.update_target(); completed += 1
        if (step + 1) % 500 == 0 or step == 0 or step + 1 == steps:
            curve.append({"update": step + 1, "loss": float(loss.detach().cpu()),
                          "gradient_norm_pre_clip": float(norm.detach().cpu()),
                          **{k: float(v.cpu()) for k, v in parts.items()}})
    if device == "cuda": torch.cuda.synchronize()
    elapsed = time.perf_counter() - start
    report = {"name": name, "candidate": candidate, "seed": seed, "planned_updates": steps,
              "completed_updates": completed, "capacity": model.capacity(), "training_seconds": elapsed,
              "training_curve": curve, "failure": fail, "data_sha256": data.hash,
              "peak_cuda_memory_bytes": torch.cuda.max_memory_allocated() if device == "cuda" else 0,
              "batch_size": batch, "gradient_clip_norm": 5., "ema_momentum": .99,
              "orthogonality_weight": .1, "target_activity_weight": .05, "online_activity_weight": .02}
    weights = None
    if fail is None and name != "supervised_graph":
        try:
            weights, report["readout"] = fit_readout(model, data, device)
        except np.linalg.LinAlgError as exc:
            fail = report["failure"] = "Readout linear algebra failed: " + str(exc)
    if fail is None:
        report["diagnostics"] = diagnostics(model, data, device)
    torch.save({"state_dict": model.state_dict(), "name": name, "readout": weights}, output / "model.pt")
    save(output / "training.json", report)
    save(output / "scalers.json", data.scaler)
    return model, weights, report


def main():
    p = argparse.ArgumentParser()
    p.add_argument("mode", choices=["smoke", "search", "confirm"])
    p.add_argument("--development", required=True)
    p.add_argument("--confirmation")
    p.add_argument("--output", required=True)
    p.add_argument("--selection")
    p.add_argument("--device", default="cuda", choices=["cuda", "cpu"])
    p.add_argument("--arm", choices=GraphModel.ARMS)
    p.add_argument("--seed", type=int)
    args = p.parse_args()
    torch.set_num_threads(2); torch.set_num_interop_threads(2)
    if args.device == "cuda": torch.cuda.set_per_process_memory_fraction(.055)
    data = Data(args.development)
    if set(data.split) - {"train", "val"}:
        raise ValueError("Development file must contain training/validation only")
    out = Path(args.output); out.mkdir(parents=True, exist_ok=True)
    env = {"python": platform.python_version(), "torch": torch.__version__, "numpy": np.__version__,
           "device": args.device, "gpu": torch.cuda.get_device_name() if args.device == "cuda" else None,
           "cuda_memory_fraction": .055, "cpu_threads": 2, "deterministic": True,
           "source_hashes": {n: sha(ROOT / n) for n in ["graph_models.py", "train_graph_models.py"]}}
    save(out / "environment.json", env)
    arms = [args.arm] if args.arm else list(GraphModel.ARMS)
    if args.mode == "smoke":
        reports = {}
        for name in arms:
            model, weights, report = train(name, CANDIDATES[1], 101, 3, data, out / name, args.device, batch=32)
            reports[name] = {"training": report, "validation": evaluate(model, weights, data, "val", args.device)}
            del model
        save(out / "smoke.json", reports)
    elif args.mode == "search":
        if args.confirmation:
            raise ValueError("Search is not permitted to accept confirmation data")
        selection = {"development_sha256": data.hash, "selection_seed": 101,
                     "selection_metric": "96-step pure-latent rollout equal-structure macro-RMSE",
                     "updates_per_candidate": 1000, "candidates_per_arm": 6,
                     "source_hashes": env["source_hashes"], "arms": {}}
        for name in arms:
            results = []
            for ci, candidate in enumerate(CANDIDATES):
                od = out / name / f"candidate_{ci}"
                model, weights, report = train(name, candidate, 101, 1000, data, od, args.device)
                val = None if report["failure"] else evaluate(model, weights, data, "val", args.device, od / "validation_predictions.npz")
                save(od / "validation.json", val)
                score = np.inf if val is None else val["pure_latent_rollout"]["macro_rmse"]
                if not np.isfinite(score): score = np.inf
                results.append({"candidate_index": ci, **candidate, "validation_macro_rmse": float(score),
                                "finite": bool(np.isfinite(score)), "training_failure": report["failure"]})
                print(json.dumps({"arm": name, "candidate": ci, "score": clean(float(score))}), flush=True)
                del model
            finite = [x for x in results if x["finite"]]
            if not finite:
                raise RuntimeError(f"All prespecified candidates failed for {name}; do not tune on heldout")
            winner = min(finite, key=lambda x: x["validation_macro_rmse"])
            selection["arms"][name] = {"all_candidates": results, "winner": winner}
            save(out / "selection_in_progress.json", selection)
        save(out / "selected_hyperparameters.json", selection)
        (out / "selected_hyperparameters.sha256").write_text(sha(out / "selected_hyperparameters.json"), encoding="utf-8")
    else:
        if not args.selection or not args.confirmation:
            raise ValueError("Confirmation requires a frozen development selection and separate confirmation NPZ")
        selected = json.loads(Path(args.selection).read_text(encoding="utf-8"))
        if selected["development_sha256"] != data.hash:
            raise ValueError("Development hash changed after selection")
        if selected["source_hashes"] != env["source_hashes"]:
            raise ValueError("Model/training code changed after selection; audited amendment needed")
        confirm = Data(args.confirmation, scaler=data.scaler)
        if set(confirm.split) & {"train", "val"}:
            raise ValueError("Confirmation file contains development labels")
        if set(data.bg) & set(confirm.bg):
            raise ValueError("Development/confirmation background overlap")
        seeds = [args.seed] if args.seed else FINAL_SEEDS
        for name in arms:
            candidate = {k: selected["arms"][name]["winner"][k] for k in ["learning_rate", "weight_decay"]}
            for seed in seeds:
                od = out / name / f"seed_{seed}"
                model, weights, report = train(name, candidate, seed, 2000, data, od, args.device)
                results = {"selection_sha256": sha(args.selection), "confirmation_sha256": confirm.hash,
                           "training": report, "splits": {}}
                if report["failure"] is None:
                    for split in sorted(set(confirm.split)):
                        results["splits"][split] = evaluate(model, weights, confirm, split, args.device, od / f"{split}_predictions.npz")
                save(od / "metrics.json", results)
                print(json.dumps({"arm": name, "seed": seed, "scores": {k: clean(v["pure_latent_rollout"]["macro_rmse"]) for k, v in results["splits"].items()}}), flush=True)
                del model


if __name__ == "__main__":
    main()
```
