## Supplementary Data - Compact audit, summary tables, code, and figure sources for "Panvascular-JEPA: Unseen-Structure Evaluation of Factorized Predictive States for Hemodynamic Networks": validate_network_physics.py

```
"""Independent numerical fixtures on training-design backgrounds only."""
from __future__ import annotations
import os
for key in ("OMP_NUM_THREADS","OPENBLAS_NUM_THREADS","MKL_NUM_THREADS"):
    os.environ.setdefault(key,"1")
import json
from pathlib import Path
import numpy as np
import networkx as nx
from scipy.integrate import solve_ivp
from generate_network_data import (ROOT, DATA, N, EMAX, CONTROL_WIDTH, freeze_design,
    background_parameters, initial_state, controls_for, reference_trajectory,
    build_system, integrate_midpoint, incidence, write_json, sha256)

def main():
    design = freeze_design()
    structs = {s["graph_id"]:s for s in design["structures"]}
    candidates = [r for r in design["backgrounds"] if r["split"]=="train" and r["replicate"]==0]
    # Fixed fixture order, independent of any model or outcome.
    selected = [next(r for r in candidates if structs[r["graph_id"]]["num_edges"]==e) for e in (5,6,7)]
    selected.extend([r for r in candidates if r not in selected][:2])
    fixtures = []
    for row in selected:
        p = background_parameters(row,structs[row["graph_id"]])
        ctrl = np.ones(CONTROL_WIDTH); ctrl[0]=p["Qmean"]
        A,b = build_system(p["node_features"],p["edge_features"],p["edge_index"],p["edge_mask"],ctrl)
        steady = np.linalg.solve(-A,b)
        steady_resid = float(np.max(np.abs(A@steady+b)))
        rng = np.random.default_rng(915+row["background_id"])
        state = np.r_[rng.uniform(20,100,N),rng.normal(0,30,EMAX)*p["edge_mask"]]
        dx = A@state+b
        C,g,inlet = p["node_features"].T
        R,L = p["edge_features"].T
        energy_dot = float(np.dot(C*state[:N],dx[:N])+np.dot(L*state[N:],dx[N:]))
        rhs_energy = float(state[:N]@inlet*ctrl[0]-np.dot(g,state[:N]**2)-np.dot(R,state[N:]**2))
        energy_error = abs(energy_dot-rhs_energy)/max(1.,abs(rhs_energy))
        # Independent closed-form complex sinusoidal solution under fixed R.
        unit = ctrl.copy();unit[0]=1.
        A,unitb = build_system(p["node_features"],p["edge_features"],p["edge_index"],p["edge_mask"],unit)
        omega = 2*np.pi/p["T"]
        dc = np.linalg.solve(-A,unitb*p["Qmean"])
        modes = [np.linalg.solve(1j*k*omega*np.eye(N+EMAX)-A,unitb*p["Qmean"]*amp)
                 for k,amp in enumerate(p["harmonics"],1)]
        t = np.linspace(0,3*p["T"],97)
        exact = np.asarray([dc+np.imag(modes[0]*np.exp(1j*omega*s))+np.imag(modes[1]*np.exp(1j*(2*omega*s+.3))) for s in t])
        def analytic_rhs(s,x):
            q = p["Qmean"]*(1+p["harmonics"][0]*np.sin(omega*s)+p["harmonics"][1]*np.sin(2*omega*s+.3))
            return A@x+unitb*q
        numerical = solve_ivp(analytic_rhs,(0,t[-1]),exact[0],method="DOP853",rtol=1e-11,atol=1e-12,t_eval=t).y.T
        analytic_error = float(np.max(np.abs(numerical-exact)))
        x0 = initial_state(p)
        controls = controls_for(p,0)
        dop,_ = reference_trajectory(x0,p,controls,p["T"]/32.)
        radau,_ = reference_trajectory(x0,p,controls,p["T"]/32.,method="Radau",rtol=1e-10,atol=1e-11)
        convergence = []
        for subdivisions in (1,2,4,8):
            y = integrate_midpoint(x0,p["node_features"],p["edge_features"],p["edge_index"],p["edge_mask"],controls,p["T"]/32.,subdivisions)
            convergence.append({"substeps":subdivisions,"state_rmse_physical_mixed_units":float(np.sqrt(np.mean((y-dop)[:,p["state_mask"]]**2)))})
        fixtures.append({"background_id":row["background_id"],"graph_id":row["graph_id"],
            "max_steady_derivative":steady_resid,"passivity_relative_residual":energy_error,
            "continuous_sinusoidal_max_abs_error":analytic_error,
            "DOP853_vs_Radau_max_abs_error":float(np.max(np.abs(dop-radau))),
            "midpoint_convergence":convergence})
    graphs = [nx.from_edgelist(s["edges"]) for s in design["structures"]]
    iso_pairs = [(i,j) for i in range(len(graphs)) for j in range(i) if nx.is_isomorphic(graphs[i],graphs[j])]
    assert not iso_pairs
    assert max(f["passivity_relative_residual"] for f in fixtures)<1e-12
    assert max(f["max_steady_derivative"] for f in fixtures)<1e-8
    assert max(f["continuous_sinusoidal_max_abs_error"] for f in fixtures)<1e-6
    assert max(f["DOP853_vs_Radau_max_abs_error"] for f in fixtures)<1e-6
    assert all(all(f["midpoint_convergence"][i+1]["state_rmse_physical_mixed_units"]<f["midpoint_convergence"][i]["state_rmse_physical_mixed_units"] for i in range(3)) for f in fixtures)
    out = {"scope":"fixed training-background fixtures; no heldout states generated or examined",
        "passed":True,"nonisomorphic_graph_count":len(graphs),"isomorphic_pairs":iso_pairs,
        "fixtures":fixtures,"generator_sha256":sha256(ROOT/"generate_network_data.py"),
        "validation_code_sha256":sha256(__file__)}
    write_json(ROOT/"physics_validation.json",out)
    print(json.dumps(out,indent=2))

if __name__=="__main__":
    main()
```
