## Supplementary figures and images for "Panvascular-JEPA: Unseen-Structure Evaluation of Factorized Predictive States for Hemodynamic Networks"

### catalogue_geometry_preview.png

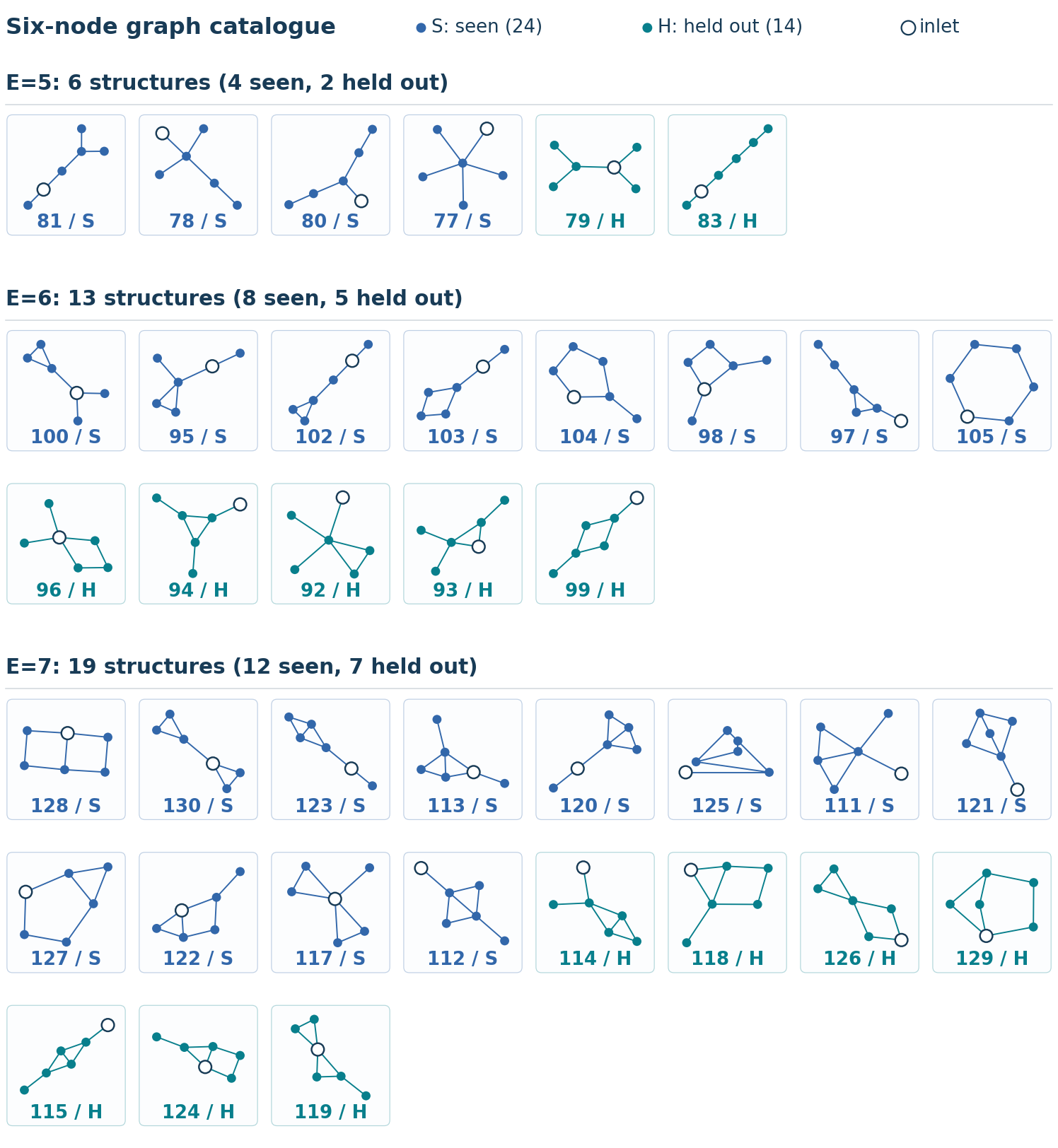
